# CloneSig can jointly infer intra-tumor heterogeneity and mutational signature activity in bulk tumor sequencing data

**DOI:** 10.1101/825778

**Authors:** Judith Abécassis, Fabien Reyal, Jean-Philippe Vert

## Abstract

Systematic DNA sequencing of cancer samples has highlighted the importance of two aspects of cancer genomics: intra-tumor heterogeneity (ITH) and mutational processes. These two aspects may not always be independent, as different mutational processes could be involved in different stages or regions of the tumor, but existing computational approaches to study them largely ignore this potential dependency. Here, we present CloneSig, a computational method to jointly infer ITH and mutational processes in a tumor from bulk-sequencing data. Extensive simulations show that CloneSig outperforms current methods for ITH inference and detection of mutational processes when the distribution of mutational signatures changes between clones. Applied to a large cohort of 8,951 tumors with whole-exome sequencing data from The Cancer Genome Atlas, and on a pan-cancer dataset of 2,632 whole-genome sequencing tumor samples from the Pan-Cancer Analysis of Whole Genomes initiative, CloneSig obtains results overall coherent with previous studies.

## 1 Introduction

The advent and recent democratization of high-throughput sequencing technologies has triggered much effort recently to identify the genomic forces that shape tumorigenesis and cancer progression. In particular, they have begun to shed light on evolutionary principles happening during cancer progression, and responsible for intra-tumor heterogeneity (ITH). Indeed, as proposed by Nowell in the 1970s, cancer cells progressively accumulate somatic mutations during tumorigenesis and the progression of the disease, following similar evolutionary principles as any biological population able to acquire heritable transformations [1]. As new mutations appear in a tumor, either because they bring a selective advantage or simply through neutral evolution, some cancer cells may undergo clonal expansion until they represent the totality of the tumor or a substantial part of it. This may result in a tumor composed of a mosaic of cell subpopulations with specific mutations. Better understanding these processes can provide valuable insights with implications in cancer detection and monitoring, patient stratification and therapeutic strategy [2, 3, 4, 5].

Bulk genome sequencing of a tumor sample allows us in particular to capture two important aspects of ITH. First, by providing an estimate of the proportion of cells harboring each single nucleotide variant (SNV), genome sequencing allows us to assess ITH in terms of presence and proportions of subclonal populations and, to some extent, to reconstruct the evolutionary history of the tumor [6, 7, 8, 9]. This estimation is challenging, both because a unique tumor sample may miss the full extent of the true tumor heterogeneity, and because the computational problem of deconvoluting a bulk sample into subclones is notoriously difficult due to noise and lack of identifiability [6, 10]. Second, beyond their frequency in the tumor, SNVs also record traces of the mutational processes active at the time of their occurrence through biases in the sequence patterns at which they arise, as characterized with the concept of mutational signature [11]. A mutational signature is a probability distribution over possible mutation types, defined by the nature of the substitution and its trinucleotide sequence context, and reflects exogenous or endogenous causes of mutations. Forty-nine such signatures have been outlined [12], and are referenced in the COSMIC database, with known or unknown aetiologies. They are sometimes denoted as SBS for single-based substitutions signatures, in opposition to signatures of other genomic alterations. Deciphering signature activities in a tumor sample, and their changes over time, can provide valuable insights about the causes of cancer, the dynamic of tumor evolution and driver events, and finally help us better estimate the patient prognosis and optimize the treatment strategy [2, 5]. A few computational methods have been proposed to estimate the activity of different predefined signatures in a tumor sample from bulk genome sequencing [12, 13], also known as the signature refitting problem; we refer the reader to the reviewing work of [14, 15] for a more formal overview of existing methods.

These two aspects of genome alterations during tumor development may not always be independent from each other. For example, if a mutation triggers subclonal expansion because it activates a particular mutational process, then new mutations in the corresponding subclone may carry the mark of this process, as observed for APOBEC mutations in human bladder [16]. Alternatively, mutational processes may change over tumor development due to varying exposures to mutagenes, or between cells of the same tumor due to different micro-environments and hence selective pressures. This may lead to additional changes in mutational signatures of SNVs not necessarily coinciding with clones, but possibly resulting in different signature activities between different subclones. Consequently, taking into account mutation types in addition to SNV frequencies may benefit ITH methods, although the extent of this dependency in human cancers is still unknown. Furthermore, identifying mutational processes specific to distinct subclones, and in particular detecting changes in mutational processes during cancer progression, may be of clinical interest since prognosis and treatment options may differ in that case. However, current computational pipelines for ITH and mutational process analysis largely ignore the potential dependency between these two aspects, and typically treat them independently from each other or sequentially. In the sequential approach, as for example implemented in Palimpsest [17], subclones are first identified by an ITH analysis, and in a second step mutational signatures active in each subclone are investigated. In such a sequential analysis, however, we can not observe changes in mutational signature composition if the initial clonal decomposition step fails to detect correct subclones, and we ignore information regarding mutational signatures during ITH inference. Recently, TrackSig [18] was proposed to combine these two steps by performing an evolution-aware signature activity deconvolution, in order to better detect changes in signature activity along tumor evolution. However, while TrackSig overcomes the need to rely on a previously computed subclonal reconstruction, it does not leverage the possible association between mutation frequency and mutation type to jointly infer ITH and mutation processes active in the tumor. Furthermore, by design TrackSig can only work if a sufficient number of SNV is available, limiting currently its use to whole-genome sequencing (WGS) data. This is an important limitation given the popularity of whole-exome sequencing (WES) to characterize tumors, particularly in the clinical setting. An extension has been proposed to better account for SNV frequencies in the change point detection, TrackSigFreq [19], but is still limited to WGS samples.

In this work, we propose CloneSig, a method that leverages both the frequency and the mutation type of SNVs to jointly perform ITH reconstruction and decipher the activity of mutational signatures in each subclone. By exploiting the possible association between subclones and mutational processes to increase its statistical power, we show that CloneSig performs accurate estimations with fewer SNVs than competing methods, and in particular that it can be used with WES data. We show through extensive simulations, and three independent simulated gold-standard datasets [20, 21, 2, 22] that CloneSig reaches state-of-the-art performance in subclonal reconstruction and signature activity deconvolution from WGS and WES data, in the presence or absence of signature activity variations between clones. We then provide a detailed analysis of 8,951 pancancer WES samples from the Cancer Genome Atlas (TCGA) with CloneSig, and on 2,632 WGS samples from the International Cancer Genome Consortium’s Pan-Cancer Analysis of Whole Genomes (PCAWG) cohort [23], where we recover results coherent with published results on WGS [23, 25, 2] as well as different findings of potential clinical relevance. CloneSig is available as a Python package at https://github.com/judithabk6/clonesig [75].

## 2 Results

### 2.1 Joint estimation of ITH and mutational processes with CloneSig

We propose CloneSig, a method to jointly infer ITH and estimate mutational processes active in different clones from bulk genome sequencing data of a tumor sample. The rationale behind CloneSig is illustrated in Figure 1, which shows a scatter-plot of all SNVs detected by WES in a sarcoma (TCGA patient TCGA-3B-A9HI) along two axes: horizontally, the mutation type of the SNV, and vertically, its cancer cell fraction (CCF) estimated from WES read counts. Following previous work on mutational processes [11, 26], we consider 96 possible mutation types, defined by the nature of the substitution involved and the two flanking nucleotides.

**Figure 1:**
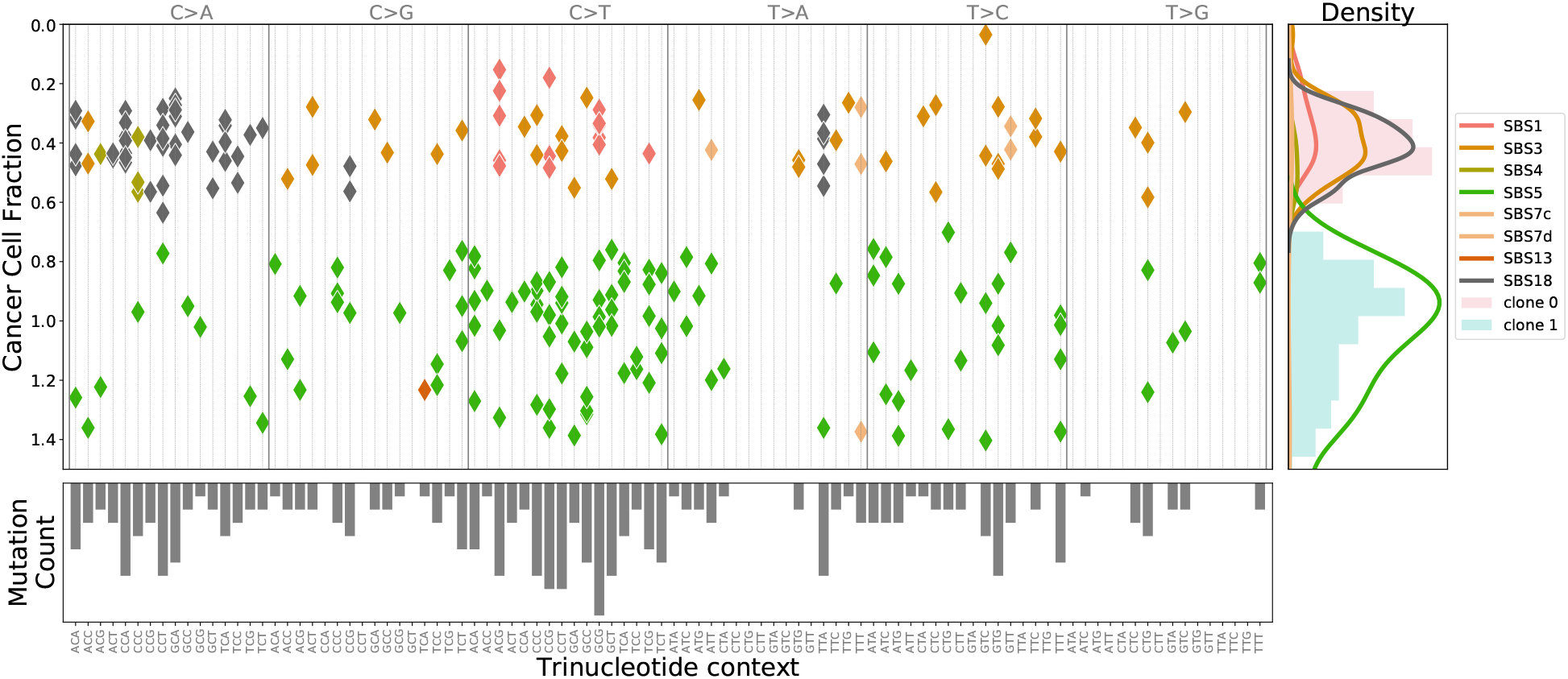
CloneSig analysis of 246 SNVs obtained by WES of a sarcoma sample (patient TCGA-3B-A9HI). The main panel displays all SNVs in 2 dimensions: horizontally the mutation type, which describes the type of substitution together with the flanking nucleotides, and vertically the estimated CCF, as corrected by CloneSig with the estimated mutation multiplicity. From these data CloneSig infers the presence of 2 clones and a number of mutational signatures active in the different clones. Each mutation in the main panel is colored according to the most likely mutational signature according to CloneSig. On the right panel, the CCF histogram is represented and colored with estimated clones, and superimposed with mutational signature density. The bottom panel represents the total mutation type profile. The changing pattern of mutation types with CCF is clearly visible, illustrating the opportunity for CloneSig to perform joint estimation of ITH and signature activity, while most methods so far explore separately those data, considering solely the CCF histogram in the right panel for ITH analysis, or the mutation profile of the bottom panel to infer mutational processes.

Standard methods for ITH assessment and clonal deconvolution only exploit the distribution of CCF values in the sample, as captured by the histogram on the right panel of Figure 1, while standard methods for mutational signature analysis only exploit the mutation profiles capturing the distribution of mutation contexts, as represented by the histogram on the bottom panel. However, we clearly see in the scatter-plot that these two parameters are not independent, e.g., C*>*A mutations tend to occur frequently at low CCF, while C*>*T mutations occur more frequently at high CCF. CloneSig exploits this association by working directly at the 2D scatter-plot level, in order to jointly infer subclones and mutational processes involved in those subclones. Intuitively, working at this level increases the statistical power of subclone detection when subclones are better separated in the 2D scatter-plot than on each horizontal or vertical axis, i.e., when the activity of mutational processes varies between subclones. Additional examples are shown in Supplementary Figures 103 to 108.

Note that while the association between the frequency and the type of a mutation may be the consequence of a co-segregation of mutational processes and clones, which itself could occur if a change in a mutational process coincides with a change in the fitness of a cell and subsequently in the clonal composition of the tumour, CloneSig’s model does not rely on such a strong hypothesis. In particular, an association may exist even if changes in mutational processes do not coincide with clonal evolution, and even if each clone contains heterogeneous populations of cells expressing different mutational processes. What CloneSig assumes is merely that, on average, the proportion of mutations associated to each mutational process may differ between clones. Importantly, if this assumption does not hold, then CloneSig can still be used and behave like a standard method for ITH inference, while we expect the performance of CloneSig to improve in cases where the assumption holds.

More precisely, CloneSig is based on a probabilistic graphical model [27], summarized graphically in Box 1, to model the distribution of allelic counts and trinucleotidic contexts of SNVs in a tumor. These observed variables are statistically associated through shared unobserved latent factors, including the number of clones in the tumor, the CCF of each clone, and the mutational processes active in each clone. CloneSig infers these latent factors for each tumor from the set of SNVs by maximum likelihood estimation, using standard machinery of probabilistic graphical models. Once the parameters of the model are inferred for a given tumor, we can read from them the estimated number of subclones together with their CCF, as well as the set of mutational processes active in each clone along with their strength. In addition, for each individual SNV, CloneSig allows us to estimate the clone and the signature that generated it, in a fully probabilistic manner; for example, in Figure 1, each SNV in the scatter-plot is colored according to the most likely mutational signature that generated it, according to CloneSig. Finally, we developed a likelihood ratio-based statistical test to assess whether mutational signatures significantly differ between subclones, in order to help characterize the evolutionary process involved in the life of the tumor. We refer the reader to the Methods section and Box 1 for all technical details regarding CloneSig.

### 2.2 Performance for subclonal reconstruction

We first assess the ability of CloneSig to correctly reconstruct the subclonal organization of a tumor on simulated data, using four different simulators: 1) the DREAM challenge dataset devised in [20, 21], consisting of 5 simulated WGS tumors with different sequencing depths; 2) PhylogicSim500, comprising 500 samples generated using the PhylogicNDT method, and 3) SimClone1000, comprising 972 samples generated with the simulator SimClone, both proposed by [2, 22]; and 4) CloneSigSim, a simulator we propose which simulates data according to the probabilistic graphical model behind CloneSig. The different simulators differ in the underlying simulation models, as well as other features such as the number of mutations simulated (DREAM, PhylogicSim500 and SimClone1000 simulate typical WGS samples while CloneSigSim simulates both WES and WGS samples), the depth of sequencing, the presence of subclonal copy number events (in DREAM and SimClone1000). Importantly, the SNVs in the DREAM dataset were simulated with a fixed distribution of signature activities across subclones, so this is a case where we do not expect a benefit for CloneSig compared to other methods. The PhylogicSim500 and SimClone1000 datasets were generated without concern about mutational signatures, so we can add a mutation type to each simulated SNV (among the 96 possibilities) using either constant or varying signature activities across clones. Similarly, for CloneSigSim, we can simulate SNV under both scenario. In order to assess the performance of CloneSig under various scenarios, we therefore consider two sets of simulations: 1) the “constant” scenario, comprising DREAM, PhylogicSim500, SimClone1000 and CloneSigSim simulated with constant signature activities across subclones, and 2) the “varying” scenario, comprising PhylogicSim500, SimClone1000 and CloneSigSim simulated with varying signature activities across subclones.

In order to measure the correctness of the subclonal reconstruction we use four different metrics adapted from [20] and described in details in the Methods section. Briefly, score1B measures how similar the true and the estimated number of clones are, score1C assesses in addition the correctness of frequency estimates for each subclone, score2A measures the adequacy between the true and predicted co-clustering matrices, and score2C the classification accuracy of clonal and subclonal mutations. We also assess the performance of eight other state-of-the-art methods for ITH estimation and compare them to CloneSig. First we evaluate TrackSig [18], that reconstructs signature activity trajectory along tumor evolution by binning mutations in groups of 100 with decreasing CCFs, and for each group performs signature activity deconvolution using an expectation-maximization (EM) algorithm. A segmentation algorithm is then applied to determine the number of breakpoints, from which we obtain subclones with different mutational processes. A recent extension integrating CCF in the segmentation algorithm to also perform subclonal reconstruction, TrackSigFreq [19] is also considered. Because of this rationale, the authors recommend to have at least 600 observed mutations to apply TrackSig or TrackSigFreq. For sake of completeness, however, we also apply TrackSig with fewer mutations in order to compare it with other methods in all settings. Third, we test Palimpsest [17], another method which associates mutational signatures and evolutionary history of a tumor. In Palimpsest, a statistical test based on the binomial distribution of variant and reference read counts for each mutation is performed, with correction for copy number, in order to classify mutations as clonal or subclonal. Then, for each of the two groups, signature activity deconvolution is performed using non-negative matrix factorization (NMF). Those methods are representative of the main approaches to the signature refitting problem: NMF-based approaches, and probability-based approaches [14, 15]. This limitation to two populations can induce a bias in the metrics 1B, 1C and 2A that are inspired from [20], so we introduce the metric 2C to account for the specificity of Palimpsest. Finally, we test five other popular methods for ITH reconstruction which do not model mutational processes: PyClone [7], PhylogicNDT [28] and DPClust [6], which are Bayesian clustering models optimized with a Markov Chain Monte Carlo (MCMC) algorithm, Ccube [8], another Bayesian clustering model, optimized with a variational inference method, and SciClone [29], also a Bayesian clustering model, optimized with a variational inference method, that only focuses on mutation in copy-number neutral regions.

##### Box 1: CloneSig’s algorithm

###### 1 Input

For a given tumor we observe

- *p*, the tumor purity of the sample, and for each SNV,
- *B_n_* and *D_n_* are respectively the variant and total read counts,
- *C_n_* is the copy number state,
- *T_n_* is the trinucleotide context.

###### 2 Probabilistic model

This network summarizes the structure of the probabilistic graphical model underlying CloneSig. Each node represents a random variable, shaded ones being observed, and edges between
two nodes describe a statistical dependency encoded as conditional distribution
in CloneSig.

- *U_n_*, the clone or subclone where the SNV occurs,
- *S_n_*, the clone-dependent mutational process that generates the mutation,
- *M_n_*, the number of chromosomal copies harboring the mutation.

and observed variables, conditionally on latent variables, with

*π* the exposures of active signatures in each subclone,
*ϕ* the cancer cell fraction of each clone,
*ξ* the proportions of SNVs in each clone,
*ρ* the overdispersion parameter of read count observations.

###### 3 Objective

CloneSig estimates the parameters of the distributions of latent (unobserved) variables We want *θ* = (*ξ*, *π*, *ρ*, *ϕ*) to maximize the likelihood of observed data.

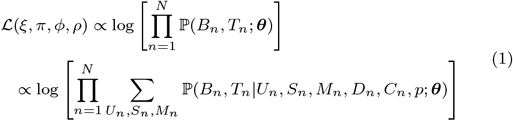

###### 4 Model fitting

Optimization with an expectation-maximization (EM) algorithm, with projected Newton method at M step.

###### 5 Output

Optimal values for *θ* = (*ξ*, *π*, *ρ*, *ϕ*). We can use CloneSig’s model to obtain a posteriori most likely values for latent variables *U_n_*, *S_n_*, *M_n_* or consider the probabilities for each possible value.

**Box 1:**
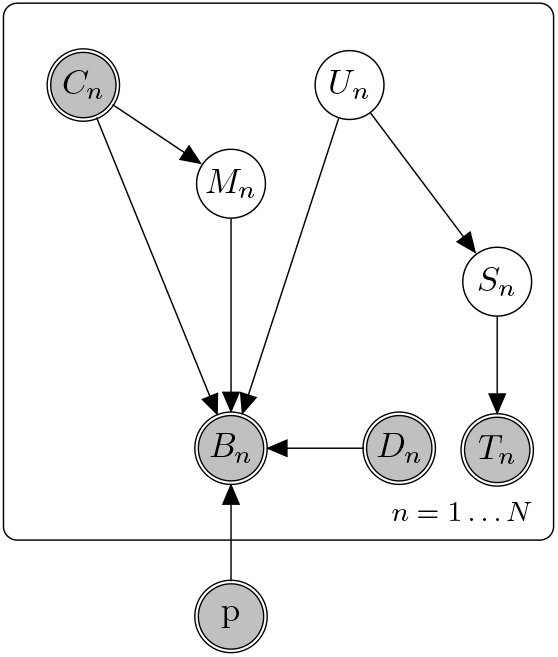
Probabilistic graphical model for CloneSig. This plot summarizes the structure of the probabilistic graphical model underlying CloneSig. Each node represents a random variable, shaded ones being observed, and edges between two nodes describe a statistical dependency encoded as conditional distribution in CloneSig. For a given tumor we observe *p*, the tumor purity of the sample, and for each SNV, *B_n_* and *D_n_* are respectively the variant and total read counts, *C_n_* is the copy number state, and *T_n_* is the trinucleotide context. Unobserved latent variable include *U_n_*, the clone or subclone where the SNV occurs, *S_n_*, the clone-dependent mutational process that generates the mutation, and *M_n_*, the number of chromosomal copies harboring the mutation. See the main text for details about the distributions and parameters of the model.

Figure 2 summarizes the subclonal reconstruction performance of CloneSig and other ITH reconstruction methods, under both the “constant” and “varying” scenarios. Each radar plot shows the average scores (1B, 1C, 2A and 2C) reached by each method on a set of simulated data. Under the “constant” scenario, we see that CloneSig is on par with or better than the best ITH methods (PhylogicNDT, DPClust and Ccube) on all scores and across all simulators, while SciClone, TrackSig and TrackSigFreq have overall poorer performance. This confirms that CloneSig reaches state-of-the-art performance even if the main assumption underlying its model is not met in the data, which is an important property since the question of how often subclonal populations with different signature activities emerge in tumors remains open. Moving to the “varying” setting, where signature activities vary across subclones, we see that, as expected, CloneSig, TrackSig and TrackSigFreq improve their performance, at least on the PhylogicSim500 and CloneSigSim simulations. Interestingly, CloneSig outperforms all methods on all simulations in that setting, confirming the potential benefits of using CloneSig on tumors where signature activities vary across subclones.

**Figure 2:**
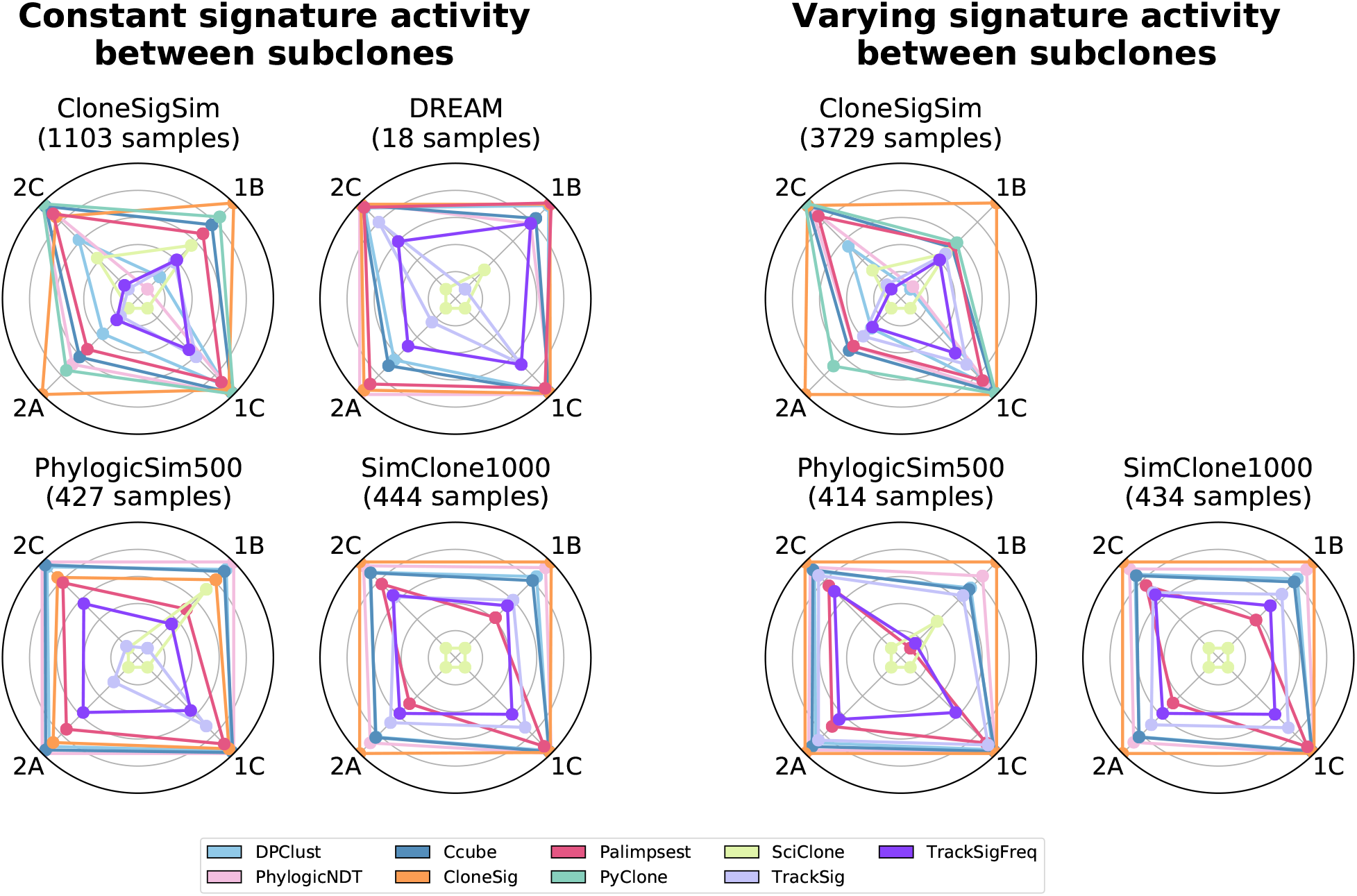
Subclonal reconstruction performance of CloneSig and eight ITH reconstruction methods on simulated data. Each radar plot summarizes the performance of the nine ITH reconstruction methods according to four performance measures on one set of simulated data. In short, score1B evaluates the number of clones found by the method, score1C the resulting mutation CCF distribution, score2A the co-clustering of mutations in the defined clones, and score2C the classification of subclonal versus clonal mutations. All scores are normalized such that the best performing method lies on the outer circle, and the worst near the center, to enhance visual distinction between methods. We have ensured that all scores are comparable by averaging them only on simulations where all methods successfully produced an estimate under reasonable computation time and memory limits; PyClone results are not shown on the DREAM, PhylogicSim500 and SimClone1000 datasets because it failed to produce an output too often. The four simulated sets on the left follow the “constant” scenario, where signature activities are the same across all subclones, while the three simulated sets on the right follow the “varying” scenario, where signature activities vary between subclones. Source data are provided as a Source Data file.

While Figure 2 shows average performances across hundreds of simulations with different parameters, it is also interesting to dig deeper into how the performance of each method fluctuates as a function of simulation parameters such as the number of clones in the tumor or the number of mutations simulated. Figure 3 summarizes this for the PhylogicSim500 simulations (in both the “varying” and “constant” settings, corresponding respectively to a “varying” or “constant” signature setting), and we refer the reader to Supplementary Note 2 for similar analyses for other simulators.

**Figure 3:**
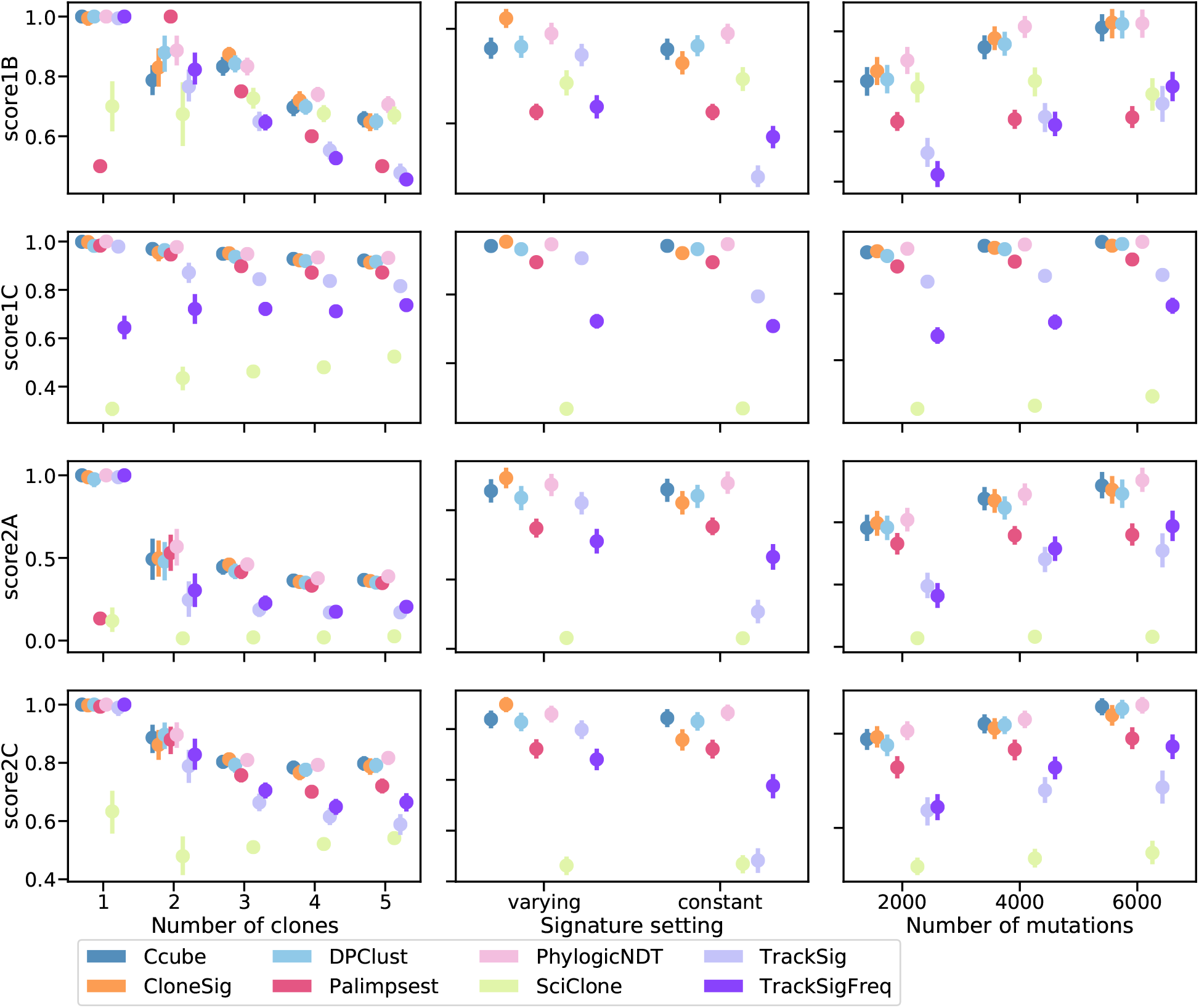
Detailed subclonal reconstruction performance on the PhylogicSim500 simulations. This figure shows the four performance scores (rows) of the different ITH reconstruction methods, when we vary one simulation parameter such as the number of clones simulated (left), the signature activity pattern to mimic the “varying” or “constant” scenario (middle), and the number of mutations observed rounded to the closest thousand (right). Each point represents the average of the scores over all simulated samples with a given parameter value. Bootstrap sampling of the scores was used to compute 95% confidence intervals, which are not visible if smaller than the dot. We have ensured that all scores are comparable by keeping only simulations where all methods output a prediction; PyClone results were excluded because they led to exclude too many samples. There are respectively 85, 47, 223, 250 and 236 samples with 1, 2, 3, 4 and 5 clones, 427 samples with constant and 414 with varying signature activities between clones, and 265, 347 and 229 samples with approximately 2000, 4000 and 6000 observed point mutations. Source data are provided as a Source Data file.

Regarding the estimation of the number of clones (score1B), CloneSig, Ccube, DPClust and Phylogic-NDT exhibit the best performances in most settings except for 2 clones where they are outperformed by Palimpsest, which however systematically predicts two clones by design. In the unfavourable “constant” scenario, PhylogicNDT has the best score, which may reflect the fact that this particular dataset is simulated according to PhylogicNDT’s model and might therefore be biased towards that method by construction. As already observed in Figure 2, CloneSig, TrackSig and TrackSigFreq see their performance increase in the favourable “varying” scenario, which is expected as they can leverage extra information to distinguish clones in cases where signature activity varies between distinct clones. SciClone tends to find a high number of clones, explaining its relatively good score1B for heterogeneous samples with 5 or 6 clones. However, other scores do not have the same positive evolution, revealing SciClone’s limits. Regarding the impact of the number of mutations on score1B, we see that CloneSig and TrackSig outperform all other comparable methods when the number of mutations is lower, illustrating the advantage of considering extra information in the ITH reconstruction process (we have already made clear that Palimpsest’s estimation of the number of clones is irrelevant, and TrackSig’s score has a very large error bar). As expected, all methods improve when the number of SNVs increases. Regarding score1C, which focuses not on the number of clones estimated by the ITH methods but on their ability to correctly recapitulate the distribution of CCF values, we see that all methods except SciClone, TrackSig and TracksigFreq have almost a perfect performance in all settings. TrackSigFreq performs slightly worse than TrackSig, but this may be explained by its poor performance when the number of mutations is too low, as performance is closer to the other methods as the number of mutations increases. Finally, SciClone is clearly the worst performing method for score1C.

Besides the ability of different methods to reconstruct the correct number of subclones and their CCF, as assessed by score1B and score1C, we measure with score2A their ability to correctly assign individual mutations to their clones, an important step for downstream analysis of mutations in each subclone. Similarly to other scores, Ccube, DPClust, PhylogicNDT and CloneSig have the best (and similar performances) in the majority of settings. For all methods, score2A decreases when the number of clones increases, and increases with more observed mutations. Again, when comparing “constant” and “varying” settings for signature activity, PhylogicNDT appears as the best performing method over all “constant” samples, and CloneSig dominates in the “varying” setting. Finally, when we assess the capacity of each method to simply discriminate clonal from subclonal mutations using score2C, a measure meant not to penalize Palimpsest which only performs that task, we see again that CloneSig is among the best methods in all scenarios, in particular with fewer observed mutations, and very close to by Ccube, DPClust and PhylogicNDT in other cases. Palimpsest is a bit below these methods, as well as TrackSig and TrackSigFreq. Again, CloneSig, TrackSig and TrackSigFreq benefit from the “varying” signature activity setting.

To further illustrate the interplay between signature change and ability to detect clones, we now test CloneSig, TrackSig, TrackSigFreq and Palimpsest on simulations with exactly two clones, and where we vary how the clones differ in terms of CCF, on the one hand, and in terms of mutational processes, on the other hand (quantified in terms of cosine distance between the two profiles of mutation type). Figure 4 shows the area under the ROC curve (AUC) for the correct classification of clonal and subclonal mutations by CloneSig as a function of these two parameters. We see an increased AUC as the distance between the mutation type profiles increases, for a constant CCF difference between the clones. For example, when two clones have similar signatures (small cosine distance), the AUC is around 0.7 when the difference between their CCF is around 0.2; when their signatures are very different (large cosine distance), the same performance can be achieved when their CCF only differ by 0.1 or slightly below. We show in Supplementary Figure 110 how other parameters (number of mutations, sequencing depth, diploid proportion of the genome, choice of input signature) also impact the performance of CloneSig in this setting. We also present similar results for other methods that both account for mutation frequency and mutational context in Supplementary Figure 111, as well as the influence of the number of observed mutations for each of them in Supplementary Figure 112.

**Figure 4:**
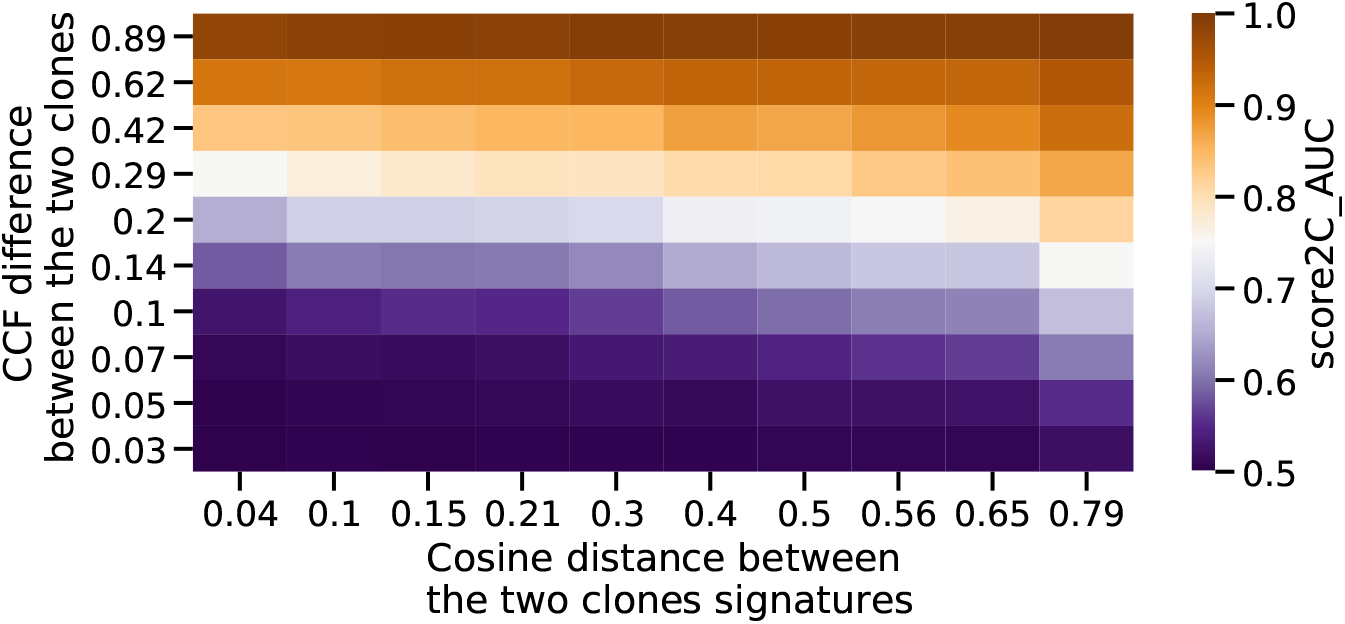
Accuracy of the number of clones estimated by CloneSig. To illustrate the factors that impact the validity of the subclonal reconstruction performed by CloneSig, we have simulated samples with two clones, varying both the difference in the cancer cell fraction (CCF) between the two clones (vertical axis) and the cosine distance between their mutational profiles (horizontal axis). The accuracy denotes the proportion of runs where CloneSig rightfully identifies two clones. Source data are provided as a Source Data file.

### 2.3 Performance for signature activity deconvolution

In addition to ITH inference in terms of subclones, CloneSig estimates the mutational processes involved in the tumor and in the different subclones. We now assess the accuracy of this estimation on simulated data, using five performance scores detailed in the Methods section. In short, score_sig_1A is the Euclidean distance between the normalized mutation type counts and the reconstructed profile (activity-weighted sum of all signatures); score_sig_1B is the Euclidean distance between the true and the reconstructed profile; score_sig_1C measures the identification of the true signatures; score_sig_1D is the proportion of signatures for which the true causal signature is correctly identified; and score_sig_1E reports the median of the distribution of the cosine distance between the true and the predicted mutation type profile that generated each mutation. We compare CloneSig to the three other methods that perform both ITH and mutational process estimation, namely, TrackSig, TrackSigFreq and Palimpsest, and add also deconstructSigs [13] in the benchmark, a method that optimizes the mixture of mutational signatures of a sample through multiple linear regressions without performing subclonal reconstruction.

Figure 5 shows the performance of the different methods according to the different metrics on the PhylogicSim500 dataset. For Score_sig_1A and Score_sig_1B, there is a clear advantage for CloneSig, TrackSig and TrackSigFreq over Palimpsest and deconstructSigs in all scenarios except when the number of mutation is the smallest, in which case all methods behave similarly. For Score_sig_1C, CloneSig, TrackSig and TrackSigFreq exhibit the best AUC to detect present signatures in all scenarios. This may be related to a better sensitivity as CloneSig and TrackSig perform signature activity deconvolution in subsets of mutations with less noise. All methods perform similarly with respect to Score_sig_1D. The median cosine distance (Score_sig_1E) is slightly better for CloneSig compared to other methods in all settings, particularly when there are three clones or more, and when signature activity varies across subclones.

**Figure 5:**
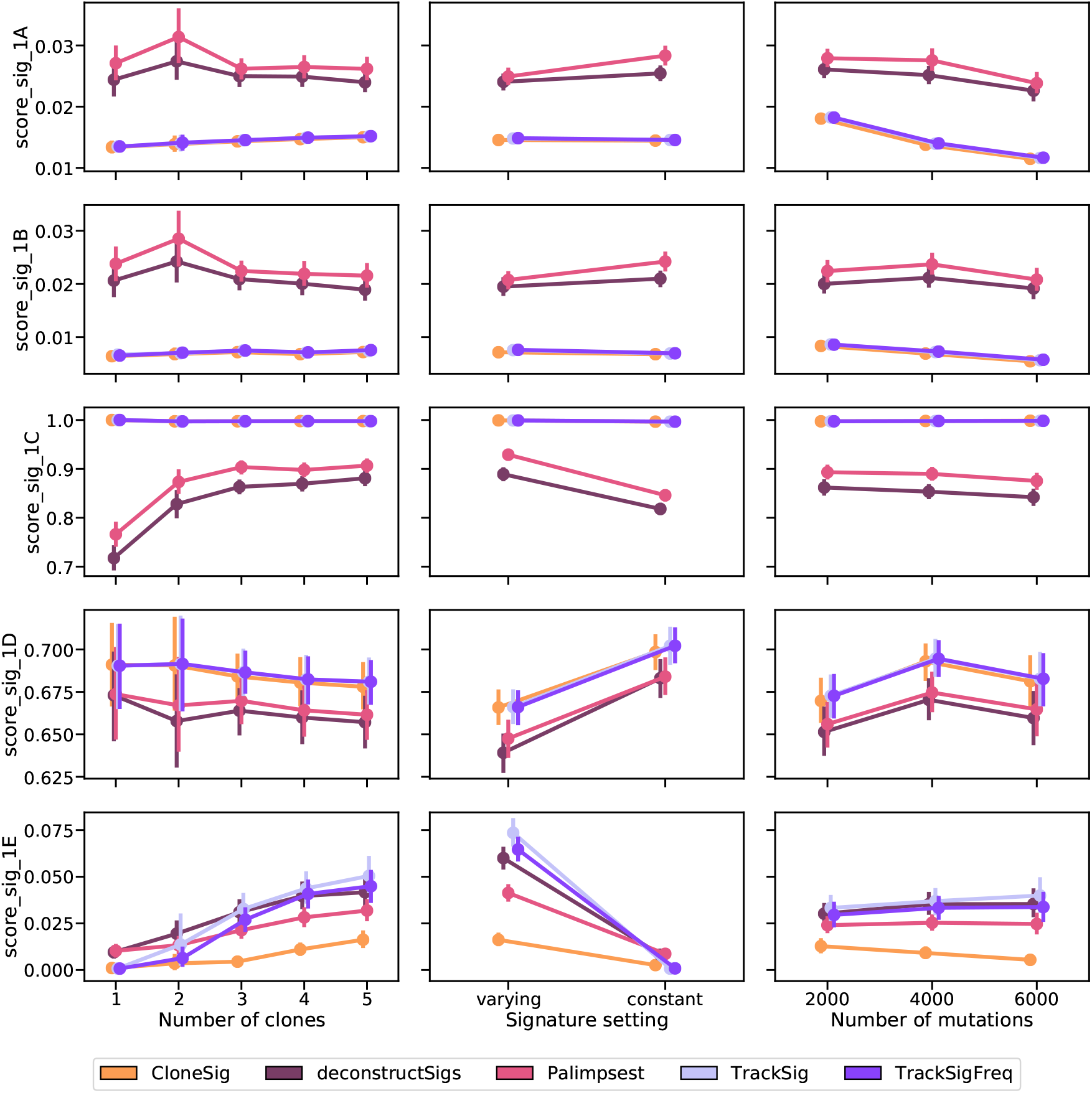
Comparison of CloneSig, deconstructSigs, Palimpsest, TrackSig and TrackSigFreq for signature activity deconvolution on the PhylogicSim500 dataset. Several metrics have been implemented, and are detailed in the main text. Scores_sig_1A and 1B are distances between the estimated mutation type profile (as defined by the signature activity proportions) to the true mutation profile (defined using the parameters used for simulations, 1A), and the empirical observed mutation profile (defined using available observed mutations, 1B), and is better when close to 0. Score_sig_1C is the area under the ROC curve for the classification of signatures as active or inactive in the sample, and is better when close to 1. Score_sig_1D is the proportion of mutations for which the correct signature was attributed, and is better when close to 1. Finally, Score_sig_1E is the median distance to the true mutation type profile of the clone to which a mutation was attributed from the true distribution of its original clone in the simulation, and is better when close to 0. The results are presented depending on several relevant covariates: the true number of clones (left), the signature activity setting (middle), and the number of mutations (right). Each point represents the average of the score over all available simulated samples. We used bootstrap sampling of the scores to compute 95% confidence intervals. There are respectively 87, 47, 223, 250 and 236 samples with 1, 2, 3, 4 and 5 clones, 428 samples with constant and 415 with varying signature activities between clones, and 267, 347 and 229 samples with approximately 2000, 4000 and 6000 observed point mutations. Source data are provided as a Source Data file.

Overall, as for ITH inference, we conclude that CloneSig is as good as or better than all other tested methods in all tested scenarios. Additional results where we vary other parameters in each methods, notably the set of mutations used as inputs or the set of signatures used as prior knowledge, can be found in Supplementary Note 2; they confirm the good performance of CloneSig in all tested settings. We also present a thorough evaluation of signature activity deconvolution on the CloneSigSim, SimClone1000 and DREAM simulated datasets in Supplementary Note 2, which overall confirm the results observed on the PhylogicSim500 dataset.

### 2.4 Pan-cancer overview of signature changes

We now use CloneSig on real pan-cancer data, to analyze ITH and mutational process changes in 8,951 tumor WES samples from the TCGA cohort spanning 31 cancer types, and in 2,632 WGS samples from 32 cancer types analyzed by the PCAWG initiative which represents the largest dataset of cancer WGS data to date.

For each sample in each cohort, we estimate with CloneSig the number of subclones present in the tumor, the signatures active in each subclone, and test for the presence of a significant signature change between clones. Based on samples exhibiting a significant signature change, we attempt to identify the signatures that are the most variant for each cancer type. To that end, we compute the absolute difference in signature activity between the largest subclone and the set of clonal mutations, neglecting cases where the absolute difference is below 0.05. Figure 6 shows a global summary of the signature changes found in the TCGA cohort. For each cancer type with at least 100 patients (see Supplementary Note 3 for the analysis over all cancers), it shows the proportion of samples where a signature change is found, and a visual summary of the proportion of samples where each individual signature is found to increase or to decrease in the largest subclone, compared to the clonal mutations. The thickness of each bar, in addition, indicates the median change of each signature. We retained only signatures found variant in more than 10% of each cohort samples. Complete results for the PCAWG samples are presented in a similar way in Supplementary Note 4. Overall, CloneSig detects a significant change in signature activity in 39% of all samples of the TCGA and 61% of the PCAWG cohort, in line with previous reports of 30% [23] or 76% [2], although these proportions vary between cancer types.

**Figure 6:**
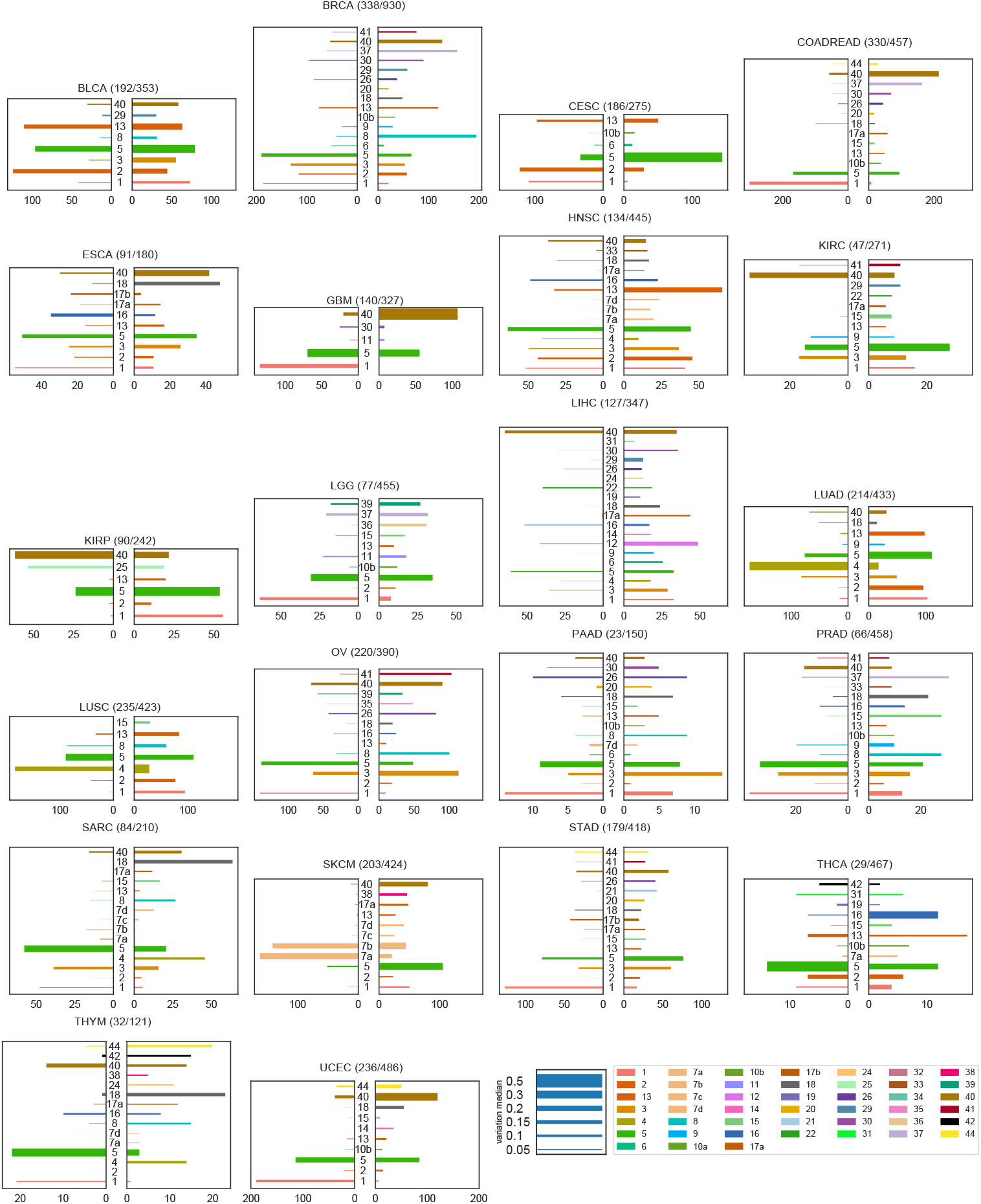
Mutational signature changes in the TCGA cohort. Each plot corresponds to one cancer type, indicates the number of samples with a significant signature change compared to the total number of samples, and shows on the right panel an increase of a signature in the largest subclone, compared to clonal mutations, and on the left panel a decrease. The length of each bar corresponds to the number of patients with such changes, and the thickness to the median absolute observed change. Only the 22 cancer types with more than 100 patients are represented: bladder urothelial carcinoma (BLCA), breast invasive carcinoma (BRCA), cervical squamous cell carcinoma and endocervical adenocarcinoma (CESC), colorectal adenocarcinoma (COADREAD), esophageal carcinoma (ESCA), glioblastoma multiforme (GBM), head and neck squamous cell carcinoma (HNSC), kidney renal clear cell carcinoma (KIRC), kidney renal papillary cell carcinoma (KIRP), brain lower grade glioma (LGG), liver hepatocellular carcinoma (LIHC), lung adenocarcinoma (LUAD), lung squamous cell carcinoma (LUSC), ovarian serous cystadenocarcinoma (OV), pancreatic adenocarcinoma (PAAD), prostate adenocarcinoma (PRAD), sarcoma (SARC), skin cutaneous melanoma (SKCM), stomach adenocarcinoma (STAD), thyroid carcinoma (THCA), thymoma (THYM), uterine corpus endometrial carcinoma (UCEC). Full results for all cancer types are available in Supplementary Note 3. Source data are provided as a Source Data file.

Let us first compare signature variations found using CloneSig in the TCGA and the PCAWG cohorts, to assess the consistency between results obtained by CloneSig on WES and WGS data. Interestingly, for most cancer types, namely BLCA, BRCA, CESC, GBM, HNSC, KIRP, LUAD, LUSC, OV, SKCM and UCEC, very similar patterns of signature changes are found in both cohorts, while some minor differences are found in other types. For example, in esophageal cancer (ESCA in the TCGA, Eso-AdenoCA in the PCAWG) a strong decrease in signature 16 activity is observed in the TCGA, but not in the PCAWG. Such a decrease pattern is however consistent with the association between signature 16 and alcohol consumption [30], and recurrent patterns of decrease of lifestyle-associated signature activity [23, 2]. For liver tumors, a strong decrease of signature 12 (unknown aetiology) in liver cancers is observed in the PCAWG (Liver-HCC), but variations in both directions are found in the TCGA (LIHC), which is coherent with a similar observation made on an independent cohort of 44 WGS samples of liver tumors [30]. Additionally, a tendency to decrease for signature 16 and increase for signature 17 activities is found by CloneSig on the TCGA cohort, which is coherent with [30] who found on a independent cohort a systematic decrease of signature 16 activity in all patients and an increase in signature 17 for one patient, while these variations are not clearly found on the PCAWG cohort. In prostate cancers, CloneSig identifies both increases and decreases of signatures 3 and 5 activities in the TCGA (PRAD), while only in one direction in the PCAWG (Prost-AdenoCA). Both increases and decreases of those two signature activities were previously reported in an independent cohort of 293 whole-genome sequenced localized prostate tumors [25]. In short, despite a few minor differences, the agreement between CloneSig’s results on the two cohorts is excellent, and in case of discrepancy the signals found on the larger TCGA cohort seem to be more coherent with current literature than those found on PCAWG. This illustrates the ability of CloneSig to detect patterns of ITH and signature activity change using WES or WGS data, and to benefit from the availability of larger WES cohorts.

To further explore the reliability of CloneSig’s results on real data, we now compare our findings in the PCAWG to results obtained on the same dataset with two different methods: TrackSig [2], and the careful distinction of early and late (and clonal and subclonal mutations) in [23]. Overall, CloneSig’s predictions are very similar to both studies for several important signals, such as lifestyle-associated signatures associated with tobacco-smoking (signature 4) and UV light exposure (signature 7) that decrease systematically in lung tumors and oral cancers (Lung-AdenoCA, Lung-SCC and Head-SCC) and skin melanoma (Skin-Melanoma) respectively. Similarly, CloneSig finds a strong decrease of signature 17 (damage by reactive oxygen species) in esophageal tumors (Eso-AdenoCA), and of signature 9 (Polymerase eta somatic hypermutation) in Chronic Lymphoid Leukemia (Lymph-CLL), which was also found by [23, 2]. Besides these coherent predictions, CloneSig finds additional differences not reported by [23, 2]. For example, CloneSig finds an increase in signature 3 (defective homologous recombination-based DNA damage repair) in bladder tumors (Bladder-TCC) unreported by [23, 2], but consistent with evidence of homologous recombination repair modulation being a marker of cancer progression in Bladder [31]. CloneSig also identifies an unreported increase in signature 8 activity in several cancer types, namely breast, pancreatic, and prostate tumors, both in TCGA and PCAWG cohorts. The same increase was reported for two out of ten multi-sample, whole-genome breast cancer cases with a local relapse or distant metastatic [32], and in some prostate tumors [25]. Also, an analysis of genomic location of signature 8 SNVs suggests that this signature arises during cancer progression [33]. However, signature 8 is deemed difficult to identify due to potential confusion with signature 3 [33], potentially contributing to discrepancies between CloneSig’s results and previous studies.

To complement the analysis of the TCGA and the PCAWG cohorts, we provide heatmaps to delineate an overview of each cancer type in Supplementary Figures 38 to 68 and 71 to 102 respectively. For each type, the first panel represents the difference between subclonal and clonal signature activities (in case of a significant change in activity), and the bottom panel represents the absolute values of each signature activity for clonal SNVs (belonging to the clone of largest CCF estimated by CloneSig), and in the main subclone (in terms of number of SNVs). This allows researchers to fully explore CloneSig’s results, and further compare their results in future studies. For the TCGA, in each panel, we have added several clinical variables, in particular, the patient’s age at diagnosis and the patient’s sex. Overall, we found no trend of association between signature activities or change in activities and those clinical characteristics, as previously observed in the particular case of prostate cancer [25]. In most types, like CESC (Figure S41), HNSC (Figure S47) and others, we observe groups of patients with different patterns of signature activity. The clinical significance of such groups remains to be further explored. Comparable clinical information is not available for the PCAWG cohort.

Altogether, we note a good agreement between findings on the TCGA WES and PCAWG WGS datasets using CloneSig, and previous similar analyses of the PCAWG WGS cohort, but also some differences which may be due to several factors including (i) the fact that both cohorts differ in size, patient clinical profiles and treatments, (ii) the fact that the algorithms used for the analysis are different, (iii) the fact that the TCGA cohort focuses on exonic mutations while PCAWG is based on WGS, and (iv) the fact that we and [2] focus on signatures with large activity changes in absolute difference while [23] focuses more on a general description of the cohort, report changes in log-fold change, and go beyond SNVs by integrating doublet-based substitutions (DBS) and small insertions and deletions (ID) signature changes. To mitigate these different effects, we provide an analysis of CloneSig’s results on the TCGA and on the PCAWG using log-fold change as a metric in Supplementary Figures 37, 70, that gives a slightly different view of the results.

## 3 Discussion

In recent years, a large number of methods have been developed to unravel ITH in tumors [7, 29, 8, 34, 6], and have been applied to different cohorts, including the TCGA. Recent analyses illustrate limits encountered when applying those methods to bulk WES [35, 10], as the number of observed mutations is small, the variance in read counts can be high, and a unique sample may miss the heterogeneity of the tumor. As sequencing costs are continuously decreasing, WGS, multi-sample sequencing and single cell sequencing will constitute relevant alternatives and simplify the study of ITH. However, to date a much larger number of tumor samples with sufficient clinical annotation is available with WES compared to other more advanced technologies, and can lead to interesting insights. Beyond the number of clones present in a tumor, another relevant aspect of tumor evolution is the presence of changes in mutational signatures activities [5], which could have clinical implications in cancer prevention and treatment, and unravel the evolutionary constraints shaping early tumor development. To the best of our knowledge, TrackSig [18], TrackSigFreq [19] and Palimpsest [17] are the only methods addressing the problem of systematic detection of signature changes, but they present serious limitations: Palimpsest first detects ITH, and then performs signature activity deconvolution, which has the major drawback that if this first step fails, no signature change can be detected. Moreover, Palimpsest simply aims to distinguish subclonal from clonal mutations, thus ignoring more complex patterns. TrackSig and TrackSigFreq are only applicable to WGS data, and though avoiding the caveat of relying on a previous detection of ITH for TrackSig, the final step of associating signature changes to the subclonal reconstruction is manual. TrackSigFreq is meant to be an extension of TrackSig to detect also clones not distinguished by a change in signature activity, but shows mitigated results on our benchmark compared to TrackSig. Finally, none of these methods efficiently leverages the changes in signature activity to inform and improve the ITH detection step. To overcome these limitations, we have developed CloneSig, the first method to offer joint inference of both subclonal reconstruction and signature activity deconvolution, which casn be applied to WGS as well as to WES data.

### 3.1 Improved ITH and signature detection in WES

CloneSig is a generative probabilistic graphical model that considers somatic mutations as derived from a mixture of clones where different mutational signatures are active. We demonstrated with a thorough simulation study the benefits of the joint inference in detecting ITH, both in WES and WGS samples. We included four different datasets to obtain a thorough evaluation of CloneSig and other ITH methods. Resorting to several datasets is necessary to obtain a complete and fair evaluation of the algorithms in a variety of situations. First, two of those datasets are generated using exactly some of the reconstruction methods: CloneSigSim and PhylogicSim500, respectively, follow the models of CloneSig and PhylogicNDT. The two other datasets do not explicitly follow the model of an evaluated method, but might also be biased towards one or more methods. Second, the DREAM and SimClone1000 have subclonal copy number events, while the other two datasets have only clonal events, which allows us to further assess the effect of neglecting them in CloneSig and other evaluated methods. Third, the simulated samples from the different datasets highly vary in number of observed SNVs, encompassing characteristics from WES and WGS data. Finally, although some relevant characteristics of our benchmark such as a very low number of observed SNVs are only covered by the CloneSigSim dataset, integration of gold-standard datasets allow us to contribute to establishing evaluation standards for ITH methods, and to facilitate comparison of CloneSig with (future) methods not considered in our study. We showed that CloneSig is competitive with or outperforms state-of-the art ITH methods, even in the absence of signature activity change between the clones, and is particularly efficient for the detection of samples with one or a few subclones. Interestingly, several other methods we considered including PyClone [7], SciClone [29], DPClust [6], PhylogicNDT [28] and Ccube [8], are fully Bayesian and choose the number of clones by maximizing of the posterior probability of the data. In those methods the prior has a regularizing role, and they exhibit a decrease of accuracy as the number of observed mutations increases. This may be related to the fact that the regularizing prior is less influential as more mutations are taken into account. We instead developed a specific adaptive criterion to estimate the number of clones, as we observed that standard statistical tools for model selection performed poorly in preliminary experiments.

Regarding the signature activity deconvolution problem, results on simulations (Score_sig_1C) suggest that CloneSig exhibits an improved sensitivity. Application to the TCGA also indicates such increased sensitivity: in the TCGA pancreatic ductal adenocarcinoma cohort (PAAD), the original study using deconstructSigs could not detect signature 3 activity in samples with somatic subclonal mutations in genes BRCA1 and BRCA2 [36], while CloneSig reports signature 3 exposure in some PAAD tumors.

### 3.2 Limits of CloneSig

It is important to explicitly state some of CloneSig’s limits, a number of them largely shared by most ITH methods. CloneSig is currently limited to SNVs, and does not account for indels or structural variants. Regarding the copy number profile of the tumor, CloneSig only considers clonal segments, but provides a complete framework to estimate a mutation’s multiplicity, which is an improvement over many existing methods. Regarding the range of application of CloneSig, the results are slightly below some pre-existing methods when fewer than 100 mutations are observed, as more stability is needed to fully benefit for robust estimates of signature activity, but exhibits very good performances for a number of mutations ranging from 100 to 10,000, as in the DREAM, SimClone1000, and PhylogicSim500 datasets, which makes it the most flexible method considered in this study. Regarding the runtime and scalability, resorting to subsampling above 10,000 observed mutations, as implemented in DPClust might be advisable.

Another point to be vigilant about when using CloneSig, as illustrated in simulations, and based on our experience with the TCGA and the PCAWG cohorts, is that the choice of the input signatures is key to CloneSig’s optimal performances. This is related to the unidentifiability of the signature activity deconvolution problem. Several solutions have been proposed: use of a pre-defined cancer-specific matrix [12, 18], selection of signatures based on other genomic information, such as patterns of indels or structural variants, or strand biases [12], or with other molecular or clinical covariates [37]. We have tested several such strategies (using all available signatures, or a well-chosen subsets), and CloneSig’s results highly depend on such a choice; we made similar observations for TrackSig, TrackSigFreq and Palimpsest. The probabilistic framework of CloneSig is well suited to integrate other mutation types (indels, structural variants), as well as prior knowledge on signature co-occurrence, and a prior based on other molecular and clinical covariates. The difficulty of this approach is the possibility to learn such association patterns. Another direction for further development would be to use CloneSig’s model to learn the signatures, or to allow some variations in the pre-defined signatures, as suggested in [38].

### 3.3 Clinical relevance of signature variations

To assess the clinical relevance of signature changes, we jointly analyzed clinical features available for the TCGA cohort and CloneSig results. We found no evidence of association between survival and the number of clones or the occurrence of a change of signature activity between subclones in any cancer type considered. We have also systematically analyzed whether we could identify an association between the exact pattern of signature change, i.e., the increase or decrease of a particular mutational signature and clinical variables, but found no significant association. However, more refined or complete analyses may be necessary to uncover the full significance of signature activity changes. Previous studies report important signature activity differences between early and metastatic tumors in endometrial and breast cancers [39, 40], with impact on the survival in the breast cancer study [40]. We could not perform a similar analysis using the TCGA with only untreated primary tumors, but this constitutes new directions and opportunities of research using CloneSig on metastatic cohorts, for instance to refine findings of [40], that compares signatures deconvoluted from the whole metastasis, and could benefit from subclonal analysis to distinguish early and late mutations.

A final potential clinical application could be usage as a marker for personalized treatment. For example, signature 3 is associated with homologous recombination repair defect (HRD), and a targeted therapy, PARP inhibitors, can successfully target cells with such defect. We may therefore use the detection of signature 3 to identify patients that can benefit from such therapy [41], and CloneSig exhibits better identification of active signatures, as illustrated in the simulation studies. Indeed, several mutations in genes like BRCA1 and 2, RAD51 are known to cause HRD, but some other mutations are less frequent, or other events may result in HRD and be undetectable using regular genome sequencing, such as epigenetic inactivation [42]. In addition, the intensity of HRD mutational process may be predictive of the treatment response. Pursuing this line of thought, the change in signature activity could also be exploited as an indicator of the current driver status of HRD in tumor development. Of course, these remain at this stage hypotheses that require further in silico and experimental exploration. In particular relating the presence or chance of a signature to the fact that it represents a driving process amenable to therapeutic intervention remains challenging, particularly in the absence of longitudinal data about tumor evolution. As the underlying processes of signatures keeps being uncovered, we nevertheless expect more examples of such clinical applications to arise. Signatures indicating sensitivity to PARP inhibitors, platinium-based chemotherapy, PD1-immunotherapy, cisplatin and resistance to tamoxifen have been identified [43].

## 4 Methods

### 4.1 CloneSig model

CloneSig is a probabilistic graphical framework, represented in Box 1, to model the joint distribution of SNV frequency and mutational context using several latent variables to capture the subclonal composition of a tumor and the mutational processes involved in each clone. For a given SNV it assumes that we observe the following variables: *D*, the total number of reads covering the SNV; *B* ≤ *D*, the number of mutated reads; *T* ∈ {1, …, 96} the index of the mutation type (i.e., the mutation and its flanking nucleotides, up to symmetry by reverse complement); and 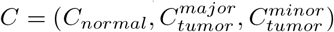 the allele-specific copy number at the SNV locus, as inferred using existing tools such as ASCAT [44]. Here *C_normal_* is the total copy number in normal cells, and 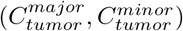 are respectively the copy number in the cancer cells of the major and minor allele, respectively. We therefore also observe 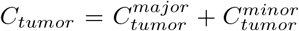, the total number in cancer cells. Finally, we assume observed the tumor sample purity *p*, i.e., the fraction of cancer cells in the sample.

In addition to those observed variables, CloneSig models the following unobserved variables: *U* ∈ {1, …, *J*}, the index of the clone where the SNV occurs (assuming a total of *J* clones); *S* ∈ {1, … *L*} the index of the mutational signature that generated the SNV (assuming a total of *L* possible signatures, given *a priori*); and 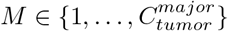, the number of chromosomes where the SNV is present. Note that here we assume that SNVs can only be present in one of the two alleles, hence the upper bound of *M* by 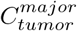. to delete

Denoting for any integer *d* by 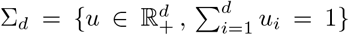 the *d*-dimensional probability simplex, and for *u* ∈ Σ*_d_* by Cat(*u*) the categorical distribution over {1, …, *d*} with probabilities *u*_1_, …, *u_d_* (i.e., *X* ∼ Cat(*u*) means that *P* (*X* = *i*) = *u_i_* for *i* = 1, …, *d*), let us now describe the probability distribution encoded by CloneSig for a single SNV; its generalization to several SNVs is simply obtained by assuming they are independent and identically distributed (i.i.d.) according to the model for a single SNV. We do not model the law of *C* and *D*, which are observed root nodes in Box 1, and therefore only explicit the conditional distribution of (*U, S, T, M, B*) given (*C, D*).

Given parameters *ξ* ∈ Σ*_J_*, π ∈ (Σ*_L_*)*^J^* and *μ* ∈ (Σ_96_)*^L^*, we simply model *U*, *S* and *T* as categorical variables:

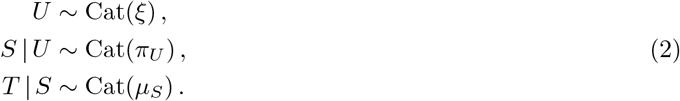

Conditionally on *C*, we assume that the number of mutated chromosomes *M* is uniformly chosen between 1 and 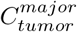, i.e.,

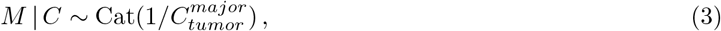

where 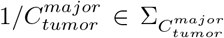 represents the vector of constant probability. Finally, to define the law of *B*, the number of mutated reads, we follow a standard approach in previous studies that represent ITH as a generative probabilistic model [7, 9, 8, 29] where the law of the mutated read counts for a given SNV must take into account the purity of the tumor, the proportion of cells in the tumor sample carrying that mutation (cancer cell fraction, CCF), as well as the various copy numbers of the normal and tumor cells. More precisely, as reviewed by [6], one can show that the expected fraction of mutated reads (variant allele frequency, VAF) satisfies

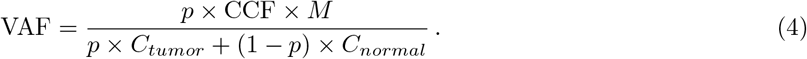

Note that this only holds under the classical simplifying assumption that all copy number events are clonal and affect all cells in the sample. If we now denote by *ϕ* ∈ [0, 1]*^J^* the vector of CCF for each clone, and introduce a further parameter 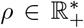 to characterize the possible overdispersion of mutated read counts compared to their expected values, we finally model the number of mutated reads using a beta binomial distribution as follows:

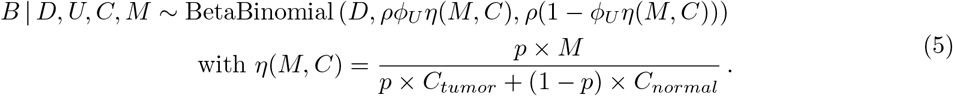

### 4.2 Parameter estimation

Besides the tumor purity *p*, we assume that the matrix of mutational processes *μ* ∈ (Σ_96_)*^L^* is known, as provided by databases like COSMIC and discussed below in Section 4.7. We note that we could consider *μ* unknown and use CloneSig to infer a new set mutational signatures from large cohorts of sequenced tumors, but prefer to build on existing work on mutational processes in order to be able to compare the results of CloneSig to the existing literature. Besides *p* and *μ*, the free parameters or CloneSig are *J*, the number of clones, and ***θ*** = (*ξ, ϕ, π, ρ*) which define the distributions of all random variables. On each tumor, we optimize ***θ*** separately for *J* = 1 to *J_max_* = 8 clones to maximize the likelihood of the observed SNV data in the tumor. The optimization is achieved approximately by an expectation-maximization (EM) algorithm [45] detailed in Supplementary Section 1.1. The number of clones *J*\* ∈ [1, *J_max_*] is then estimated by maximizing an adaptive model selection criterion, detailed in Supplementary Section 1.2.

### 4.3 Test of mutational signature changes

We use a likelihood ratio test to determine the significance of a signature change, by comparing a regular CloneSig fit to a fit with a single mixture of signatures common to all clones. To adapt the test, the parameter of the chi-squared distribution needs a calibration, that we perform on simulated data under the null hypothesis (without change of signatures between clones). We obtain the optimal parameter using a ridge regression model with the number of clones and the degree of freedom of the input signature matrix as covariates. The coefficient values are averaged over 10-fold cross-validation to ensure robustness. We provide more details about this test in Supplementary Section 1.3.

### 4.4 Simulations

We use several simulation strategies to evaluate the performance of CloneSig and other methods in various situations. We also use simulations to adjust several aspects of CloneSig, in particular the setting of a custom stopping criterion and the calibration of the statistical test to detect a significant signature change along tumor evolution.

#### 4.4.1 Default simulations

We implemented a class `SimLoader` to perform data simulation in CloneSig package. The user sets the number of clones *J*, the number of observed mutations *N*, and the matrix of *L* possible signatures *μ*. She can also specify the desired values for the CCF of each clone *ϕ* ∈ [0, 1]*^J^*, the proportion of each clone *ξ* ∈ Σ*_J_*, the exposure of each signature in each clone *π* ∈ (Σ*_L_*)*^J^*, and the overdispersion parameter *ρ* ∈ ℝ^+^* for the beta-binomial distribution, as well as the proportion of the genome that is diploid. If the user does not provide values for one or several parameters, we generate them randomly as follows:

*π* the number of active signatures follows a *Poisson*(7) + 1 distribution, and the signatures are chosen uniformly among the *L* available signatures. Then for each subclone, the exposures of active signatures follow a Dirichlet distribution of parameter 1 for each active signature;
*ϕ* the cancer cell fraction of each clone is set such that the largest clone has a CCF of 1, and each subsequent CCF is uniformly drawn in decreasing order to be greater than 0.1, and at a distance at least 0.05 from the previous clone;
*ξ* the proportions of clones are drawn from a Dirichlet distribution of parameter 1 for each clone. The proportions are repeatedly drawn until the minimal proportion of a clone is greater than 0.05;
*ρ* follows a normal distribution of mean 60 and of variance 5.

The same strategy is used for random initialization of the parameters for the EM algorithm.

The total copy number status is drawn for a user-set diploid proportion of the genome with a bell-like distribution centered in 2, and skewed towards the right (see Supplementary Figure 109 for examples), or from a rounded log-normal distribution of parameters 1 and 0.3. The minor copy number is then drawn as the rounded product between a beta distribution of parameters 5 and 3 and the total copy number. The multiplicity of each mutation *n* is uniformly drawn between 1 and 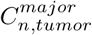. The purity is drawn as the minimum between a normal variable of mean 0.7 and of variance 0.1, and 0.99. The other observed variables (*T*, *B*, *D*) are drawn according to CloneSig probabilistic model.

#### 4.4.2 Simulations for comparison with other ITH and signature methods

To calibrate the custom stopping criterion and for further evaluation of CloneSig, we simulated a dataset comprising 7, 812 samples using the previously described setting, with a few adjustments: we set the minimal proportion of each clone to 0.1, the minimal difference between 2 successive clone CCFs to 0.1, and we chose the active signatures among the active signatures for each of the 31 cancer types from the TCGA, and curated using literature knowledge, as described later in Methods, in section 4.7. We draw the number of active signatures as the minimum of a *Pois*(7) + 1 distribution and the number of active signatures for this cancer type. We required a cosine distance of at least 0.05 between the mutational profiles of two successive clones.

In total, for each of the 31 cancer types, we generated a simulated sample for each combination of a number of mutations from the set {20, 50, 100, 300, 600, 1000, 5000} covering the range observed in WES and WGS data, a percentage of the genome that is diploid from the set {0%, 20%, 40%, 60%, 80%, 100%} to assess the impact of copy number variations, and finally, between 1 and 6 clones.

#### 4.4.3 Simulations without signature change between clones

We generated a set of simulations similar in all points to the one for comparison with other ITH and signature methods, except that there is a unique signature mixture common to all clones. We used this dataset in two contexts: (i) to evaluate CloneSig in comparison to other methods in the absence of signature change, and (ii) to design a statistical test to assess the significance of a change in mutational signatures. For the latter, the dataset was limited to the first eleven cancer types to avoid unnecessary computations (2,772 samples).

#### 4.4.4 Simulations to assess the separating power of CloneSig

To assess the separating power of CloneSig, we generated a dataset of 12,000 simulated tumor samples with two clones, where each clone represents 50% of the observed SNVs. Our objective was to explore the set of the distance between two clones, in terms of CCF distance, and of cosine distance between the two mutational profiles. For that purpose we first drew ten possible CCF distances evenly on a log scale between 0 and 1, and set to 1 the largest clone CCF. We also generated 50 matrices *π* with cosine distances covering regularly the possible cosine distances; to obtain them, we first generated 10,000 such *π* matrices to estimate an empirical distance distribution, and we implemented a rejection sampling strategy to obtain 50 samples from a uniform distribution. For each pair of CCF distance and *π* matrix, several samples were generated with the number of mutations varying among {30, 100, 300, 1000}, the diploid proportion of the genome among {0.1, 0.5, 0.9}, and the sequencing depth among {100, 500}.

#### 4.4.5 Simulations to assess the sensitivity of the statistical test

To measure the sensitivity of the statistical test to detect a significant signature change along tumor evolution, we generated a dataset of 3,600 simulated tumor samples with 2 to 6 clones. We used again a rejection sampling strategy to explore the space of the maximal distance between the profiles between any 2 clones, but the target distribution is here a beta distribution of parameters 1.5 and 8 as a target distribution, as the objective was to sample more thoroughly the small cosine distances. We repeated the sampling of 30 *π* matrices for 2 to 6 clones, and in each case, and generated several samples with the number of mutations varying among {30, 100, 300, 1000}, the diploid proportion of the genome among {0.1, 0.5, 0.9}, and the sequencing depth among {100, 500}.

### 4.5 Evaluation metrics

We use several evaluation metrics to assess the quality of CloneSig and other comparable methods. Some assess specifically the accuracy of the subclonal decomposition, while others assess the performance of signature activity deconvolution.

#### 4.5.1 Metrics evaluating the subclonal decomposition

The metrics described in this section evaluate the accuracy of the subclonal deconvolution. They are adapted from [20].

**Score1B** measures the adequacy between the true number of clones *J_true_* and the estimated number of clones *J_pred_*. It is computed as 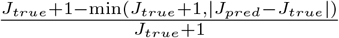
**Score1C** is the Wasserstein similarity, defined as 1 minus the Wasserstein distance between the true and the predicted clustering, defined by the CCFs of the different clones and their associated weights (proportion of mutations), implemented as the function `stats.wasserstein distance` in the Python package `scipy`.
**Score2A** measures the correlation between the true and predicted binary co-clustering matrices in a vector form, *M_true_* and *M_pred_*. It is the average of 3 correlation coefficients:

**Pearson correlation coefficient** 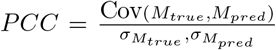, implemented as the function `pearsonr` in the Python package `scipy`,
**Matthews correlation coefficient** 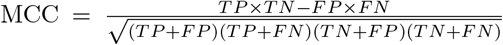, implemented as the function `metrics.matthews_corrcoef` in the Python package `scikit-learn`,
**V-measure** is the harmonic mean of a homogeneity score that quantifies the fact that each cluster contains only members of a single class, and a completeness score measuring if all members of a given class are assigned to the same cluster [46]; here the classes are the true clustering. We used the function `v_measure_score` in the Python package `scikit-learn`. Before averaging, all those scores were rescaled between 0 and 1 using the score of the minimal score between two “bad scenarios”: all mutations are in the same cluster, or all mutations are in their own cluster (*M_pred_* = **1**_*N*×*N*_ or 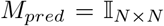). This score follows the strategy initially proposed in the preprint version [47] of [20], and suffers from a high memory usage, making it impractical beyond 20,000 mutations.
**Score2C** quantifies the accuracy of each method prediction of clonal and subclonal mutations. We report the accuracy, and the area under the ROC curve (implemented in function `metrics.roc auc score` in the Python package `scikit-learn`), sensitivity and specificity in Supplementary Note 2.

#### 4.5.2 Metrics evaluating the identification of mutational signatures

The metrics described in this section evaluate the accuracy of the mutational signature activity deconvolution.

**Score_sig_1A** computes the Euclidean distance between normalized mutation type counts (empirical), and the reconstituted profile. This is the objective function of most signature reconstruction approaches (including deconstructSigs[13] and Palimpsest [17]).
**Score_sig_1B** is the Euclidean distance between simulated and estimated signature profiles (weighted sum over all clones). This is closer to the objective of CloneSig, TrackSig [18] and TrackSigFreq [19].
**Score_sig_1C** measures the ability of each method to correctly identify present signatures. For CloneSig, no signature has a null contribution to the mixture, so for each clone, the signatures are considered in the decreasing order of their contribution to the mixture, and selected until the cumulative sum reaches 0.95. This rule is applied to all methods. For that metric, the area under the ROC curve (implemented in function `metrics.roc auc score` in the Python package `scikit-learn`) is reported, as well as the accuracy, sensitivity, and specificity in Supplementary Note 2
**Score_sig_1D** is the percent of mutations with the right signature. For each mutation, the most likely signature is found by taking into account the distribution of each mutation type in each signature, and the contribution of the signature to the mixture.
**Score_sig_1E** measures for each mutation the cosine distance between the clonal mutation type distribution that generated the mutation and the reconstituted one. We consider a unique global distribution for deconstructSigs. This allows us to measure the relevance of the reconstruction even if the wrong signatures are selected, as several signatures have very similar profiles. The result is a distribution of distances over all mutations, and we report the median of this distribution. We also report in Supplementary Note 2 more results with the minimum, the maximum, and the standard deviation of this distribution (`max diff distrib mut, median diff distrib mut`), as well as the proportions of mutations with a distance below 0.05 or 0.1 (`perc dist 5 and perc dist 10`).

### 4.6 Implementation

CloneSig is implemented in Python, and is available as a Python package at https://github.com/judithabk6/clonesig [75]. A wrapper function implements the successive optimization of CloneSig with increasing number of clones. For two clones and more, the model is initialized using results from the precedent run with one fewer clone, by splitting the subclone with the largest contribution to the mixture entropy as described in [48]. This process is stopped when the maximum number of subclones is reached, or when the selection criterion decreases for two successive runs. A class for simulating data according to the CloneSig model is also implemented, as detailed above.

### 4.7 Construction of a curated list of signatures associated with each cancer type

A very important input for CloneSig is the signature matrix. For application to the TCGA data, we restrict ourselves to signatures known to be active in each subtype. To that end, we downloaded the signatures found in the TCGA using SigProfiler [26] from synapse table syn11801497. The resulting list was not satisfactory as it lacked important known patterns; for instance signature 3, associated with homologous recombination repair deficiency was not found to be active in any tumor of the prostate cohort, while signature 3 in prostate cancer is well described in the literature [25, 55]. We therefore completed the signatures present in each cancer type based on the literature [56, 57, 30, 58, 59, 60, 61, 25, 62, 39, 63, 64, 12], and used the resulting matrix in all CloneSig runs on the TCGA. Our curated list of signatures present in each cancer type is provided in Table S4.

### 4.8 Quantification of signature changes

For analysis of the TCGA and the PCAWG cohorts, it can be of interest to go beyond the simple assessment of a significant signature change between clones, in particular to observe different patterns of variations for distinct cancer types. Using CloneSig’s results, such observations remain qualitative, as the statistical framework does not analyze changes at the individual signature activity level. Qualitatively, it might be difficult to strongly assess whether a signature’s activity increases or decreases because both patterns can occur in the case where more than two clones are found, and additionally, there might be complex branching evolution patterns between those clones that go beyond CloneSig’s abilities. To provide a first approximation, we measure such variations between the clonal clone (defined by having the highest cellular prevalence) and the largest subclone (defined as having the largest number of mutations). We have used two metrics to report such changes:

**absolute difference** *A_subclonal_ − A_clonal_*, where *A* is defined as the proportion of the activity (between 0 and 1). In that case, we have considered only cases where this difference is greater than 0.05 to avoid noisy reports.
**relative difference** 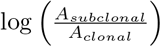; with exclusion of cases where both *A_subclonal_* < 10^−4^ and *A_clonal_* < 10^−4^, and otherwise, replacement of smaller values by 10^−4^ to avoid very large or small undue values

Each metric provides a different viewpoint of the data, with the absolute difference stressing variations of some importance, while the relative difference, though providing some focus on important variations on the logarithmic scale between two signature activities below 10^−2^.

For the graphical representations of Figures 7, S36 and S69, we have additionally applied two thresholds: only absolute differences greater than 0.05 in absolute value were considered, and only signatures varying in more than 10% of the cohort. Those thresholds were not applied in the additional representations as heatmaps (Supplementary Figures 38 to 68, and 71 to 102).

### 4.9 Survival analysis

We used the package survival in R for the survival analysis.

### 4.10 Data collection and processing

#### 4.10.1 TCGA cohort

We obtained CNV and CNA data for the TCGA cohort from the Genomic Data Commons (GDC). We downloaded annotated SNV for 33 cancer types obtained with four variant callers (MuSe, Mutect2, VarScan2 and SomaticSniper) and aligned to the GRCh38 human genome assembly. The GDC provides two sets, one denoted “protected” with complete outputs from the variant callers, with restricted access, and a “public” one, corresponding to a heavily filtered version of the “protected” set to ensure that all germline variants are removed. The documentation suggests to use the “protective” set if “omission of true-positive somatic mutations is a concern”. For the sake of completeness, we therefore applied CloneSig to both sets. Variant calling is known to be a challenging problem, and it is common practice to filter variant callers output, as ITH methods are deemed to be highly sensitive to false positive SNVs. We therefore filtered out indels from the public dataset, and considered the union of the 4 variant callers output SNVs. For the protected data, we also removed indels, and then filtered SNVs on the `FILTER` columns output by the variant caller (“PASS” only VarScan2, SomaticSniper, “PASS” or “panel of normals” for Mutect2, and “Tier1” to “Tier5” for MuSe). In addition, for all variant callers, we removed SNVs with a frequency in 1000 genomes or Exac greater than 0.01, except if the SNV was reported in COSMIC. We added a coverage filter, and we kept SNVs with at least 6 reads at the position in the normal sample, of which 1 maximum reports the alternative nucleotide (or with a variant allele frequency (VAF) <0.01), and for the tumor sample, at least 8 reads covering the position, of which at least 3 reporting the variant, or a VAF>0.2. The relative amount of excluded SNVs from protected to public SNV sets varied significantly between the 3 cancer types (see Table S3). All annotations are the ones downloaded from the TCGA, using VEP v84, and GENCODE v.22, sift v.5.2.2, ESP v.20141103, polyphen v.2.2.2, dbSNP v.146, Ensembl genebuild v.2014-07, Ensembl regbuild v.13.0, HGMD public v.20154, ClinVar v.201601. We further denote the filtered raw mutation set as “Protected SNVs” and the other one, which is publicly available, as “Public SNVs”. For CNA, we collected data from the ASCAT complete results on TCGA data partly reported on the COSMIC database [44, 50]. We then converted ASCAT results on hg19 to GRCh38 coordinates using the `segment_liftover` Python package [51]. ASCAT results also provide an estimate of purity, which we used as input to ITH methods when possible. Other purity measures are available [52]; however we selected the ASCAT estimate to ensure consistency with CNV (copy number variant) data. For all patients, we downloaded clinical data from the cBioPortal [49]

#### 4.10.2 PCAWG cohort

For the PCAWG cohort, we downloaded SNV and copy number calls from the ICGC data portal (data access DACO-1086821). Variant calls were generated by three pipelines run independently on each sample, with subsequent merging into a consensus set of high-quality calls. As described in [24], each of the three pipelines—labelled “Sanger” [65, 66, 67, 68], “EMBL/DKFZ” [69, 70] and “Broad” [53, 71, 72, 73] after the computational biology groups that created or assembled them—consisted of multiple software packages for calling somatic SNVs, small indels, CNAs and somatic SVs. Purity values represent the estimated fraction of cells in the sample derived from the tumour clone; ploidy values represent estimated average copy number in tumour cells. No further filtering or treatment was applied, except exclusion of indels, and SNVs outside of documented CNV segments (without CNV information). Matching of cancer types to the TCGA types was done using the provided table downloadable at https://dcc.icgc.org/api/v1/download?fn=/PCAWG/clinical_and_histology/tumour_subtype_consolidation_map.xlsx, and referenced in the ICGC data portal, using the second sheet (“Unique List of Tumour Types Aug”), as the “Live Version” sheet did not contain the contributing projects field, allowing mapping with the TCGA cohorts. For types absent from the TCGA (and hence missing the associated list of present signatures), we manually used the signatures reported in Figure 3 of the associated publication [12].

#### 4.10.3 DREAM simulated cohort

We obtained the preprocessed files and associated truth for the DREAM dataset [20, 21] from Synapse storage, accession ID syn2813581. Processed data includes mutation calling by Mutect2 [53], and the Battenberg algorithm [54] for CNV deciphering, and purity and ploidy estimation. We performed no further filtering, except neglecting the subclonal copy number events, as none of the evaluated method considered them (in that case, we considered the event with the largest cellular prevalence as clonal), and excluding indels, and SNVs without CNV information. To run signature methods in “cancertype” mode, that is with considering only a subset of all available 47 signatures, we attributed the tumors to a cancer type that contained all true signatures based on [12], and when several were possible, choosing a frequent one. T2 was considered of the type “Liver-HCC”, T3 “Lung-AdenoCA”, T4 and T5 “Breast”, and T6 “Colorect-AdenoCA”. All methods were run on this dataset, on a single CPU node per sample, with a time limit of 48 hours, and a memory limit of 10GB.

#### 4.10.4 PhylogicSim500 and SimClone1000

We obtained the 500 samples in the PhylogicSim500 dataset, and the 972 samples in the SimClone1000 dataset as described in [2, 22]. The data comprises the mutations called and CNV information, with subclonal CNV events in the PhylogicSim500 data but not in the SimClone1000 set. We ignored subclonal CNV events, and for each segment, we kept only the one with largest CCF in the input provided to all methods, as none of them deals with those events. As signature activity was not considered in the simulation, we used the ground truth to generate two versions of each sample, a “constant” setting with a signature activity distribution common to all clones (corresponding to the unfavourable scenario for CloneSig), and a “varying” setting with a different signature activity distribution for each clone (corresponding to the favourable scenario for CloneSig). In each sample, we drew active signatures according to the PCAWG cancer type specified for samples of the PhylogicSim500 dataset, and a random PCAWG cancer type, as it was not set for the SimClone1000 dataset. For the PhylogicSim500 dataset, each method was run on a single CPU node per sample, with a time limit of 48 hours, and a memory limit of 5GB, and for the SimClone1000, each method was run on a single CPU node per sample, with a time limit of 48 hours, and a memory limit of 15GB, only for samples with fewer than 10,000 SNVs to avoid unnecessary computations; this limit in SNV number was set based on observations on the DREAM dataset.

## 5 Data availability

The SNV and CNA data for the TCGA cohort are available from the GDC data portal https://portal.gdc.cancer.gov. For SNV, we used the query `cases.project.program.name in [“TCGA“] and files.data type in [“Aggregated Somatic Mutation”, “Masked Somatic Mutation“] and files.experimental_strategy in [“WXS“]` to obtain the list of files to download from GDC. We provide that list in the file `external_data/gdc_manifest.201` of [74], which can be used directly with the GDC download tools. In order to download the restricted mutation files, which are considered potentially identifying, researchers will need to apply to the TCGA Data Access Committee (DAC) via dbGaP (https://dbgap.ncbi.nlm.nih.gov). Regarding CNA, ASCAT results were only available on the hg19 genome alignment version, so we provide the raw CNV and purity data, as well as the final file converted to GRCh38 (see Methods) in the supplementary Source Data file to exactly reproduce our results. Alternatively, the pipeline was recently re-run using the GRCh38 genome alignement, and the data can be downloaded from GDC with the query `files.analysis.workflow type in [“ASCAT2“] and files.data type in [“Allele-specific Copy Number Segment“]`.

For the PCAWG cohort, the mutations are available from the ICGC data portal at https://dcc.icgc.org/releases/PCAWG/consensus_snv_indel, and the CNV data are available at https://dcc.icgc.org/releases/PCAWG/consensus_cnv. A comprehensive list of the files used is presented in Table 1, as well as their accession IDs on the Synapse platform (https://www.synapse.org), where the data is mirrored. To access information that could potentially identify participants, researchers need to apply to the ICGC Data Access Compliance Office (DACO; http://icgc.org/daco) before downloading the PCAWG data.

**Table 1:**
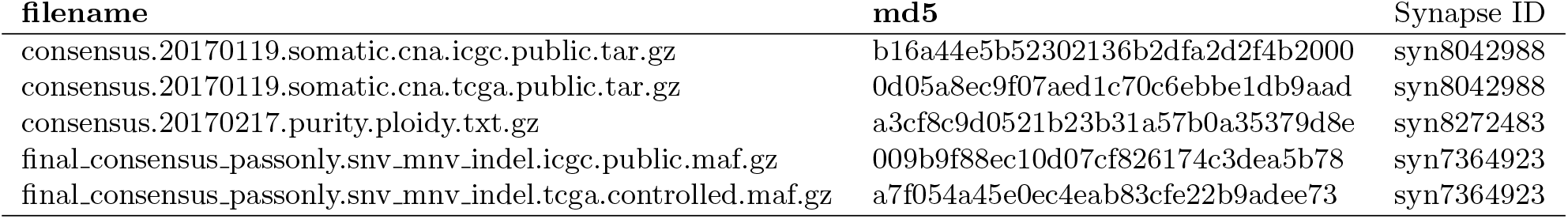
PCAWG files downloaded from the ICGC data portal

The signature matrices used in this work are provided in the supplementary Source Data file, and in the folder external data of the companion github repository [74]. The preprocessed files for the DREAM simulated cohort [21] are publicly available at https://www.synapse.org/#!Synapse:syn2813581/files/, and the PhylogicSim500 and SimClone1000 [22] files are publicly available at https://data.mendeley.com/datasets/by4gbgr9gd/1.

## 6 Code availability

CloneSig is available as a Python package at https://github.com/judithabk6/clonesig [75]. All the code for the analyses performed in this article, aggregated results from simulated and real data, and code to reproduce all figures is available at https://github.com/judithabk6/Clonesig_analysis [74].

## 7 Acknowledgments

The authors thank Stefan Dentro and Adriana Salcedo for their precious help in reusing gold-standard simulated datasets for ITH method evaluation.

## 8 Author Contributions

JPV designed and supervised the project, FR supervised the application to real data and clinical interpretation of the results, JA designed and implemented the method and performed the experiments. JA and JPV wrote the manuscript, with the assistance of FR. All authors read and approved the final manuscript.

