## Supplementary information for "CloneSig can jointly infer intra-tumor heterogeneity and mutational signature activity in bulk tumor sequencing data"

**Supplementary Note 3: Complete overview of TCGA results** **41**

**Supplementary Note 4: Complete overview of PCAWG results** **75**

**Supplementary Figures** **110**

**Supplementary Tables** **117**

### Supplementary Note 1: Supplementary methods

#### 1.1 EM algorithm for parameter estimation

In this section we detail the EM algorithm used to estimate the parameters $\boldsymbol{\theta} = (\xi, \phi, \pi, \rho)$ of CloneSig, for a given number of clones $J$. To lighten notations, we use in this section the notation $M_{max_n} = (C_{tumor}^{major})_n$ for the maximum value that $M_n$ can take. We do not model the distributions of the observed variables $C_n$ (copy number information) and $D_n$ (total read count), and therefore only consider the following complete conditional log-likelihood:

$$\begin{aligned}
\mathcal{L}(\boldsymbol{\theta}) &= \log\left[\prod_{n=1}^{N} \mathbb{P}(B_n, T_n, U_n, S_n, M_n | D_n, C_n; \boldsymbol{\theta}, p, \mu)\right] \\
&= \log\Bigg[\prod_{n=1}^{N}\prod_{u=1}^{J}\prod_{s=1}^{L}\prod_{m=1}^{M_{max_n}} (\mathbb{P}(U_n = u; \boldsymbol{\theta})\mathbb{P}(S_n = s | U_n = u; \boldsymbol{\theta})\mathbb{P}(T_n = t | S_n = s; \mu) \\
&\qquad \mathbb{P}(M_n = m | C_n)\mathbb{P}(B_n | D_n, C_n, M_n = m, U_n = u; \boldsymbol{\theta}, p))^{\mathbb{I}(S_n = s, U_n = u, M_n = m)}\Bigg] \\
&= \sum_{n=1}^{N}\sum_{u=1}^{J}\sum_{s=1}^{L}\sum_{m=1}^{M_{max_n}} \mathbb{I}(S_n = s, U_n = u, M_n = m) \log\left[\xi_u \pi_{us} \mu_{st} M_{max_n}^{-1} \operatorname{BB}(B_n\,;\,D_n, \rho\phi_u\eta_{nm}, \rho(1-\phi_u\eta_{nm}))\right],
\end{aligned} \tag{6}$$

where BB is the beta-binomial density:

$$\operatorname{BB}(k\,;\,n, \alpha, \beta) = \binom{n}{k} \frac{\Gamma(k+\alpha)\Gamma(n-k+\beta)\Gamma(\alpha+\beta)}{\Gamma(n+\alpha+\beta)\Gamma(\alpha)\Gamma(\beta)}, \tag{7}$$

and

$$\eta_{nm} = \frac{pm}{p \times (C_{tumor})_n + (1-p) \times (C_{normal})_n}. \tag{8}$$

To maximize $\mathcal{L}(\boldsymbol{\theta})$, we introduce the auxiliary function $\mathcal{Q}(\boldsymbol{\theta}, \boldsymbol{\theta'})$ as the expected value of the loglikelihood function of $\boldsymbol{\theta}$ when the latent variables follow the law with parameters $\boldsymbol{\theta'}$, that will be alternatively computed and maximized in the two steps of the EM algorithm. For that purpose, let us denote by $\mathbf{X}_n = (C_n, T_n, B_n, D_n)$ the set observed variables for the $n$-th SNV, and $\mathcal{D} = (\mathbf{X}_1, \ldots, \mathbf{X}_N)$ the totality of observed variables. Then we define:

$$\begin{aligned}
\mathcal{Q}(\boldsymbol{\theta}, \boldsymbol{\theta'}) &= \mathbb{E}(\mathcal{L}(\boldsymbol{\theta}) | \mathcal{D}; \boldsymbol{\theta'}, p, \mu) \\
&= \sum_{n=1}^{N}\sum_{u=1}^{J}\sum_{s=1}^{L}\sum_{m=1}^{M_{max_n}} q_{nu} r_{nus} v_{mnu} \log\left[\xi_u \pi_{us} \mu_{st} M_{max_n}^{-1} \operatorname{BB}(B_n\,;\,D_n, \rho\phi_u\eta_{nm}, \rho(1-\phi_u\eta_{nm}))\right],
\end{aligned} \tag{9}$$

with

$$q_{nu} = \mathbb{P}(U_n = u | \mathbf{X}_n; \boldsymbol{\theta'}), \tag{10}$$

$$r_{nus} = \mathbb{P}(S_n = s | U_n = u, \mathbf{X}_n; \boldsymbol{\theta'}), \tag{11}$$

$$v_{mnu} = \mathbb{P}(M_n = m | U_n = u, \mathbf{X}_n; \boldsymbol{\theta'}). \tag{12}$$

The EM algorithm iteratively builds a sequence of estimate $\boldsymbol{\theta^1}, \boldsymbol{\theta^2}, \ldots$ by solving recursively

$$\boldsymbol{\theta^i} = \operatorname{argmax}_{\boldsymbol{\theta}}\, \mathcal{Q}(\boldsymbol{\theta}, \boldsymbol{\theta^{i-1}}). \tag{13}$$

For that purpose, at each iteration $i$, the expectation (E) step first consists in computing the function $\mathcal{Q}(\boldsymbol{\theta}, \boldsymbol{\theta^{i-1}})$ with the current parameters $\boldsymbol{\theta^{i-1}}$. In other words, we must estimate the variables (10)-(12).

Given the conditional independence relationships encoded in the graphical model (Figure 2), one easily gets:

$$q_{nu} = \frac{\sum_{s=1}^{L}\sum_{m=1}^{M_{max_n}} \xi_u^{i-1}\,\mathrm{BB}(B_n|D_n,\rho^{i-1}\phi_u^{i-1}\eta_{nm}^{i-1},\rho^{i-1}(1-\phi_u^{i-1}\eta_{nm}^{i-1}))\mu_{sT_n}\pi_{us}^{i-1}}{\sum_{u'=1}^{J}\sum_{s=1}^{L}\sum_{m=1}^{M_{max_n}} \xi_{u'}^{i-1}\,\mathrm{BB}(B_n|D_n,\rho^{i-1}\phi_{u'}^{i-1}\eta_{nm}^{i-1},\rho^{i-1}(1-\phi_{u'}^{i-1}\eta_{nm}^{i-1}))\mu_{sT_n}\pi_{u's}^{i-1}}\,, \tag{14}$$

$$r_{nus} = \frac{\mu_{sT_n}\pi_{us}^{i-1}}{\sum_{s'=1}^{M}\mu_{s'T_n}\pi_{us'}^{i-1}}\,, \tag{15}$$

$$v_{mnu} = \frac{\sum_{s=1}^{L}\xi_u^{i-1}\,\mathrm{BB}(B_n|D_n,\rho^{i-1}\phi_u^{i-1}\eta_{nm}^{i-1},\rho^{i-1}(1-\phi_u^{i-1}\eta_{nm}^{i-1}))\mu_{sT_n}\pi_{us}^{i-1}}{\sum_{s=1}^{L}\sum_{m'=1}^{M_{max_n}}\xi_u^{i-1}\,\mathrm{BB}(B_n|D_n,\rho^{i-1}\phi_u^{i-1}\eta_{nm'}^{i-1},\rho^{i-1}(1-\phi_u^{i-1}\eta_{nm'}^{i-1}))\mu_{sT_n}\pi_{us}^{i-1}}\,. \tag{16}$$

In the maximization (M) step, we compute $\boldsymbol{\theta^i}$ by plugging the estimates of the E-step (14)-(16) onto (9) and maximizing $\mathcal{Q}(\boldsymbol{\theta},\boldsymbol{\theta^{i-1}})$ separately for each component of $\boldsymbol{\theta}$. The maximization in $\xi$ and $\pi$ are easily obtained as:

$$\forall u \in (1\dots J), \xi_u^i = \sum_{n=1}^{N}\frac{q_{nu}}{N}, \tag{17}$$

$$\forall u \in (1\dots J), \forall s \in (1\dots L), \pi_{us}^i = \frac{\sum_{n=1}^{N} r_{nus}q_{nu}}{\sum_{n'=1}^{N} q_{n'u}}. \tag{18}$$

The optimization of $\phi$ and $\rho$ inside the beta-binomial density term are not computable using a close formula. We therefore resort to numerical optimization and use a projected Newton method, with line search to set the Newton step at each iteration [1], in order to compute approximations of $\phi^i$ and $\rho^i$ that respect constraints on their domain. Indeed, $\rho$ must be non-negative and $\phi$ is a proportion so in the unit interval. For that purpose, we now compute the first and second derivatives of $\mathcal{Q}$ with respect to $\phi$ and $\tau = 1/\rho$:

$$\begin{aligned}\mathcal{Q}(\boldsymbol{\theta},\boldsymbol{\theta^{i-1}}) = \sum_{n=1}^{N}\sum_{u=1}^{J}\sum_{s=1}^{L}\sum_{m=1}^{M_{max_n}} r_{nus}q_{nu}v_{mnu}\Bigg[&\log(\xi_u\mu_{st}\pi_{us}M_{max_n}^{-1}) + \log(\binom{d_n}{b_n})\\ &+\log(\Gamma(b_n+\frac{\phi_u\eta_{nm}}{\tau})) + \log(\Gamma(\frac{1-\phi_u\eta_{nm}}{\tau}+d_n-b_n)) + \log(\Gamma(\frac{1}{\tau}))\\ &-\log(\Gamma(\frac{1}{\tau}+d_n)) - \log(\Gamma(\frac{\phi_u\eta_{nm}}{\tau})) - \log(\Gamma(\frac{1-\phi_u\eta_{nm}}{\tau}))\Bigg]\end{aligned} \tag{19}$$

Let's now compute derivatives. $\psi_0$ and $\psi_1$ denote the digamma and trigamma functions respectively.

$$\frac{\partial \mathcal{Q}(\boldsymbol{\theta}, \boldsymbol{\theta^{i-1}})}{\partial \tau} = \sum_{n=1}^{N}\sum_{u=1}^{J}\sum_{m=1}^{M_{max_n}} \frac{q_{nu}v_{mnu}}{\tau^2}\Big[-\eta_{nm}\phi_u\psi_0(b_n + \frac{\phi_u\eta_{nm}}{\tau}) - (1-\eta_{nm}\phi_u)\psi_0(\frac{1-\phi_u\eta_{nm}}{\tau} + d_n - b_n)$$
$$-\psi_0(\frac{1}{\tau}) + \psi_0(\frac{1}{\tau} + d_n) + \eta_{nm}\phi_u\ \psi_0(\frac{\eta_{nm}\phi_u}{\tau}) + (1-\eta_{nm}\phi_u)\ \psi_0(\frac{1-\eta_{nm}\phi_u}{\tau})\Big] \tag{20}$$

$$\frac{\partial^2 \mathcal{Q}(\boldsymbol{\theta}, \boldsymbol{\theta^{i-1}})}{\partial \tau^2} = \sum_{n=1}^{N}\sum_{u=1}^{J}\sum_{m=1}^{M_{max_n}} q_{nu}v_{mnu}\Bigg[\frac{2}{\tau^3}\bigg(\eta_{nm}\phi_u\psi_0(b_n + \frac{\phi_u\eta_{nm}}{\tau}) + (1-\eta_{nm}\phi_u)\psi_0(\frac{1-\phi_u\eta_{nm}}{\tau} + d_n - b_n)$$
$$+\psi_0(\frac{1}{\tau}) - \psi_0(\frac{1}{\tau} + d_n) - \eta_{nm}\phi_u\ \psi_0(\frac{\eta_{nm}\phi_u}{\tau}) - (1-\eta_{nm}\phi_u)\ \psi_0(\frac{1-\eta_{nm}\phi_u}{\tau})\bigg)$$
$$+\frac{1}{\tau^4}\bigg(\eta_{nm}2\phi_u2\psi_1(b_n + \frac{\phi_u\eta_{nm}}{\tau}) + (1-\eta_{nm}\phi_u)2\psi_1(\frac{1-\phi_u\eta_{nm}}{\tau} + d_n - b_n) + \psi_1(\frac{1}{\tau})$$
$$-\psi_1(\frac{1}{\tau} + d_n) - \eta_{nm}2\phi_u2\ \psi_1(\frac{\eta_{nm}\phi_u}{\tau}) - (1-\eta_{nm}\phi_u)2\ \psi_1(\frac{1-\eta_{nm}\phi_u}{\tau})\bigg)\Bigg] \tag{21}$$

$$\frac{\partial \mathcal{Q}(\boldsymbol{\theta}, \boldsymbol{\theta^{i-1}})}{\partial \phi_u} = \sum_{n=1}^{N}\sum_{m=1}^{M_{max_n}} q_{nu}v_{mnu}\frac{\eta_{nm}}{\tau}\Big[\psi_0(b_n + \frac{\phi_u\eta_{nm}}{\tau}) - \psi_0(\frac{1-\phi_u\eta_{nm}}{\tau} + d_n - b_n)$$
$$-\ \psi_0(\frac{\eta_{nm}\phi_u}{\tau}) +\ \psi_0(\frac{1-\eta_{nm}\phi_u}{\tau})\Big] \tag{22}$$

$$\frac{\partial^2 \mathcal{Q}(\boldsymbol{\theta}, \boldsymbol{\theta^{i-1}})}{\partial \phi_u^2} = \sum_{n=1}^{N}\sum_{m=1}^{M_{max_n}} q_{nu}v_{mnu}\frac{\eta_{nm}^2}{\tau^2}\Big[\psi_1(b_n + \frac{\phi_u\eta_{nm}}{\tau}) + \psi_1(\frac{1-\phi_u\eta_{nm}}{\tau} + d_n - b_n)$$
$$-\ \psi_1(\frac{\eta_{nm}\phi_u}{\tau}) -\ \psi_1(\frac{1-\eta_{nm}\phi_u}{\tau})\Big] \tag{23}$$

$$\frac{\partial^2 \mathcal{Q}(\boldsymbol{\theta}, \boldsymbol{\theta^{i-1}})}{\partial \phi_u \partial \tau} = \sum_{n=1}^{N}\sum_{m=1}^{M_{max_n}} q_{nu}v_{mnu}\frac{\eta_{nm}}{\tau^2}\Big[-\psi_0(b_n + \frac{\phi_u\eta_{nm}}{\tau}) - \frac{\eta_{nm}\ \phi_u}{\tau}\psi_1(b_n + \frac{\phi_u\eta_{nm}}{\tau}) + \psi_0(\frac{1-\phi_u\eta_{nm}}{\tau} + d_n - b_n)$$
$$+\frac{(1-\eta_{nm}\phi_u)}{\tau}\psi_1(\frac{1-\phi_u\eta_{nm}}{\tau} + d_n - b_n) + \psi_0(\frac{\eta_{nm}\phi_u}{\tau}) + \frac{\phi_u\eta_{nm}}{\tau}\psi_1(\frac{\eta_{nm}\phi_u}{\tau})$$
$$-\psi_0(\frac{1-\eta_{nm}\phi_u}{\tau}) - \frac{1-\phi_u\eta_{nm}}{\tau}\psi_1(\frac{1-\eta_{nm}\phi_u}{\tau})\Big] \tag{24}$$

$$\frac{\partial^2 \mathcal{Q}(\boldsymbol{\theta}, \boldsymbol{\theta^{i-1}})}{\partial \phi_u \partial \phi_{u'}} = 0 \tag{25}$$

For the sake of completeness, we provide below a second, equivalent computation using another formulation following [2].

$$
\begin{aligned}
\mathcal{Q}(\boldsymbol{\theta},\boldsymbol{\theta^{i-1}}) &= \sum_{n=1}^{N}\sum_{u=1}^{J}\sum_{s=1}^{L}\sum_{m=1}^{M_{max_n}} r_{nus}q_{nu}v_{mnu} \\
&\log\left[\xi_u\mu_{st}\pi_{us}M_{max_n}^{-1}\binom{d_n}{b_n}\frac{\Gamma(b_n+\rho\phi_u\eta_{nm})\Gamma(\rho(1-\phi_u\eta_{nm})+d_n-b_n)}{\Gamma(\rho+d_n)}\frac{\Gamma(\rho)}{\Gamma(\rho\phi_u\eta_{nm})\Gamma(\rho(1-\phi_u\eta_{nm})}\right] \\
&= \sum_{n=1}^{N}\sum_{u=1}^{J}\sum_{s=1}^{L}\sum_{m=1}^{M_{max_n}} r_{nus}q_{nu}v_{mnu} \\
&\log\left[\xi_u\mu_{st}\pi_{us}M_{max_n}^{-1}\binom{d_n}{b_n}\frac{\prod_{i=0}^{b_n-1}(\phi_u\eta_{nm}+\frac{i}{\rho})\prod_{i=0}^{d_n-b_n-1}(1-\phi_u\eta_{nm}+\frac{i}{\rho})}{\prod_{i=0}^{d_n-1}(1+\frac{i}{\rho})}\right] \\
&= \sum_{n=1}^{N}\sum_{u=1}^{J}\sum_{s=1}^{L}\sum_{m=1}^{M_{max_n}} r_{nus}q_{nu}v_{mnu}\left[\log(\xi_u\mu_{st}\pi_{us}M_{max_n}^{-1})+\log(\binom{d_n}{b_n})+\sum_{i=0}^{b_n-1}\left[\log(\phi_u\eta_{nm}+\frac{i}{\rho})\right]\right. \\
&\left.+\sum_{i=0}^{d_n-b_n-1}\left[\log(1-\phi_u\eta_{nm}+\frac{i}{\rho})\right]-\sum_{i=0}^{d_n-1}\left[\log(1+\frac{i}{\rho})\right]\right] \qquad (26)
\end{aligned}
$$

Let's set $\tau=\frac{1}{\rho}$. We are trying to compute maximum likelihood estimates for $\phi_u$ and $\tau$.

$$\frac{\partial\mathcal{Q}(\boldsymbol{\theta},\boldsymbol{\theta^{i-1}})}{\partial\tau} = \sum_{n=1}^{N}\sum_{u=1}^{J}\sum_{m=1}^{M_{max_n}} q_{nu}v_{mnu}\left[\sum_{i=0}^{b_n-1}\left[\frac{i}{\phi_u\eta_{nm}+i\tau}\right]+\sum_{i=0}^{d_n-b_n-1}\left[\frac{i}{1-\phi_u\eta_{nm}+i\tau}\right]-\sum_{i=0}^{d_n-1}\left[\frac{i}{1+i\tau}\right]\right] \qquad (27)$$

$$\frac{\partial^2\mathcal{Q}(\boldsymbol{\theta},\boldsymbol{\theta^{i-1}})}{\partial\tau^2} = \sum_{n=1}^{N}\sum_{u=1}^{J}\sum_{m=1}^{M_{max_n}} q_{nu}v_{mnu}\left[-\sum_{i=0}^{b_n-1}\left[\frac{i^2}{(\phi_u\eta_{nm}+i\tau)}\right]-\sum_{i=0}^{d_n-b_n-1}\left[\frac{i^2}{(1-\phi_u\eta_{nm}+\tau)^2}\right]+\sum_{i=0}^{d_n-1}\left[\frac{i^2}{(1+\tau)^2}\right]\right] \qquad (28)$$

$$\frac{\partial^2\mathcal{Q}(\boldsymbol{\theta},\boldsymbol{\theta^{i-1}})}{\partial\phi_u\partial\tau} = \sum_{n=1}^{N}\sum_{m=1}^{M_{max_n}} q_{nu}v_{mnu}\left[-\sum_{i=0}^{b_n-1}\left[\frac{i\eta_{nm}}{(\phi_u\eta_{nm}+i\tau)^2}\right]+\sum_{i=0}^{d_n-b_n-1}\left[\frac{i\eta_{nm}}{(1-\phi_u\eta_{nm}+i\tau)^2}\right]\right] \qquad (29)$$

$$\frac{\partial\mathcal{Q}(\boldsymbol{\theta},\boldsymbol{\theta^{i-1}})}{\partial\phi_u} = \sum_{n=1}^{N}\sum_{m=1}^{M_{max_n}} q_{nu}v_{mnu}\left[\sum_{i=0}^{b_n-1}\left[\frac{\eta_{nm}}{\phi_u\eta_{nm}+i\tau}\right]+\sum_{i=0}^{d_n-b_n-1}\left[\frac{-\eta_{nm}}{1-\phi_u\eta_{nm}+i\tau}\right]\right] \qquad (30)$$

$$\frac{\partial^2\mathcal{Q}(\boldsymbol{\theta},\boldsymbol{\theta^{i-1}})}{\partial\phi_u^2} = \sum_{n=1}^{N}\sum_{m=1}^{M_{max_n}} q_{nu}v_{mnu}\left[-\sum_{i=0}^{b_n-1}\left[\frac{\eta_{nm}}{\phi_u\eta_{nm}+i\tau}\right]^2-\sum_{i=0}^{d_n-b_n-1}\left[\frac{\eta_{nm}}{1-\phi_u\eta_{nm}+i\tau}\right]^2\right] \qquad (31)$$

$$\frac{\partial^2\mathcal{Q}(\boldsymbol{\theta},\boldsymbol{\theta^{i-1}})}{\partial\phi_{u'}\partial\phi_u} = 0 \qquad (32)$$

We can then plug these formulas in the projected Newton algorithm to estimate $\phi^i$ and $\rho^i$. We repeat the E and M steps until $\|\boldsymbol{\theta^i}-\boldsymbol{\theta^{i-1}}\| < 10^{-5}\times J\times L$.

#### 1.2 Selecting the number of clones

As explained in Supplementary Section 1.1, the EM algorithm allows us to optimize all parameters of the CloneSig model for a given number of clones $J$. Here we explain how to estimate $J$. A first idea to automatize that choice is to rely on a model selection heuristics, such as the widely used Bayesian Information Criterion (BIC) [3], an asymptotic Bayesian criterion aiming at selecting the model best supported by the data. BIC is defined as

$$BIC(J) = \ell(\mathcal{D};\boldsymbol{\theta}_J) - \frac{D_J}{2}\log N, \qquad (33)$$

where $\ell(\mathcal{D};\boldsymbol{\theta}_J)$ is the maximum log-likelihood as estimated by the EM procedure with $J$ clones, and $D_J$ is the degree of freedom of the model; by default, we take it equal to the number of free parameters, namely, $D_J = J * (L - 1 + 2)$ for $J$ clones, where $L$ is the number of signatures. Indeed, for each clone, we have $L - 1$ parameters for the signature proportions ($\pi$), the frequency of the clone ($\phi_u$), and the proportion of the clone $\xi_u$. We have to remove 1 because $\sum_{u=1}^{J} \xi_u = 1$, and add 1 for the overdispersion parameter $\tau$.

On simulations, however, we found that while BIC correctly identifies the number of clones when the number of SNVs is large, it tends to performs poorly when the number of mutations is low (a few hundreds) in which case it quasi systematically selects a single clone. On the other hand, when we observe the variation of the log-likelihood with the number of components $J$ as for example in Supplementary Figure 1, we clearly see an "elbow" for some $J > 1$, suggesting that the information about $J$ is properly captured by CloneSig's likelihood but not by BIC.

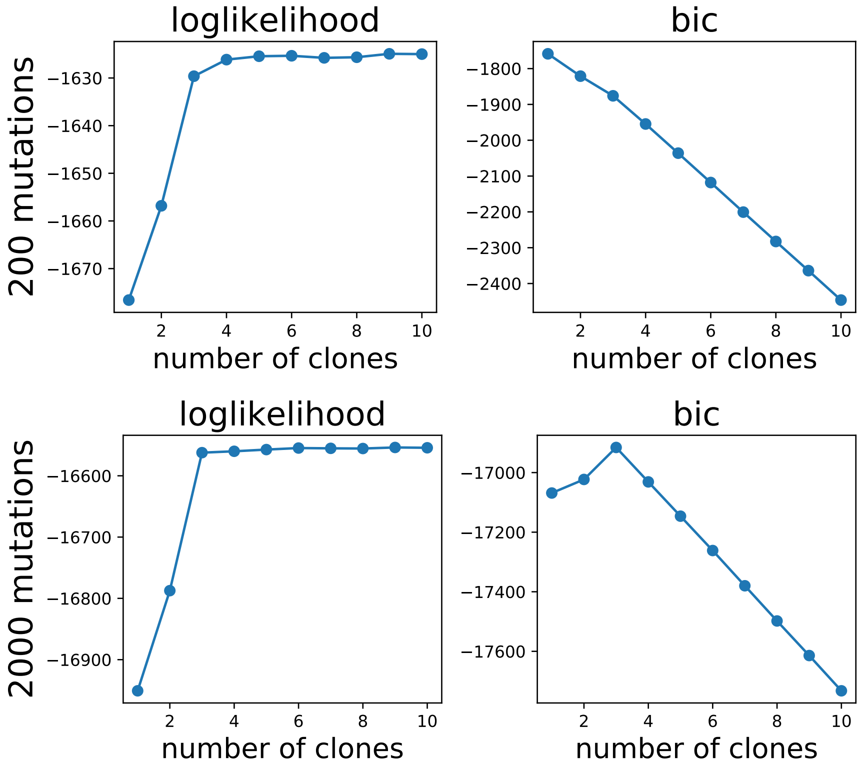

Supplementary Figure 1: **Evolution of the loglikelihood and BIC criterion for 2 simulated samples**, with the same parameters and 200 mutations (up panels), and 2000 mutations (bottom panels). In both cases, the loglikelihood has an "elbow" at 3 clones indicating that the likelihood of the data increases much less at the addition of an additional mixture component beyond 3 components. The BIC criterion is maximal at 3 clones when 2000 mutations are observed, but at 1 clone in the case with 200 mutations.

We observed similar behaviors with other classical criteria such as the Akaike Information Criterion (AIC) [4], the Integrated Classification Likelihood (ICL) [5]), or the slope heuristics as described in [6]. This difficulty can be related to results from statistical theory of model selection and penalization suggesting that asymptotic results are known up to a factor when applied to smaller datasets [7], and therefore propose now as an alternative an empirical criterion that can be fit on data with known model, such as simulations. More precisely, we consider the following criterion:

$$BIC_\alpha(J) = \ell(\mathcal{D};\boldsymbol{\theta}_J) - \alpha D_J \log N\,. \tag{34}$$

with $\alpha > 0$ is a free parameter to be user-defined or estimated, and $D_J$ is a measure complexity of the model.

While we leave $\alpha$ as a user-defined parameter in the CloneSig software, we now propose a systematic approach to estimate it when we can simulate samples. For each simulated sample, we fit CloneSig for 1 to

10 clones. The objective is to estimate a parameter $\alpha$ such that $BIC_{\alpha,J}$ is maximal for the true number of clones $J_{true}$ on all or most simulations. To achieve that, we formulate it as a standard supervised classification problem where for each simulation and each $J \neq J_{true}$, we want $BIC_\alpha(J_{true}) > BIC_\alpha(J)$; since $BIC_\alpha(J)$ is itself a linear function of $\alpha$, we estimate $\alpha$ by minimizing a convex proxy to the number of errors, namely,

$$\min_\alpha \sum_{\mathcal{D}} \sum_{J \neq J_{true}} \phi\left(BIC_\alpha(J_{true}) - BIC_\alpha(J)\right) , \tag{35}$$

where $\phi(u) = \max(0, 1-u)$ is the hinge loss that pushes its argument to be larger than one when minimized; solving (35) is a simple support vector machine (SVM) problem that we solve with a standard SVM solver.

The second important aspect of (34) is $D_J$, that measures the complexity of the model with $J$ clones. The original BIC penalizes the "dimension of the model" [3], that can be interpreted as the degree of freedom of the model, and we now discuss different possible definitions for it. The parameters $\phi$, $\xi$ and $\rho$ determining the CCFs and proportions of the different clones in the mixture must clearly be counted as in BIC. Regarding the signatures however, one can notice that the signatures are neither orthogonal (some signatures are very similar), nor independent (some signatures are associated with the same underlying biological process). Instead of just counting the number of signatures, we therefore propose to estimate the degree of freedom $\text{dof}_L$ of the matrix with $L$ signatures by the number of eigenvalues of the cosine similarity matrix greater than 0.5 in absolute value. As shown in Figure 2, $\text{dof}_L$ is roughly proportional to $L$, at least for $L$ up to 20. Another source of degree of freedom is the copy number. Indeed, for each observed mutation, several values of the number of mutated copies are considered, so if the maximal average multiplicity for mutations in the sample is $M_{max_{avg}}$, a unique clone CCF corresponds in average to $M_{max_{avg}}$ possible VAFs, adding some freedom to the model. We therefore consider four possible definitions for $D_J$, indexed with letters A to D.

$$D_J^A = J \times (L+1) \times M_{max_{avg}} , \tag{36}$$

$$D_J^B = J \times (L+1) , \tag{37}$$

$$D_J^C = J \times (\text{dof}_L + 1) \times M_{max_{avg}} , \tag{38}$$

$$D_J^D = J \times (\text{dof}_L + 1) . \tag{39}$$

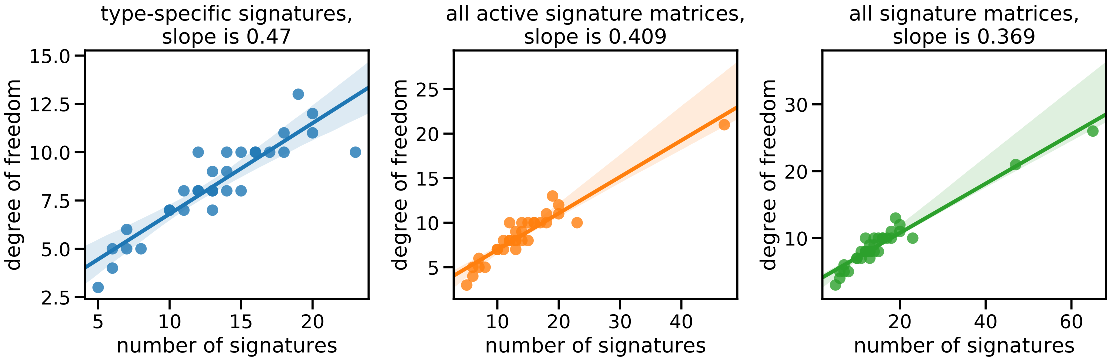

Supplementary Figure 2: **Variation of the degree of freedom for different signature sets** We considered three possible input signature sets: subset of cancer type-specific signatures (31 distinct types illustrated in Supplementary Table 4) or all available signatures depending on the number of signatures (65, or 47 if the potential artefactual signatures are excluded). The left panel shows the dependence for type-specific subsets only, and the middle and right panel for the 47 and 65 signatures additionally. We see that the dependence with the number of signature (slope) is different in the two cases, but that the removal of the 12 artefactual signatures has a moderate impact on the slope compared to the use of only the subsets. The translucent bands around the regression line represent the 95% confidence interval obtained by bootstrap sampling. Source data are provided as a Source Data file.

Moreover, if we consider the variations of the degree of freedom associated with $L$ signatures, $\text{dof}_L$, as a function of $L$ for the 31 available cancer types, and for the all 47 signatures, we note that there is a gap, as the maximal number of signatures in one cancer type is 23, and that the slope seems different for a subset or for all the signatures (see Supplementary Figure 2). The dependency being quite different, this raises the question of whether we should estimate a single $\alpha$ for all situations (i.e., a unique BIC model), or whether we should fit two BIC models: one for the cases where CloneSig is run with only cancer type-specific signatures, and one for the case where CloneSig is run with all the 47 signatures.

For each possible definition of $D_J$ (36)-(39), and for each setting (estimating a unique or two separate BIC models), we ran simulations to estimate the value of $\alpha$ such that $BIC_{\alpha,j}, j \in \{1, \dots, 10\}$ is maximal for the true value of $J$, by solving (35). To evaluate the results, we split the dataset into a train (80% of data) and a test set (20%), and assess the accuracy of $J$ estimation on the test set. To evaluate the stability of the learnt parameter $\alpha$, we compute the 95% confidence interval over 10 independent train-test splits. The values for learnt coefficients, averaged over 10 independent train/test splits for each case are presented in Table 1. The test accuracies for different criteria and different learning settings are presented in Figure 3.

| | $D_J^A$ | $D_J^B$ | $D_J^C$ | $D_J^D$ |
|---|---|---|---|---|
| **separate model (subset)** | $-0.026 \pm 0.000171$ | $-0.043 \pm 0.000200$ | $-0.040 \pm 0.000294$ | $-0.065 \pm 0.000357$ |
| **unique model** | $-0.013 \pm 0.000228$ | $-0.021 \pm 0.000101$ | $-0.026 \pm 0.000160$ | $-0.042 \pm 0.000199$ |
| **separate model (47 signatures)** | $-0.0098 \pm 0.000048$ | $-0.016 \pm 0.000068$ | $-0.021 \pm 0.000109$ | $-0.034 \pm 0.000149$ |

Supplementary Table 1: **Values for the coefficients $\alpha$ for different penalty shapes and training subset.** We see that the coefficients for the whole dataset and for the 47 signatures examples are close. Overall, the confidence interval for the coefficients are small. Source data are provided as a Source Data file.

We first see that, as mentioned earlier, standard model selection criteria (BIC, AIC, ICL) perform overall

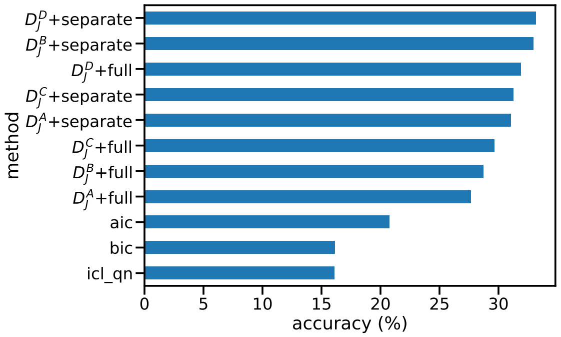

Supplementary Figure 3: **Test accuracy of various model selection criteria.** BIC, AIC and ICL are standard model selection. The others are attempts to learn a valid criterion on simulated data. Source data are provided as a Source Data file.

poorly. Second, we notice that the "separate" strategy is usually slightly better than the "full" strategy, i.e., learning a single $\alpha$ for CloneSig with all 47 signatures or only a subset is not as good as learning two different $\alpha$'s. As for the definition of $D_J$, we see in both cases that using the degree of freedom of the signature matrix is better than counting the number of columns, and that taking into account the variations in copy numbers through $M_{max_{avg}}$ does not bring any benefit. A complete overview of the number of clones found over the test set for each penalization strategy is given in Figure 4. In conclusion, we use in all our experiments an adaptive BIC criterion based on $D_j^D$ as a measure of degree of freedom, and $\alpha$ estimated separately when CloneSig is fitted with 47 signatures or with a cancer-specific subset.

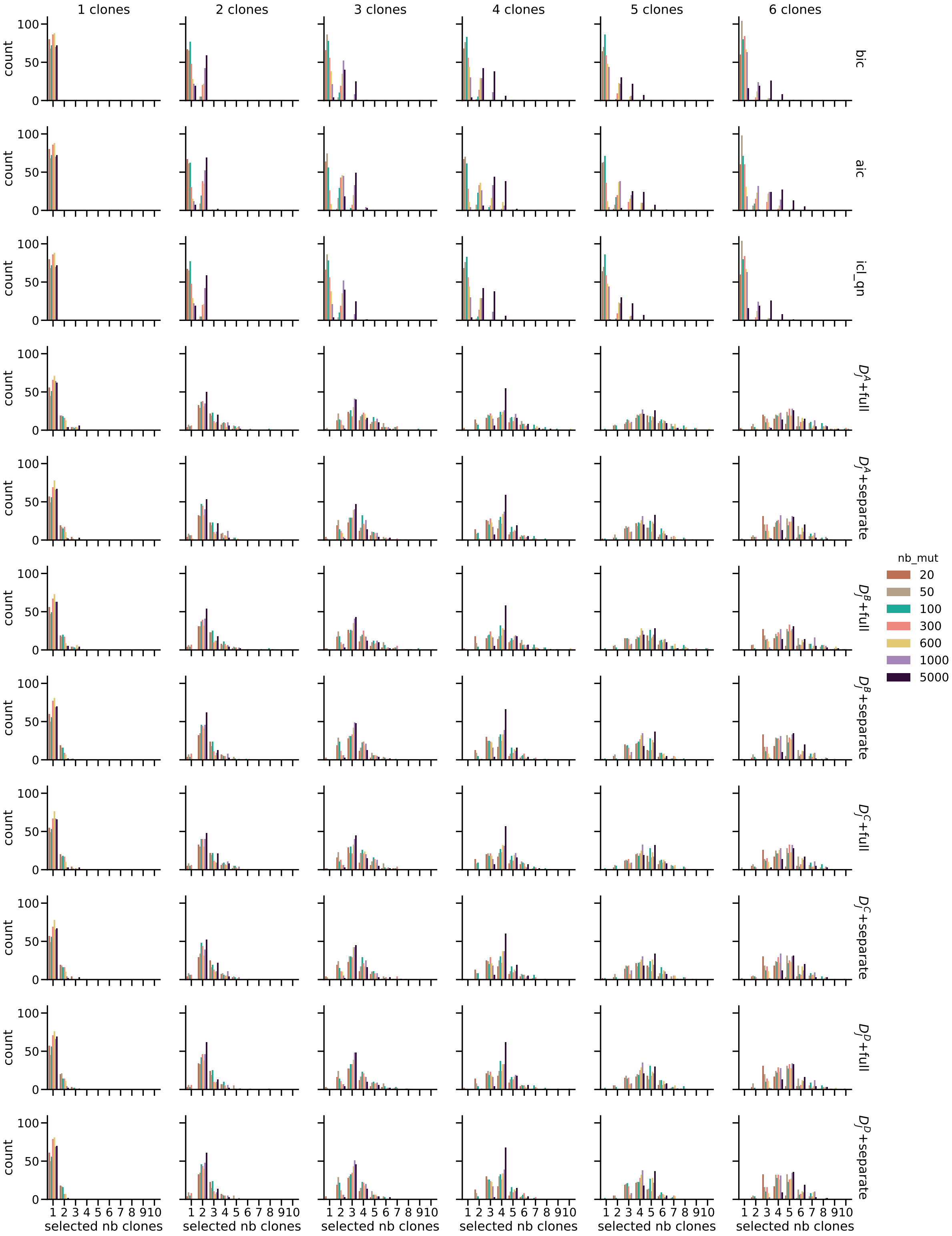

Supplementary Figure 4: **Number of clones found with different model selection criteria on the test set.** The test set was not used to fit the model selection criteria. This illustrates the improved accuracy of the adapted BIC criterion compared to classical criteria. Source data are provided as a Source Data file.

#### 1.3 Statistical test for signature change

To assess whether a signature change between clones is statistically significant, we design and calibrate a statistical test. To that end, we compare the likelihood of a CloneSig model with $J$ clones as determined by the model selection criterion, and the likelihood of a model with the same clones but a single mixture of signatures common to all the clones (and found by fitting all observed mutations together). The objective of the test is to determine whether the difference between the two likelihoods is significant. To that end, we implement a likelihood-ratio test based on the statistics:

$$\lambda = \frac{\ell_{sigCst}}{\ell_{sigChange}} . \tag{40}$$

Following Neyman-Pearson lemma [8] one can set a threshold $c$ to reject the null hypothesis that there is no signature change if $\lambda$ is lower or equal to $c$ with a certain level of significance $\alpha$ determined by the distributions of the likelihood of the model. As this distribution is unknown, we apply the Wilks theorem stating that asymptotically, $-2\log(\lambda)$ follows a chi-squared distribution of parameter the difference in dimensionality between the two alternative models [9].

As previously illustrated for the model selection criterion, the number of parameters is different from the degree of freedom in the case of CloneSig, so we resort to simulations to fit the degree of freedom of the test. We simulate a dataset with a similar mixture of signatures for all clones of each sample, and focused on samples with at least 2 clones, as described in the Methods section. For the purpose of calibration, we use the true number of clones to fit the two alternative models. The objective of this approach is to fit a chi-squared distribution on the empirical distribution of $-2\log(\lambda)$ obtained in simulations. This is achieved again in two settings: fitting with all 47 signatures or with a cancer type-specific subset of signatures. In both cases, the distribution for each number of clones $J$ evokes indeed a chi-squared distribution (Figure 5)

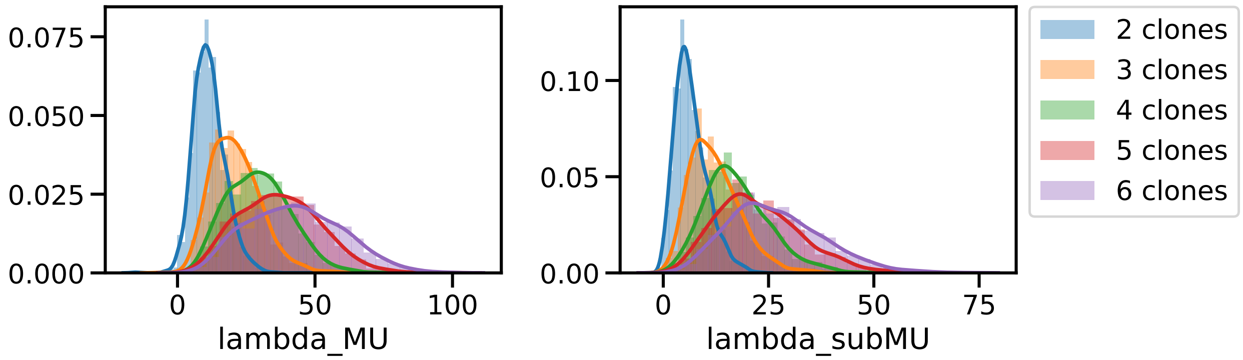

Supplementary Figure 5: **Empirical distribution of** $-2\log(\lambda)$. Practically, $\lambda = \frac{\ell_{sigCst}}{\ell_{sigChange}}$ was obtained by fitting CloneSig with the true number of clones on simulated data, either with all 47 signatures (left), or with a subset of cancer type-specific signatures. The distribution is estimated separately for each number of clones. Source data are provided as a Source Data file.

To fit the degree of freedom to use in the implementation of the test, as the degree of freedom of a chi-squared-distributed variable is its mean, we train a linear ridge regression model to fit $-2\log(\lambda)$ to relevant covariates. Four covariates were initially considered: the number of clones, the degree of freedom of the input signature matrix, the number of mutations, and the diploid proportion of the genome. We found that the last variable has no visible correlation with the target variable (see Supplementary Figure 6). Additionally, when added to the model, with standard scaling of input variables, it has coefficients more than ten times smaller than the ones of the number of clones, the signature degree of freedom, and the number of mutations. We therefore compute the final model on the three retained (unscaled) variables, and we average the values of the coefficients over 10-fold cross-validation. The resulting coefficients are reported in Table 2.

To finally ascertain the validity of the test, we now check the uniform distribution of the p-values for negative samples in Figure 7. There is a slight deviation from the uniform distribution, probably due to the

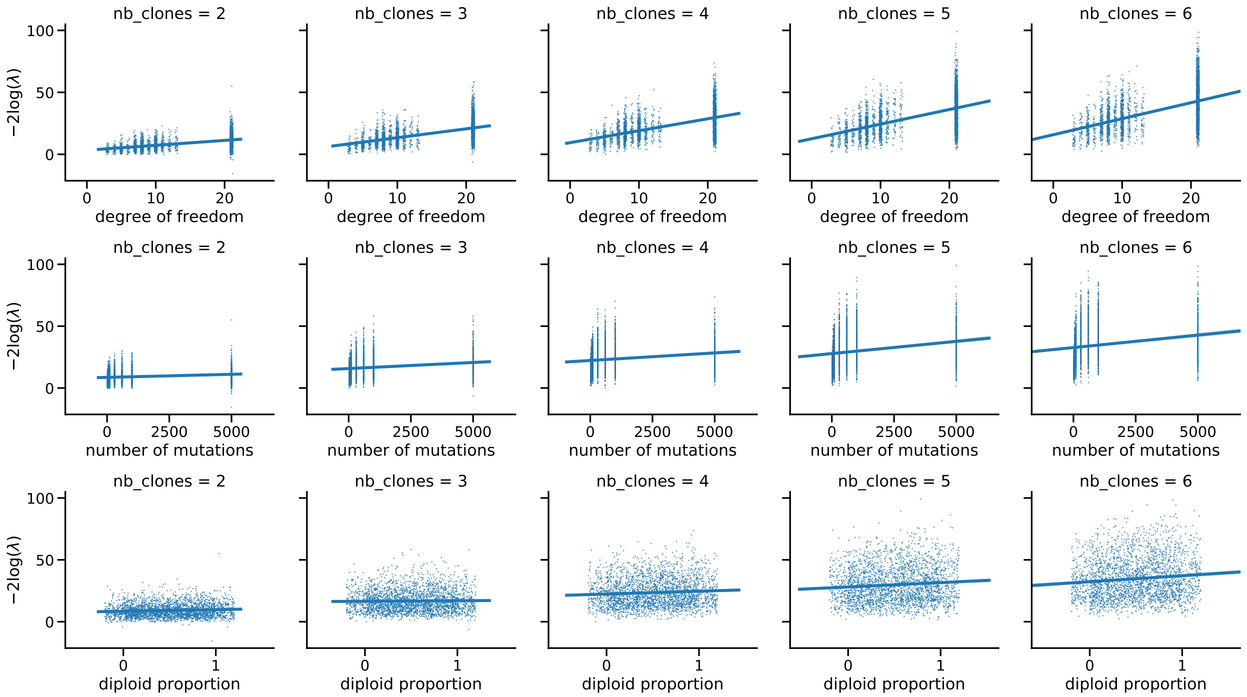

Supplementary Figure 6: **Correlation of $-2\log(\lambda)$, with $\lambda = \frac{\ell_{sigCst}}{\ell_{sigChange}}$ with potentially relevant covariates.** Source data are provided as a Source Data file.

| | Intercept | Number of Clones coefficient | Degree of freedom coefficient | Number of mutations coefficient |
|---|---|---|---|---|
| **separate model (subset)** | $-14.235 \pm 0.0901$ | $5.038 \pm 0.0125$ | $1.233 \pm 0.00916$ | $0.000992 \pm 0.000014$ |
| **unique model** | $-17.620 \pm 0.0577$ | $6.436 \pm 0.0142$ | $0.909 \pm 0.00220$ | $0.00134 \pm 0.000012$ |
| **separate model (47 signatures)** | $-4.606 \pm 0.06111$ | $5.038 \pm 0.0183$ | $0 \pm 0$ | $0.00168 \pm 0.000017$ |

Supplementary Table 2: **Values for the coefficient $\alpha$ for different penalty shapes and training subset.** We see that the coefficients for the whole dataset and for the 47 signatures examples are close. Overall, the confidence interval for the coefficients are small. Source data are provided as a Source Data file.

fact that CloneSig does not necessarily converge to the true model likelihood (and instead to a local maxima), and thus does not respect the conditions of application of Wilks theorem.

We finally explore the sensitivity of the test on the maximum cosine distance between signatures. The dataset used for that purpose consists of 3,600 samples with the number of clones varying between 2 and 6. For each number of clones, we drew 30 distinct $\pi$ matrices with distinct maximal cosine distances between the mutation type profiles. For each number of clones and $\pi$ matrix, we generated a sample with varying number of observed mutations, diploid percent of the genome, and sequencing depth. Figure 8 illustrates the proportion of samples where the test p-value is below 0.05 depending on the maximal distance between two subclones. We observe that detection is more efficient as the distance between clones becomes larger. Dependence on other variables is explored in Supplementary Figure 9. We see in particular the importance of the number of observed mutations, and of the choice of input signatures.

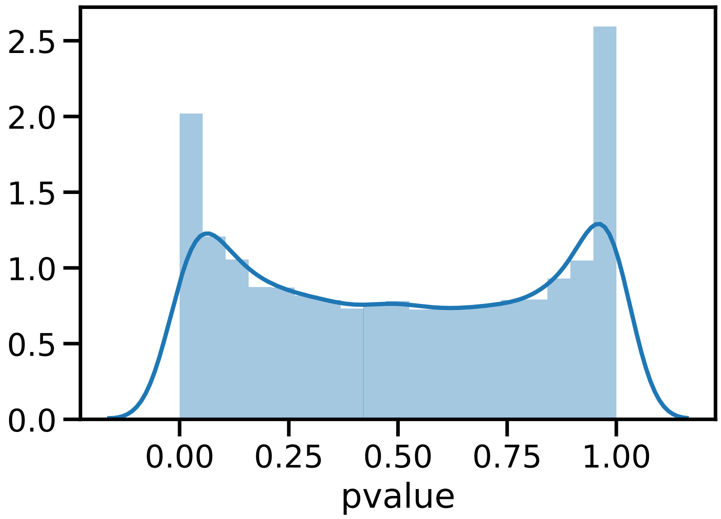

Supplementary Figure 7: **Empirical distribution of the p-values of the calibrated test of significance of signature change for negative simulated samples.** The statistical test presented in 1.3, a likelihood-ratio-based test, to assess a significant change in signature activity was applied to 2,772 simulated samples with no change in signature activity between clones to assess the validity of the test (the null hypothesis). The p-values distribution slightly deviates from the expected uniform distribution. Source data are provided as a Source Data file.

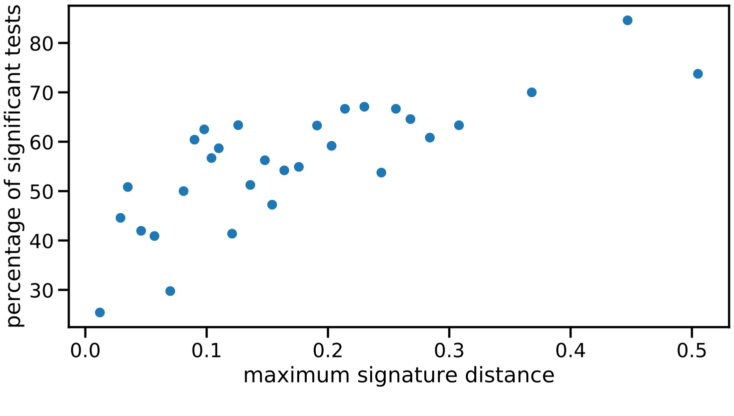

Supplementary Figure 8: **Percentage of significant tests depending on the max distance between 2 clones, quantized in 30 bins.** Source data are provided as a Source Data file.

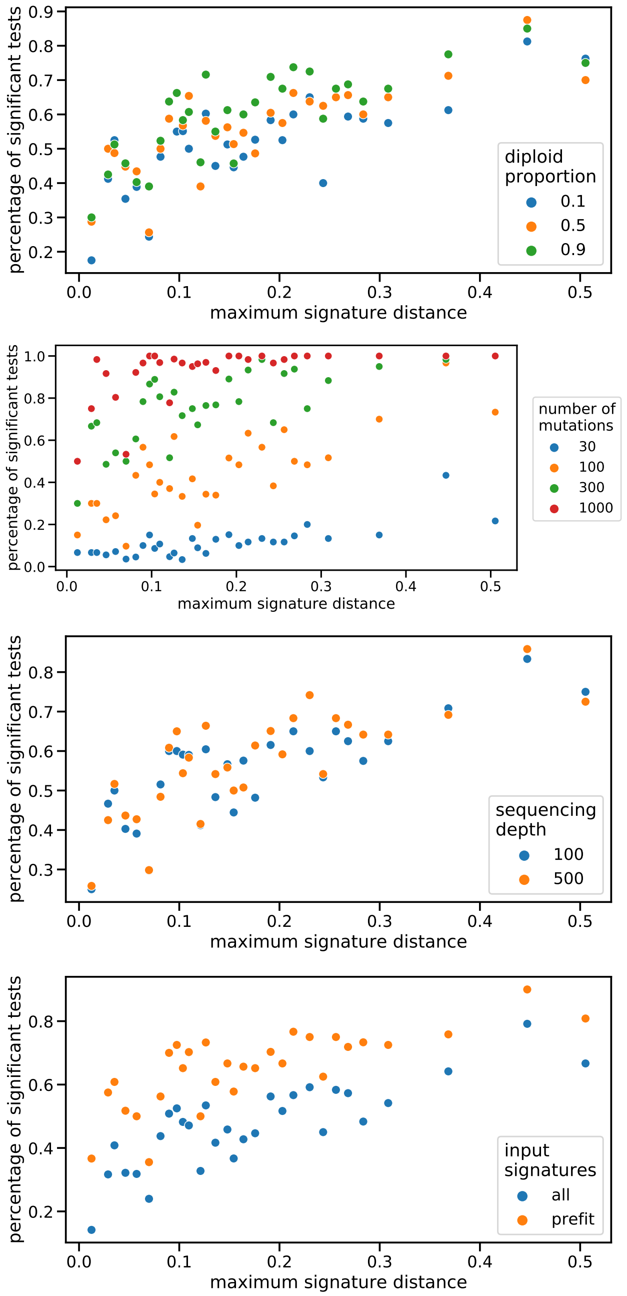

Supplementary Figure 9: **Percentage of significant tests depending several variables.** Considered variables are the number of mutations, and percentage of diploid genome, the sequencing depth, and input signature choice setting. Source data are provided as a Source Data file.

#### 1.4 Several "modes" to run CloneSig

A crucial difficulty in performing mutational signature activity deconvolution is the identifiability of the problem. Indeed, several mixtures of signatures may provide satisfying results. The most common approach to address this issue is to reduce the number of candidate signatures, in particular by using only signatures known to be active in the cancer type of the considered tumor sample [10] (approach `cancer_type`). An alternative approach is to perform two successive fits, the first one on all mutations in the sample in order to select potentially active signatures by keeping those with a contribution greater than a threshold, and the second one to refit those selected signatures with varying number of clones. This avoids the situation where a lot of signatures have very small contributions to the final mixture [11] (approach `prefit`). Those two alternatives are implemented in CloneSig (see Figure 10) and also tested for all methods tested (see supplementary Figures 13-26). For the subclonal reconstruction problem, we see that the two approaches that limit the number of signatures have similar performance and improve the accuracy of CloneSig, especially in cases with few mutations. However, for the signature activity deconvolution problem, even though the `prefit` approach exhibits improved performance compared to taking all signatures, the `cancer_type` approach shows significantly better results. The results were similar for the other signature activity deconvolution methods, so for the rest of the analysis, we retain the `cancer_type` approach, and report only one result per method, to simplify the interpretation.

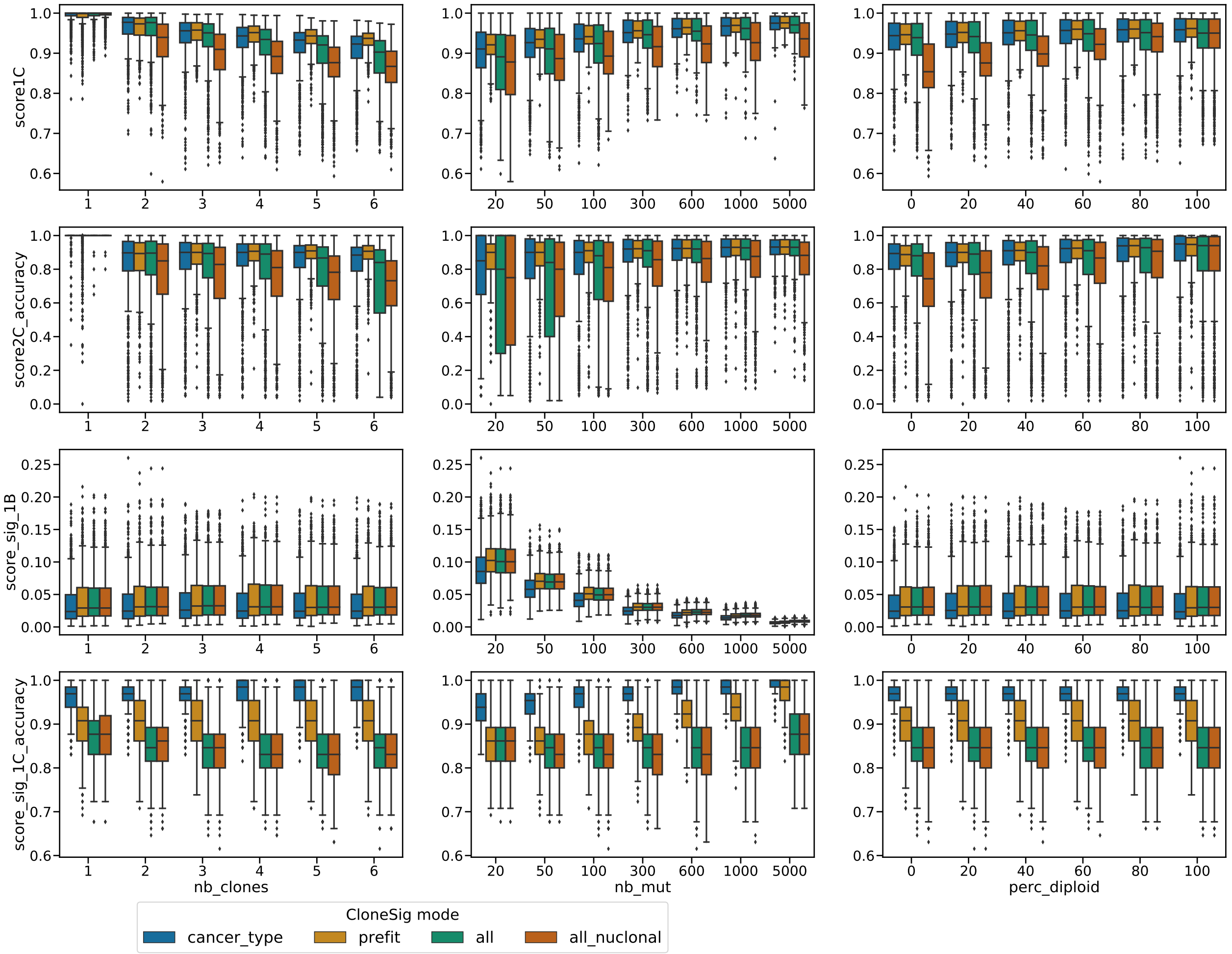

Supplementary Figure 10: **CloneSig's performance for 3 different input signature strategies:** use all available signature (all), a subset of cancer type-specific signatures (cancer_type), or proceed in two steps by first fitting all mutations together to select potential signatures, and then actally run CloneSig with the selected subset (prefit). Additionally, the contribution of CloneSig'a approach for accounting for copy number was evaluated, by implementing the simpler approach from Palimpsest [12] (all_nuclonal). Each boxplot represents the three quartile values of the distribution along with extreme values. The "whiskers" extend to points that lie within 1.5 interquartile ranges of the lower and upper quartile, and then observations that fall outside this range are displayed independently. This means that each value in the boxplot corresponds to an actual observation in the data. Results presented in the figure include respectively 1302, 1299, 1297, 1284, 1267 and 1253 samples with 1 to 6 clones (out of 1302 simulated samples), 1009, 1114, 1116, 1116, 1116, 1116, and 1115 samples with 20, 50, 100, 300, 600, 1000 and 5000 observed mutations (out of 1116 simulated samples), and finally 1285, 1287, 1279, 1283, 1289 and 1979 samples with 0, 20, 40, 60, 80 and 100 percent of the genome being diploid without alteration (out of 1302 simulated samples). Discrepancies between the final results and the initial number of simulated data correspond to unsuccessful runs, for various reasons, including memory limits, encounter of degenerate cases with clones without SNVs. They might complete using a different seed when running CloneSig, but are overall very rare. Source data are provided as a Source Data file.

### Supplementary Note 2: Full benchmarking results

We considered four datasets for the evaluation of CloneSig, and comparison of its performances to other relevant methods. Representative results obtained on the PhylogicSim500 dataset are presented in the main text, and results on other datasets in this note, for brevity.

#### 2.1 CloneSigSim dataset

The CloneSigSim dataset was generated using CloneSig's statistical model, and provides a very thorough analysis of CloneSig and other relevant methods in different situations, representing both WES and WGS data. A total of 10,584 samples were generated, with varying number of mutations, number of clones, percentage of diploid genome, active signatures, and variation of signature activity between clones or not.

Figure 11 summarizes the performance of the different methods according to the different metrics, and under different scenarios, where we vary respectively the number of clones in the simulation (more clones should be more challenging), the number of mutations available (more mutations should help), and the percentage of diploid genome (a higher percentage should be easier).

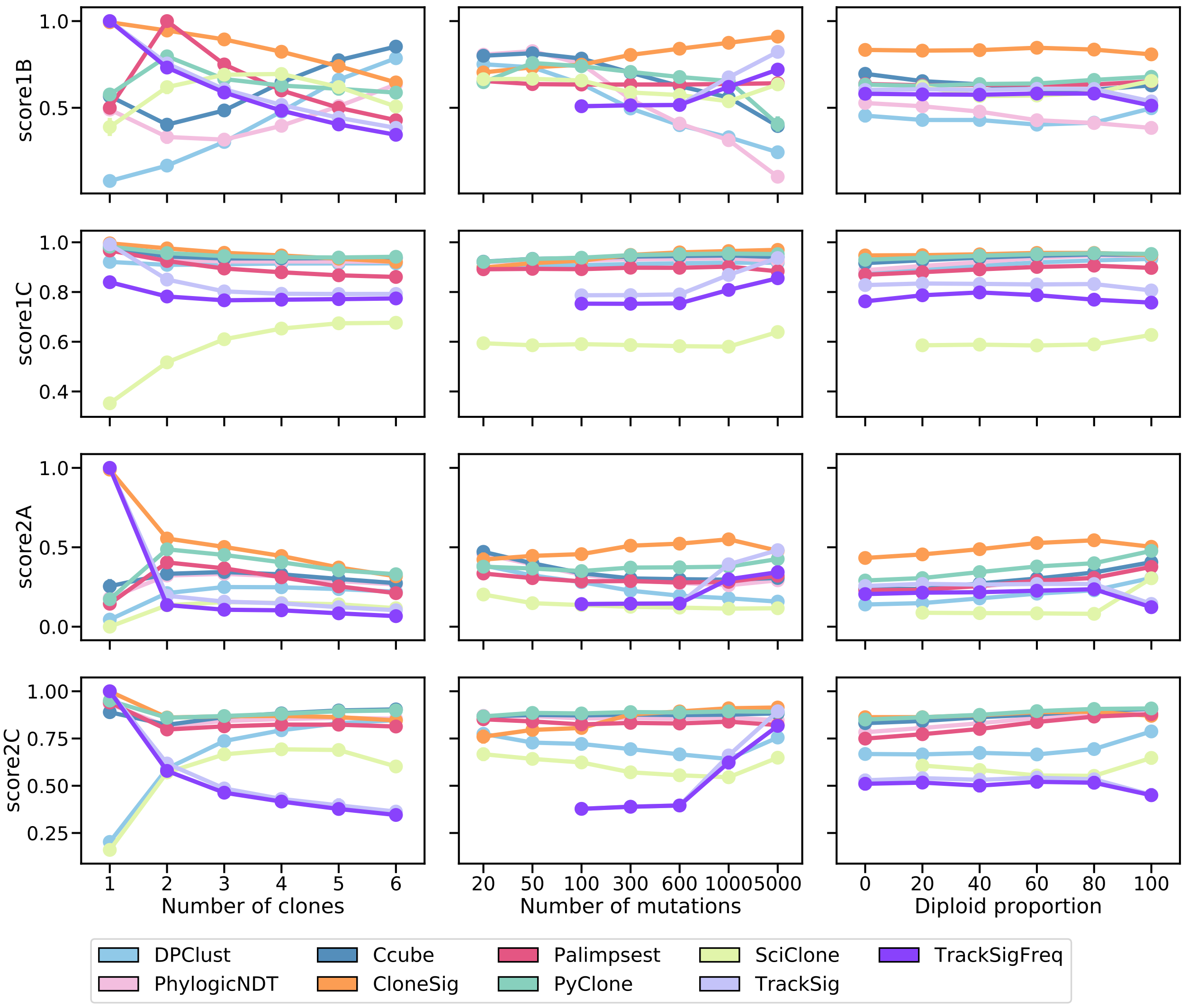

Supplementary Figure 11: **Comparison of CloneSig, TrackSig, TrackSigFreq, Palimpsest, PyClone, SciClone, DPClust, PhylogicNDT and Ccube for subclonal reconstruction on the CloneSigSim dataset.** Each row corresponds to one score, as detailed in the main text. In short, score1B evaluates the number of clones found by the method, score1C the resulting mutation CCF distribution, score2A the co-clustering of mutations in the defined clones, and score2C the classification of subclonal versus clonal mutations. All scores are normalized between 0 and 1, with 1 being the best and 0 the worst. Each column corresponds to a setting where one parameter in the simulation varies: the true number of clones (left), the observed number of mutations (middle), and the diploid proportion of the genome (right). Each point represents the average of the score over all available simulated samples. Bootstrap sampling of the scores was used to compute 95% confidence intervals, which are not visible if smaller than the dot indicating each value in each plot. We have ensured that all scores are comparable by removing samples where at least one method fails for all other methods the corresponding sample sues (such as SciClone and entirely non-diploid genomes, or TrackSig and TrackSigFreq for under 100 mutations). As a result, there are respectively 442, 648, 655, 665, 660 and 659 samples with 1 to 6 clones (out of 1302 simulated samples), 434, 732, 795, 816, 813, 679, and 626 samples with 20, 50, 100, 300, 600, 1000 and 5000 observed mutations (out of 1116 simulated samples), and finally 751, 773, 768, 780, 759 and 649 samples with 0, 20, 40, 60, 80 and 100 percent of the genome being diploid without alteration (out of 1302 simulated samples). Discrepancies between the final results and the initial number of simulated data correspond to unsuccessful runs for at least one of the methods. Source data are provided as a Source Data file.

Regarding the estimation of the number of clones (score1B), CloneSig is the best method in all settings, except in the presence of 5 clones or more, and except for 2 clones where it is outperformed by Palimpsest, which by design systematically predicts two clones. It is in particular the only method achieving a perfect accuracy in identifying samples with one or two clones, and exhibits the best performance for score1B up to 4 clones (with the exception of Palimpsest for two clones). CloneSig, TrackSig and TrackSigFreq see their performance decrease with the number of clones, as expected, while surprisingly Ccube, DPClust and PhylogicNDT have the opposite behavior and achieve better results when the number of clones is large. During the experiments we noticed that PyClone tends to find large numbers of clones with only one mutation, so we ignore these clones when we compute score1B in order not to excessively penalize PyClone for this problematic behavior. PyClone, SciClone and Palimpsest have overall a stable performance with varying numbers of clones. Regarding the impact of the number of mutations on score1B, we see that CloneSig outperforms all other methods when enough mutations are observed. As expected, CloneSig, TrackSig and TrackSigFreq improve when the number of SNV increases, and we confirm that TrackSig requires at least 1,000 SNVs to be competitive with other methods in this experiment, while CloneSig reaches the best performance of TrackSig with as few as 100 SNVs. A surprising result is that for PyClone, SciClone, DPClust, PhylogicNDT and Ccube, score1B decreases with the number of observed mutations, which may suggest a bad calibration of the clone number estimate for large numbers of SNVs; for CloneSig we designed a specific, adaptive estimator for the number of clones since we observed that standard statistical approaches for model selection perform poorly in this setting (see Methods and Supplementary Section 1.2). The percentage of diploid genome has no visible impact on the performance of any method. Regarding score1C, which focuses not on the number of clones estimated but on their ability to correctly recapitulate the distribution of CCF values, we also see that all methods except SciClone, TrackSig and TracksigFreq have almost perfect performance in all settings. TrackSig and TrackSigFreq perform slightly worse, especially as the number of clones increases, but this may be explained by its poor performance when the number of mutations is too low, as performance matches the other methods for 5,000 mutations. Finally, SciClone is clearly the worse method for score1C, particularly with 1 to 3 clones.

Besides the ability of different methods to reconstruct the correct number of subclones and their CCF, as assessed by score1B and score1C, we measure with score2A their ability to correctly assign individual mutations to their clones, an important step for downstream analysis of mutations in each subclone. According to score2A, CloneSig outperforms all other methods in almost all scenarios, illustrating the improved accuracy of accounting for both CCF and mutational signatures when achieving ITH reconstruction. For all methods, score2A decreases when the number of clones increases and when the percentage of diploid genomes decreases, as expected, but the relative order of methods does not change, with CloneSig followed by a group of three methods with similar performances: PyClone, Ccube and Palimpsest. DPClust is slightly behind, and SciClone performs poorly except when the genome is fully diploid, in which case it gets competitive with Palimpsest but still below CloneSig, PyClone and Ccube. The number of mutations has a limited impact on the performance of all methods except for TrackSig and TrackSigFreq, which only becomes competitive after 1,000 mutations. CloneSig with 100 mutations still outperforms TrackSig with 1,000 mutations, though. Finally, when we assess the capacity of each method to simply discriminate clonal from subclonal mutations using score2C, a measure meant not to penalize Palimpsest which only performs that task, we see again that CloneSig is among the best in all scenarios, except under 100 mutations, and very close to by Ccube, PyClone Palimpsest, as well as TrackSig and TrackSigFreq with 5,000 mutations. Palimpsest is a bit below these methods, while SciClone, DPClust, PhylogicNDT TrackSig and TrackSigFreq with 1,000 mutations or less are clearly not competitive for this metric.

Overall, these experiments show that CloneSig performs as well as or better than the state-of-the-art according to all metrics considered and in most simulated scenarios, confirming that accounting for the mutation type for each mutation, in addition to its CCF, improves the accuracy of subclonal reconstruction. We also confirm that TrackSig and TrackSigFreq, the only existing methods that combine CCF and mutational signature information to detect subclones, require at least 1,000 mutations to obtain results competitive with other methods in our benchmark, while CloneSig reaches good accuracy in all scores with as few as 100 mutations. For smaller number of mutations, state-of-the art subclonal reconstruction methods outperform CloneSig, indicating that the signature signal may bring more noise than help at such a small resolution.

In addition to ITH inference in terms of subclones, CloneSig estimates the mutational processes involved in the tumor and in the different subclones. We now assess the accuracy of this estimation on simulated data, using six performance scores detailed in the Methods section. In short, score_sig_1A is the Euclidean distance between the normalized mutation type counts and the reconstructed profile (activity-weighted sum of all signatures); score_sig_1B is the Euclidean distance between the true and the reconstructed profile; score_sig_1C measures the identification of the true signatures; score_sig_1D is the proportion of signatures for which the true causal signature is correctly identified; and score_sig_1E reports the median of the distribution of the cosine distance between the true and the predicted mutation type profile that generated each mutation. We compare CloneSig to the two other methods that perform both ITH and mutational process estimation, namely, TrackSig and Palimpsest, and add also deconstructSigs [13] in the benchmark, a method that optimizes the mixture of mutational signature of a sample through multiple linear regressions without performing subclonal reconstruction.

Figure 12 shows the performance of the different methods according to the different metrics. For Score_sig_1A and Score_sig_1B, all methods exhibit overall similar performances, with a small advantage for CloneSig, TrackSig and TrackSigFreq over Palimpsest and deconstructSigs in several scenarios. For Score_sig_1C, CloneSig, TrackSig and TrackSigFreq exhibit the best AUC to detect present signatures. It may be related to a better sensitivity as CloneSig and TrackSig perform signature activity deconvolution in smaller subsets of mutations. All methods perform similarly with respect to Score_sig_1D. The median cosine distance (Score_sig_1E) is also slightly better for CloneSig than for other methods in all settings.

Overall, as for ITH inference, we conclude that CloneSig is as good as or better than all other methods in all scenarios tested. Further results where we vary other parameters in each methods, notably the set of mutations used as inputs or the set of signatures used as prior knowledge, can be found in Supplementary Figures 13 to 27; they confirm the good performance of CloneSig in all settings tested.

To fully assess CloneSig's performance on the CloneSigSim data, in comparison with other state-of-the-art approaches for subclonal reconstruction and signature activity deconvolution, we report here the full results with all tested "modes" (all signatures, a subset of cancer-type-specific signatures, or a pre-fit step where only the most prominent signatures found on the whole set of mutations are then retained for the true signature activity deconvolution for CloneSig, TrackSig and Palimpsest). In this extensive version of the results, we report all metrics used to create score2C (AUC, specificity, sensitivity), score_sig_1C and score_sig_1E (`max_diff_distrib_mut`, `median_diff_distrib_mut`, `perc_dist_5` and `perc_dist_10`).

Regarding the subclonal reconstruction problem, for all metrics, there is little difference between the different modes of each signature-aware method, except for `score2C_sensitivity` for CloneSig, where the use of the cancer-type-specific subset exhibits better results. For signature activity deconvolution, there is a higher variability of results with respect to the run mode. CloneSig is the best performing method, except for one metric: `max_diff_distrib_mut`. For `Score_sig_1C`, the mode cancer-type-specific subset for CloneSig achieves a very good specificity, but the other modes have a high proportion of false positive signatures.

Additionally, we conduct a similar benchmark in the case where there is no signature change between subclones, and present results in Supplementary Figures 13 to 27, panels b, c, f. The improvement of CloneSig over other methods in subclonal reconstruction is partially lost in this setting, but CloneSig remains competitive, and the best performing method for score 1B up to 3 clones. A similar trend is visible for all scores for the subclonal reconstruction problem, with slightly worse scores, and higher inter-quartile space when there is no signature variation between clones. For the signature activity deconvolution problem, most metrics are unaffected, except for score_sig_1E, where all methods perform better and close the gap with CloneSig. Overall, CloneSig performs better than other methods when there are differences of signature activities between subclones, and remains competitive with other approaches in the absence of signature change.

The runtimes of all methods for those simulations are presented in Figure 27. The main determinant of runtime is the number of input mutations for all methods. CloneSig is slower than methods involving variational inference for the subclonal reconstruction problem, but is significantly faster than PyClone, especially for high numbers of mutations, thus illustrating its scalability to both WES and WGS data.

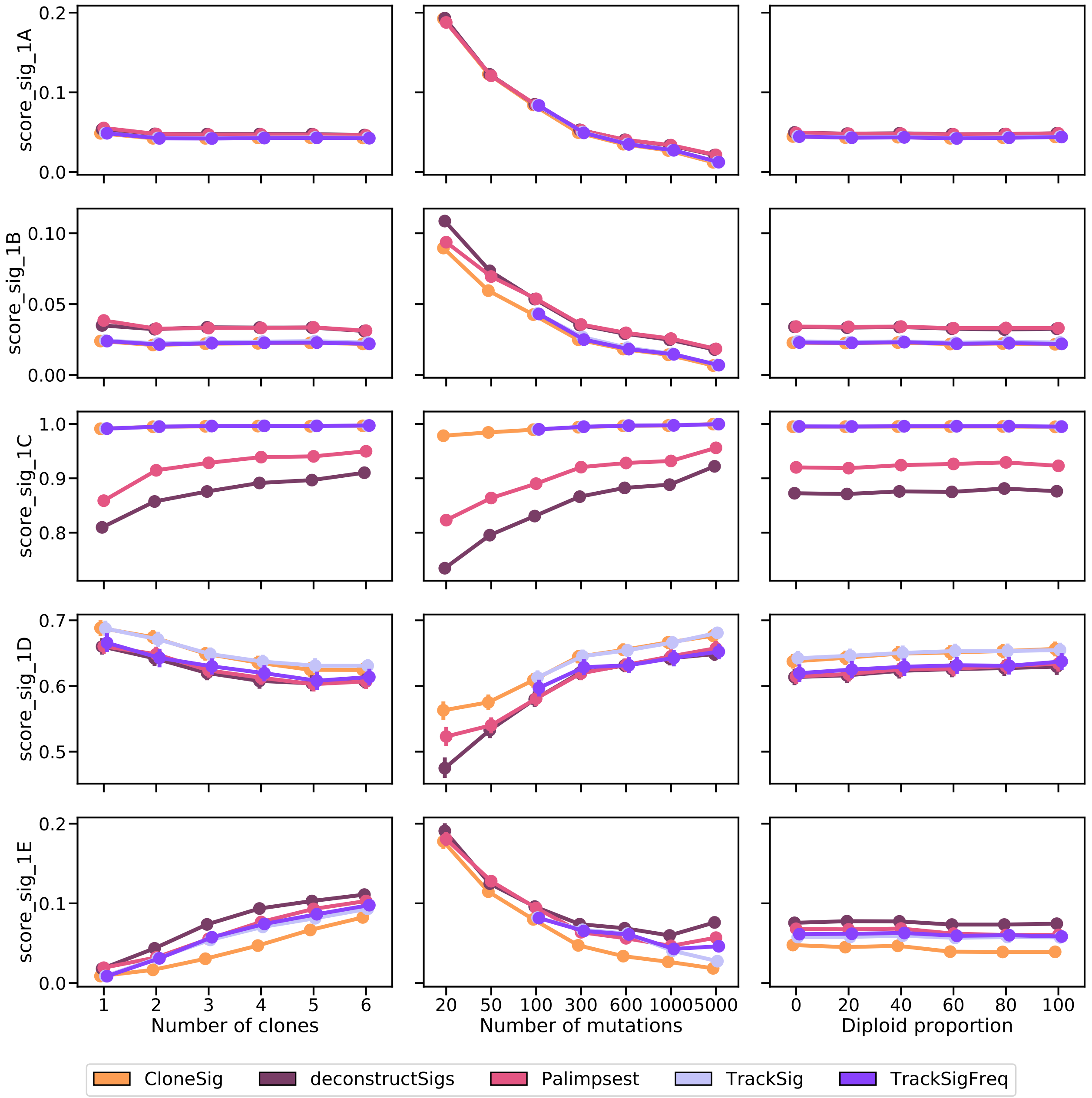

Supplementary Figure 12: **Comparison of CloneSig, TrackSig, TrackSigFreq, Palimpsest, and deconstructSigs for signature activity deconvolution on the CloneSigSim dataset.** Several metrics have been implemented, and are detailed in the main text. Scores_sig_1A and 1B are distances between the estimated mutation type profile (as defined by the signature activity proportions) to the true mutation profile (defined using the parameters used for simulations, 1A), and the empirical observed mutation profile (defined using available observed mutations, 1B), and is better when close to 0. Score_sig_1C is the area under the ROC curve for the classification of signatures as active or inactive in the sample, and is better when close to 1. Score_sig_1D is the proportion of mutations for which the correct signature was attributed, and is better when close to 1. Finally, Score_sig_1E is the median distance to the true mutation type profile of the clone to which a mutation was attributed from the true distribution of its original clone in the simulation, and is better when close to 0. The results are presented depending on several relevant covariates: the true number of clones (left), the number of mutations (middle), and the diploid proportion of the genome (right). Each point represents the average of the score over all available simulated samples. Bootstrap sampling of the scores was used to compute 95% confidence intervals. Results for each setting are presented for samples for which all methods ran successfully; there are respectively 659, 791, 795, 791, 793 and 782 samples with 1 to 6 clones (out of 1302 simulated samples), 861, 995, 992, 1010, 1012, 835, and 762 samples with 20, 50, 100, 300, 600, 1000 and 5000 observed mutations (out of 1116 simulated samples), and finally 755, 781, 769, 782, 759 and 765 samples with 0, 20, 40, 60, 80 and 100 percent of the genome being diploid without alteration (out of 1302 simulated samples). Discrepancies between the final results and the initial number of simulated data correspond to unsuccessful runs for at least one of the methods. Source data are provided as a Source Data file.

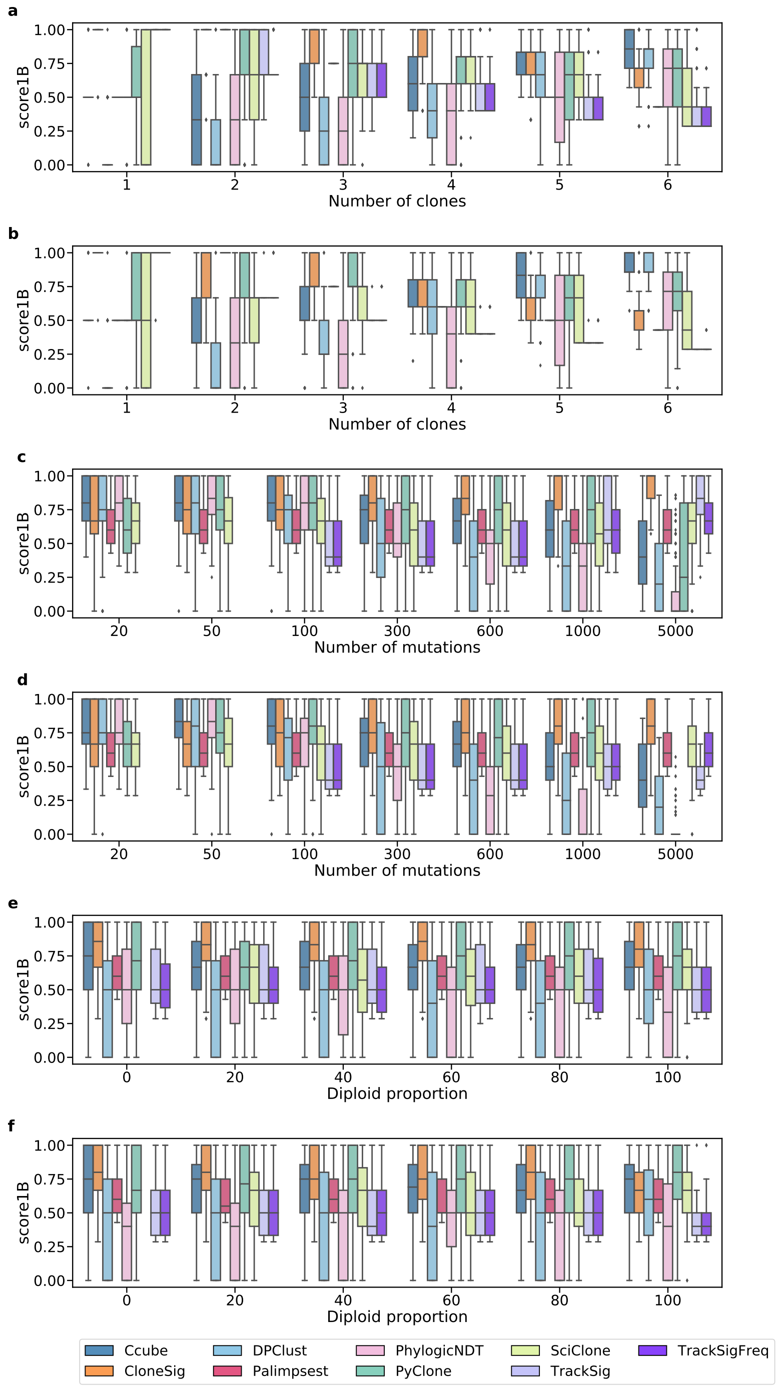

Supplementary Figure 13: **Score_1B for ITH methods on the CloneSigSim dataset**, with varying number of clones (a,b), number of observed mutations (c,d) and diploid percent of the genome (e,f). Panels a, c and e correspond to simulations with varying signature between clones, and b, d, f to simulations with constant signatures. Each boxplot represents the three quartile values of the distribution along with extreme values. The "whiskers" extend to points that lie within 1.5 interquartile ranges of the lower and upper quartile, and then observations that fall outside this range are displayed independently. This means that each value in the boxplot corresponds to an actual observation in the data. We have ensured that all scores are comparable by removing samples where at least one method fails for all other methods the corresponding sample wues (such as SciClone and entirely non-diploid genomes, or TrackSig and TrackSigFreq for under 100 mutations, and PyClone with 5000 mutations in the constant setting). Results for each setting are presented for samples for which all methods ran successfully; there are respectively 442, 648, 655, 665, 660 and 659 samples with 1 to 6 clones (out of 1302 simulated samples) in the varying setting (panel a), and 162, 192, 188, 195, 184 and 182 (out of 420 simulated samples) in the constant setting (panel b), 434, 732, 795, 816, 813, 679, and 626 samples with 20, 50, 100, 300, 600, 1000 and 5000 observed mutations (out of 1116 simulated samples) in the varying setting (panel c), and 153, 265, 289, 290, 287, 237 and 224 (out of 360 simulated samples) in the constant setting (panel d), and finally 751, 773, 768, 780, 759 and 649 samples with 0, 20, 40, 60, 80 and 100 percent of the genome being diploid without alteration (out of 1302 simulated samples) in the varying setting (panel e), and 225, 224, 225, 234, 229 and 191 (out of 420 simulated samples) in the constant setting (panel f). Source data are provided as a Source Data file.

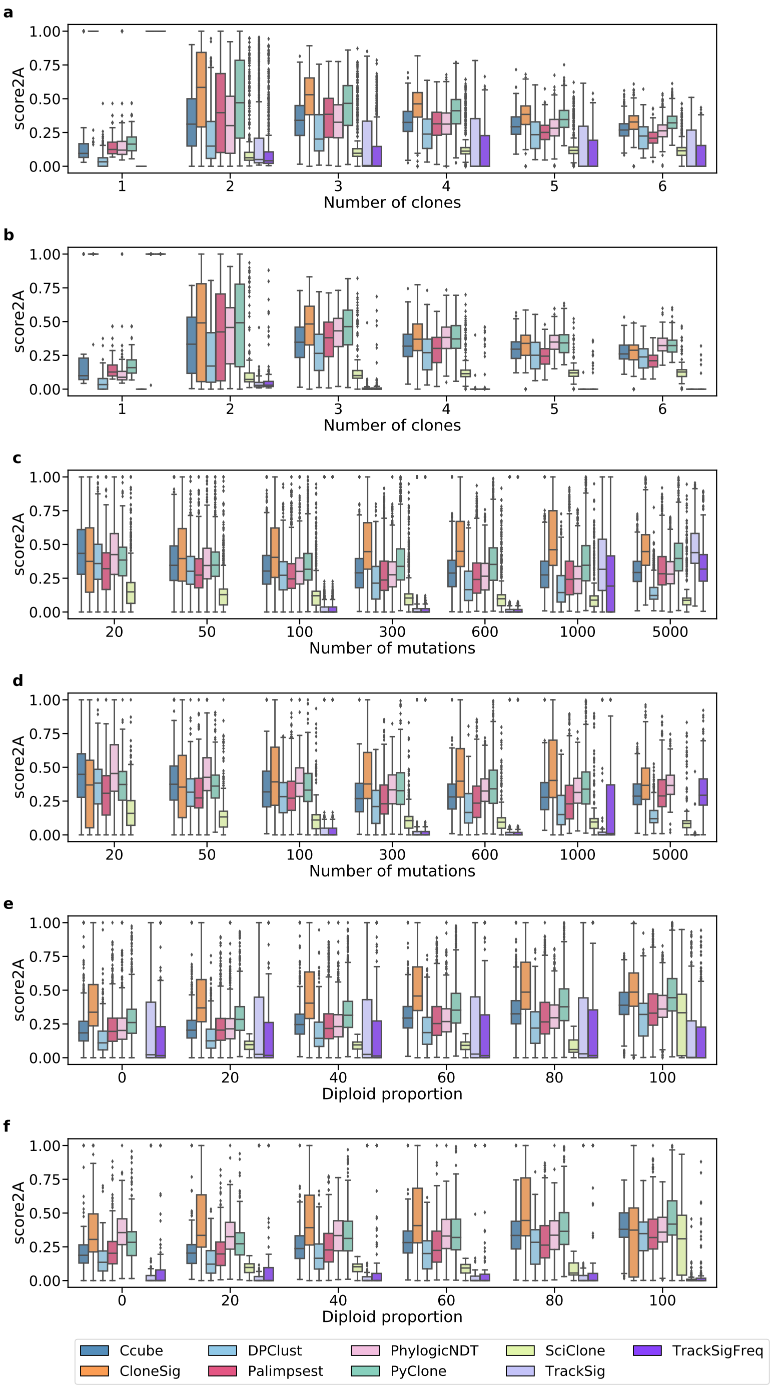

Supplementary Figure 14: **Score_2A for ITH methods on the CloneSigSim dataset**, with varying number of clones (a, b), number of observed mutations (c, d) and diploid percent of the genome (e, f). Panels a, c and e correspond to simulations with varying signature between clones, and b, d, f to simulations with constant signatures. Each boxplot represents the three quartile values of the distribution along with extreme values. The "whiskers" extend to points that lie within 1.5 interquartile ranges of the lower and upper quartile, and then observations that fall outside this range are displayed independently. This means that each value in the boxplot corresponds to an actual observation in the data. We have ensured that all scores are comparable by removing samples where at least one method fails for all other methods the corresponding stic issues (such as SciClone and entirely non-diploid genomes, or TrackSig and TrackSigFreq for under 100 mutations, and PyClone with 5000 mutations in the constant setting). Results for each setting are presented for samples for which all methods ran successfully; there are respectively 442, 648, 655, 665, 660 and 659 samples with 1 to 6 clones (out of 1302 simulated samples) in the varying setting (panel a), and 162, 192, 188, 195, 184 and 182 (out of 420 simulated samples) in the constant setting (panel b), 434, 732, 795, 816, 813, 679, and 626 samples with 20, 50, 100, 300, 600, 1000 and 5000 observed mutations (out of 1116 simulated samples) in the varying setting (panel c), and 153, 265, 289, 290, 287, 237 and 224 (out of 360 simulated samples) in the constant setting (panel d), and finally 751, 773, 768, 780, 759 and 649 samples with 0, 20, 40, 60, 80 and 100 percent of the genome being diploid without alteration (out of 1302 simulated samples) in the varying setting (panel e), and 225, 224, 225, 234, 229 and 191 (out of 420 simulated samples) in the constant setting (panel f). Source data are provided as a Source Data file.

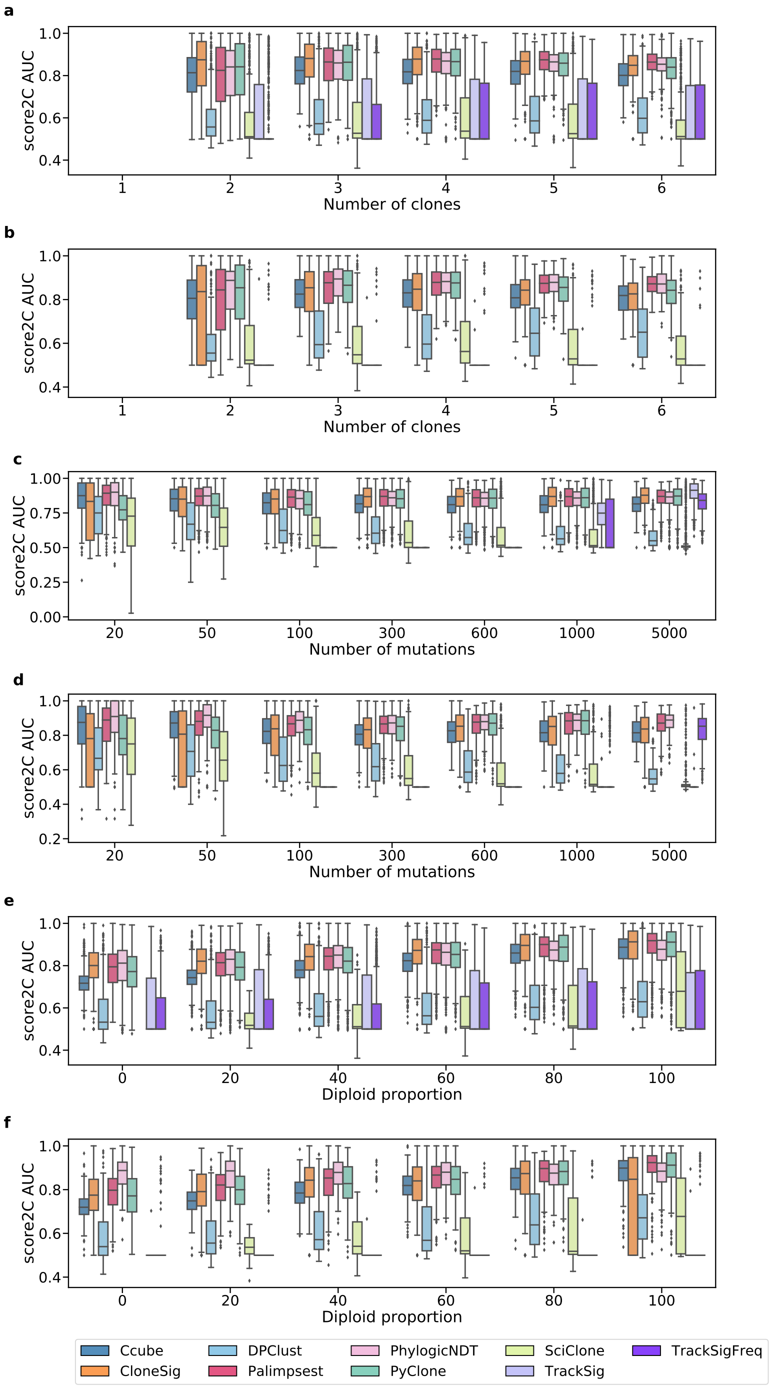

Supplementary Figure 15: **Score_2C (area under the curve) for ITH methods on the CloneSigSim dataset**, with varying number of clones (a, b), number of observed mutations (c, d) and diploid percent of the genome (e, f). Panels a, c and e correspond to simulations with varying signature between clones, and b, d, f to simulations with constant signatures. Each boxplot represents the three quartile values of the distribution along with extreme values. The "whiskers" extend to points that lie within 1.5 interquartile ranges of the lower and upper quartile, and then observations that fall outside this range are displayed independently. This means that each value in the boxplot corresponds to an actual observation in the data. We have ensured that all scores are comparable by removing samples where at least one method fails for all other methods the corresponding sample sues (such as SciClone and entirely non-diploid genomes, or TrackSig and TrackSigFreq for under 100 mutations, and PyClone with 5000 mutations in the constant setting). Results for each setting are presented for samples for which all methods ran successfully; there are respectively 442, 648, 655, 665, 660 and 659 samples with 1 to 6 clones (out of 1302 simulated samples) in the varying setting (panel a), and 162, 192, 188, 195, 184 and 182 (out of 420 simulated samples) in the constant setting (panel b), 434, 732, 795, 816, 813, 679, and 626 samples with 20, 50, 100, 300, 600, 1000 and 5000 observed mutations (out of 1116 simulated samples) in the varying setting (panel c), and 153, 265, 289, 290, 287, 237 and 224 (out of 360 simulated samples) in the constant setting (panel d), and finally 751, 773, 768, 780, 759 and 649 samples with 0, 20, 40, 60, 80 and 100 percent of the genome being diploid without alteration (out of 1302 simulated samples) in the varying setting (panel e), and 225, 224, 225, 234, 229 and 191 (out of 420 simulated samples) in the constant setting (panel f). Source data are provided as a Source Data file.

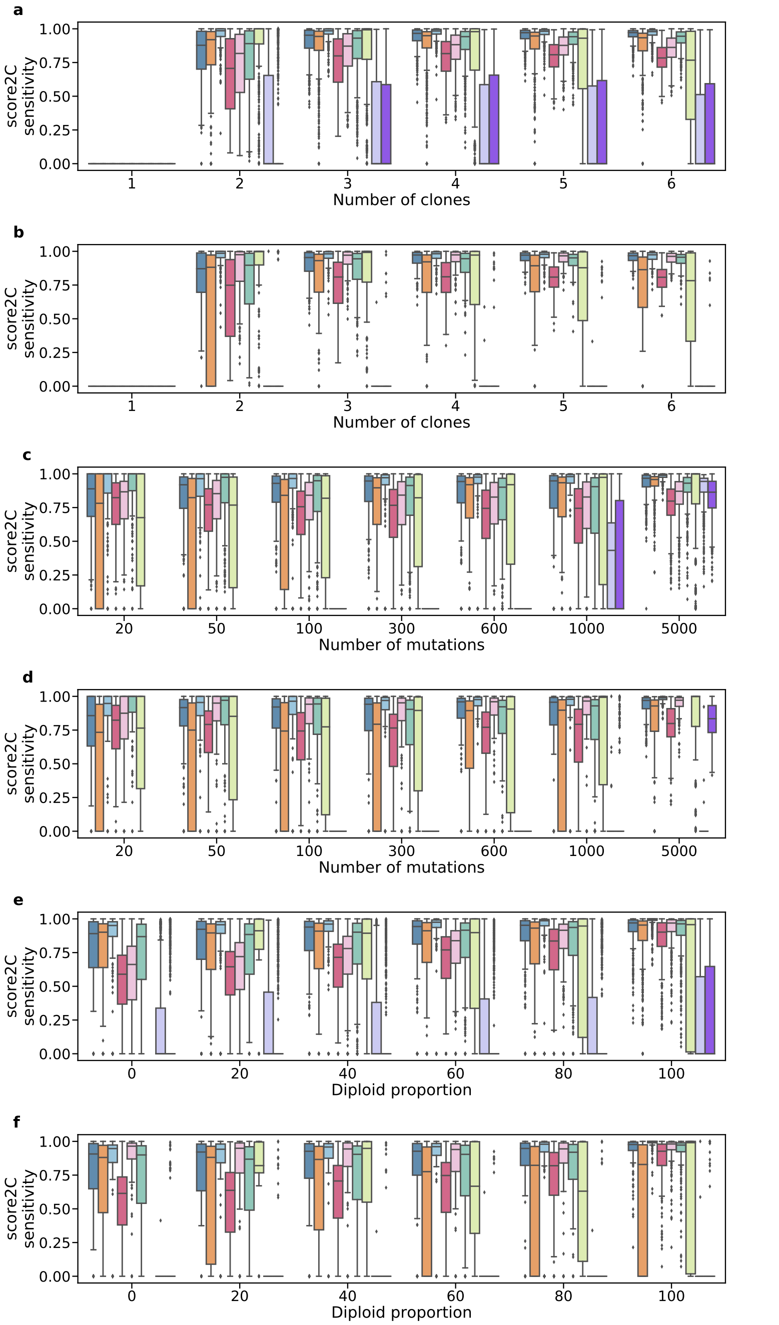

Supplementary Figure 16: **Score_2C (sensitivity) for ITH methods on the CloneSigSim dataset**, with varying number of clones (a, b), number of observed mutations (c, d) and diploid percent of the genome (e, f). Panels a, c and e correspond to simulations with varying signature between clones, and b, d, f to simulations with constant signatures. Each boxplot represents the three quartile values of the distribution along with extreme values. The "whiskers" extend to points that lie within 1.5 interquartile ranges of the lower and upper quartile, and then observations that fall outside this range are displayed independently. This means that each value in the boxplot corresponds to an actual observation in the data. We have ensured that all scores are comparable by removing samples where at least one method fails for all other methods the corresponding sample sues (such as SciClone and entirely non-diploid genomes, or TrackSig and TrackSigFreq for under 100 mutations, and PyClone with 5000 mutations in the constant setting). Results for each setting are presented for samples for which all methods ran successfully; there are respectively 442, 648, 655, 665, 660 and 659 samples with 1 to 6 clones (out of 1302 simulated samples) in the varying setting (panel a), and 162, 192, 188, 195, 184 and 182 (out of 420 simulated samples) in the constant setting (panel b), 434, 732, 795, 816, 813, 679, and 626 samples with 20, 50, 100, 300, 600, 1000 and 5000 observed mutations (out of 1116 simulated samples) in the varying setting (panel c), and 153, 265, 289, 290, 287, 237 and 224 (out of 360 simulated samples) in the constant setting (panel d), and finally 751, 773, 768, 780, 759 and 649 samples with 0, 20, 40, 60, 80 and 100 percent of the genome being diploid without alteration (out of 1302 simulated samples) in the varying setting (panel e), and 225, 224, 225, 234, 229 and 191 (out of 420 simulated samples) in the constant setting (panel f). Source data are provided as a Source Data file.

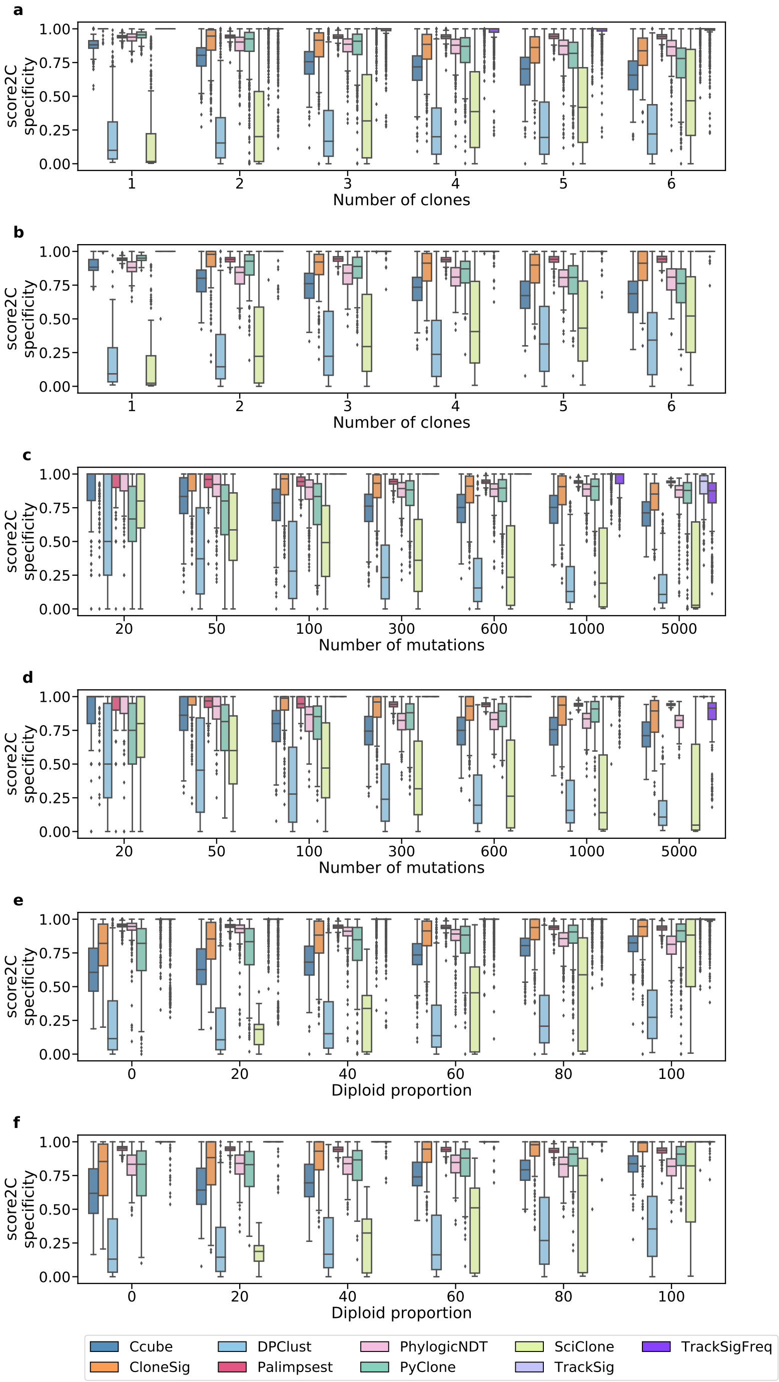

Supplementary Figure 17: **Score_2C (specificity) for ITH methods on the CloneSigSim dataset**, with varying number of clones (a, b), number of observed mutations (c, d) and diploid percent of the genome (e, f). Panels a, c and e correspond to simulations with varying signature between clones, and b, d, f to simulations with constant signatures. Each boxplot represents the three quartile values of the distribution along with extreme values. The "whiskers" extend to points that lie within 1.5 interquartile ranges of the lower and upper quartile, and then observations that fall outside this range are displayed independently. This means that each value in the boxplot corresponds to an actual observation in the data. We have ensured that all scores are comparable by removing samples where at least one method fails for all other methods the corresponding sample sues (such as SciClone and entirely non-diploid genomes, or TrackSig and TrackSigFreq for under 100 mutations, and PyClone with 5000 mutations in the constant setting). Results for each setting are presented for samples for which all methods ran successfully; there are respectively 442, 648, 655, 665, 660 and 659 samples with 1 to 6 clones (out of 1302 simulated samples) in the varying setting (panel a), and 162, 192, 188, 195, 184 and 182 (out of 420 simulated samples) in the constant setting (panel b), 434, 732, 795, 816, 813, 679, and 626 samples with 20, 50, 100, 300, 600, 1000 and 5000 observed mutations (out of 1116 simulated samples) in the varying setting (panel c), and 153, 265, 289, 290, 287, 237 and 224 (out of 360 simulated samples) in the constant setting (panel d), and finally 751, 773, 768, 780, 759 and 649 samples with 0, 20, 40, 60, 80 and 100 percent of the genome being diploid without alteration (out of 1302 simulated samples) in the varying setting (panel e), and 225, 224, 225, 234, 229 and 191 (out of 420 simulated samples) in the constant setting (panel f). Source data are provided as a Source Data file.

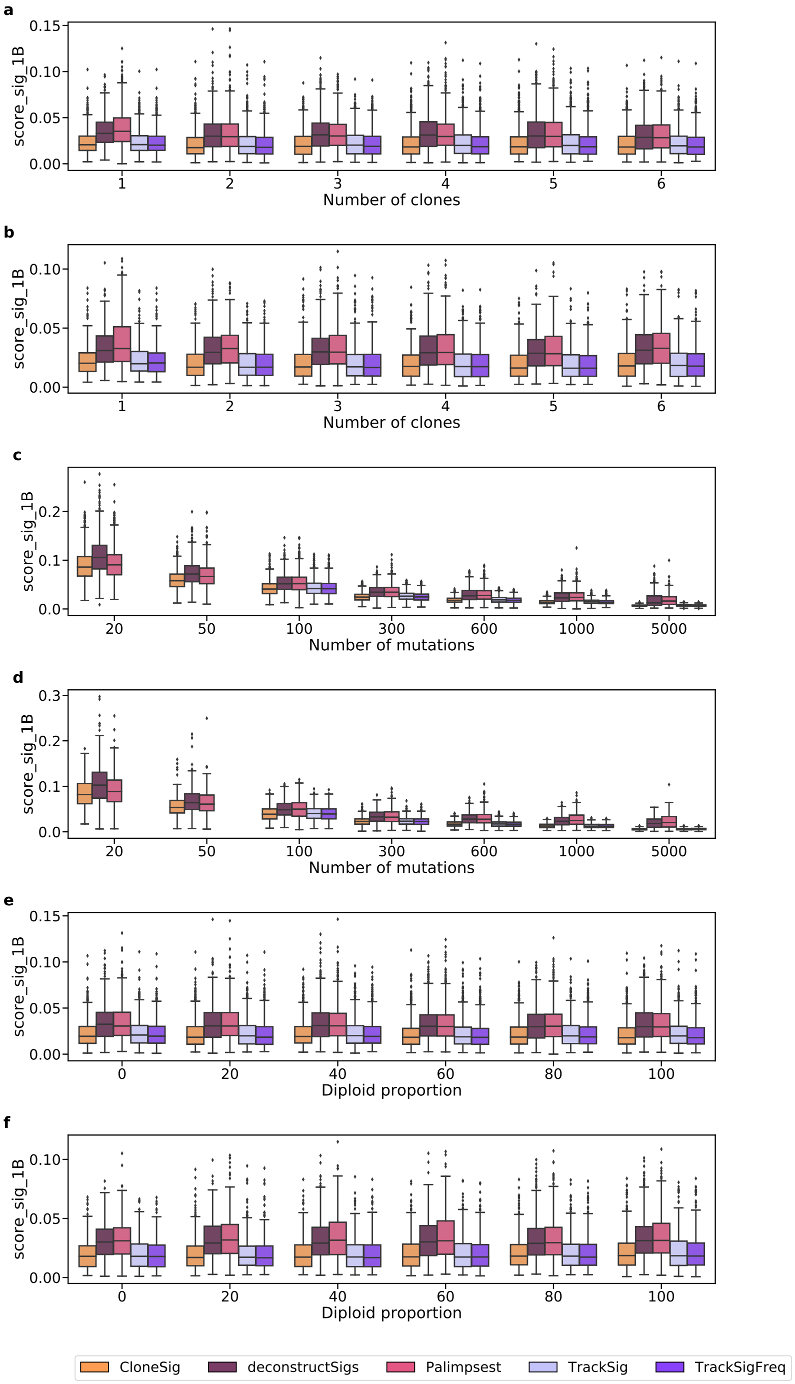

Supplementary Figure 18: **Score_sig_1B for signature activity deconvolution methods on the CloneSigSim dataset**, with varying number of clones (a, b), number of observed mutations (c, d) and diploid percent of the genome (e, f). Panels a, c and e correspond to simulations with varying signature between clones, and b, d, f to simulations with constant signatures. Each boxplot represents the three quartile values of the distribution along with extreme values. The "whiskers" extend to points that lie within 1.5 interquartile ranges of the lower and upper quartile, and then observations that fall outside this range are displayed independently. This means that each value in the boxplot corresponds to an actual observation in the data. We have ensured that all scores are comparable by removing samples where at least one method fails for all other methods the corresponding sample sues (such as SciClone and entirely non-diploid genomes, or TrackSig and TrackSigFreq for under 100 mutations, and PyClone with 5000 mutations in the constant setting). Results for each setting are presented for samples for which all methods ran successfully; there are respectively 659, 791, 795, 791, 793 and 782 samples with 1 to 6 clones (out of 1302 simulated samples) in the varying setting (panel a), and 233, 281, 284, 292, 277 and 275 (out of 420 simulated samples) in the constant setting (panel b), 861, 995, 992, 1010, 1012, 835, and 762 samples with 20, 50, 100, 300, 600, 1000 and 5000 observed mutations (out of 1116 simulated samples) in the varying setting (panel c), and 303, 366, 357, 360, 358, 291 and 276 (out of 360 simulated samples) in the constant setting (panel d), and finally 755, 781, 769, 782, 759 and 765 samples with 0, 20, 40, 60, 80 and 100 percent of the genome being diploid without alteration (out of 1302 simulated samples) in the varying setting (panel e), and 277, 272, 269, 280, 271 and 273 (out of 420 simulated samples) in the constant setting (panel f). Source data are provided as a Source Data file.

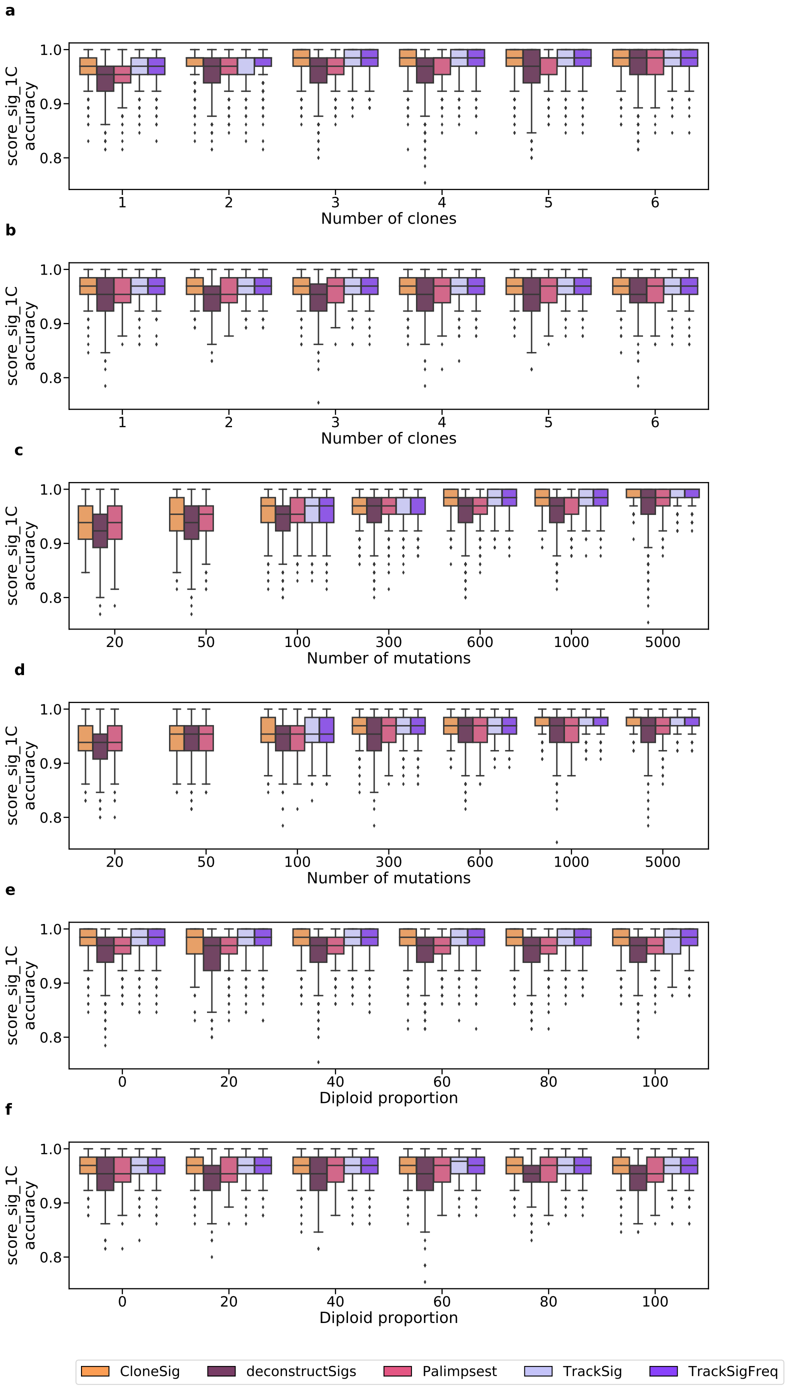

Supplementary Figure 19: **Score_sig_1C (accuracy) for signature activity deconvolution methods on the CloneSigSim dataset**, with varying number of clones (a, b), number of observed mutations (c, d) and diploid percent of the genome (e, f). Panels a, c and e correspond to simulations with varying signature between clones, and b, d, f to simulations with constant signatures. Each boxplot represents the three quartile values of the distribution along with extreme values. The "whiskers" extend to points that lie within 1.5 interquartile ranges of the lower and upper quartile, and then observations that fall outside this range are displayed independently. This means that each value in the boxplot corresponds to an actual observation in the data. We have ensured that all scores are comparable by removing samples where at least one method fails for all other methods the corresponding sample whes (such as SciClone and entirely non-diploid genomes, or TrackSig and TrackSigFreq for under 100 mutations, and PyClone with 5000 mutations in the constant setting). Results for each setting are presented for samples for which all methods ran successfully; there are respectively 659, 791, 795, 791, 793 and 782 samples with 1 to 6 clones (out of 1302 simulated samples) in the varying setting (panel a), and 233, 281, 284, 292, 277 and 275 (out of 420 simulated samples) in the constant setting (panel b), 861, 995, 992, 1010, 1012, 835, and 762 samples with 20, 50, 100, 300, 600, 1000 and 5000 observed mutations (out of 1116 simulated samples) in the varying setting (panel c), and 303, 366, 357, 360, 358, 291 and 276 (out of 360 simulated samples) in the constant setting (panel d), and finally 755, 781, 769, 782, 759 and 765 samples with 0, 20, 40, 60, 80 and 100 percent of the genome being diploid without alteration (out of 1302 simulated samples) in the varying setting (panel e), and 277, 272, 269, 280, 271 and 273 (out of 420 simulated samples) in the constant setting (panel f). Source data are provided as a Source Data file.

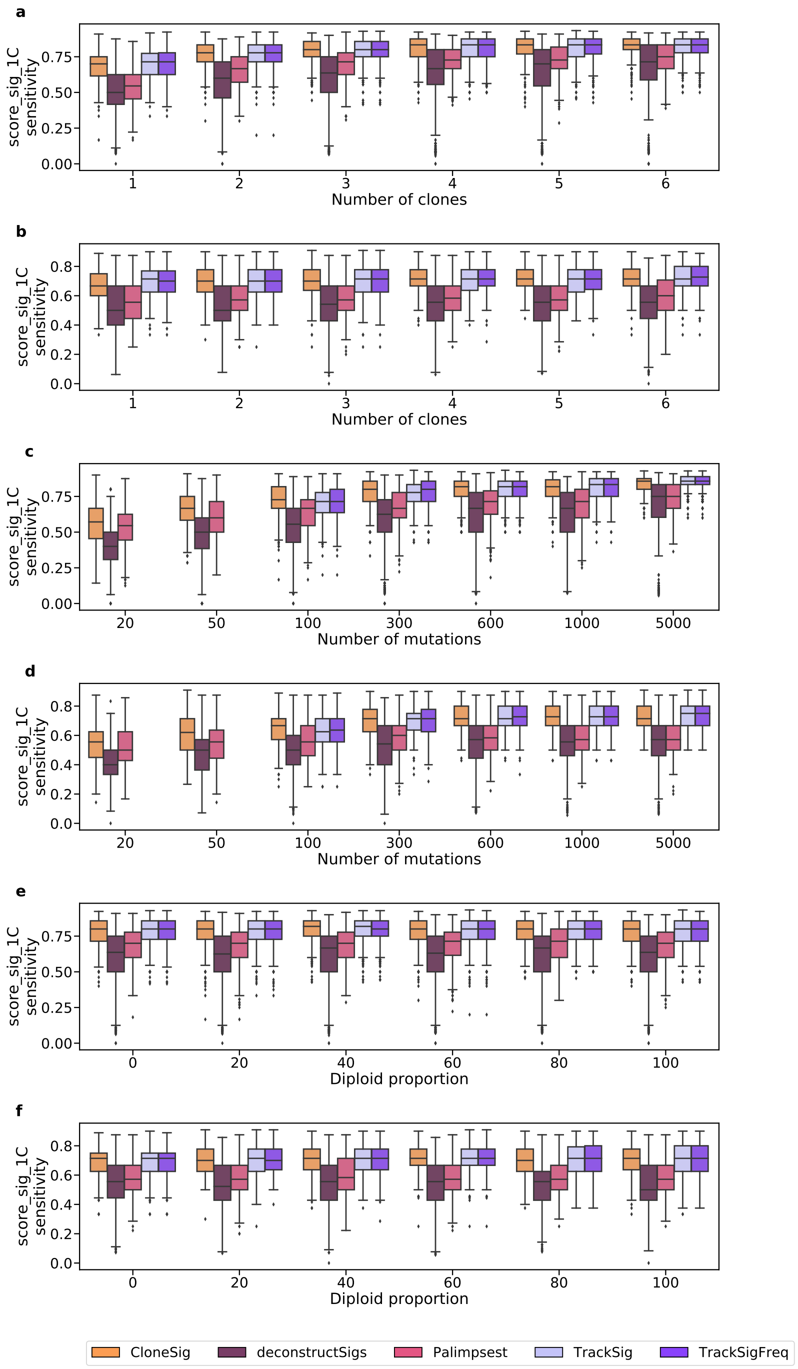

Supplementary Figure 20: **Score_sig_1C (sensitivity) for signature activity deconvolution methods on the CloneSigSim dataset**, with varying number of clones (a, b), number of observed mutations (c, d) and diploid percent of the genome (e, f). Panels a, c and e correspond to simulations with varying signature between clones, and b, d, f to simulations with constant signatures. Each boxplot represents the three quartile values of the distribution along with extreme values. The "whiskers" extend to points that lie within 1.5 interquartile ranges of the lower and upper quartile, and then observations that fall outside this range are displayed independently. This means that each value in the boxplot corresponds to an actual observation in the data. We have ensured that all scores are comparable by removing samples where at least one method fails for all other methods the corresponding sample whes (such as SciClone and entirely non-diploid genomes, or TrackSig and TrackSigFreq for under 100 mutations, and PyClone with 5000 mutations in the constant setting). Results for each setting are presented for samples for which all methods ran successfully; there are respectively 659, 791, 795, 791, 793 and 782 samples with 1 to 6 clones (out of 1302 simulated samples) in the varying setting (panel a), and 233, 281, 284, 292, 277 and 275 (out of 420 simulated samples) in the constant setting (panel b), 861, 995, 992, 1010, 1012, 835, and 762 samples with 20, 50, 100, 300, 600, 1000 and 5000 observed mutations (out of 1116 simulated samples) in the varying setting (panel c), and 303, 366, 357, 360, 358, 291 and 276 (out of 360 simulated samples) in the constant setting (panel d), and finally 755, 781, 769, 782, 759 and 765 samples with 0, 20, 40, 60, 80 and 100 percent of the genome being diploid without alteration (out of 1302 simulated samples) in the varying setting (panel e), and 277, 272, 269, 280, 271 and 273 (out of 420 simulated samples) in the constant setting (panel f). Source data are provided as a Source Data file.

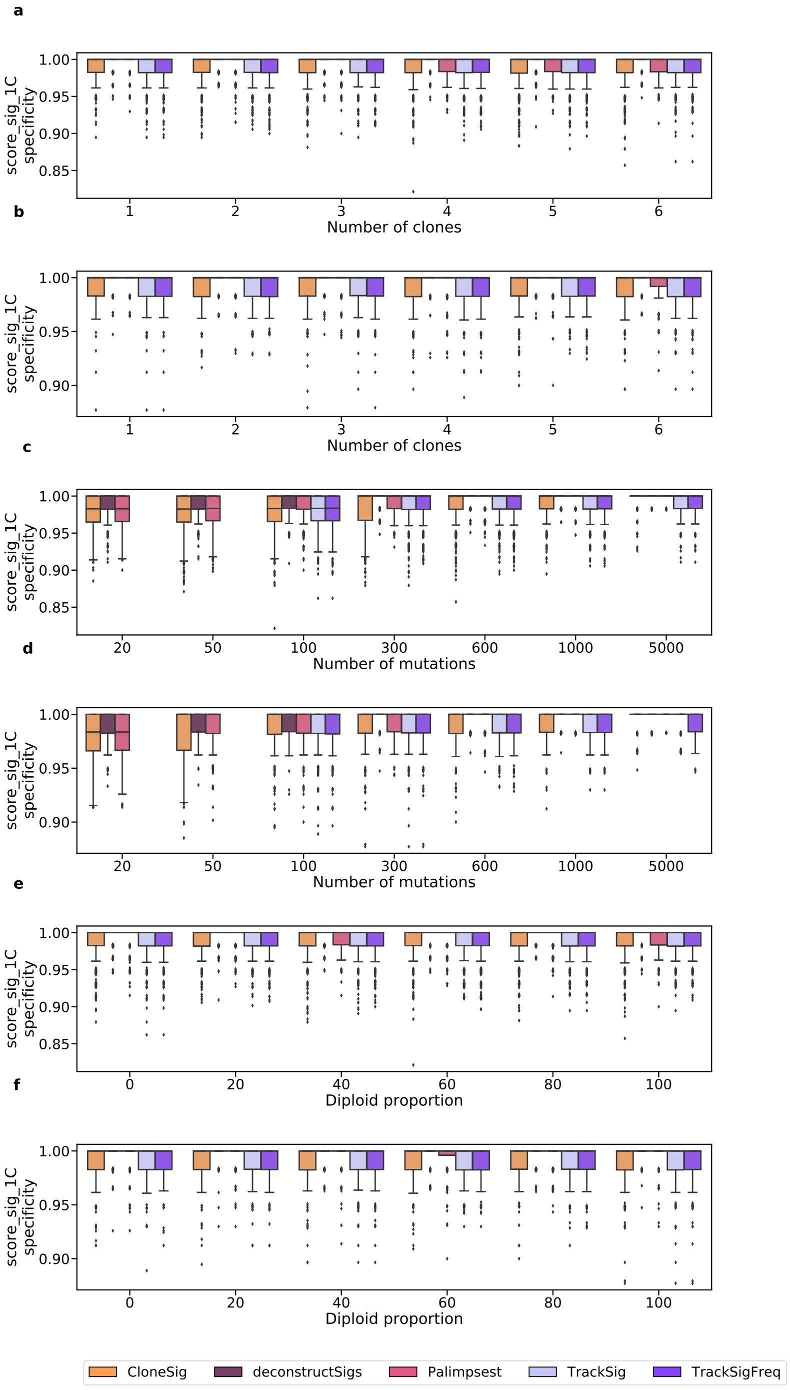

Supplementary Figure 21: **Score_sig_1C (specificity) for signature activity deconvolution methods on the CloneSigSim dataset**, with varying number of clones (a, b), number of observed mutations (c, d) and diploid percent of the genome (e, f). Panels a, c and e correspond to simulations with varying signature between clones, and b, d, f to simulations with constant signatures. Each boxplot represents the three quartile values of the distribution along with extreme values. The "whiskers" extend to points that lie within 1.5 interquartile ranges of the lower and upper quartile, and then observations that fall outside this range are displayed independently. This means that each value in the boxplot corresponds to an actual observation in the data. We have ensured that all scores are comparable by removing samples where at least one method fails for all other methods the corresponding sample whes (such as SciClone and entirely non-diploid genomes, or TrackSig and TrackSigFreq for under 100 mutations, and PyClone with 5000 mutations in the constant setting). Results for each setting are presented for samples for which all methods ran successfully; there are respectively 659, 791, 795, 791, 793 and 782 samples with 1 to 6 clones (out of 1302 simulated samples) in the varying setting (panel a), and 233, 281, 284, 292, 277 and 275 (out of 420 simulated samples) in the constant setting (panel b), 861, 995, 992, 1010, 1012, 835, and 762 samples with 20, 50, 100, 300, 600, 1000 and 5000 observed mutations (out of 1116 simulated samples) in the varying setting (panel c), and 303, 366, 357, 360, 358, 291 and 276 (out of 360 simulated samples) in the constant setting (panel d), and finally 755, 781, 769, 782, 759 and 765 samples with 0, 20, 40, 60, 80 and 100 percent of the genome being diploid without alteration (out of 1302 simulated samples) in the varying setting (panel e), and 277, 272, 269, 280, 271 and 273 (out of 420 simulated samples) in the constant setting (panel f). Source data are provided as a Source Data file.

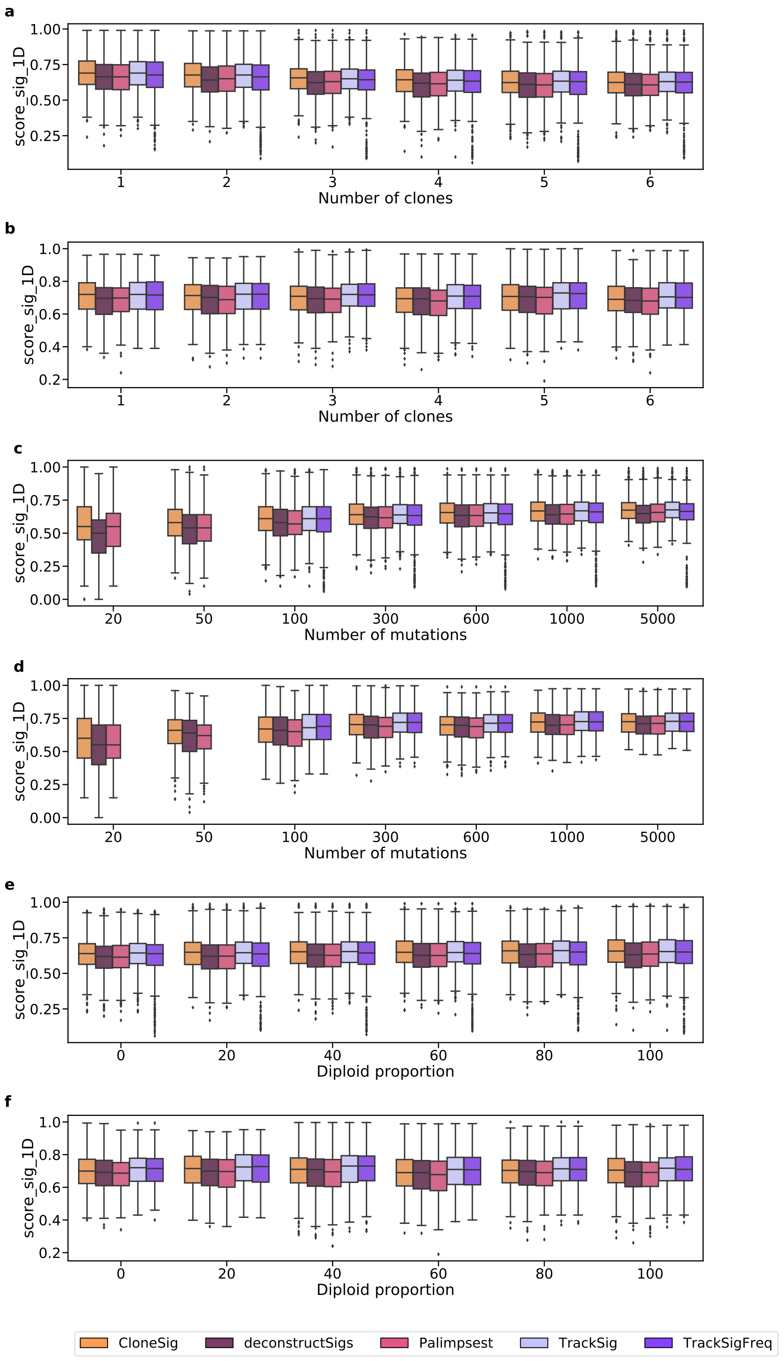

Supplementary Figure 22: **Score_sig_1D for signature activity deconvolution methods on the CloneSigSim dataset**, with varying number of clones (a, b), number of observed mutations (c, d) and diploid percent of the genome (e, f). Panels a, c and e correspond to simulations with varying signature between clones, and b, d, f to simulations with constant signatures. Each boxplot represents the three quartile values of the distribution along with extreme values. The "whiskers" extend to points that lie within 1.5 interquartile ranges of the lower and upper quartile, and then observations that fall outside this range are displayed independently. This means that each value in the boxplot corresponds to an actual observation in the data. We have ensured that all scores are comparable by removing samples where at least one method fails for all other methods the corresponding sample sues (such as SciClone and entirely non-diploid genomes, or TrackSig and TrackSigFreq for under 100 mutations, and PyClone with 5000 mutations in the constant setting). Results for each setting are presented for samples for which all methods ran successfully; there are respectively 659, 791, 795, 791, 793 and 782 samples with 1 to 6 clones (out of 1302 simulated samples) in the varying setting (panel a), and 233, 281, 284, 292, 277 and 275 (out of 420 simulated samples) in the constant setting (panel b), 861, 995, 992, 1010, 1012, 835, and 762 samples with 20, 50, 100, 300, 600, 1000 and 5000 observed mutations (out of 1116 simulated samples) in the varying setting (panel c), and 303, 366, 357, 360, 358, 291 and 276 (out of 360 simulated samples) in the constant setting (panel d), and finally 755, 781, 769, 782, 759 and 765 samples with 0, 20, 40, 60, 80 and 100 percent of the genome being diploid without alteration (out of 1302 simulated samples) in the varying setting (panel e), and 277, 272, 269, 280, 271 and 273 (out of 420 simulated samples) in the constant setting (panel f). Source data are provided as a Source Data file.

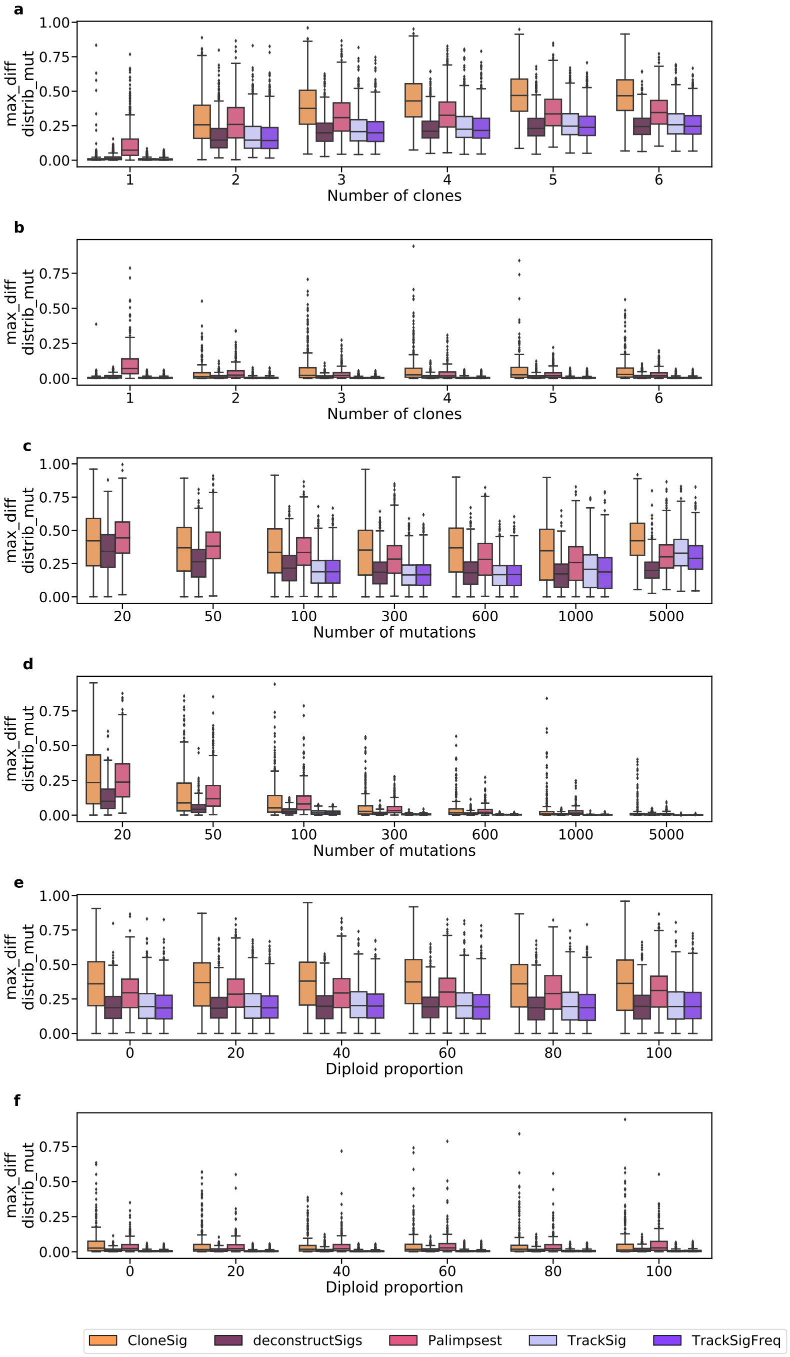

Supplementary Figure 23: **Maximal cosine distance between the true and estimated mutation type profile for signature activity deconvolution methods on the CloneSigSim dataset**, with varying number of clones (a, b), number of observed mutations (c, d) and diploid percent of the genome (e, f). Panels a, c and e correspond to simulations with varying signature between clones, and b, d, f to simulations with constant signatures. Each boxplot represents the three quartile values of the distribution along with extreme values. The "whiskers" extend to points that lie within 1.5 interquartile ranges of the lower and upper quartile, and then observations that fall outside this range are displayed independently. This means that each value in the boxplot corresponds to an actual observation in the data. We have ensured that all scores are comparable by removing samples where at least one method fails for all other methods the corresponding sample wues (such as SciClone and entirely non-diploid genomes, or TrackSig and TrackSigFreq for under 100 mutations, and PyClone with 5000 mutations in the constant setting). Results for each setting are presented for samples for which all methods ran successfully; there are respectively 659, 791, 795, 791, 793 and 782 samples with 1 to 6 clones (out of 1302 simulated samples) in the varying setting (panel a), and 233, 281, 284, 292, 277 and 275 (out of 420 simulated samples) in the constant setting (panel b), 861, 995, 992, 1010, 1012, 835, and 762 samples with 20, 50, 100, 300, 600, 1000 and 5000 observed mutations (out of 1116 simulated samples) in the varying setting (panel c), and 303, 366, 357, 360, 358, 291 and 276 (out of 360 simulated samples) in the constant setting (panel d), and finally 755, 781, 769, 782, 759 and 765 samples with 0, 20, 40, 60, 80 and 100 percent of the genome being diploid without alteration (out of 1302 simulated samples) in the varying setting (panel e), and 277, 272, 269, 280, 271 and 273 (out of 420 simulated samples) in the constant setting (panel f). Source data are provided as a Source Data file.

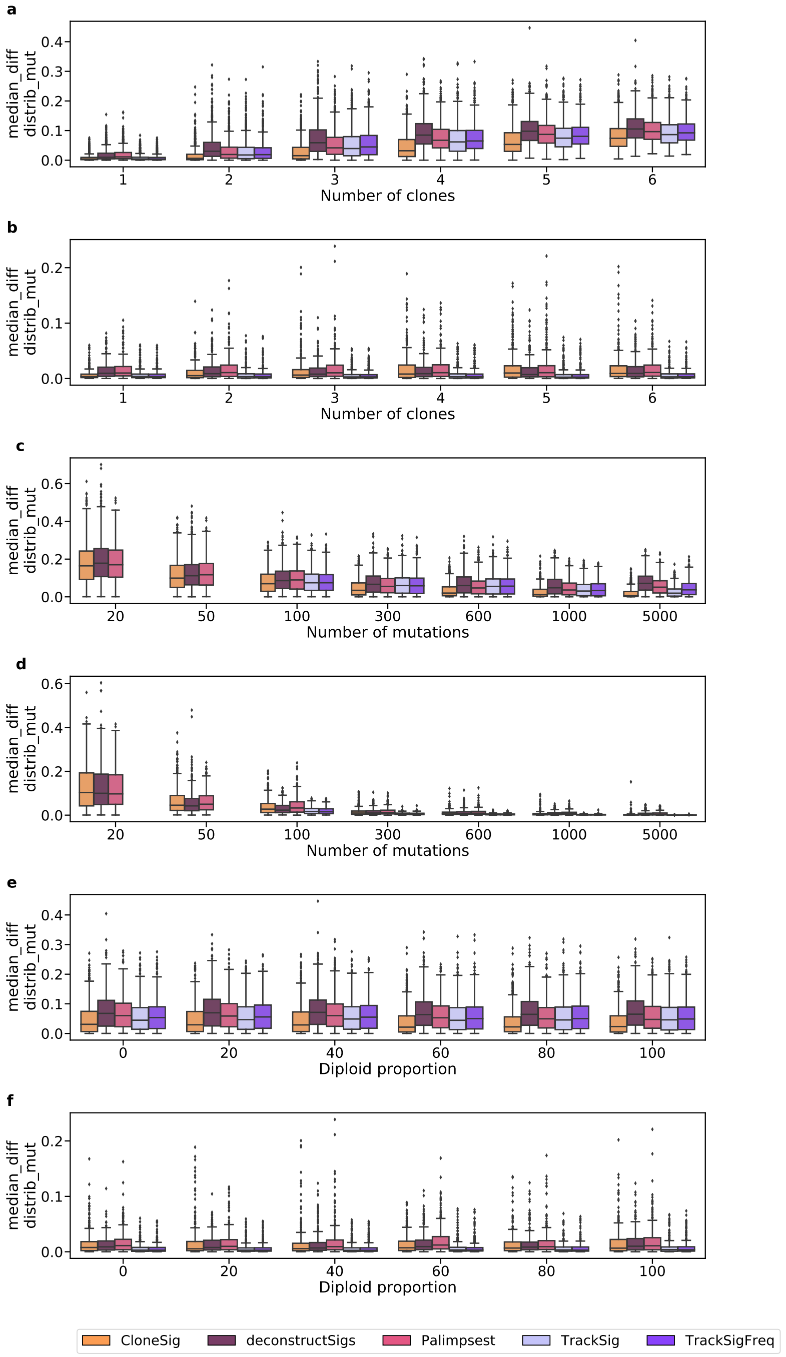

Supplementary Figure 24: **Median cosine distance between the true and estimated mutation type profile for signature activity deconvolution methods on the CloneSigSim dataset**, with varying number of clones (a, b), number of observed mutations (c, d) and diploid percent of the genome (e, f). Panels a, c and e correspond to simulations with varying signature between clones, and b, d, f to simulations with constant signatures. Each boxplot represents the three quartile values of the distribution along with extreme values. The "whiskers" extend to points that lie within 1.5 interquartile ranges of the lower and upper quartile, and then observations that fall outside this range are displayed independently. This means that each value in the boxplot corresponds to an actual observation in the data. We have ensured that all scores are comparable by removing samples where at least one method fails for all other methods the corresponding sample whes (such as SciClone and entirely non-diploid genomes, or TrackSig and TrackSigFreq for under 100 mutations, and PyClone with 5000 mutations in the constant setting). Results for each setting are presented for samples for which all methods ran successfully; there are respectively 659, 791, 795, 791, 793 and 782 samples with 1 to 6 clones (out of 1302 simulated samples) in the varying setting (panel a), and 233, 281, 284, 292, 277 and 275 (out of 420 simulated samples) in the constant setting (panel b), 861, 995, 992, 1010, 1012, 835, and 762 samples with 20, 50, 100, 300, 600, 1000 and 5000 observed mutations (out of 1116 simulated samples) in the varying setting (panel c), and 303, 366, 357, 360, 358, 291 and 276 (out of 360 simulated samples) in the constant setting (panel d), and finally 755, 781, 769, 782, 759 and 765 samples with 0, 20, 40, 60, 80 and 100 percent of the genome being diploid without alteration (out of 1302 simulated samples) in the varying setting (panel e), and 277, 272, 269, 280, 271 and 273 (out of 420 simulated samples) in the constant setting (panel f). Source data are provided as a Source Data file.

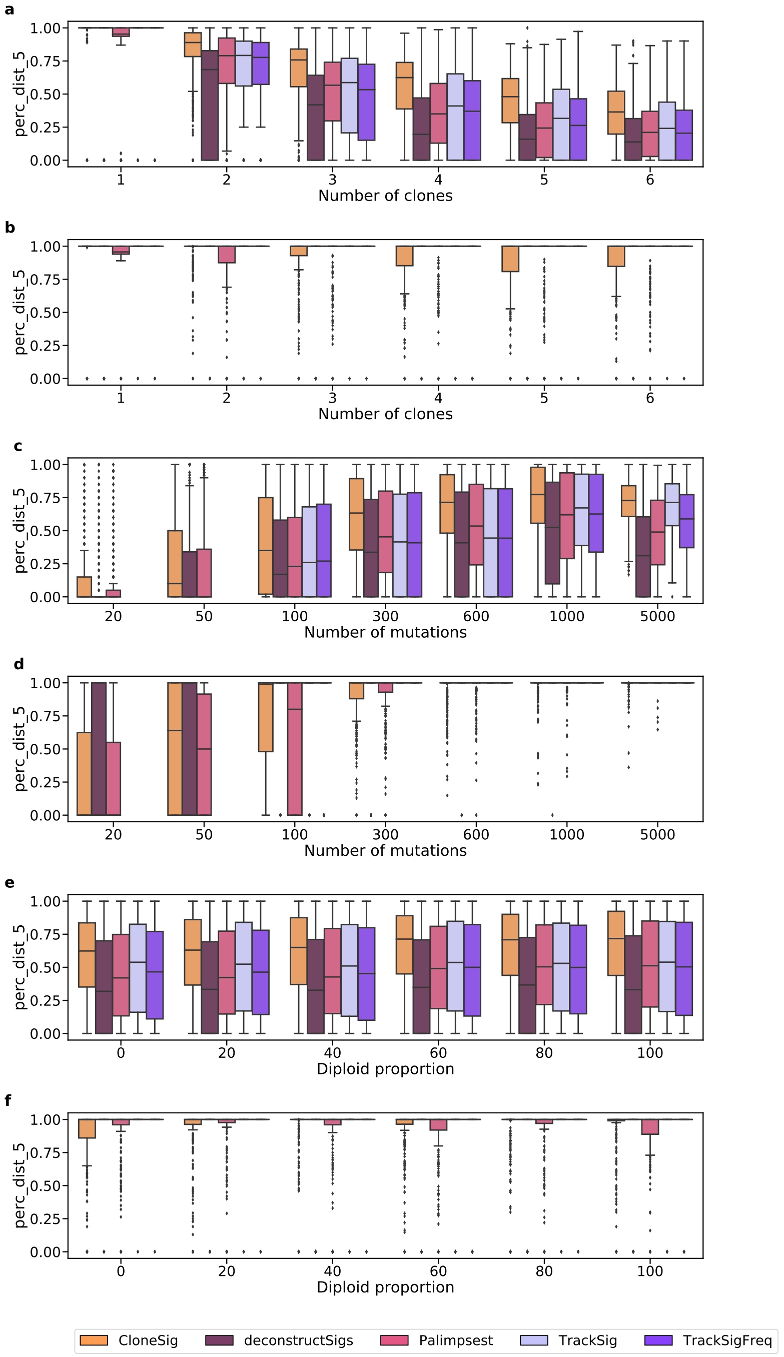

Supplementary Figure 25: **Proportion of SNVs with cosine distance between the true and estimated mutation type profile under 0.05 for signature activity deconvolution methods on the CloneSigSim dataset**, with varying number of clones (a, b), number of observed mutations (c, d) and diploid percent of the genome (e, f). Panels a, c and e correspond to simulations with varying signature between clones, and b, d, f to simulations with constant signatures. Each boxplot represents the three quartile values of the distribution along with extreme values. The "whiskers" extend to points that lie within 1.5 interquartile ranges of the lower and upper quartile, and then observations that fall outside this range are displayed independently. This means that each value in the boxplot corresponds to an actual observation in the data. We have ensured that all scores are comparable by removing samples where at least one method fails for all other methods the corresponding sample sues (such as SciClone and entirely non-diploid genomes, or TrackSig and TrackSigFreq for under 100 mutations, and PyClone with 5000 mutations in the constant setting). Results for each setting are presented for samples for which all methods ran successfully; there are respectively 659, 791, 795, 791, 793 and 782 samples with 1 to 6 clones (out of 1302 simulated samples) in the varying setting (panel a), and 233, 281, 284, 292, 277 and 275 (out of 420 simulated samples) in the constant setting (panel b), 861, 995, 992, 1010, 1012, 835, and 762 samples with 20, 50, 100, 300, 600, 1000 and 5000 observed mutations (out of 1116 simulated samples) in the varying setting (panel c), and 303, 366, 357, 360, 358, 291 and 276 (out of 360 simulated samples) in the constant setting (panel d), and finally 755, 781, 769, 782, 759 and 765 samples with 0, 20, 40, 60, 80 and 100 percent of the genome being diploid without alteration (out of 1302 simulated samples) in the varying setting (panel e), and 277, 272, 269, 280, 271 and 273 (out of 420 simulated samples) in the constant setting (panel f). Source data are provided as a Source Data file.

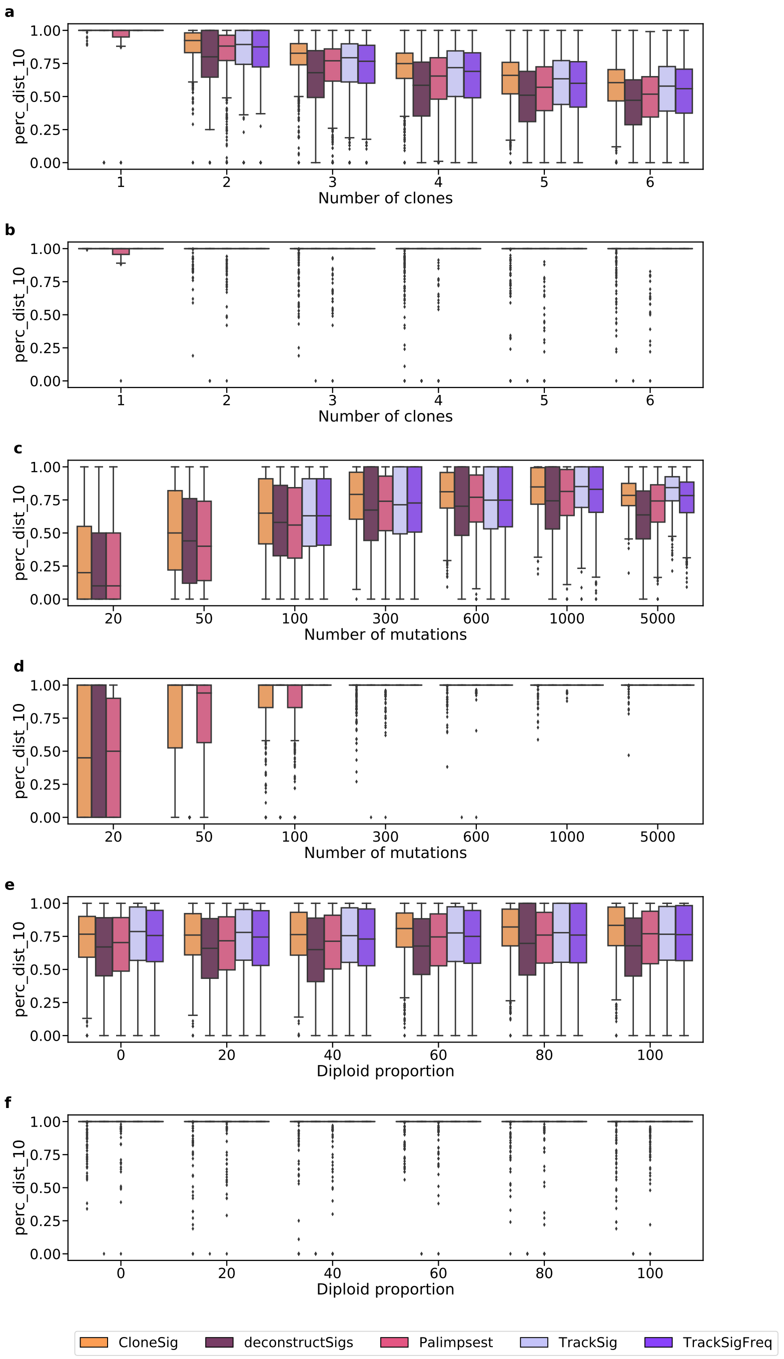

Supplementary Figure 26: **Proportion of SNVs with cosine distance between the true and estimated mutation type profile under 0.10 for signature activity deconvolution methods on the CloneSigSim dataset**, with varying number of clones (a, b), number of observed mutations (c, d) and diploid percent of the genome (e, f). Panels a, c and e correspond to simulations with varying signature between clones, and b, d, f to simulations with constant signatures. Each boxplot represents the three quartile values of the distribution along with extreme values. The "whiskers" extend to points that lie within 1.5 interquartile ranges of the lower and upper quartile, and then observations that fall outside this range are displayed independently. This means that each value in the boxplot corresponds to an actual observation in the data. We have ensured that all scores are comparable by removing samples where at least one method fails for all other methods the corresponding sample sues (such as SciClone and entirely non-diploid genomes, or TrackSig and TrackSigFreq for under 100 mutations, and PyClone with 5000 mutations in the constant setting). Results for each setting are presented for samples for which all methods ran successfully; there are respectively 659, 791, 795, 791, 793 and 782 samples with 1 to 6 clones (out of 1302 simulated samples) in the varying setting (panel a), and 233, 281, 284, 292, 277 and 275 (out of 420 simulated samples) in the constant setting (panel b), 861, 995, 992, 1010, 1012, 835, and 762 samples with 20, 50, 100, 300, 600, 1000 and 5000 observed mutations (out of 1116 simulated samples) in the varying setting (panel c), and 303, 366, 357, 360, 358, 291 and 276 (out of 360 simulated samples) in the constant setting (panel d), and finally 755, 781, 769, 782, 759 and 765 samples with 0, 20, 40, 60, 80 and 100 percent of the genome being diploid without alteration (out of 1302 simulated samples) in the varying setting (panel e), and 277, 272, 269, 280, 271 and 273 (out of 420 simulated samples) in the constant setting (panel f). Source data are provided as a Source Data file.

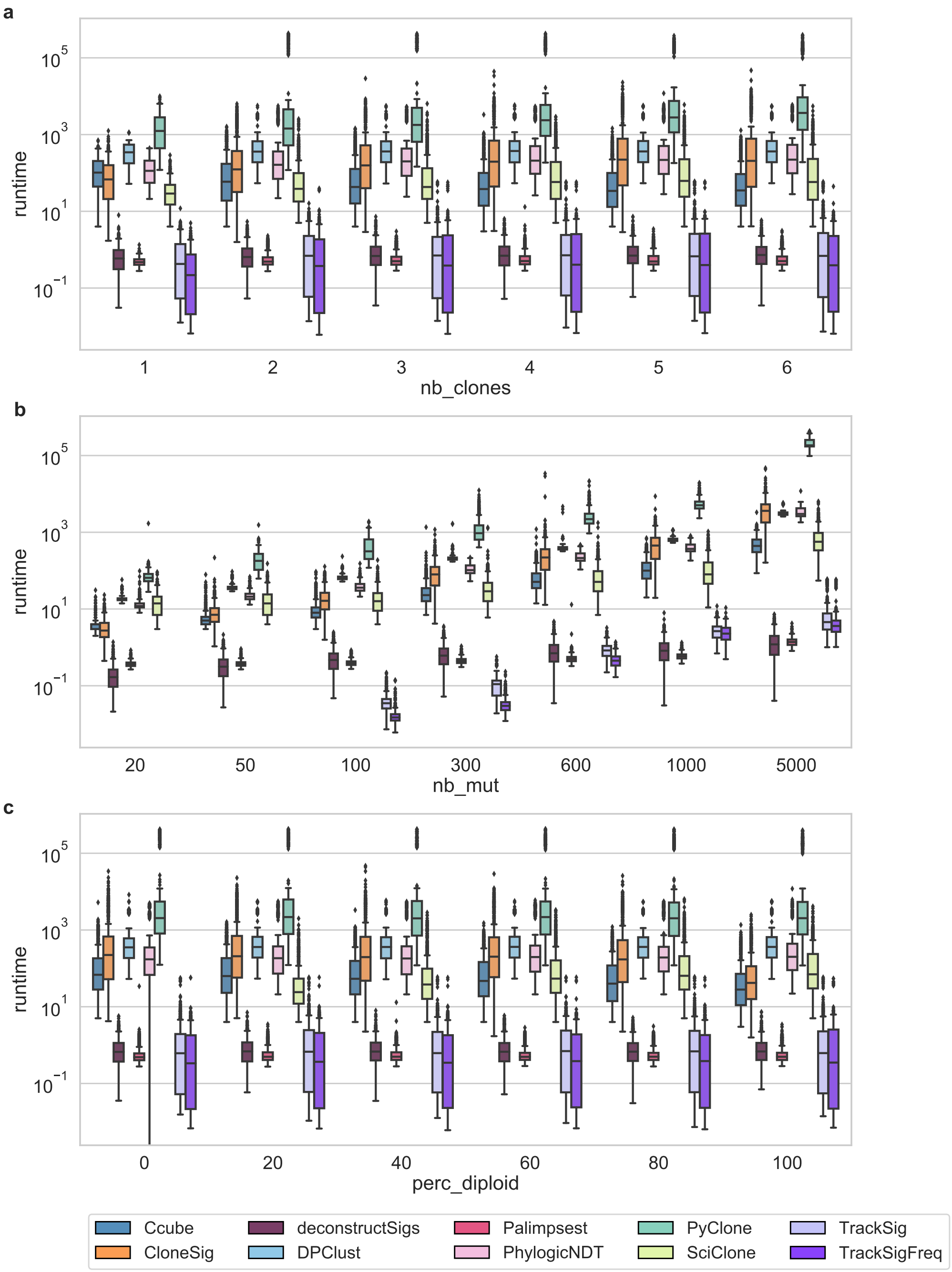

Supplementary Figure 27: **Runtime for ITH reconstruction and signature activity deconvolution methods on the CloneSigSim dataset**, with varying number of clones (a), number of observed mutations (b) and diploid percent of the genome (c). Results with varying signature between clones only are shown but similar results were obtained on simulations with constant signatures. Each boxplot represents the three quartile values of the distribution along with extreme values. The "whiskers" extend to points that lie within 1.5 interquartile ranges of the lower and upper quartile, and then observations that fall outside this range are displayed independently. This means that each value in the boxplot corresponds to an actual observation in the data. We have ensured that all scores are comparable by removing samples where at least one method fails for all other methods the corresponding sample sues (such as SciClone and entirely non-diploid genomes, or TrackSig and TrackSigFreq for under 100 mutations, and PyClone with 5000 mutations in the constant setting). Results for each setting are presented for samples for which all methods ran successfully; there are respectively 442, 648, 655, 665, 660 and 659 samples with 1 to 6 clones (out of 1302 simulated samples) in the varying setting (panel a), and 162, 192, 188, 195, 184 and 182 (out of 420 simulated samples) in the constant setting (panel b), 434, 732, 795, 816, 813, 679, and 626 samples with 20, 50, 100, 300, 600, 1000 and 5000 observed mutations (out of 1116 simulated samples) in the varying setting (panel c), and 153, 265, 289, 290, 287, 237 and 224 (out of 360 simulated samples) in the constant setting (panel d), and finally 751, 773, 768, 780, 759 and 649 samples with 0, 20, 40, 60, 80 and 100 percent of the genome being diploid without alteration (out of 1302 simulated samples) in the varying setting (panel e), and 225, 224, 225, 234, 229 and 191 (out of 420 simulated samples) in the constant setting (panel f). Source data are provided as a Source Data file.

#### 2.2 DREAM challenge dataset

We tested all ITH inference methods on the DREAM Challenge dataset [14, 15], with a time limit of 48 hours. Figure 28 shows the subclonal reconstruction performance metrics. We observe overall very similar performances for CloneSig, Ccube, Palimpsest, DPClust and PhylogicNDT in most cases, except for score2A where Ccube, CloneSig, PhylogicNDT and Palimpsest clearly outperform other methods. PyClone failed to run in less than 48 hours for most samples and was not represented on this Figure. Complete results per sample are available in Supplementary Figures 29 and 30. Results for signature reconstruction metrics are shows in Supplementary Figures 31-33.

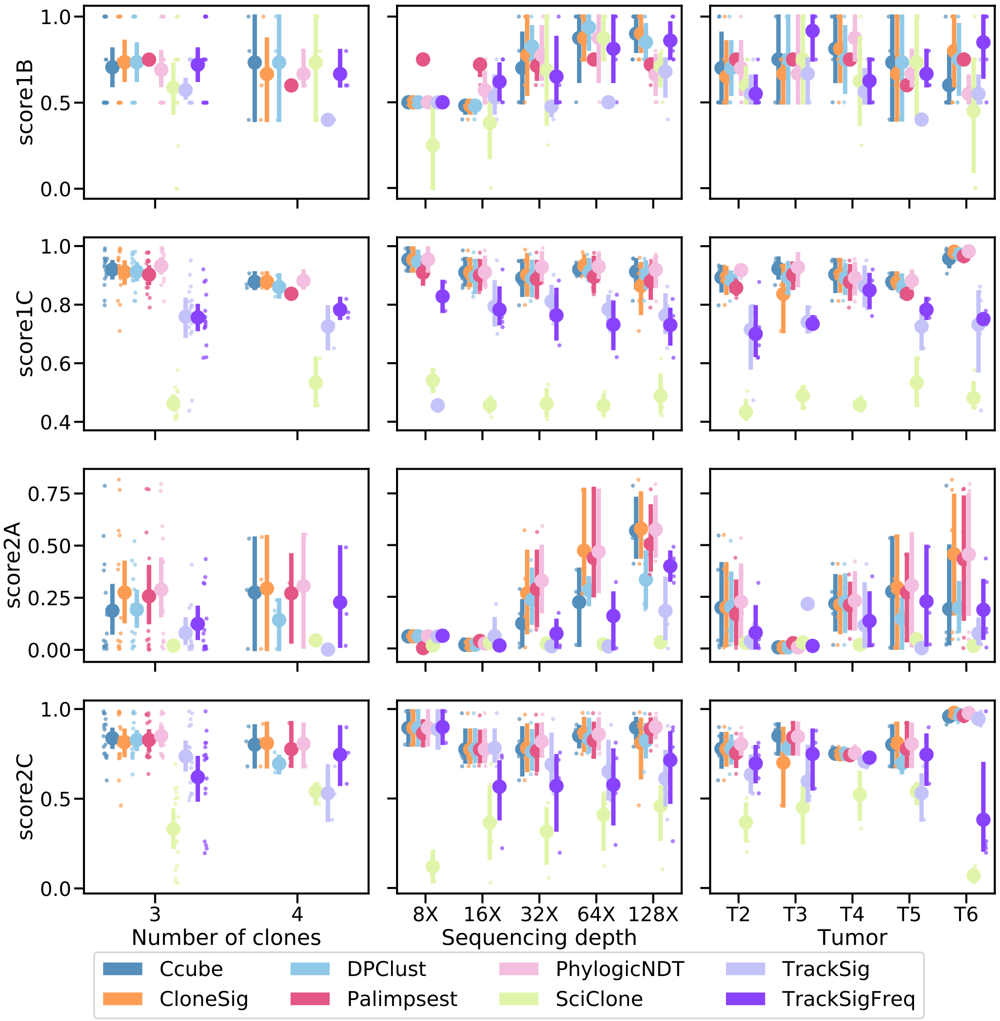

Supplementary Figure 28: **Comparison of CloneSig, TrackSig, TrackSigFreq, Palimpsest, SciClone, DPClust, PhylogicNDT and Ccube for subclonal reconstruction on the DREAM dataset.** Each row corresponds to one score, as detailed in the main text. Score1B evaluates the number of clones found by the method, score1C the resulting mutation CCF distribution, score2A the co-clustering of mutations in the defined clones, and score2C the classification of subclonal versus clonal mutations. All scores are normalized between 0 and 1, with 1 being the best and 0 the worst. Each column corresponds to a setting where one parameter in the simulation varies: the true number of clones (left), the sequencing depth (middle), and the simulated tumor (right). Each point represents the average of the score over all available simulated samples. Bootstrap sampling of the scores was used to compute 95% confidence intervals of the mean. We have ensured that all scores are comparable by removing samples where at least one method fails for all other methods the corresponding stic issues (such as SciClone and entirely non-diploid genomes, or TrackSig and TrackSigFreq for under 100 mutations). Score2A requires heavy computations in our implementation, and was not computed in some cases. As a result, there are respectively 17 (15 for score2A) with 3 clones and 3 samples with 4 clones, 2, 5, 4, 4 (3 for score2A) and 5 (4 for score2A) samples with a depth of 8X, 16X, 32X, 64X and 128X, and 5, 3 (1 for score2A), 4, 3 and 5 samples for tumors T2, T3, T4, T5 and T6. Source data are provided as a Source Data file.

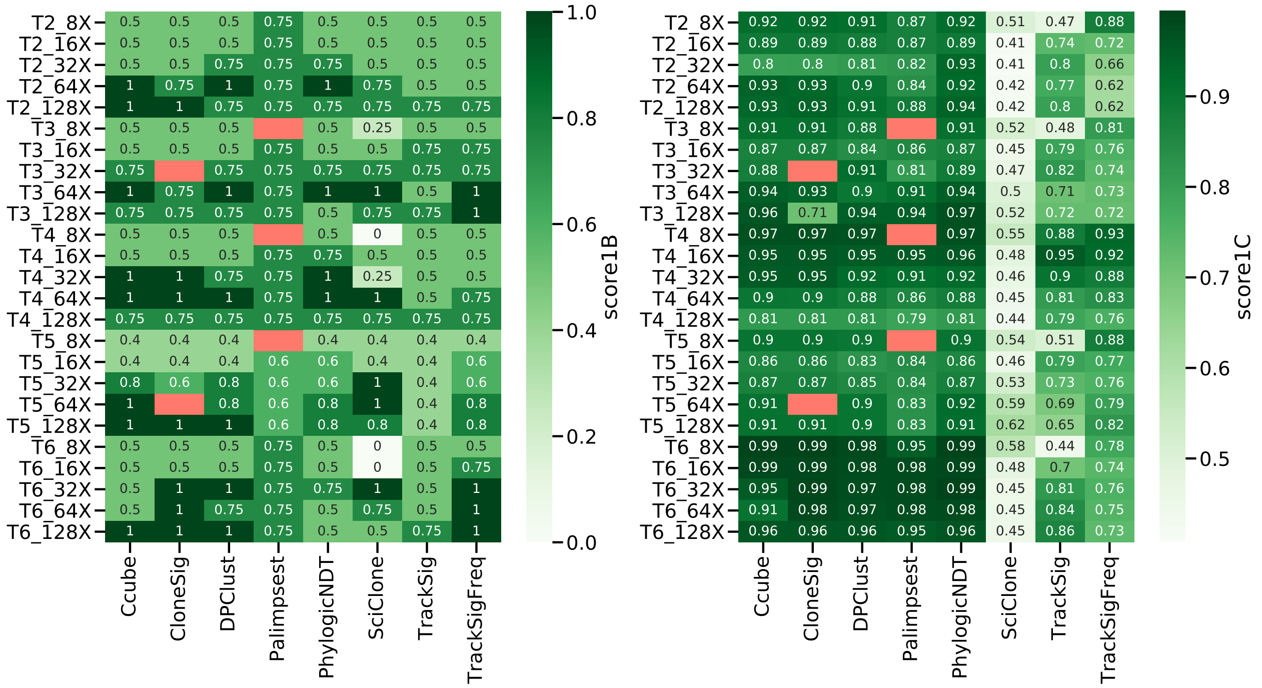

Supplementary Figure 29: **Complete results for the DREAM dataset, score1B and score1C.** Source data are provided as a Source Data file.

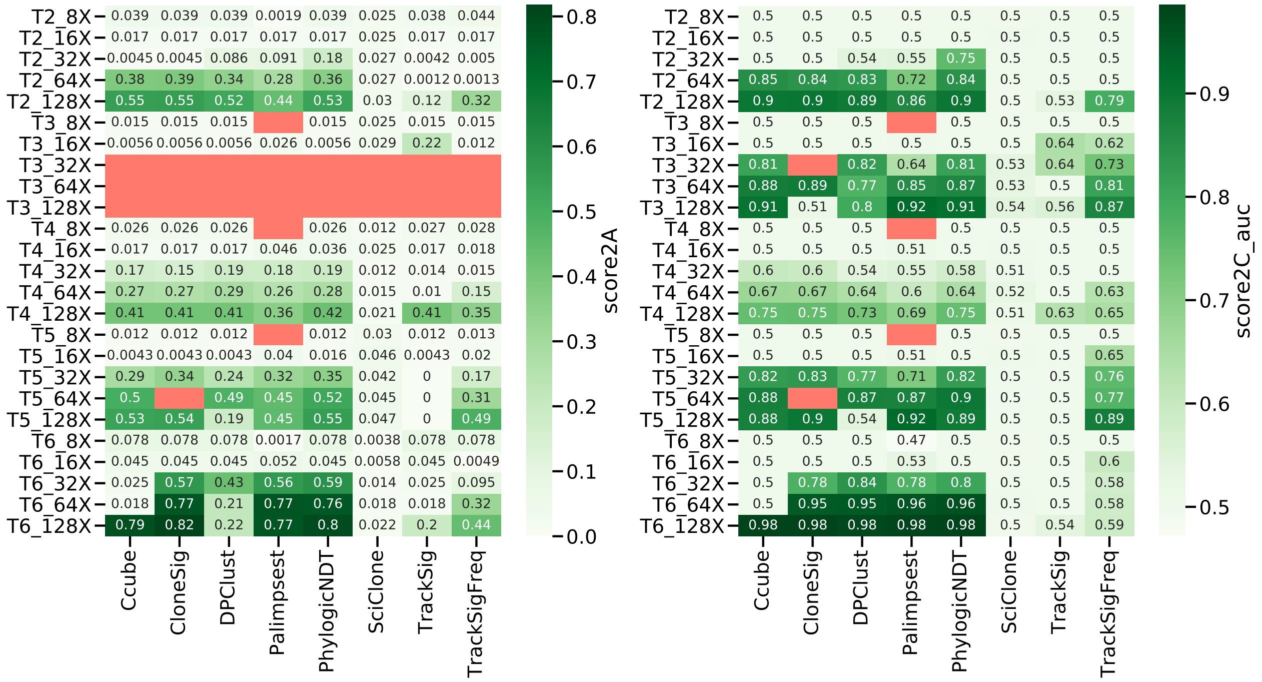

Supplementary Figure 30: **Complete results for the DREAM dataset, score2A and score2C AUC.** Source data are provided as a Source Data file.

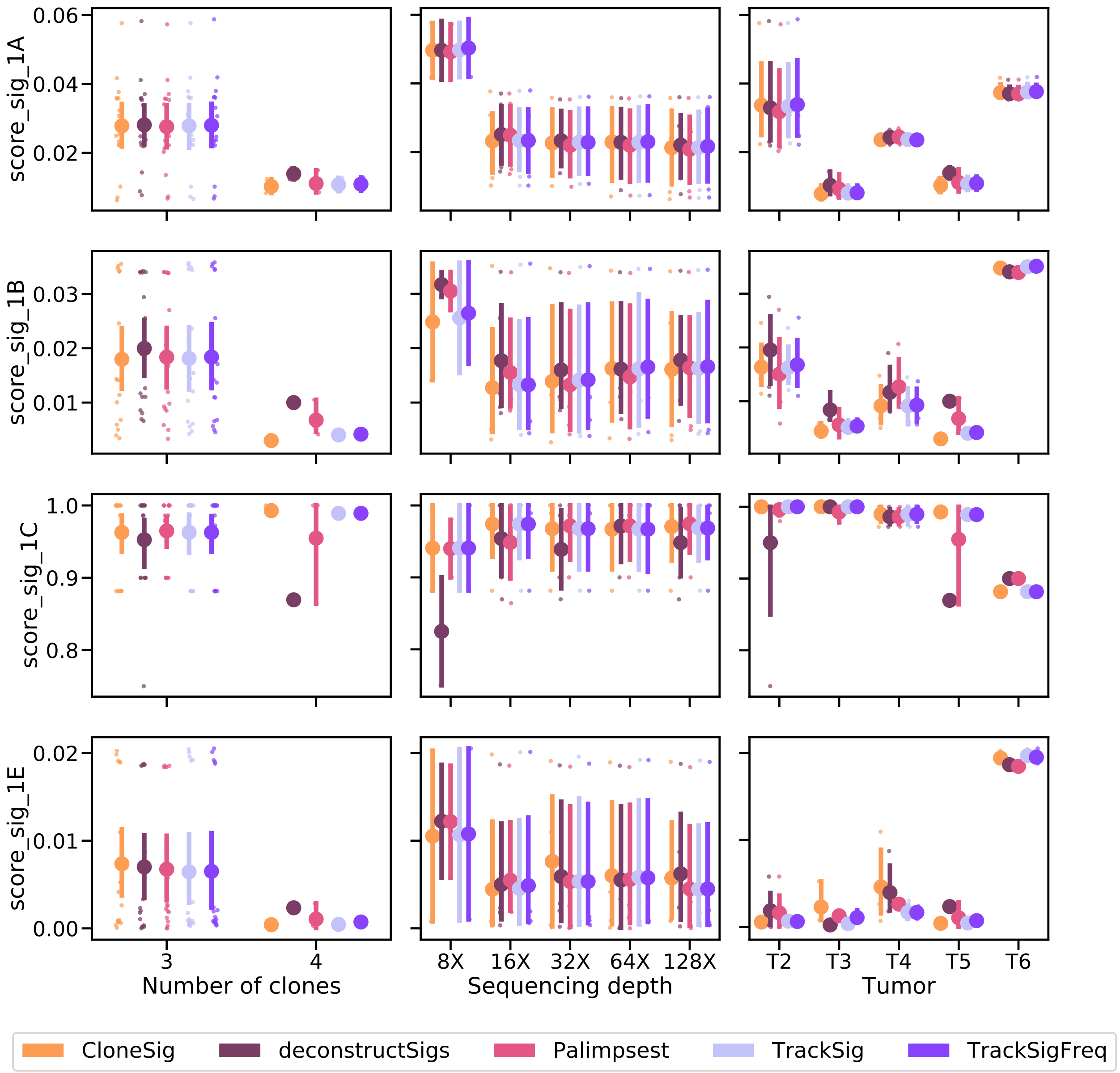

Supplementary Figure 31: **Comparison of CloneSig, TrackSig, TrackSigFreq, Palimpsest and deconstructSigs for signature activity deconvolution on the DREAM dataset.** Scores_sig_1A and 1B are distances between the estimated mutation type profile (as defined by the signature activity proportions) to the true mutation profile (defined using the parameters used for simulations, 1A), and the empirical observed mutation profile (defined using available observed mutations, 1B), and is better when close to 0. Score_sig_1C is the area under the ROC curve for the classification of signatures as active or inactive in the sample, and is better when close to 1. Finally, Score_sig_1E is the median distance to the true mutation type profile of the clone to which a mutation was attributed from the true distribution of its original clone in the simulation, and is better when close to 0. The results are presented depending on several relevant covariates: the true number of clones (left), the sequencing depth (middle), and the diploid proportion of the genome (right). Each point represents the average of the score over all available simulated samples. Bootstrap sampling of the scores was used to compute 95% confidence intervals of the mean.Results for each setting are presented for samples for which all methods ran successfully; there are respectively 17 with 3 clones and 3 samples with 4 clones, 2, 5, 4, 4 and 5 samples with a depth of 8X, 16X, 32X, 64X and 128X, and 5, 3, 4, 3 and 5 samples for tumors T2, T3, T4, T5 and T6. Source data are provided as a Source Data file.

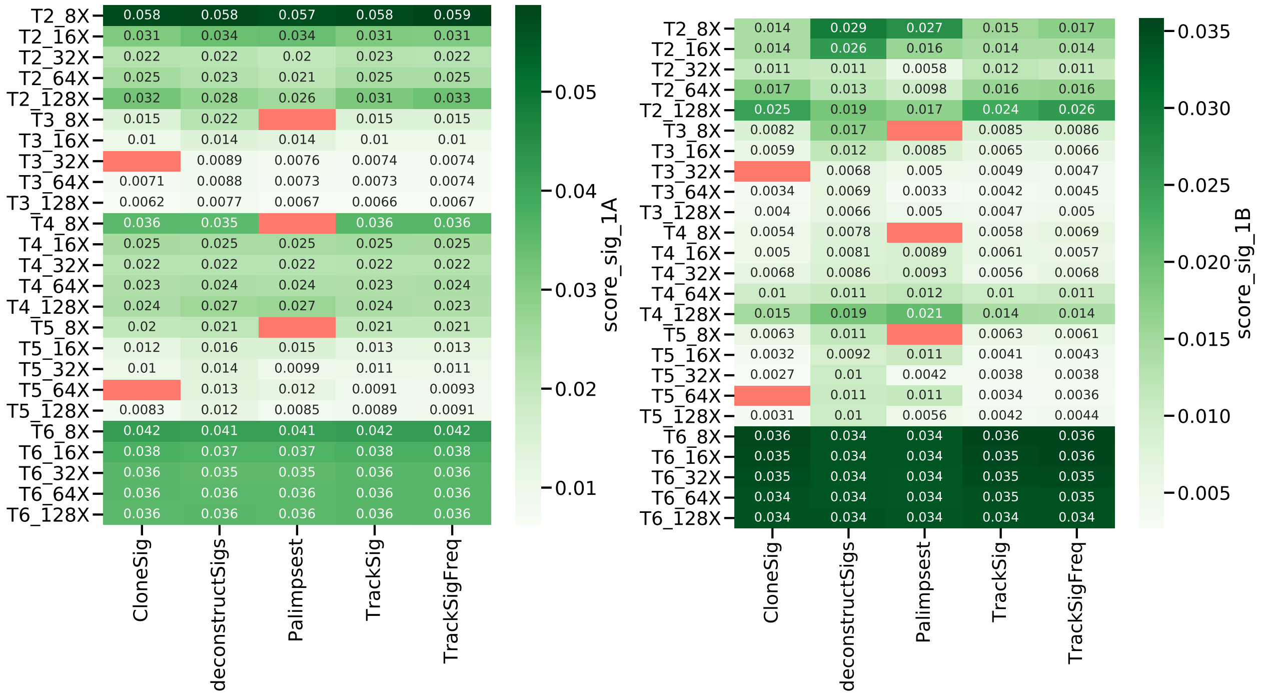

Supplementary Figure 32: **Complete results for the DREAM dataset, score_sig_1A and score_sig_1B.** Source data are provided as a Source Data file.

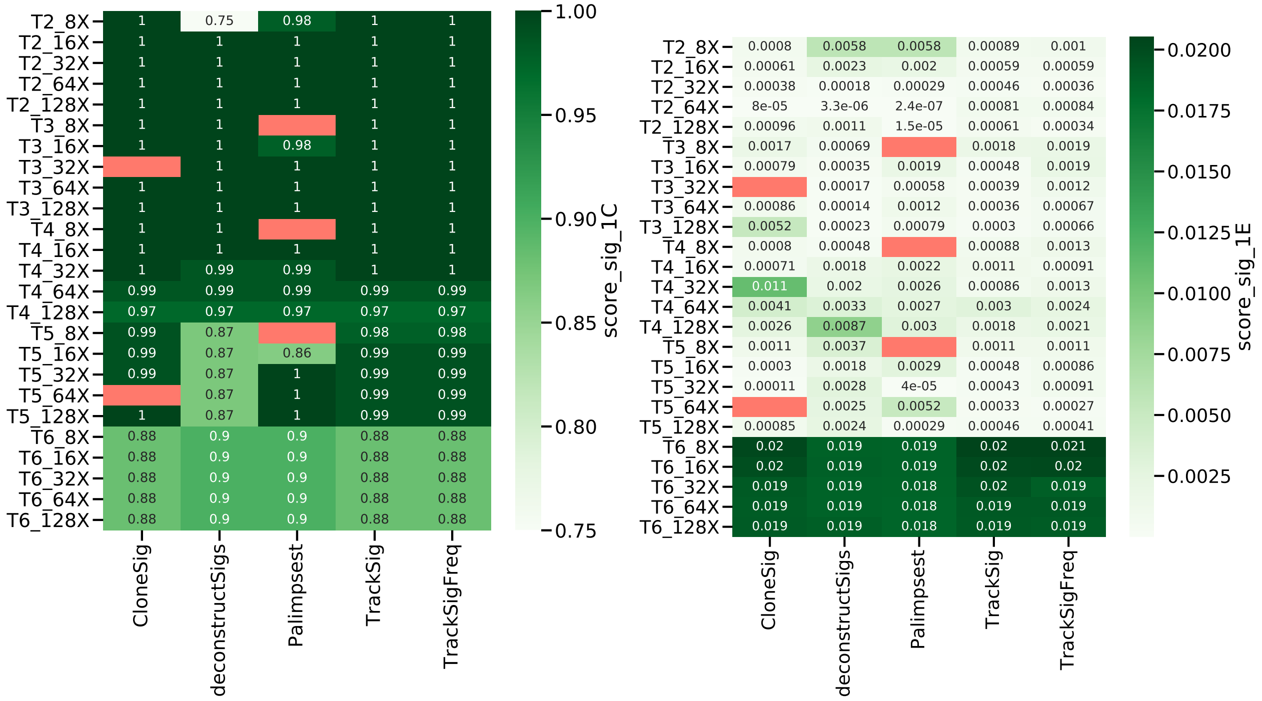

Supplementary Figure 33: **Complete results for the DREAM dataset, score_sig_1C and score_sig_1E.** Source data are provided as a Source Data file.

#### 2.3 SimClone1000 dataset

We tested all evaluated methods on the SimClone1000 simulated dataset generated in [16, 17], for samples with less than 10,000 mutations. Results for samples in which all methods ran successfully are presented in Figure 34 for the subclonal reconstruction task, and in Figure 35 for the signature activity deconvolution task. This illustrates CloneSig's competitive performances for evaluated tasks on an independent dataset.

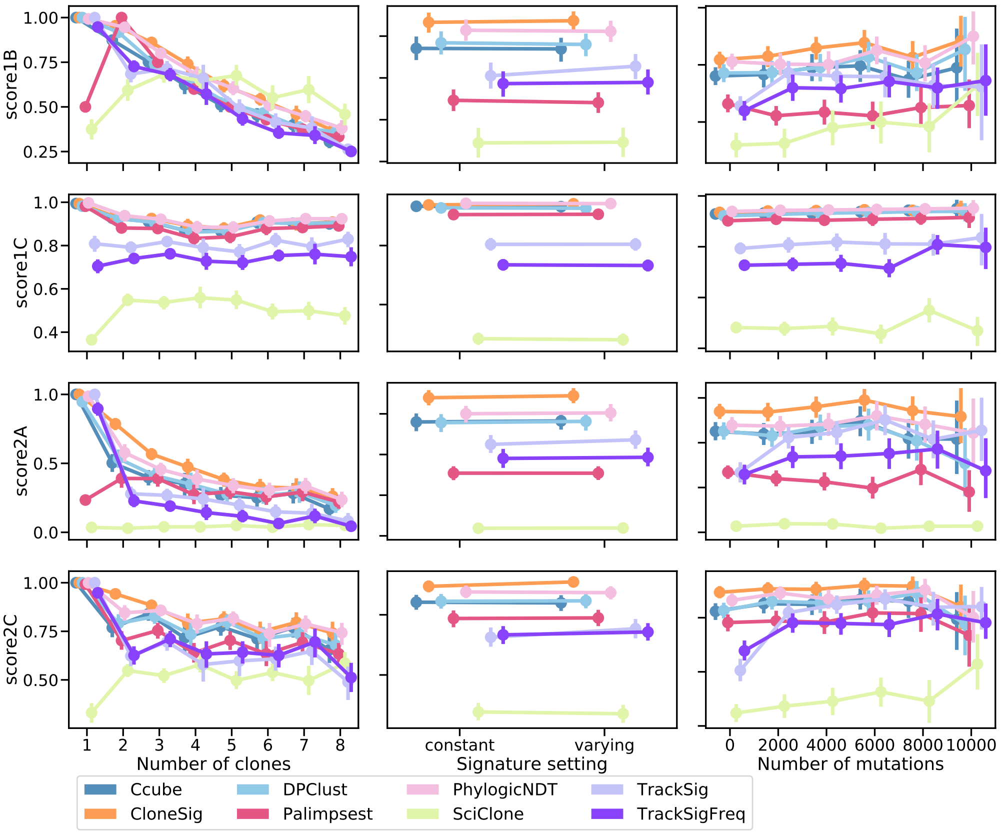

Supplementary Figure 34: **Comparison of CloneSig, TrackSig, TrackSigFreq, Palimpsest, SciClone, DPClust, PhylogicNDT and Ccube for subclonal reconstruction on the SimClone1000 dataset.** Each row corresponds to one score, as detailed in the main text. All scores are normalized between 0 and 1, with 1 being the best and 0 the worst. Each column corresponds to a setting where one parameter in the simulation varies: the true number of clones (left), the simulation setting (middle), and the number of mutations (right). Each point represents the average of the score over all available simulated samples. Bootstrap sampling of the scores was used to compute 95% confidence intervals of the mean. Score1B evaluates the number of clones found by the method, score1C the resulting mutation CCF distribution, score2A the co-clustering of mutations in the defined clones, and score2C the classification of subclonal versus clonal mutations. We have ensured that all scores are comparable by removing samples where at least one method fails for all other methods. As a result, we considered respectively 264, 190, 171, 43, 77, 50, 36 and 47 samples with 1, 2, 3, 4, 5, 6, 7 and 8 clones, 444 samples with the constant signature, and 434 with the varying setting, and 373, 216, 132, 94, 50 and 13 samples with a number of mutations closest to 0, 2000, 4000, 6000, 8000 or 10000. Source data are provided as a Source Data file.

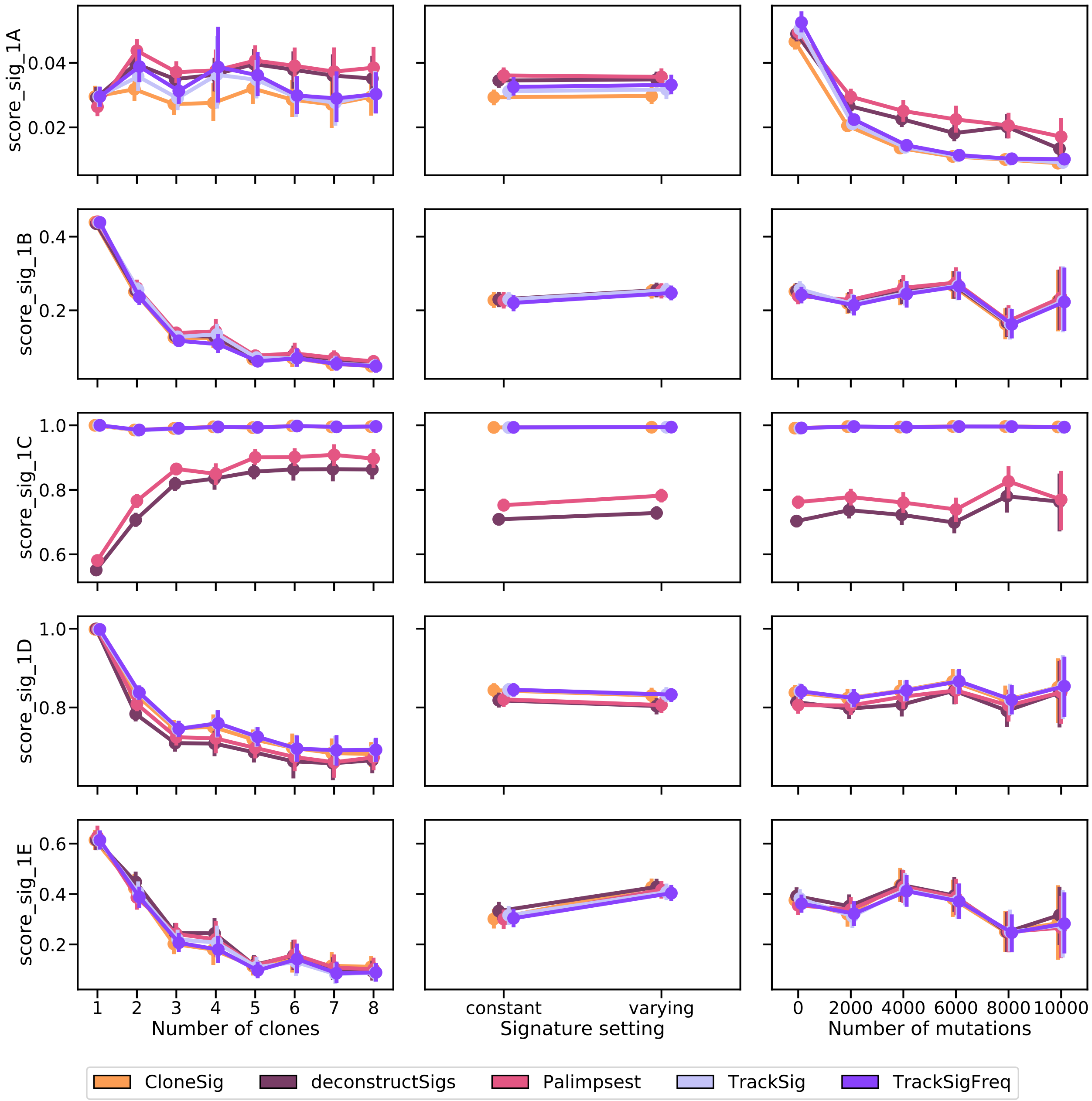

Supplementary Figure 35: **Comparison of CloneSig, TrackSig, TrackSigFreq, Palimpsest, and deconstructSigs for signature activity deconvolution on the SimClone1000 dataset.** Several metrics have been implemented, and are detailed in the main text. Scores_sig_1A and 1B are distances between the estimated mutation type profile (as defined by the signature activity proportions) to the true mutation profile (defined using the parameters used for simulations, 1A), and the empirical observed mutation profile (defined using available observed mutations, 1B), and is better when close to 0. Score_sig_1C is the area under the ROC curve for the classification of signatures as active or inactive in the sample, and is better when close to 1. Score_sig_1D is the proportion of mutations for which the correct signature was attributed, and is better when close to 1. Finally, Score_sig_1E is the median distance to the true mutation type profile of the clone to which a mutation was attributed from the true distribution of its original clone in the simulation, and is better when close to 0. The results are presented depending on several relevant covariates: the true number of clones (left), the simulation setting (middle), and the number of mutations (right). Each point represents the average of the score over all available simulated samples. Bootstrap sampling of the scores was used to compute 95% confidence intervals of the mean. We have ensured that all scores are comparable by removing samples where at least one method fails for all other methods. As a result, we considered respectively 314, 201, 182, 49, 77, 52, 38 and 49 samples with 1, 2, 3, 4, 5, 6, 7 and 8 clones, 487 samples with the constant signature, and 475 with the varying setting, and 433, 226, 134, 103, 53 and 13 samples with a number of mutations closest to 0, 2000, 4000, 6000, 8000 or 10000. Source data are provided as a Source Data file.

### Supplementary Note 3: Complete overview of TCGA results

We now use CloneSig on real data, to analyze ITH and mutational process changes in a large cohort of 8,951 tumor WES samples from the TCGA spanning 31 cancer types. An overview of the main characteristics of the cohort is presented in Table 3.

For each sample in the cohort, we estimate with CloneSig the number of subclones present in the tumor, the signatures active in each subclone, and test for the presence of a signature change between clones. Figure 36 shows a global summary of the signature changes found in the cohort. For each cancer type, it shows the proportion of samples where a signature change is found, and a visual summary of the proportion of samples where each individual signature is found to increase or to decrease in the largest subclone, compared to the clonal mutations. The thickness of each bar, in addition, indicates the median change of each signature.

A multivariate Cox model was fitted to the TCGA data to compare three groups of tumors from the TCGA: homogeneous tumors with one clone, tumors with multiple clones but no significant signature change, and tumors with multiple clones and a significant signature change, with a stratification with respect to the tumor type. It indicates for 2 clones a hazard ratio (HR) of 1.02 (95% confidence interval (CI): $[0.91, 1.13]$, $p = 0.74$), and for 3 clones a HR of 0.97 (CI= $[0.87, 1.09]$, $p = 0.65$), hence no significant difference in survival between the three groups.

Finally, detailed results on each cancer type are presented in Supplementary Figures 38 to 68.

We additionally generated similar plots using a log-fold change in Figure 37, a relative variation of signature activity measure.

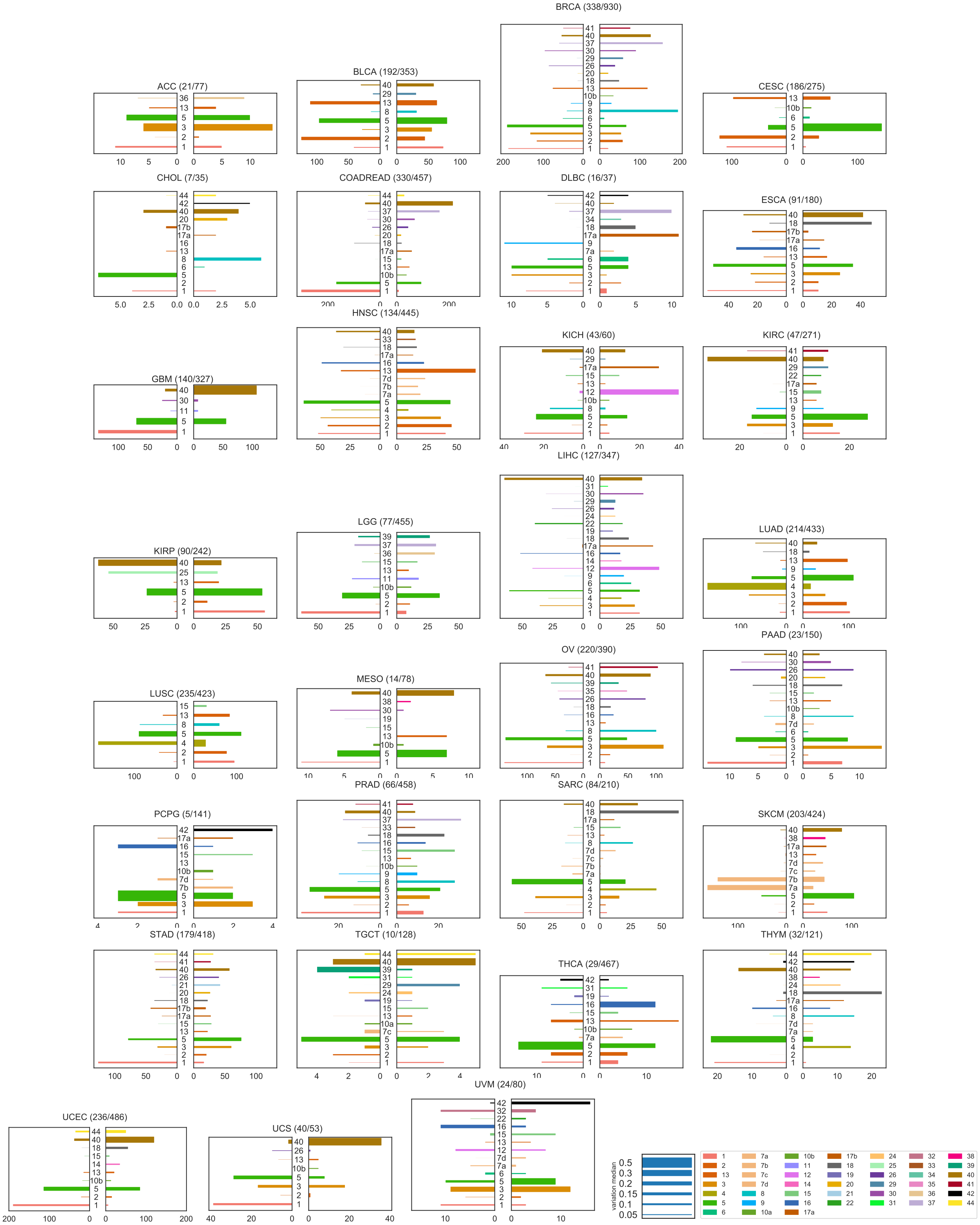

Supplementary Figure 36: **Mutational signature changes in the TCGA cohort.** Each plot corresponds to one cancer type, indicates the number of samples with a significant signature change compared to the total number of samples, and shows on the right panel an increase of a signature in the largest subclone, compared to clonal mutations, and on the left panel a decrease. The length of each bar corresponds to the number of patients with such changes, and the thickness to the median absolute observed change. Source data are provided as a Source Data file.

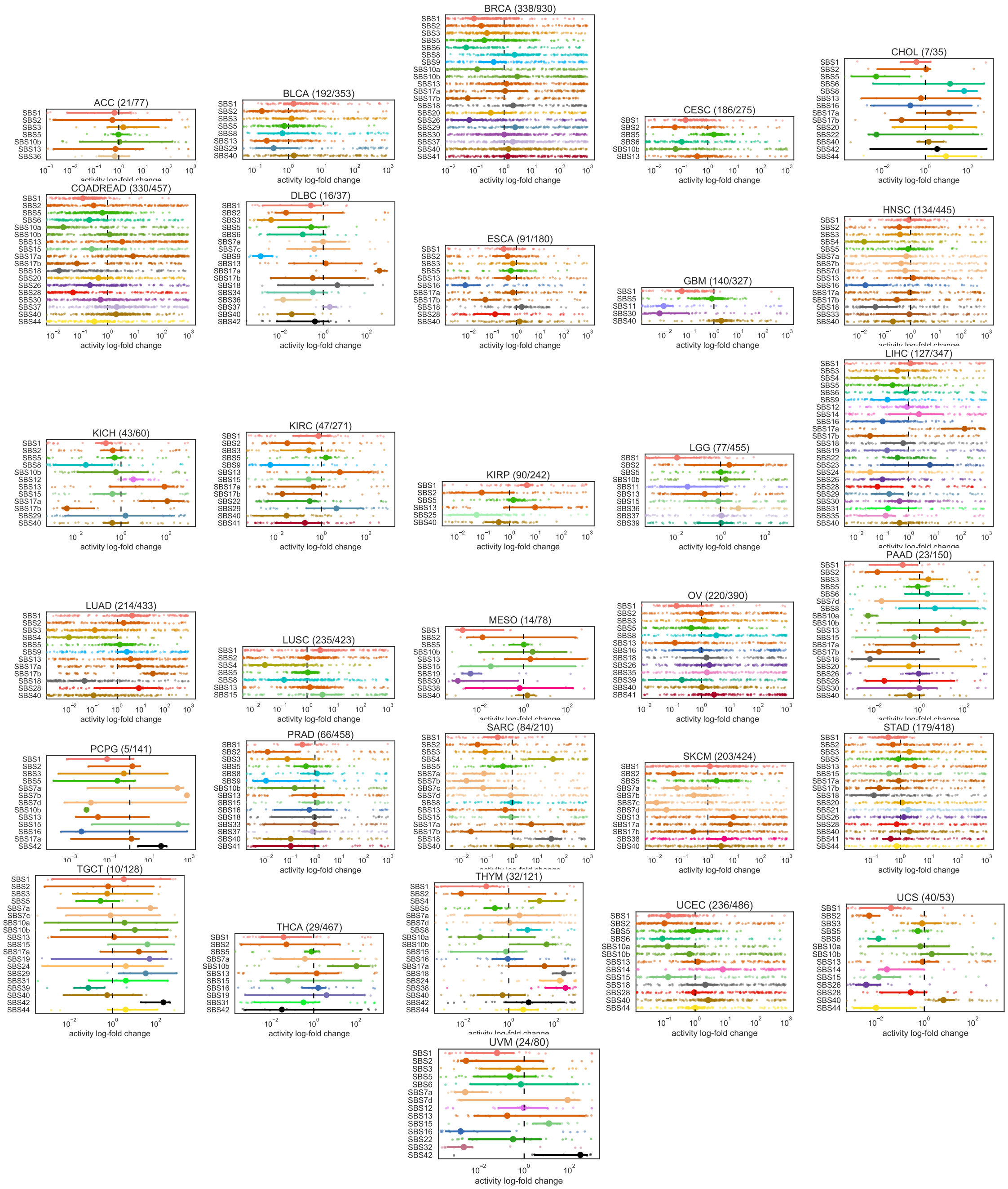

Supplementary Figure 37: **Mutational signature changes in the TCGA cohort.** Each plot corresponds to one cancer type, indicates the number of samples with a significant signature change compared to the total number of samples, and shows log fold change of signature activity in the largest subclone, compared to clonal mutations. The small dots correspond to individual samples, and the larger dot to the median change. Bootstrap sampling of the scores was used to compute 95% confidence intervals of the median. The vertical dotted line in each plot represent no change. Source data are provided as a Source Data file.

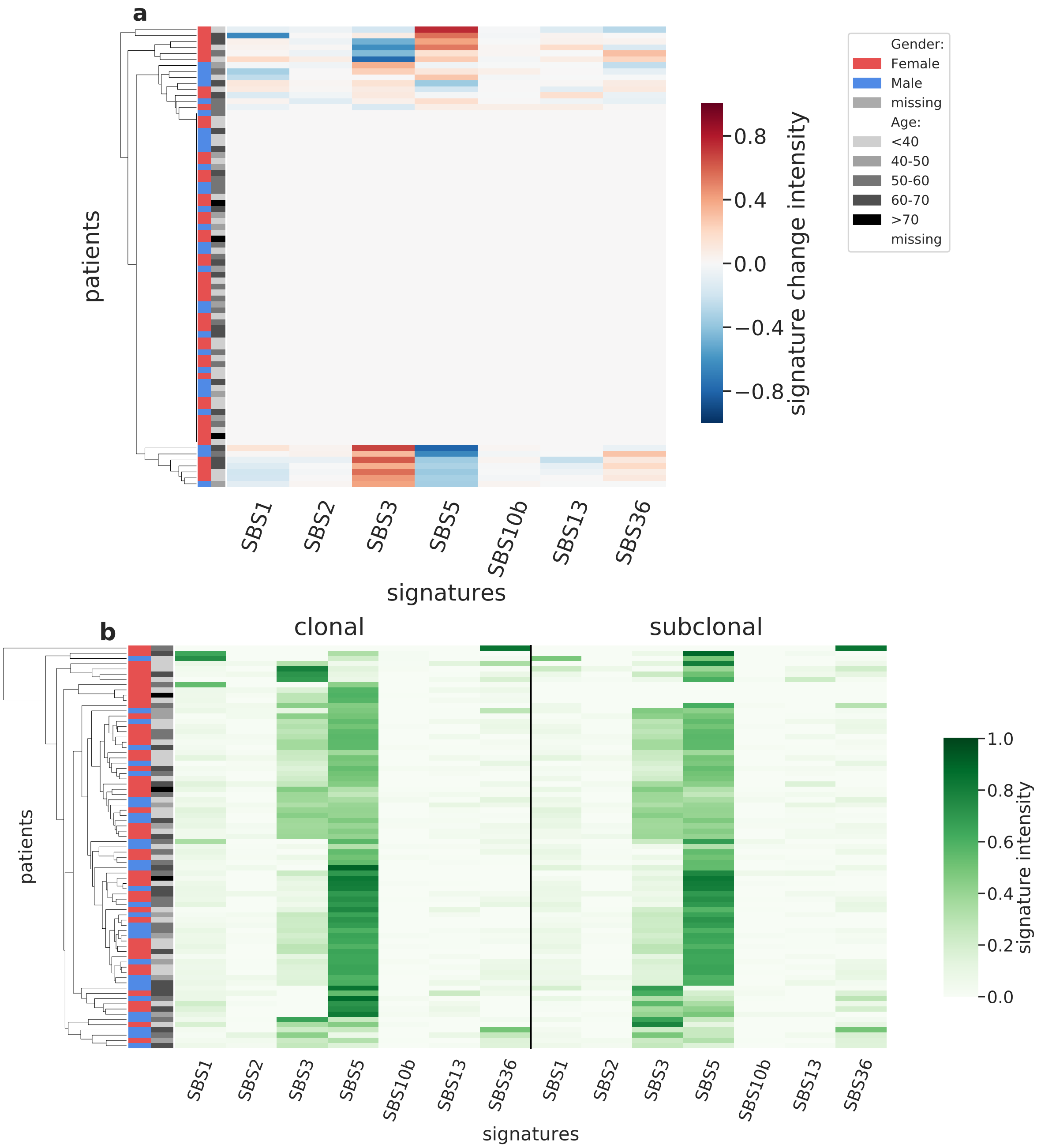

Supplementary Figure 38: **Overview of CloneSig results for the TCGA sub-cohort ACC (n=77).** Panel a: Stratification of patients depending on their pattern of signature change for ACC patients (77 patients, including 21 with a significant signature change). The heatmap represents the difference between the signature activity in the largest subclone (in terms of number of mutations) and the clonal mutations (defined as belonging to the clone of highest CCF).Panel b: Stratification of patients depending on their complete pattern of signature exposure. The heatmap represents the signature activity in the largest subclone (in terms of number of mutations) and the clonal mutations (defined as belonging to the clone of highest CCF). Source data are provided as a Source Data file.

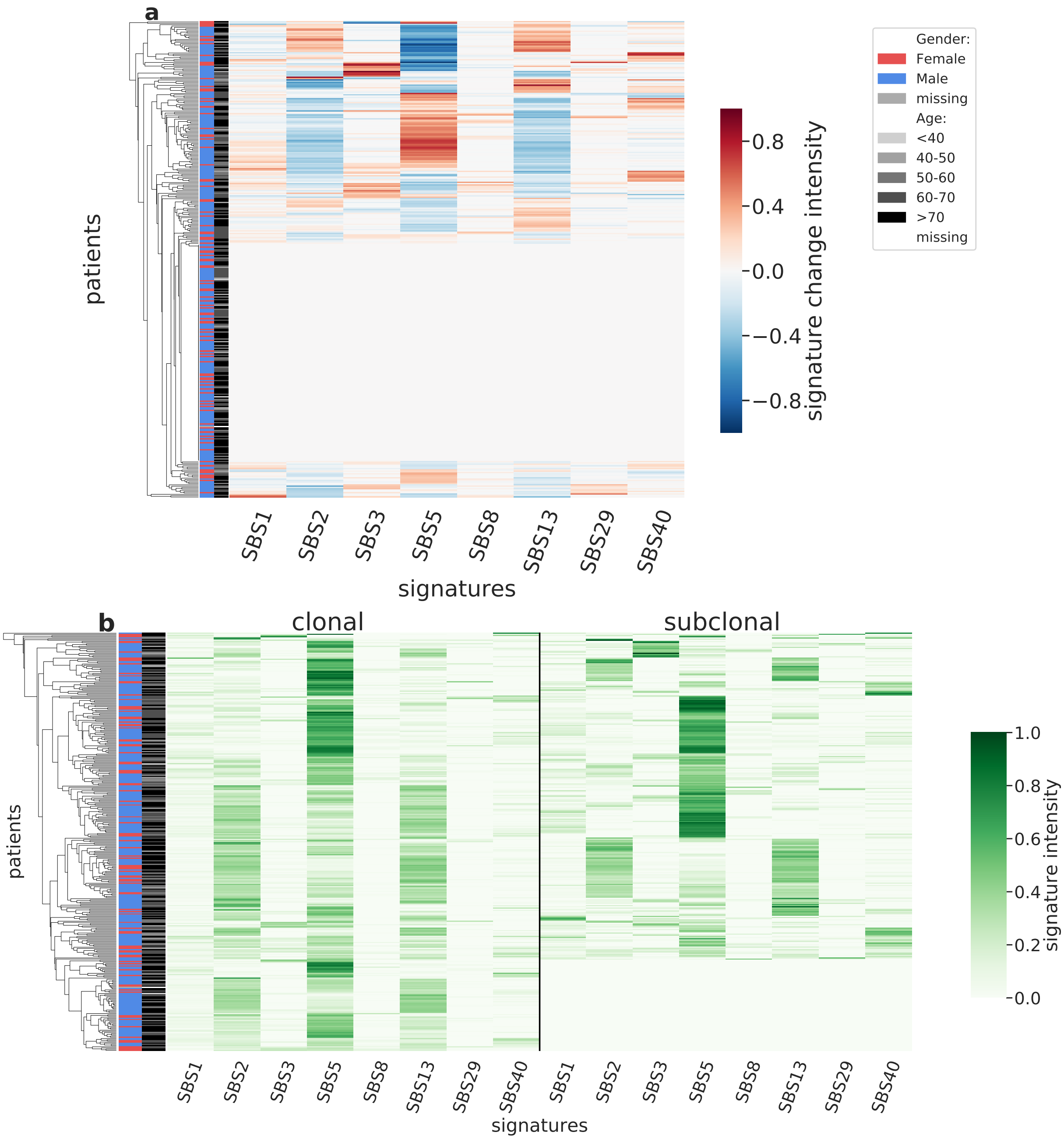

Supplementary Figure 39: **Overview of CloneSig results for the TCGA sub-cohort BLCA (n=353).** Panel a: Stratification of patients depending on their pattern of signature change for BLCA patients (353 patients, including 192 with a significant signature change). The heatmap represents the difference between the signature activity in the largest subclone (in terms of number of mutations) and the clonal mutations (defined as belonging to the clone of highest CCF).Panel b: Stratification of patients depending on their complete pattern of signature exposure. The heatmap represents the signature activity in the largest subclone (in terms of number of mutations) and the clonal mutations (defined as belonging to the clone of highest CCF). Source data are provided as a Source Data file.

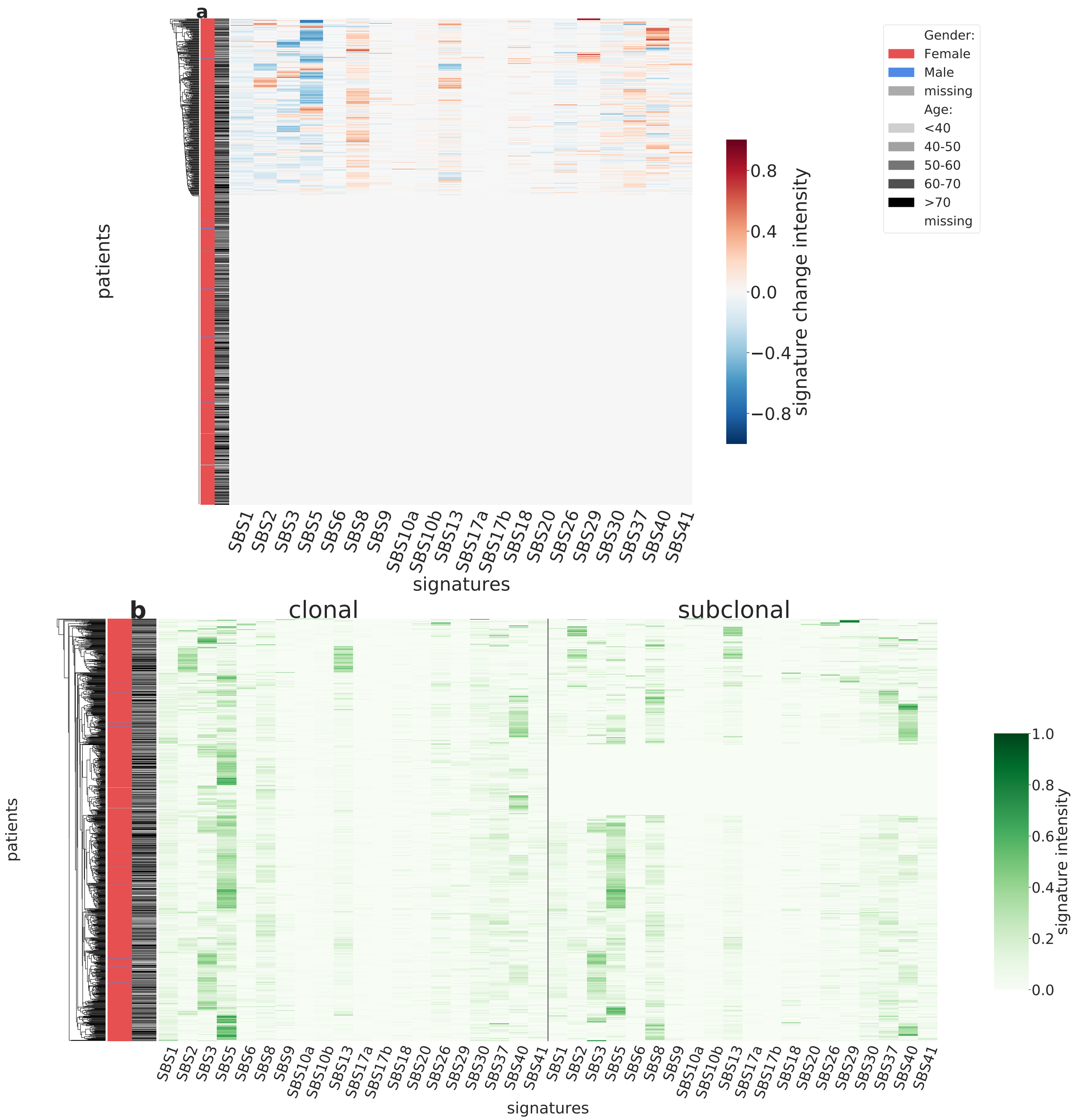

Supplementary Figure 40: **Overview of CloneSig results for the TCGA sub-cohort BRCA (n=930).** Panel a: Stratification of patients depending on their pattern of signature change for BRCA patients (930 patients, including 338 with a significant signature change). The heatmap represents the difference between the signature activity in the largest subclone (in terms of number of mutations) and the clonal mutations (defined as belonging to the clone of highest CCF).Panel b: Stratification of patients depending on their complete pattern of signature exposure. The heatmap represents the signature activity in the largest subclone (in terms of number of mutations) and the clonal mutations (defined as belonging to the clone of highest CCF). Source data are provided as a Source Data file.

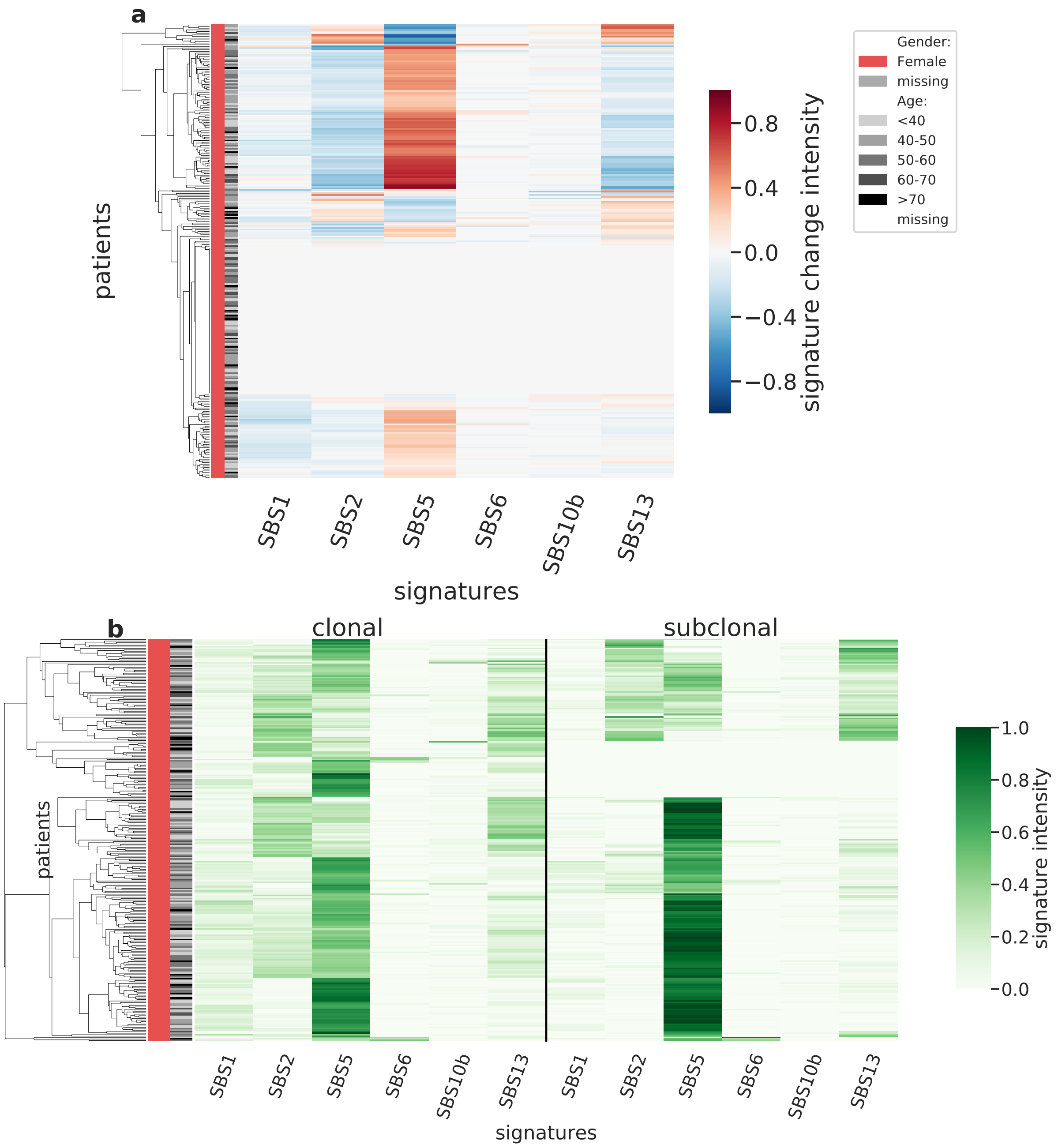

Supplementary Figure 41: **Overview of CloneSig results for the TCGA sub-cohort CESC (n=275).** Panel a: Stratification of patients depending on their pattern of signature change for CESC patients (275 patients, including 186 with a significant signature change). The heatmap represents the difference between the signature activity in the largest subclone (in terms of number of mutations) and the clonal mutations (defined as belonging to the clone of highest CCF).Panel b: Stratification of patients depending on their complete pattern of signature exposure. The heatmap represents the signature activity in the largest subclone (in terms of number of mutations) and the clonal mutations (defined as belonging to the clone of highest CCF). Source data are provided as a Source Data file.

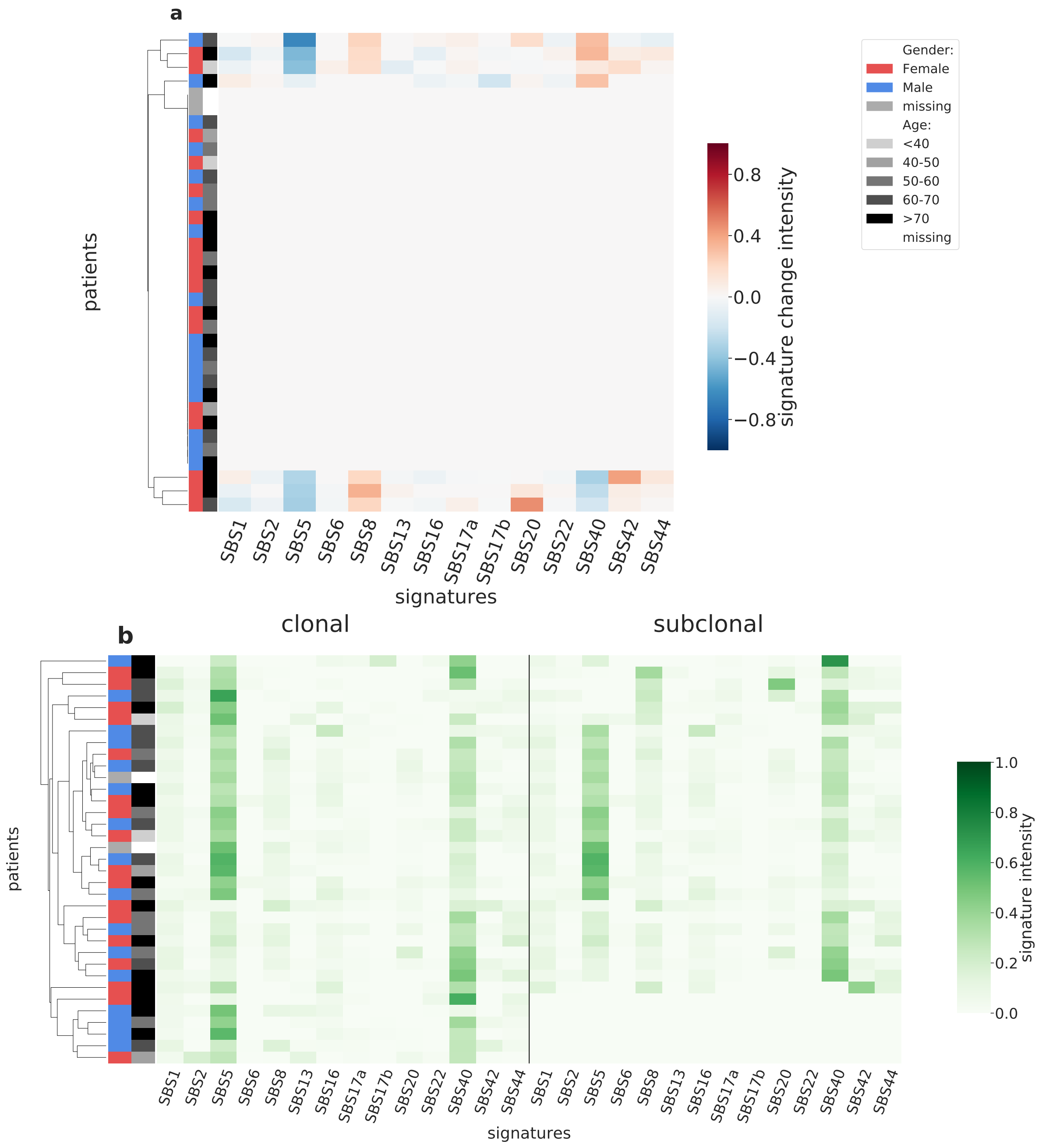

Supplementary Figure 42: **Overview of CloneSig results for the TCGA sub-cohort CHOL (n=35).** Panel a: Stratification of patients depending on their pattern of signature change for CHOL patients (35 patients, including 7 with a significant signature change). The heatmap represents the difference between the signature activity in the largest subclone (in terms of number of mutations) and the clonal mutations (defined as belonging to the clone of highest CCF).Panel b: Stratification of patients depending on their complete pattern of signature exposure. The heatmap represents the signature activity in the largest subclone (in terms of number of mutations) and the clonal mutations (defined as belonging to the clone of highest CCF). Source data are provided as a Source Data file.

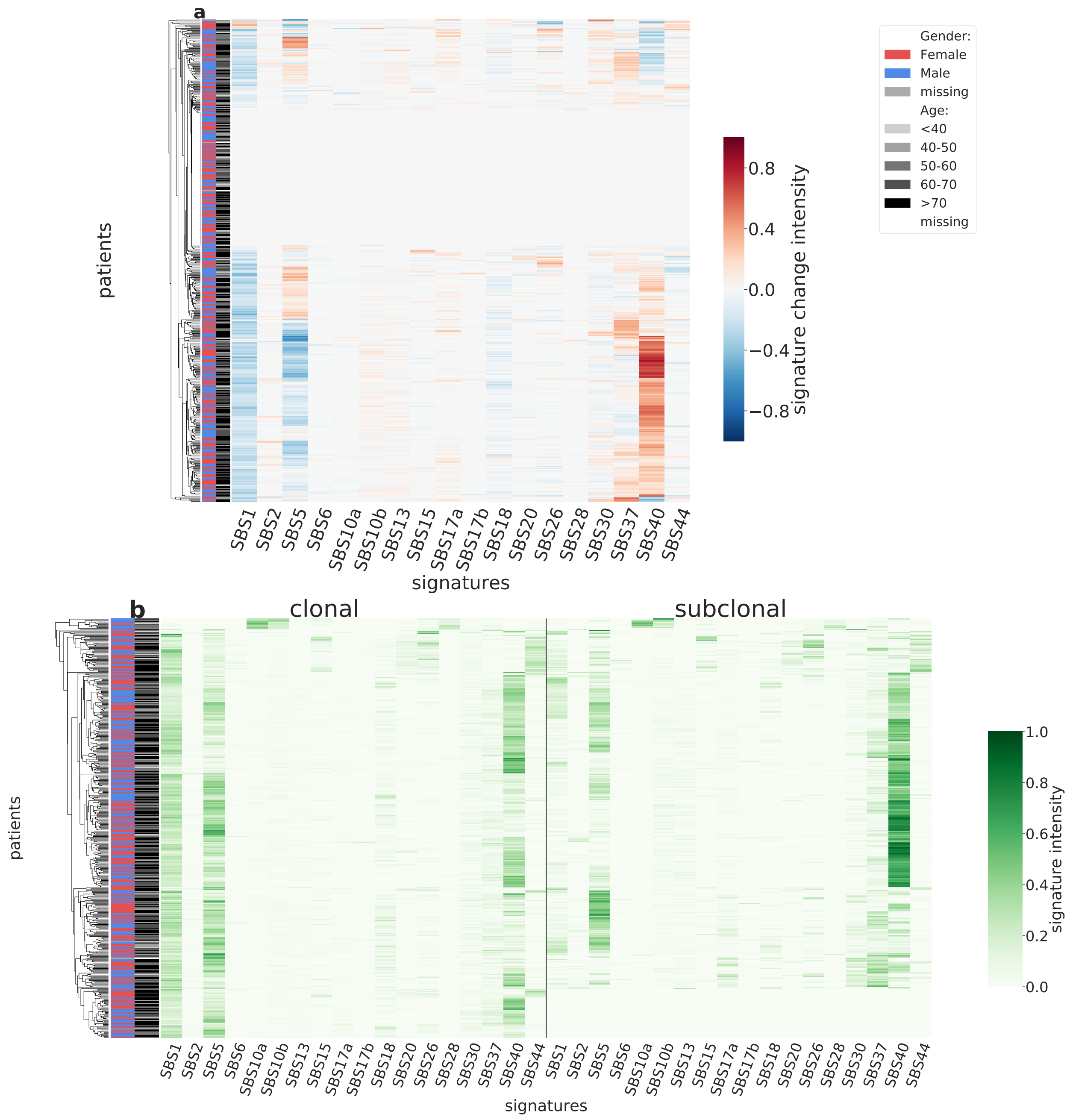

Supplementary Figure 43: **Overview of CloneSig results for the TCGA sub-cohort COADREAD (n=457).** Panel a: Stratification of patients depending on their pattern of signature change for COADREAD patients (457 patients, including 330 with a significant signature change). The heatmap represents the difference between the signature activity in the largest subclone (in terms of number of mutations) and the clonal mutations (defined as belonging to the clone of highest CCF).Panel b: Stratification of patients depending on their complete pattern of signature exposure. The heatmap represents the signature activity in the largest subclone (in terms of number of mutations) and the clonal mutations (defined as belonging to the clone of highest CCF). Source data are provided as a Source Data file.

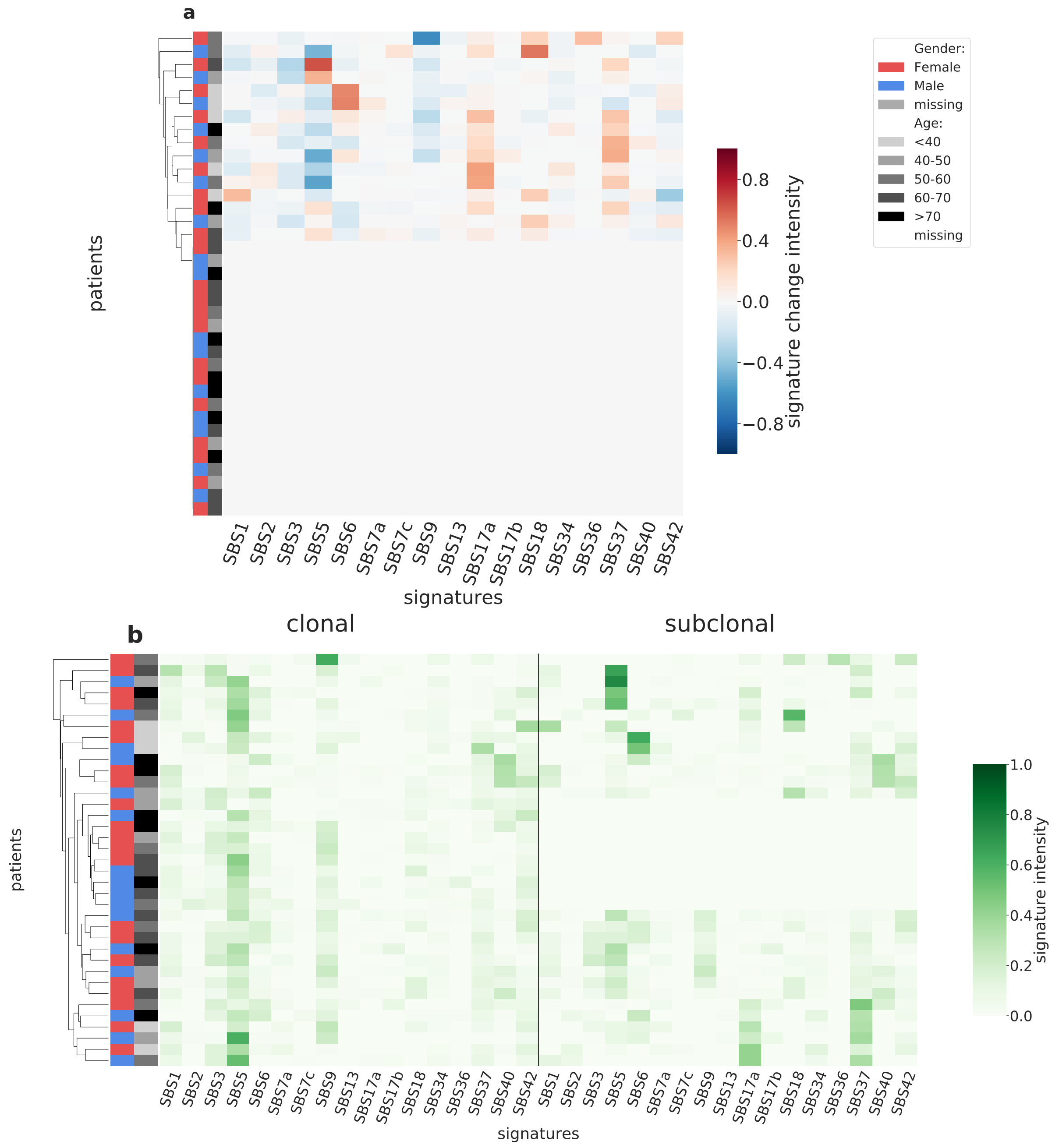

Supplementary Figure 44: **Overview of CloneSig results for the TCGA sub-cohort DLBC (n=37).** Panel a: Stratification of patients depending on their pattern of signature change for DLBC patients (37 patients, including 16 with a significant signature change). The heatmap represents the difference between the signature activity in the largest subclone (in terms of number of mutations) and the clonal mutations (defined as belonging to the clone of highest CCF).Panel b: Stratification of patients depending on their complete pattern of signature exposure. The heatmap represents the signature activity in the largest subclone (in terms of number of mutations) and the clonal mutations (defined as belonging to the clone of highest CCF). Source data are provided as a Source Data file.

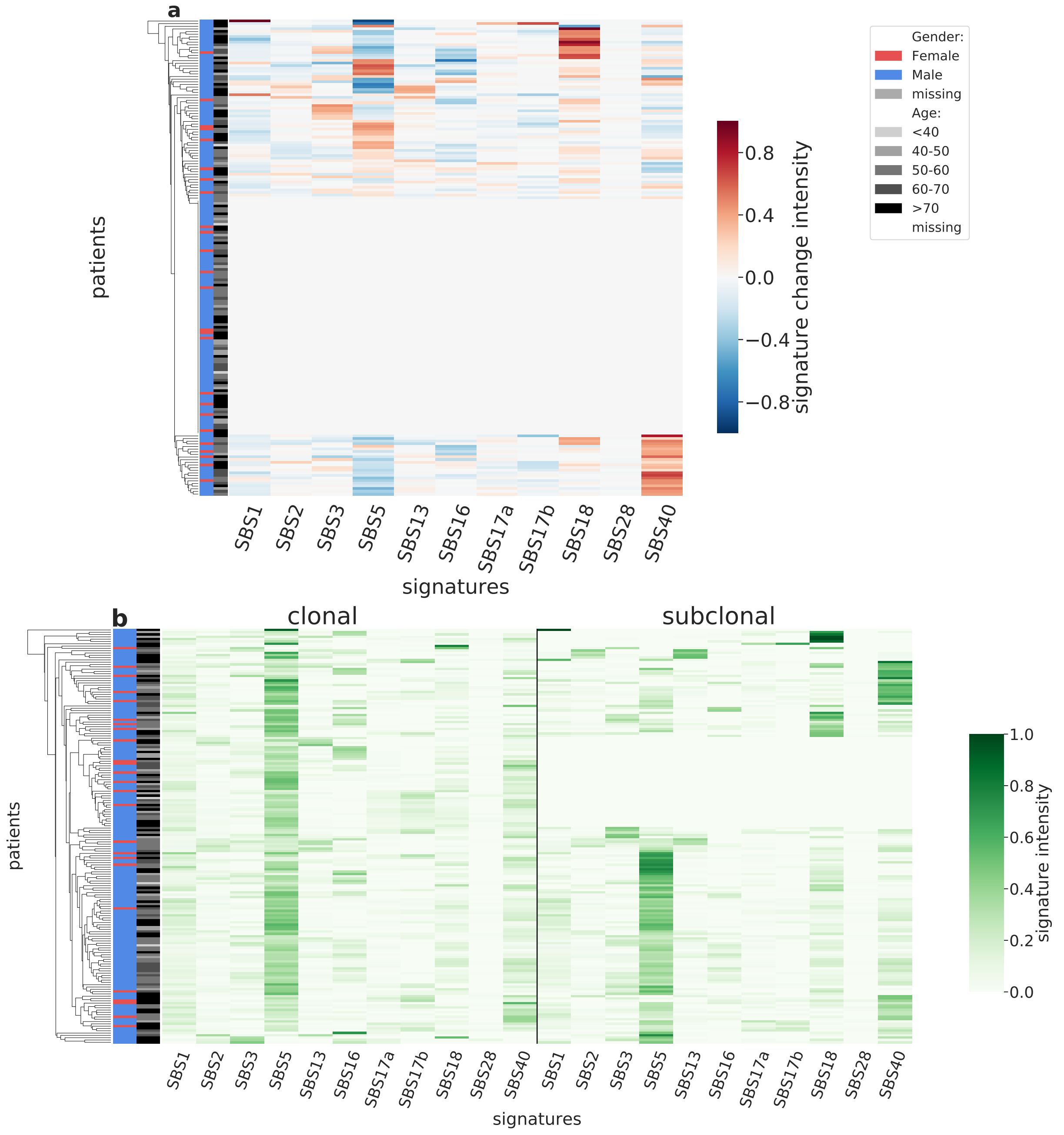

Supplementary Figure 45: **Overview of CloneSig results for the TCGA sub-cohort ESCA (n=180).** Panel a: Stratification of patients depending on their pattern of signature change for ESCA patients (180 patients, including 91 with a significant signature change). The heatmap represents the difference between the signature activity in the largest subclone (in terms of number of mutations) and the clonal mutations (defined as belonging to the clone of highest CCF).Panel b: Stratification of patients depending on their complete pattern of signature exposure. The heatmap represents the signature activity in the largest subclone (in terms of number of mutations) and the clonal mutations (defined as belonging to the clone of highest CCF). Source data are provided as a Source Data file.

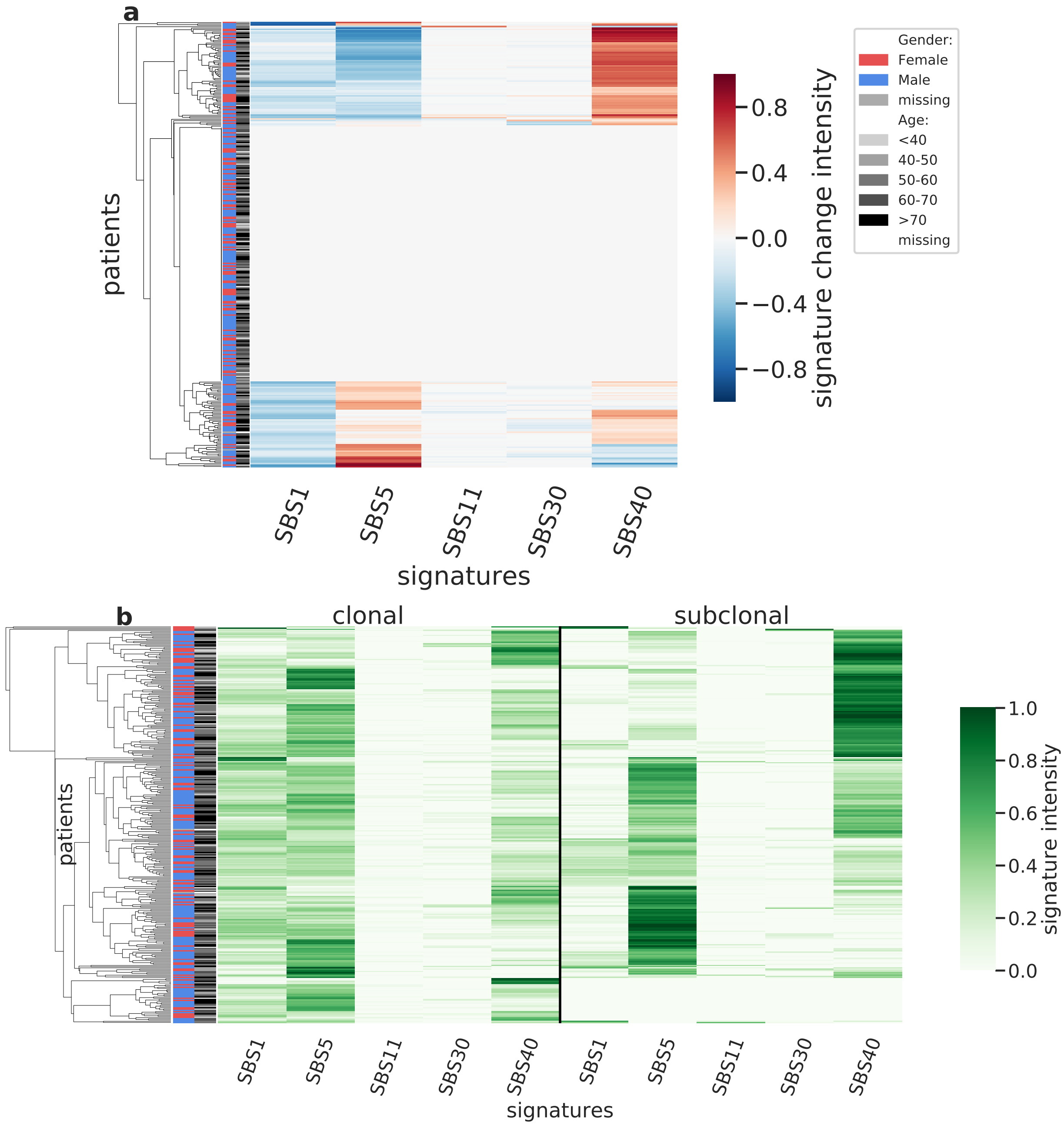

Supplementary Figure 46: **Overview of CloneSig results for the TCGA sub-cohort GBM (n=327).** Panel a: Stratification of patients depending on their pattern of signature change for GBM patients (327 patients, including 140 with a significant signature change). The heatmap represents the difference between the signature activity in the largest subclone (in terms of number of mutations) and the clonal mutations (defined as belonging to the clone of highest CCF).Panel b: Stratification of patients depending on their complete pattern of signature exposure. The heatmap represents the signature activity in the largest subclone (in terms of number of mutations) and the clonal mutations (defined as belonging to the clone of highest CCF). Source data are provided as a Source Data file.

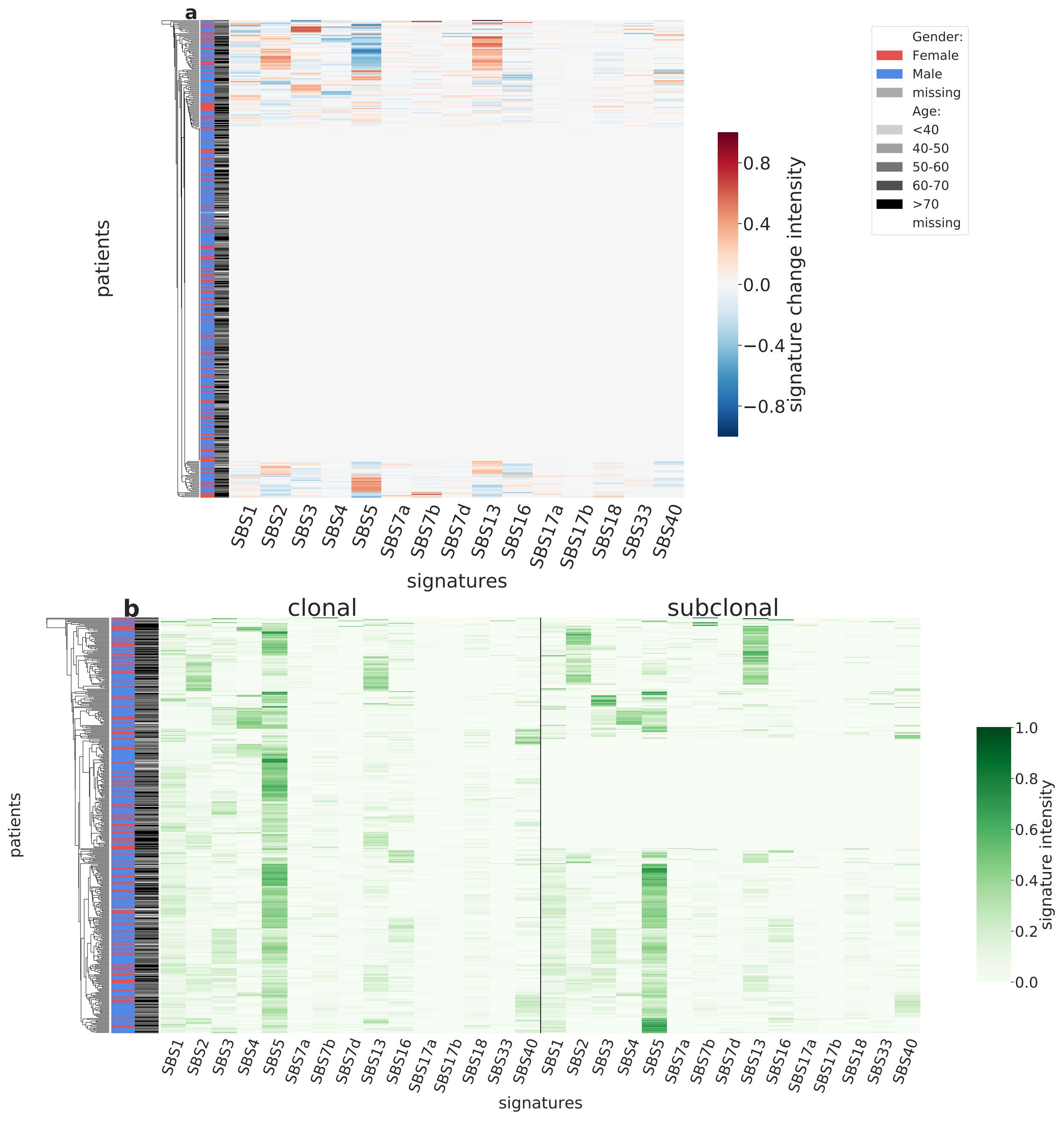

Supplementary Figure 47: **Overview of CloneSig results for the TCGA sub-cohort HNSC (n=445).** Panel a: Stratification of patients depending on their pattern of signature change for HNSC patients (445 patients, including 134 with a significant signature change). The heatmap represents the difference between the signature activity in the largest subclone (in terms of number of mutations) and the clonal mutations (defined as belonging to the clone of highest CCF).Panel b: Stratification of patients depending on their complete pattern of signature exposure. The heatmap represents the signature activity in the largest subclone (in terms of number of mutations) and the clonal mutations (defined as belonging to the clone of highest CCF). Source data are provided as a Source Data file.

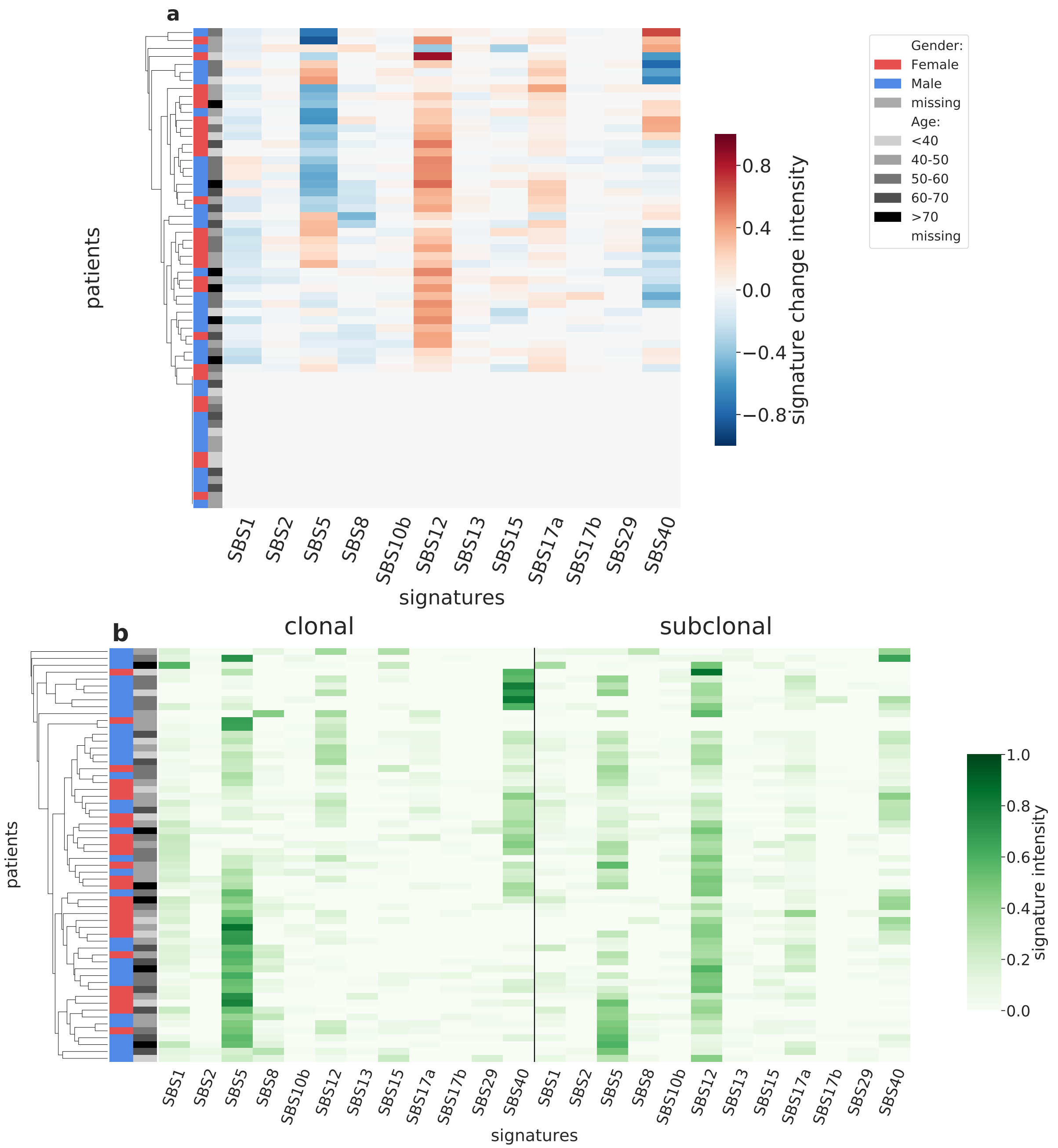

Supplementary Figure 48: **Overview of CloneSig results for the TCGA sub-cohort KICH (n=60).** Panel a: Stratification of patients depending on their pattern of signature change for KICH patients (60 patients, including 43 with a significant signature change). The heatmap represents the difference between the signature activity in the largest subclone (in terms of number of mutations) and the clonal mutations (defined as belonging to the clone of highest CCF).Panel b: Stratification of patients depending on their complete pattern of signature exposure. The heatmap represents the signature activity in the largest subclone (in terms of number of mutations) and the clonal mutations (defined as belonging to the clone of highest CCF). Source data are provided as a Source Data file.

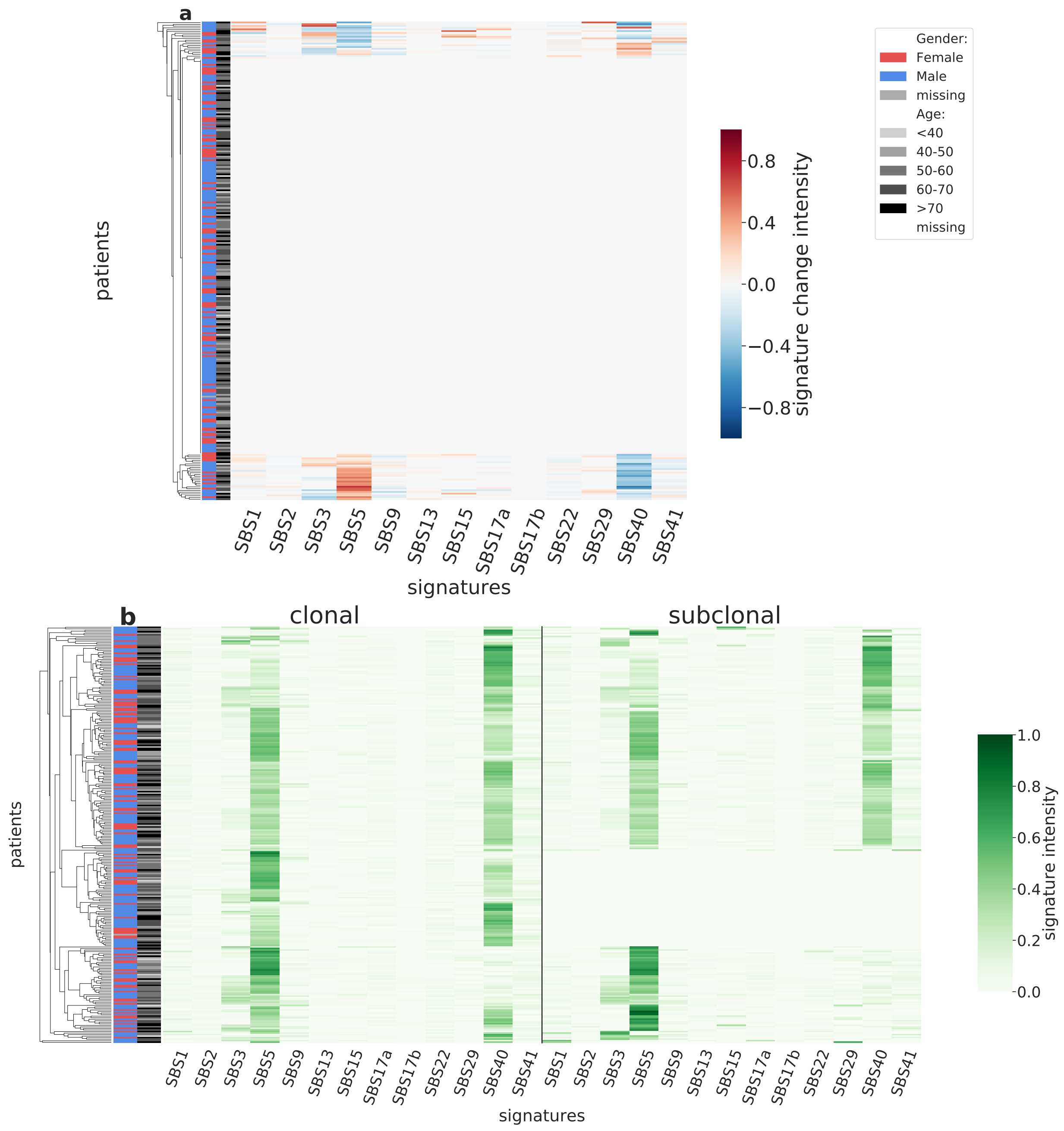

Supplementary Figure 49: **Overview of CloneSig results for the TCGA sub-cohort KIRC (n=271).** Panel a: Stratification of patients depending on their pattern of signature change for KIRC patients (271 patients, including 47 with a significant signature change). The heatmap represents the difference between the signature activity in the largest subclone (in terms of number of mutations) and the clonal mutations (defined as belonging to the clone of highest CCF).Panel b: Stratification of patients depending on their complete pattern of signature exposure. The heatmap represents the signature activity in the largest subclone (in terms of number of mutations) and the clonal mutations (defined as belonging to the clone of highest CCF). Source data are provided as a Source Data file.

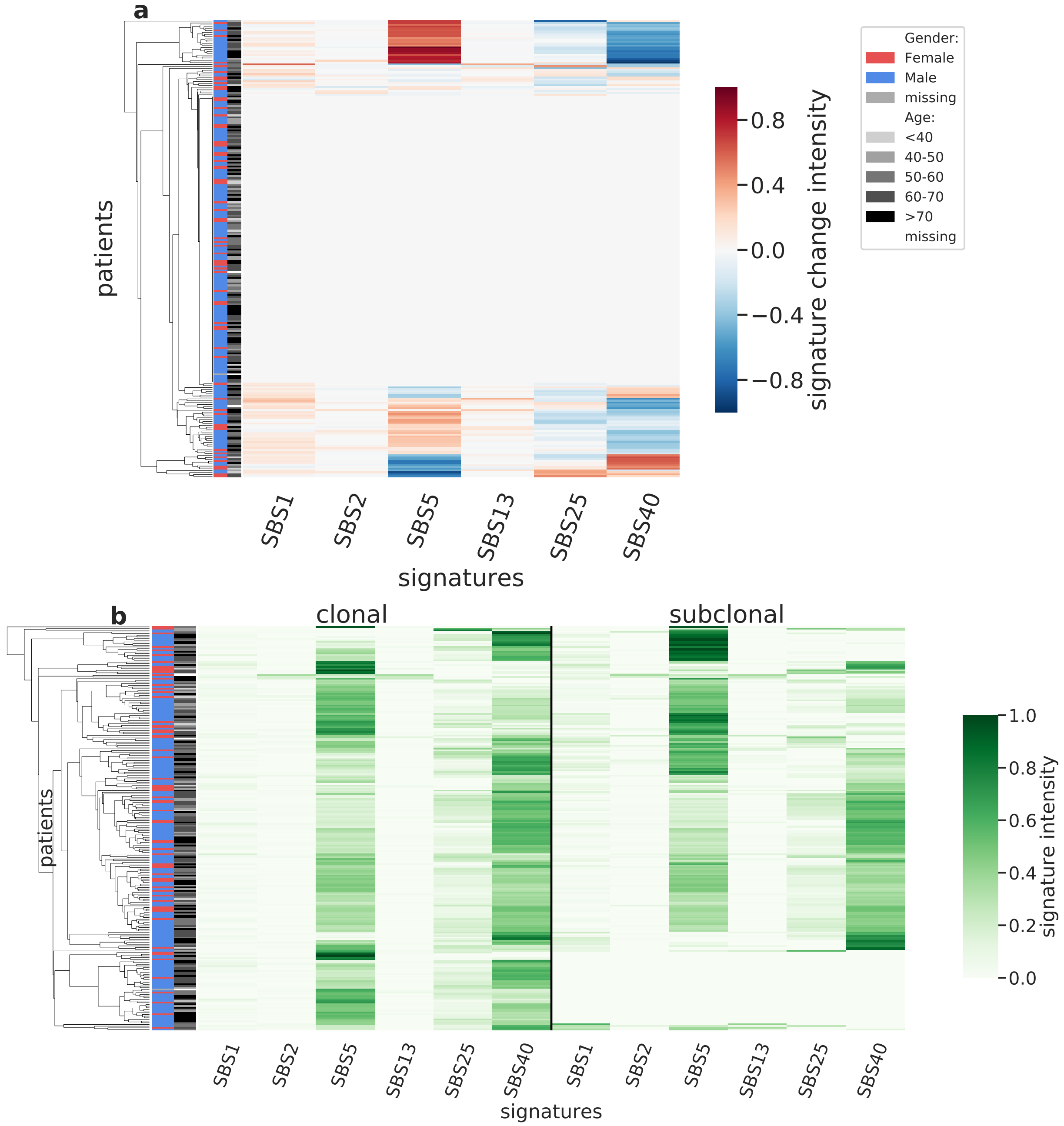

Supplementary Figure 50: **Overview of CloneSig results for the TCGA sub-cohort KIRP (n=242).** Panel a: Stratification of patients depending on their pattern of signature change for KIRP patients (242 patients, including 90 with a significant signature change). The heatmap represents the difference between the signature activity in the largest subclone (in terms of number of mutations) and the clonal mutations (defined as belonging to the clone of highest CCF).Panel b: Stratification of patients depending on their complete pattern of signature exposure. The heatmap represents the signature activity in the largest subclone (in terms of number of mutations) and the clonal mutations (defined as belonging to the clone of highest CCF). Source data are provided as a Source Data file.

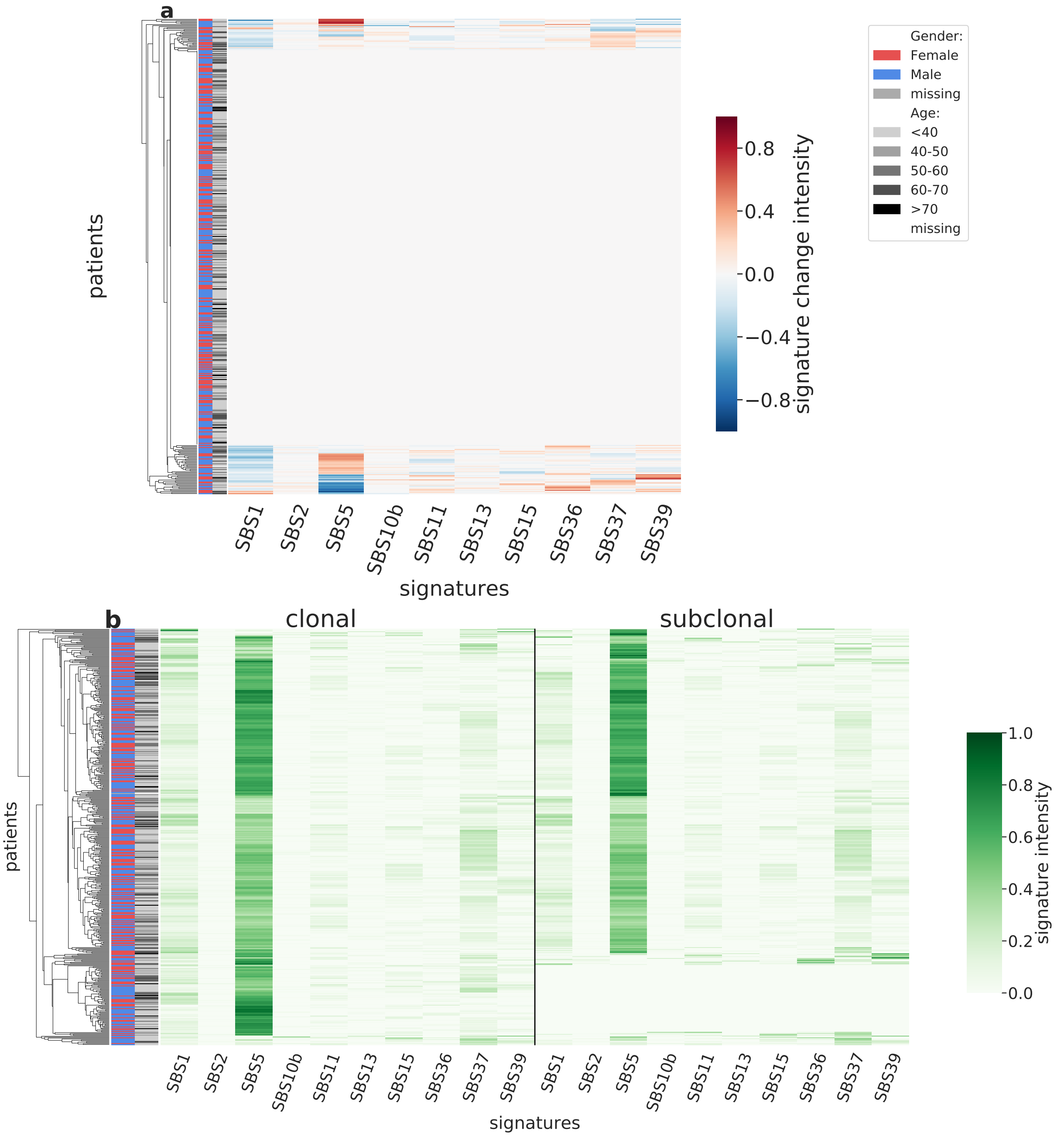

Supplementary Figure 51: **Overview of CloneSig results for the TCGA sub-cohort LGG (n=455).** Panel a: Stratification of patients depending on their pattern of signature change for LGG patients (455 patients, including 77 with a significant signature change). The heatmap represents the difference between the signature activity in the largest subclone (in terms of number of mutations) and the clonal mutations (defined as belonging to the clone of highest CCF).Panel b: Stratification of patients depending on their complete pattern of signature exposure. The heatmap represents the signature activity in the largest subclone (in terms of number of mutations) and the clonal mutations (defined as belonging to the clone of highest CCF). Source data are provided as a Source Data file.

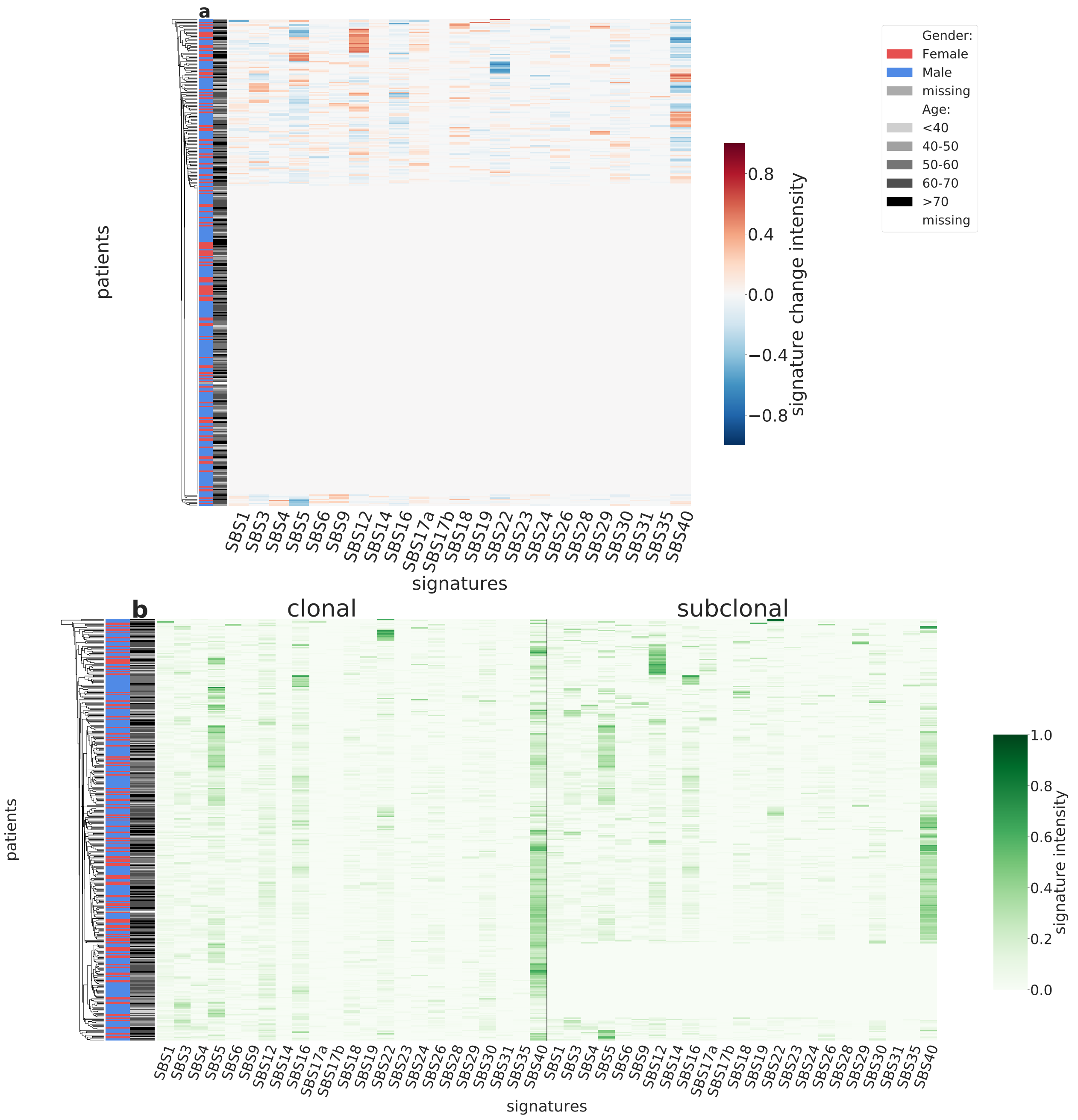

Supplementary Figure 52: **Overview of CloneSig results for the TCGA sub-cohort LIHC (n=347).** Panel a: Stratification of patients depending on their pattern of signature change for LIHC patients (347 patients, including 127 with a significant signature change). The heatmap represents the difference between the signature activity in the largest subclone (in terms of number of mutations) and the clonal mutations (defined as belonging to the clone of highest CCF).Panel b: Stratification of patients depending on their complete pattern of signature exposure. The heatmap represents the signature activity in the largest subclone (in terms of number of mutations) and the clonal mutations (defined as belonging to the clone of highest CCF). Source data are provided as a Source Data file.

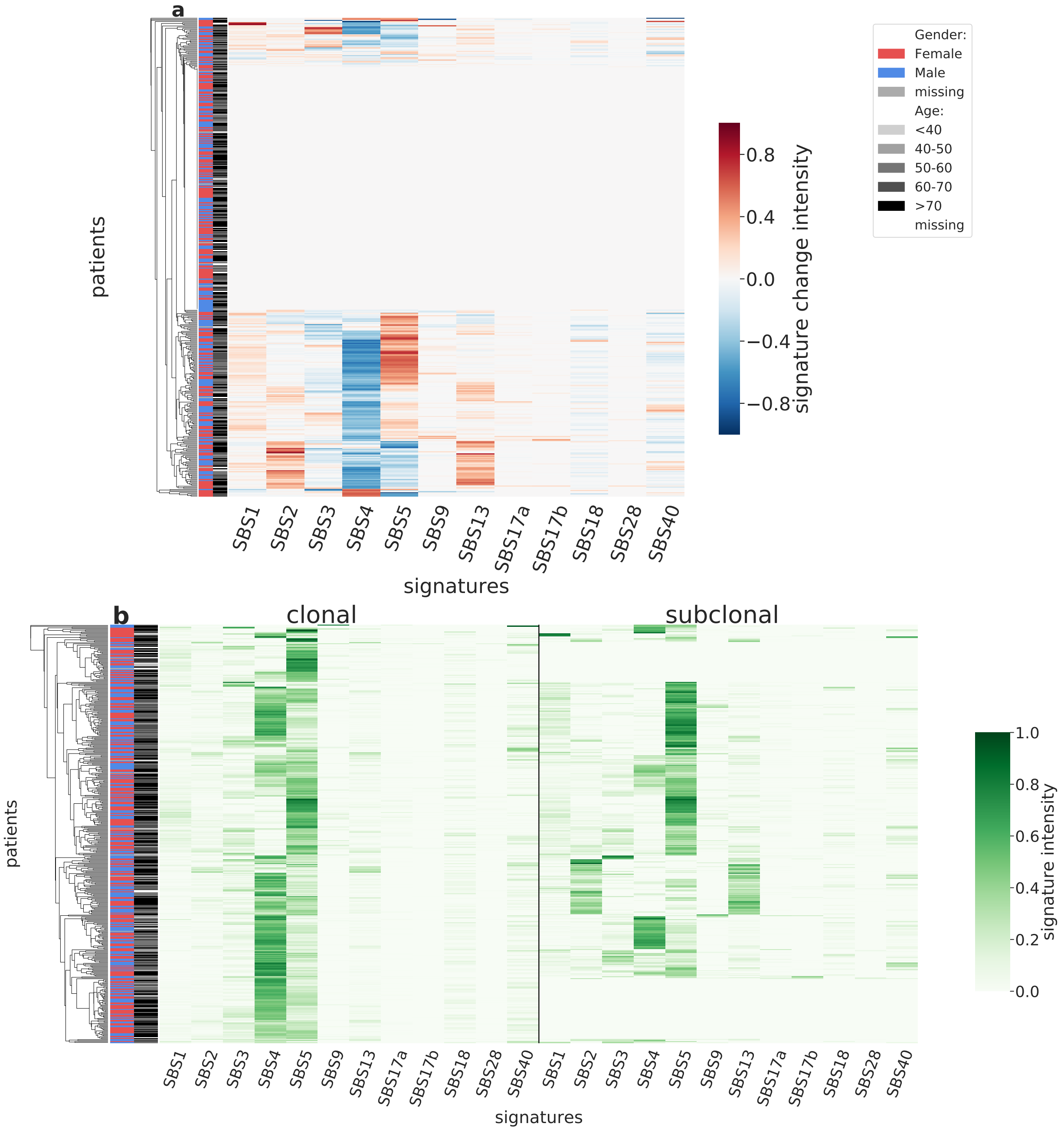

Supplementary Figure 53: **Overview of CloneSig results for the TCGA sub-cohort LUAD (n=433).** Panel a: Stratification of patients depending on their pattern of signature change for LUAD patients (433 patients, including 214 with a significant signature change). The heatmap represents the difference between the signature activity in the largest subclone (in terms of number of mutations) and the clonal mutations (defined as belonging to the clone of highest CCF).Panel b: Stratification of patients depending on their complete pattern of signature exposure. The heatmap represents the signature activity in the largest subclone (in terms of number of mutations) and the clonal mutations (defined as belonging to the clone of highest CCF). Source data are provided as a Source Data file.

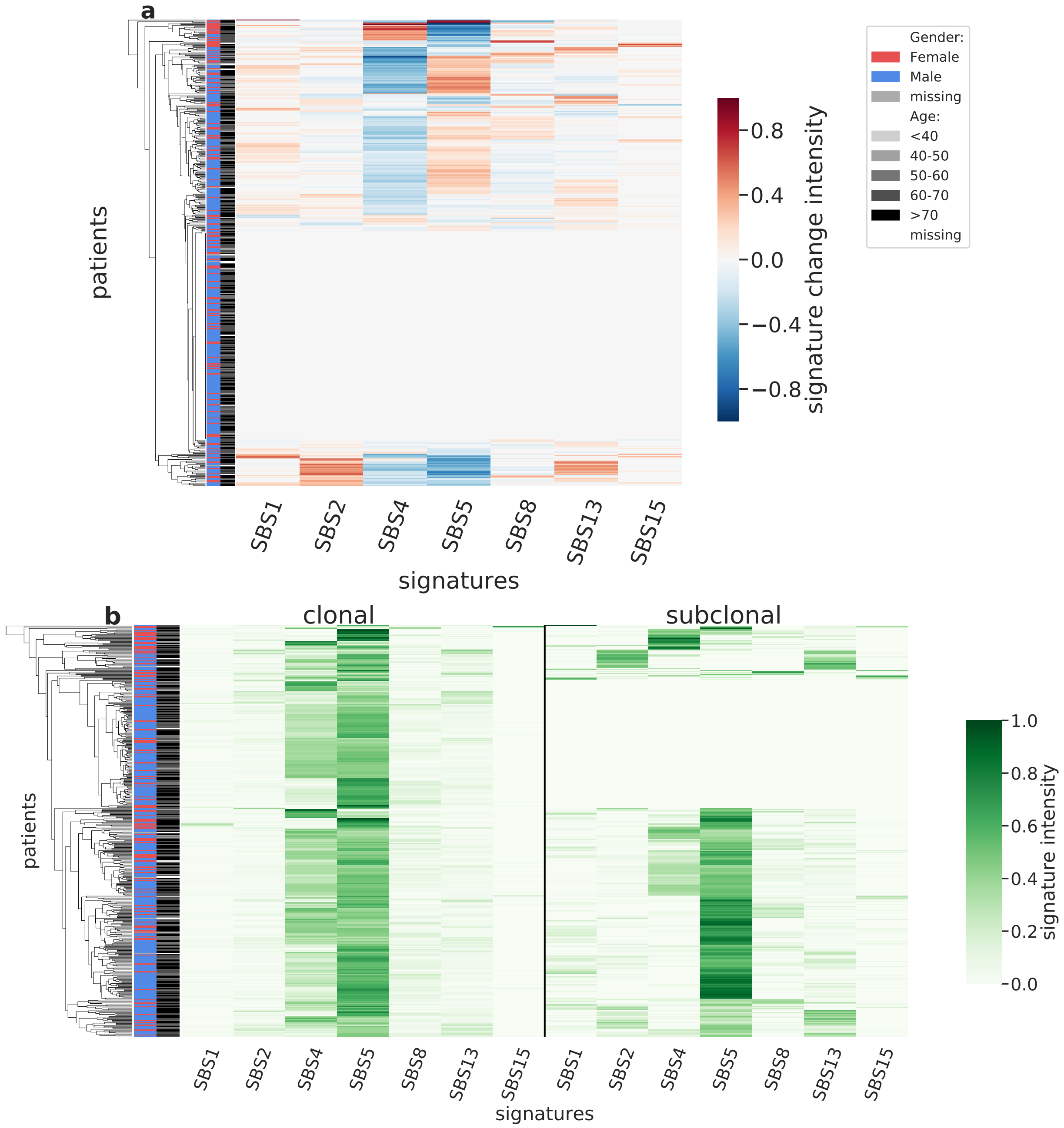

Supplementary Figure 54: **Overview of CloneSig results for the TCGA sub-cohort LUSC (n=423).** Panel a: Stratification of patients depending on their pattern of signature change for LUSC patients (423 patients, including 235 with a significant signature change). The heatmap represents the difference between the signature activity in the largest subclone (in terms of number of mutations) and the clonal mutations (defined as belonging to the clone of highest CCF).Panel b: Stratification of patients depending on their complete pattern of signature exposure. The heatmap represents the signature activity in the largest subclone (in terms of number of mutations) and the clonal mutations (defined as belonging to the clone of highest CCF). Source data are provided as a Source Data file.

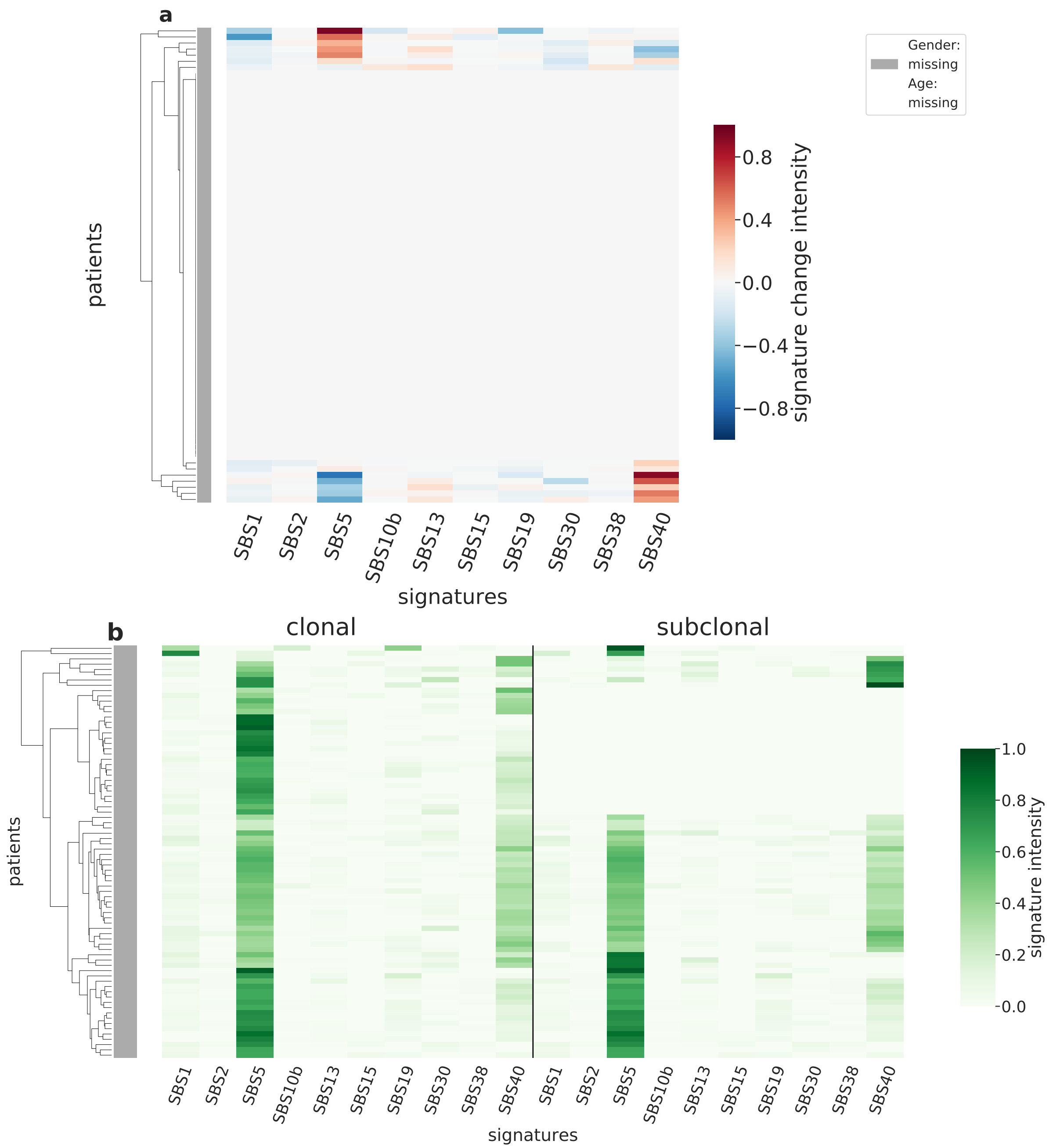

Supplementary Figure 55: **Overview of CloneSig results for the TCGA sub-cohort MESO (n=78).** Panel a: Stratification of patients depending on their pattern of signature change for MESO patients (78 patients, including 14 with a significant signature change). The heatmap represents the difference between the signature activity in the largest subclone (in terms of number of mutations) and the clonal mutations (defined as belonging to the clone of highest CCF).Panel b: Stratification of patients depending on their complete pattern of signature exposure. The heatmap represents the signature activity in the largest subclone (in terms of number of mutations) and the clonal mutations (defined as belonging to the clone of highest CCF). Source data are provided as a Source Data file.

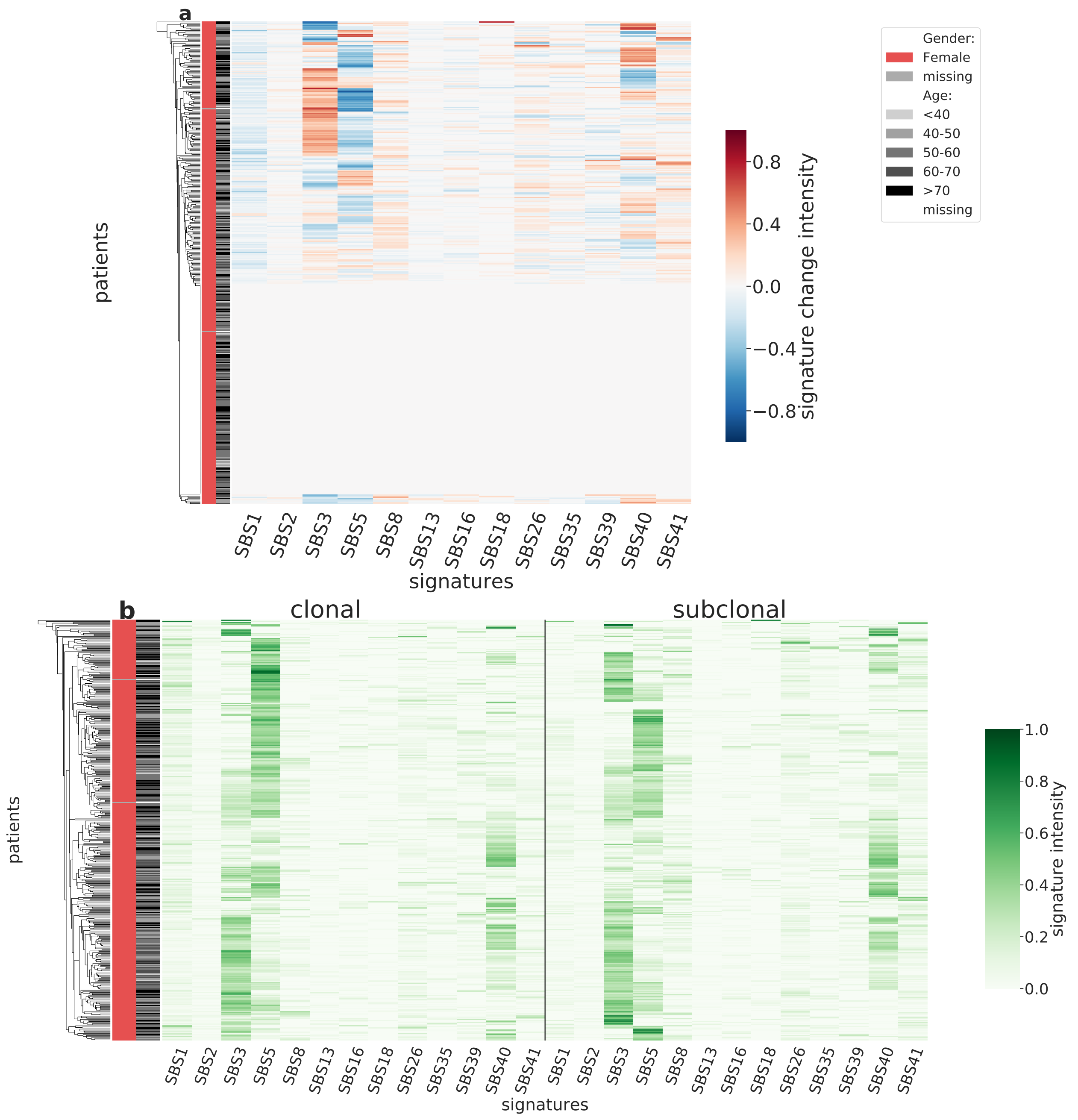

Supplementary Figure 56: **Overview of CloneSig results for the TCGA sub-cohort OV (n=390).** Panel a: Stratification of patients depending on their pattern of signature change for OV patients (390 patients, including 220 with a significant signature change). The heatmap represents the difference between the signature activity in the largest subclone (in terms of number of mutations) and the clonal mutations (defined as belonging to the clone of highest CCF).Panel b: Stratification of patients depending on their complete pattern of signature exposure. The heatmap represents the signature activity in the largest subclone (in terms of number of mutations) and the clonal mutations (defined as belonging to the clone of highest CCF). Source data are provided as a Source Data file.

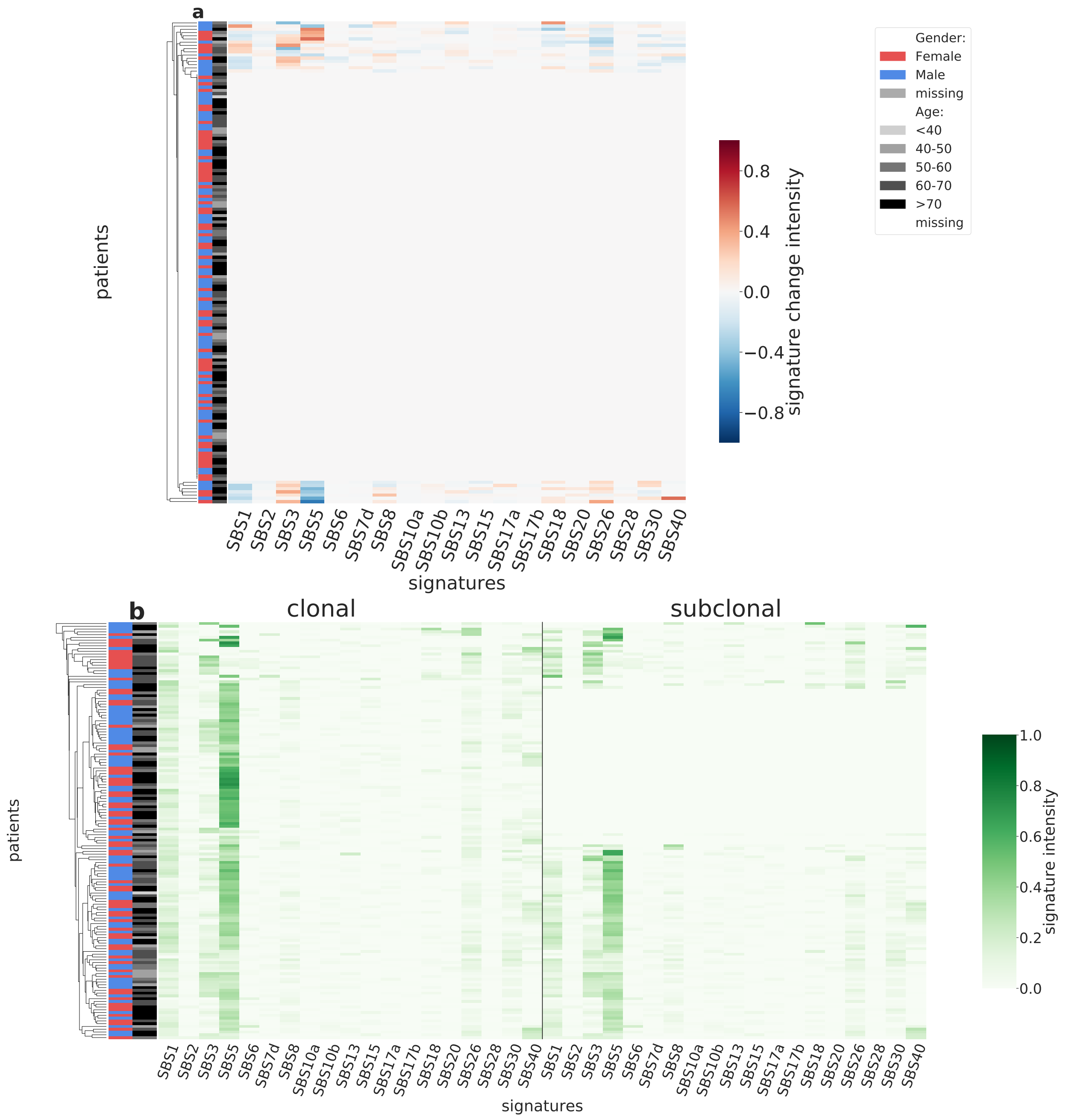

Supplementary Figure 57: **Overview of CloneSig results for the TCGA sub-cohort PAAD (n=150).** Panel a: Stratification of patients depending on their pattern of signature change for PAAD patients (150 patients, including 23 with a significant signature change). The heatmap represents the difference between the signature activity in the largest subclone (in terms of number of mutations) and the clonal mutations (defined as belonging to the clone of highest CCF).Panel b: Stratification of patients depending on their complete pattern of signature exposure. The heatmap represents the signature activity in the largest subclone (in terms of number of mutations) and the clonal mutations (defined as belonging to the clone of highest CCF). Source data are provided as a Source Data file.

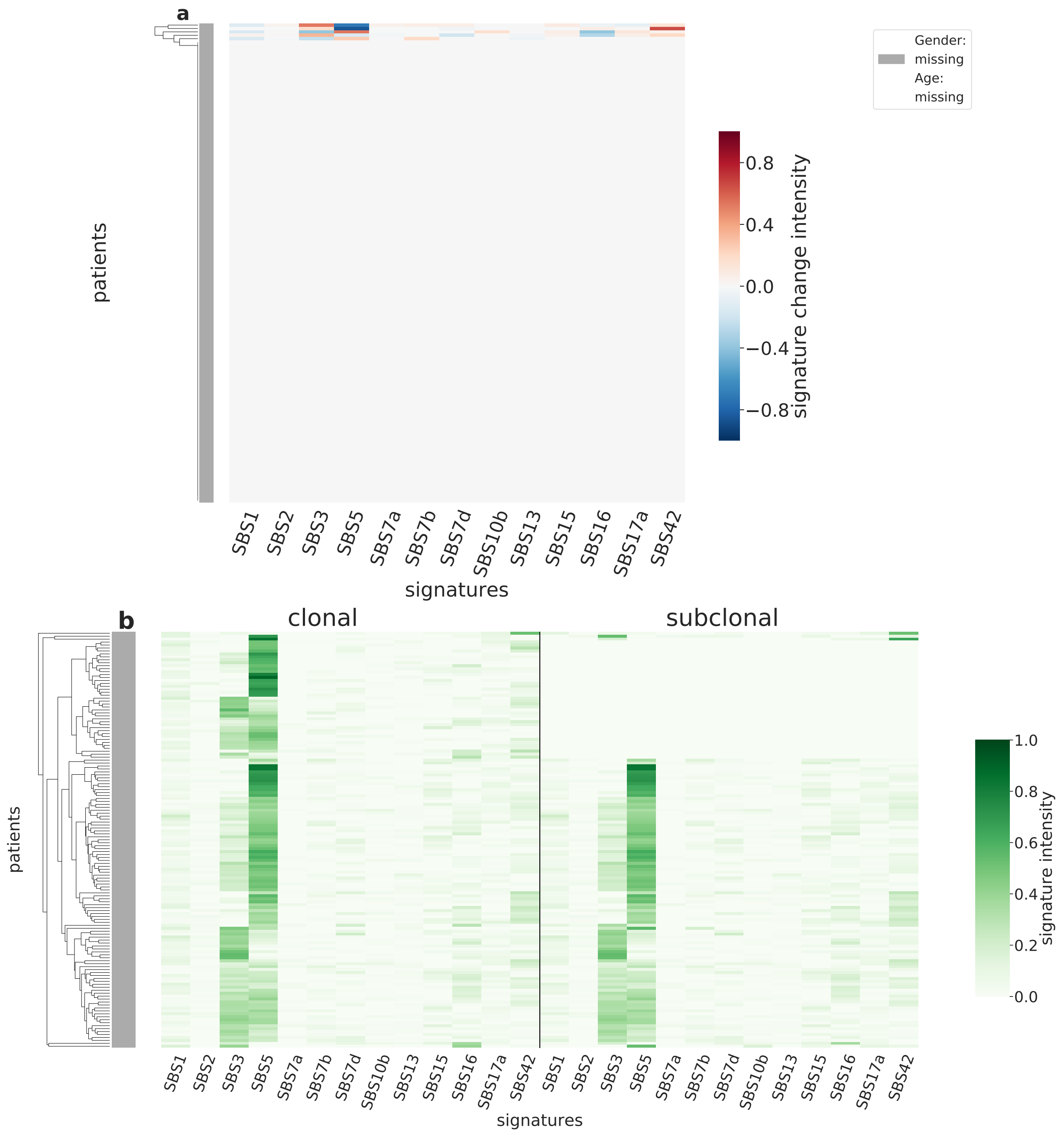

Supplementary Figure 58: **Overview of CloneSig results for the TCGA sub-cohort PCPG (n=141).** Panel a: Stratification of patients depending on their pattern of signature change for PCPG patients (141 patients, including 5 with a significant signature change). The heatmap represents the difference between the signature activity in the largest subclone (in terms of number of mutations) and the clonal mutations (defined as belonging to the clone of highest CCF).Panel b: Stratification of patients depending on their complete pattern of signature exposure. The heatmap represents the signature activity in the largest subclone (in terms of number of mutations) and the clonal mutations (defined as belonging to the clone of highest CCF). Source data are provided as a Source Data file.

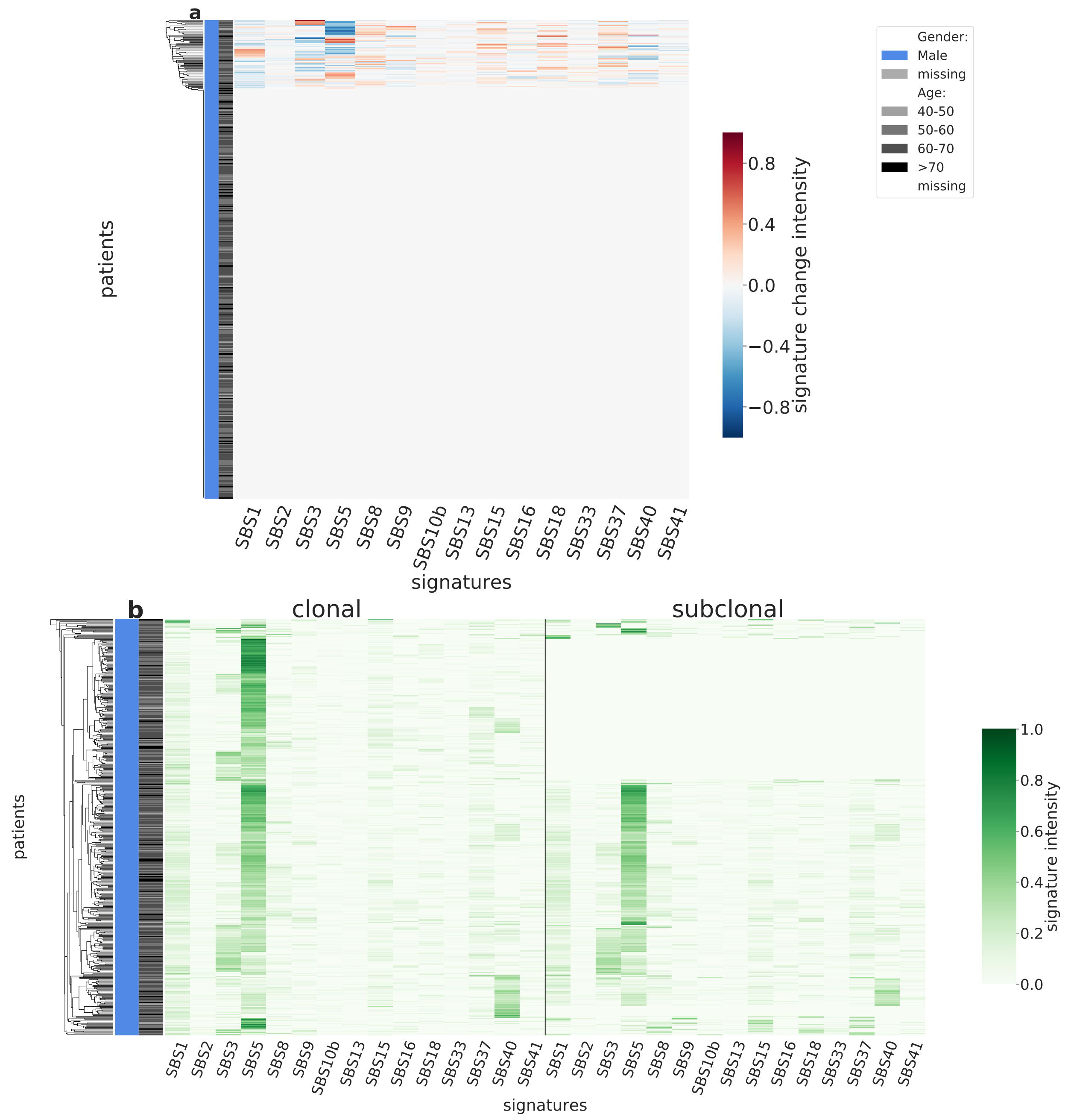

Supplementary Figure 59: **Overview of CloneSig results for the TCGA sub-cohort PRAD (n=458).** Panel a: Stratification of patients depending on their pattern of signature change for PRAD patients (458 patients, including 66 with a significant signature change). The heatmap represents the difference between the signature activity in the largest subclone (in terms of number of mutations) and the clonal mutations (defined as belonging to the clone of highest CCF).Panel b: Stratification of patients depending on their complete pattern of signature exposure. The heatmap represents the signature activity in the largest subclone (in terms of number of mutations) and the clonal mutations (defined as belonging to the clone of highest CCF). Source data are provided as a Source Data file.

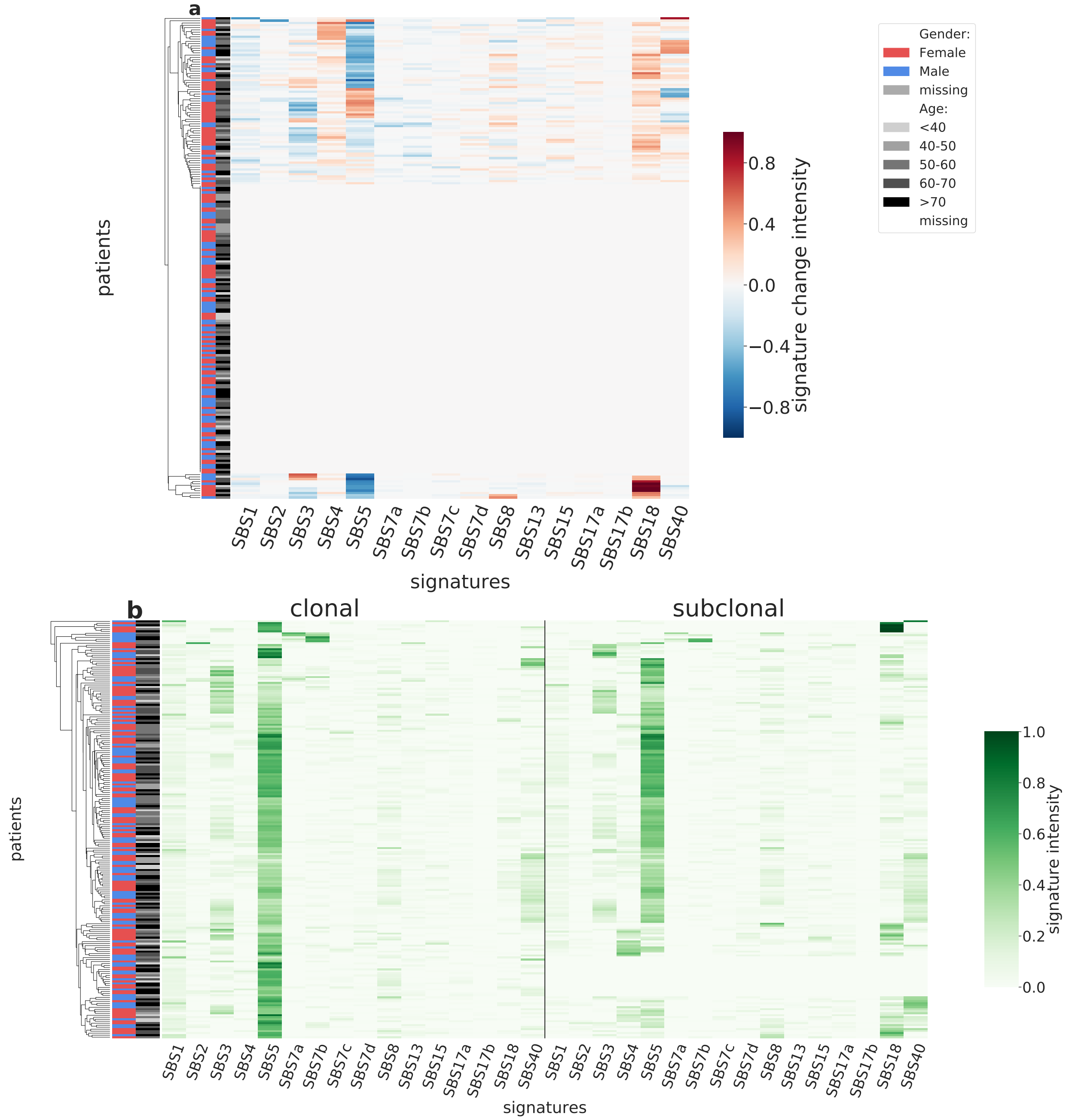

Supplementary Figure 60: **Overview of CloneSig results for the TCGA sub-cohort SARC (n=210).** Panel a: Stratification of patients depending on their pattern of signature change for SARC patients (210 patients, including 84 with a significant signature change). The heatmap represents the difference between the signature activity in the largest subclone (in terms of number of mutations) and the clonal mutations (defined as belonging to the clone of highest CCF).Panel b: Stratification of patients depending on their complete pattern of signature exposure. The heatmap represents the signature activity in the largest subclone (in terms of number of mutations) and the clonal mutations (defined as belonging to the clone of highest CCF). Source data are provided as a Source Data file.

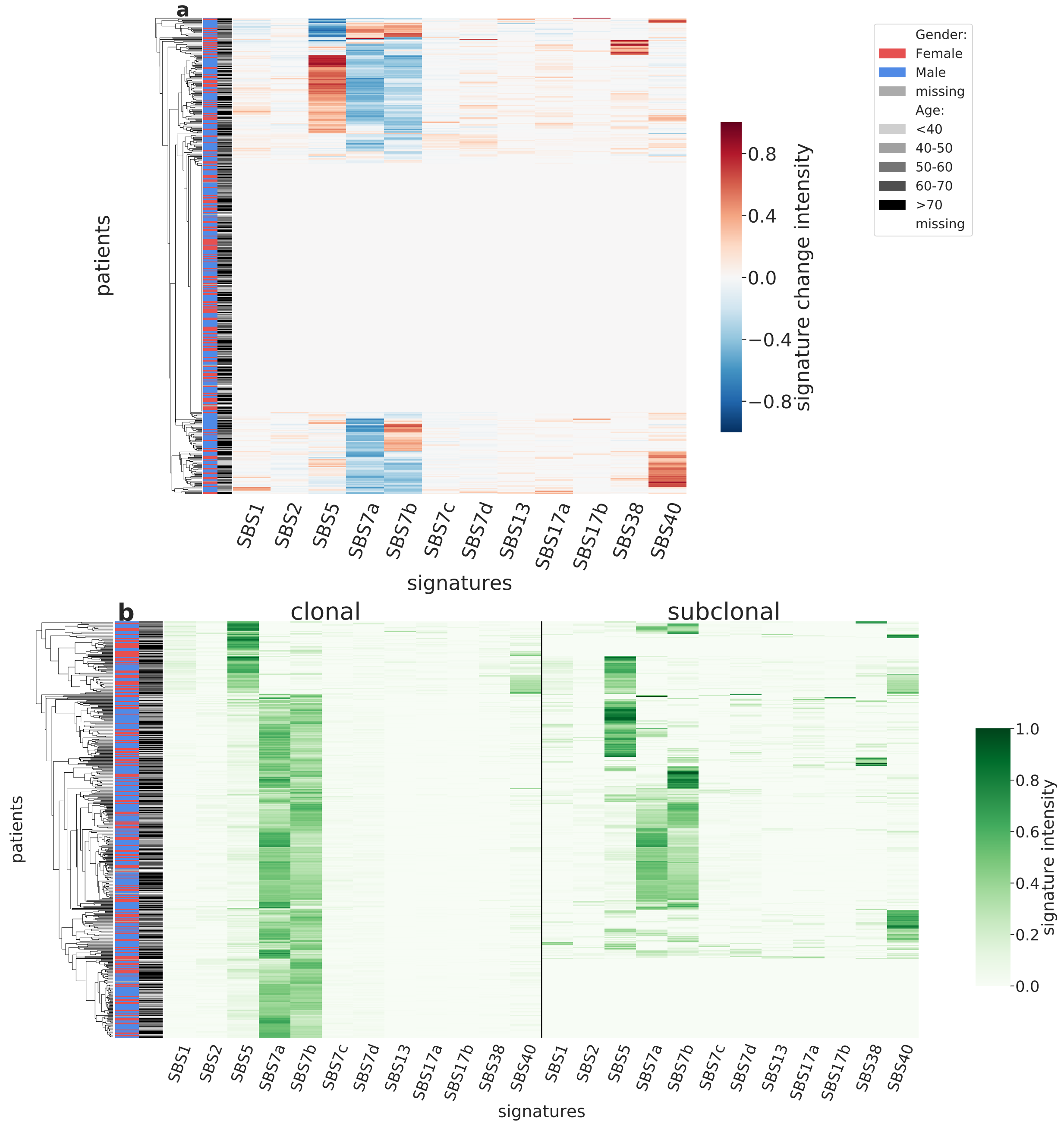

Supplementary Figure 61: **Overview of CloneSig results for the TCGA sub-cohort SKCM (n=424).** Panel a: Stratification of patients depending on their pattern of signature change for SKCM patients (424 patients, including 203 with a significant signature change). The heatmap represents the difference between the signature activity in the largest subclone (in terms of number of mutations) and the clonal mutations (defined as belonging to the clone of highest CCF).Panel b: Stratification of patients depending on their complete pattern of signature exposure. The heatmap represents the signature activity in the largest subclone (in terms of number of mutations) and the clonal mutations (defined as belonging to the clone of highest CCF). Source data are provided as a Source Data file.

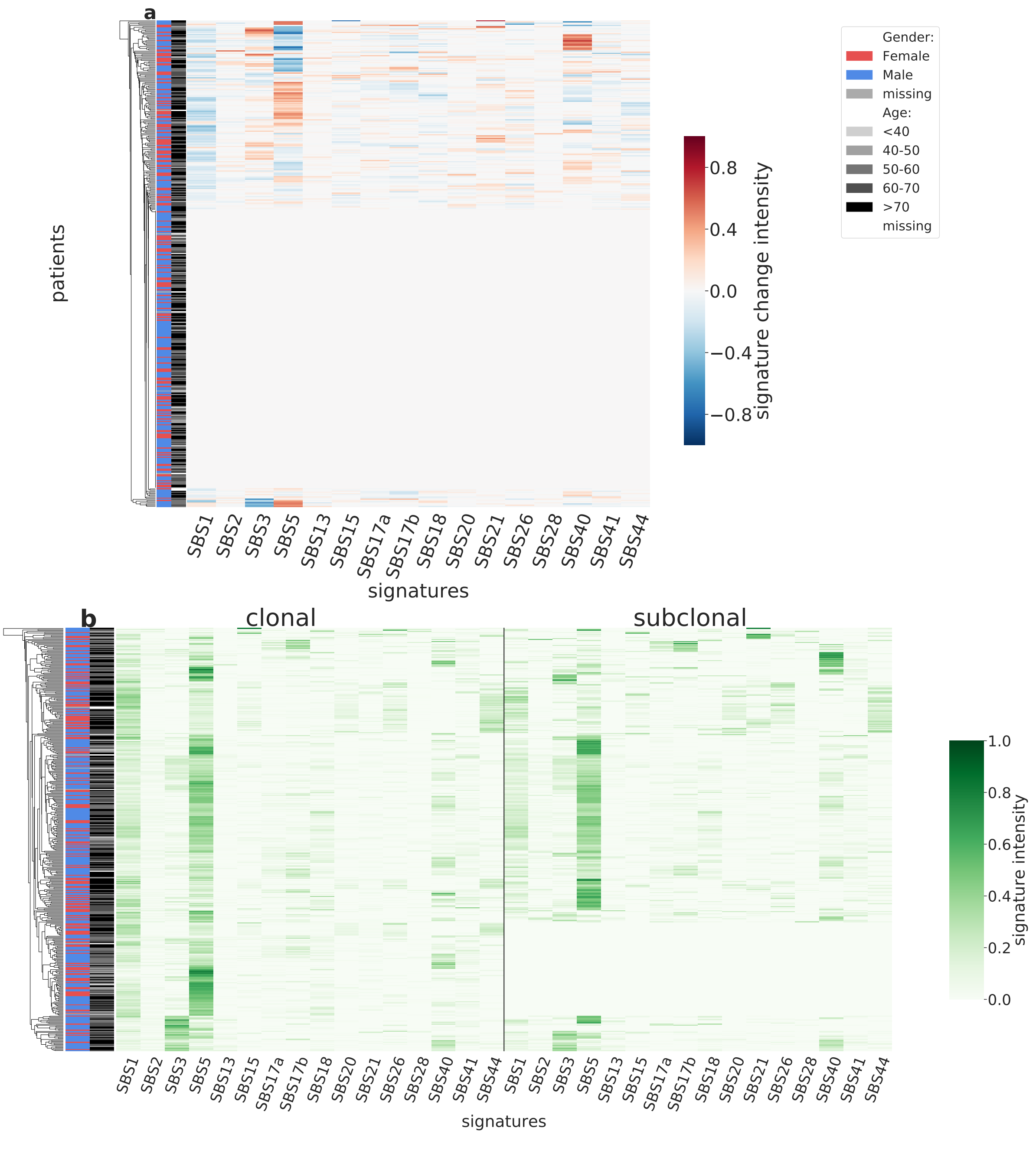

Supplementary Figure 62: **Overview of CloneSig results for the TCGA sub-cohort STAD (n=418).** Panel a: Stratification of patients depending on their pattern of signature change for STAD patients (418 patients, including 179 with a significant signature change). The heatmap represents the difference between the signature activity in the largest subclone (in terms of number of mutations) and the clonal mutations (defined as belonging to the clone of highest CCF).Panel b: Stratification of patients depending on their complete pattern of signature exposure. The heatmap represents the signature activity in the largest subclone (in terms of number of mutations) and the clonal mutations (defined as belonging to the clone of highest CCF). Source data are provided as a Source Data file.

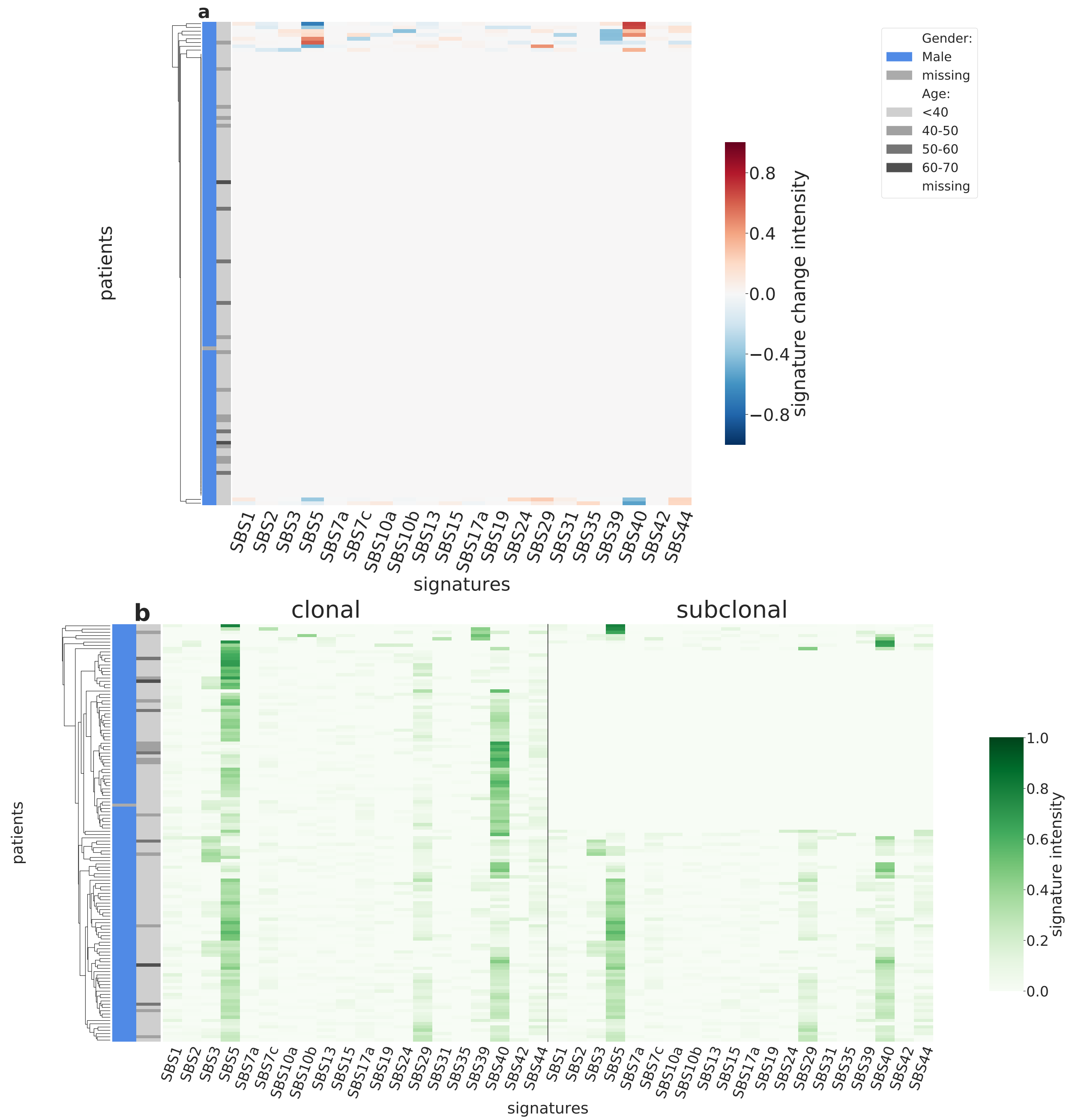

Supplementary Figure 63: **Overview of CloneSig results for the TCGA sub-cohort TGCT (n=128).** Panel a: Stratification of patients depending on their pattern of signature change for TGCT patients (128 patients, including 10 with a significant signature change). The heatmap represents the difference between the signature activity in the largest subclone (in terms of number of mutations) and the clonal mutations (defined as belonging to the clone of highest CCF).Panel b: Stratification of patients depending on their complete pattern of signature exposure. The heatmap represents the signature activity in the largest subclone (in terms of number of mutations) and the clonal mutations (defined as belonging to the clone of highest CCF). Source data are provided as a Source Data file.

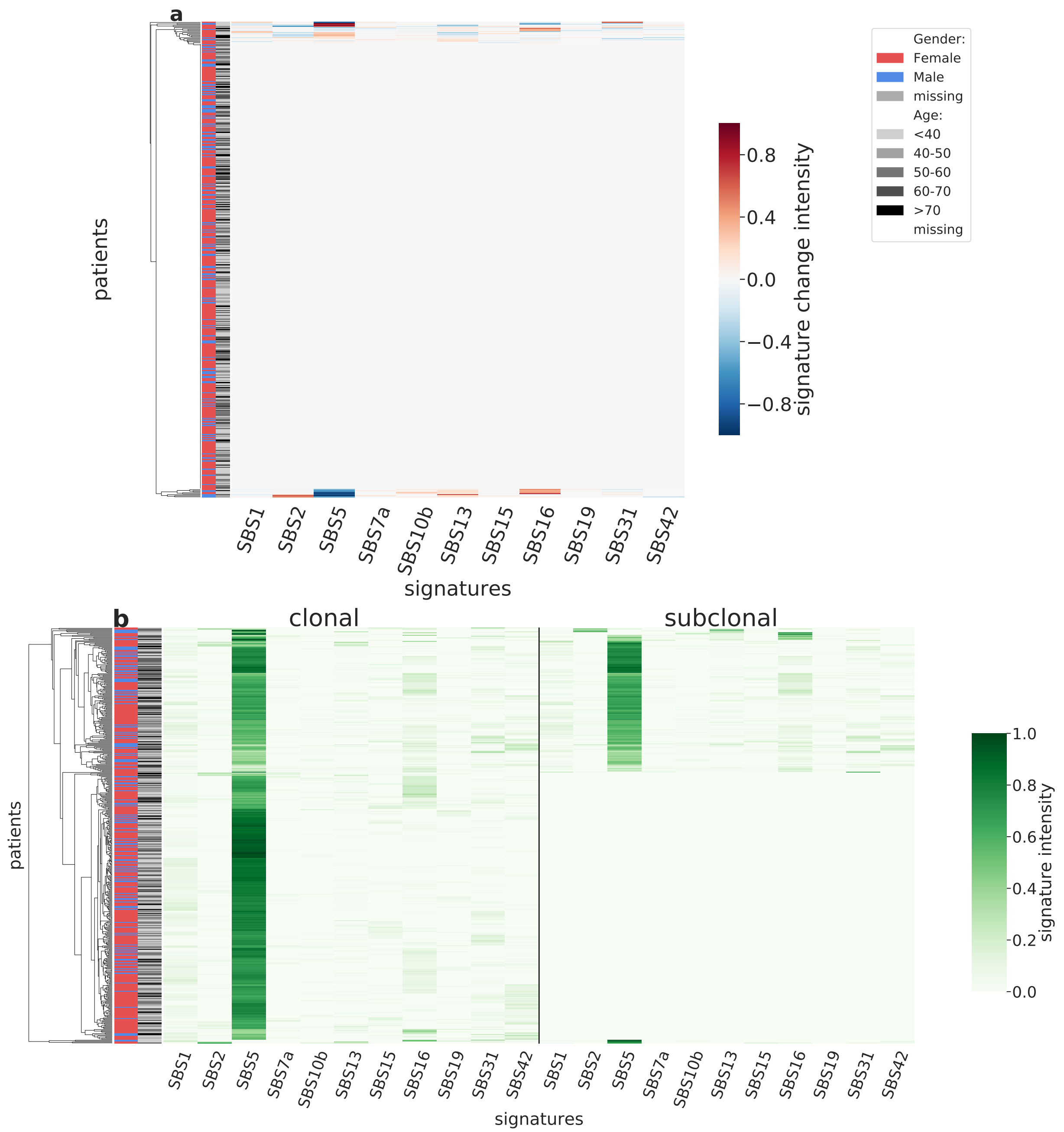

Supplementary Figure 64: **Overview of CloneSig results for the TCGA sub-cohort THCA (n=467).** Panel a: Stratification of patients depending on their pattern of signature change for THCA patients (467 patients, including 29 with a significant signature change). The heatmap represents the difference between the signature activity in the largest subclone (in terms of number of mutations) and the clonal mutations (defined as belonging to the clone of highest CCF).Panel b: Stratification of patients depending on their complete pattern of signature exposure. The heatmap represents the signature activity in the largest subclone (in terms of number of mutations) and the clonal mutations (defined as belonging to the clone of highest CCF). Source data are provided as a Source Data file.

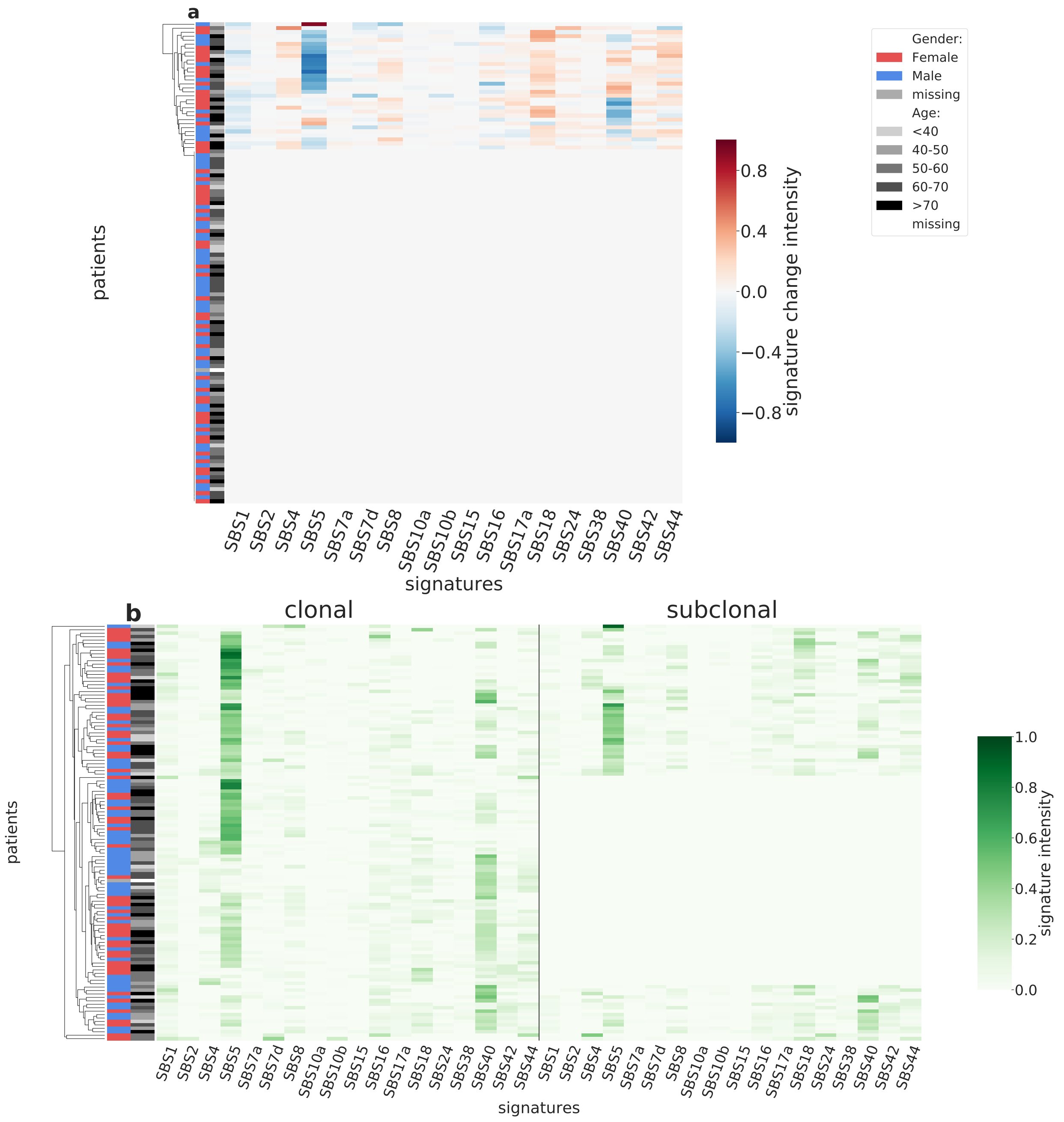

Supplementary Figure 65: **Overview of CloneSig results for the TCGA sub-cohort THYM (n=121).** Panel a: Stratification of patients depending on their pattern of signature change for THYM patients (121 patients, including 32 with a significant signature change). The heatmap represents the difference between the signature activity in the largest subclone (in terms of number of mutations) and the clonal mutations (defined as belonging to the clone of highest CCF).Panel b: Stratification of patients depending on their complete pattern of signature exposure. The heatmap represents the signature activity in the largest subclone (in terms of number of mutations) and the clonal mutations (defined as belonging to the clone of highest CCF). Source data are provided as a Source Data file.

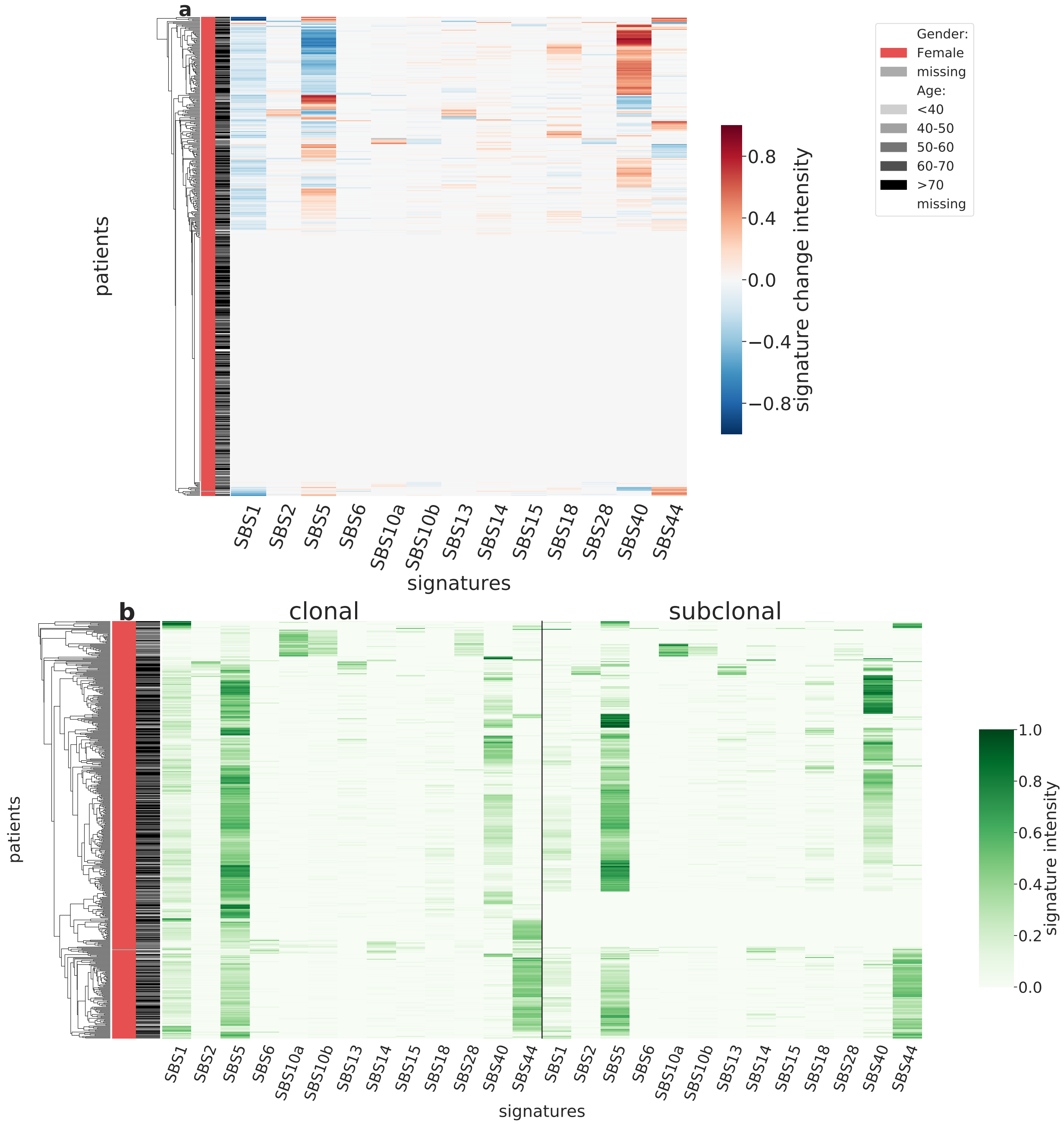

Supplementary Figure 66: **Overview of CloneSig results for the TCGA sub-cohort UCEC (n=486).** Panel a: Stratification of patients depending on their pattern of signature change for UCEC patients (486 patients, including 236 with a significant signature change). The heatmap represents the difference between the signature activity in the largest subclone (in terms of number of mutations) and the clonal mutations (defined as belonging to the clone of highest CCF).Panel b: Stratification of patients depending on their complete pattern of signature exposure. The heatmap represents the signature activity in the largest subclone (in terms of number of mutations) and the clonal mutations (defined as belonging to the clone of highest CCF). Source data are provided as a Source Data file.

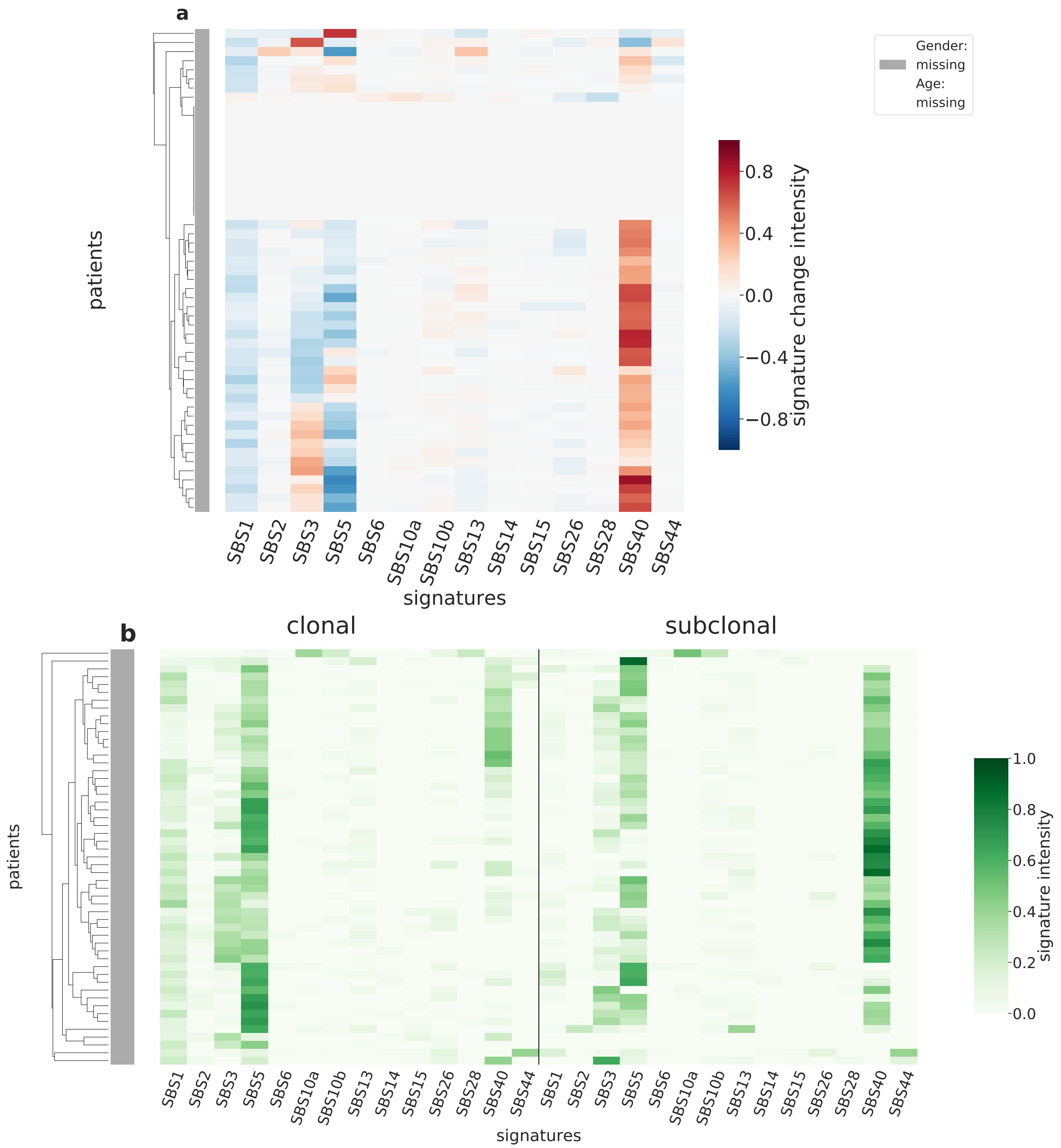

Supplementary Figure 67: **Overview of CloneSig results for the TCGA sub-cohort UCS (n=53).** Panel a: Stratification of patients depending on their pattern of signature change for UCS patients (53 patients, including 40 with a significant signature change). The heatmap represents the difference between the signature activity in the largest subclone (in terms of number of mutations) and the clonal mutations (defined as belonging to the clone of highest CCF).Panel b: Stratification of patients depending on their complete pattern of signature exposure. The heatmap represents the signature activity in the largest subclone (in terms of number of mutations) and the clonal mutations (defined as belonging to the clone of highest CCF). Source data are provided as a Source Data file.

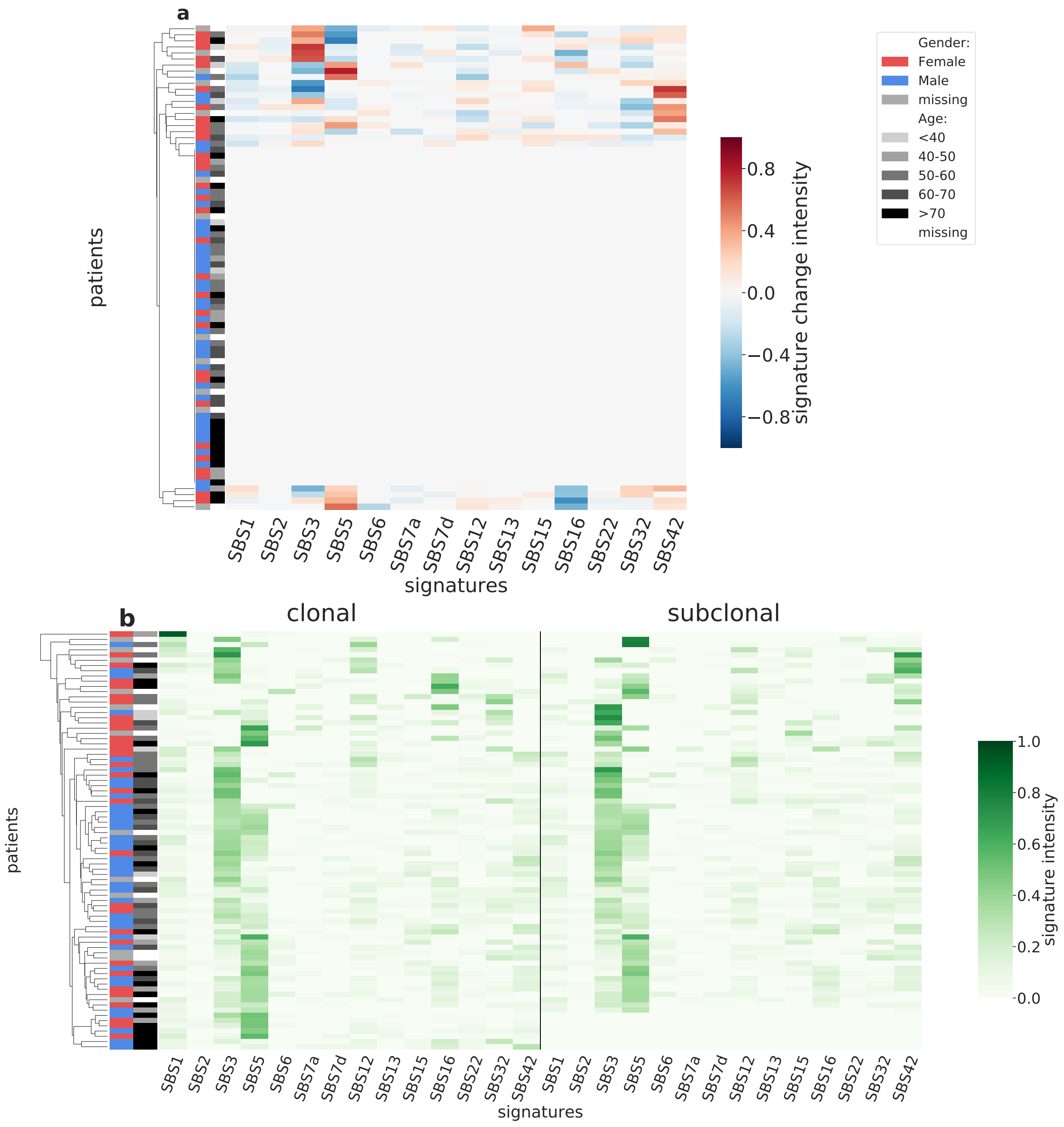

Supplementary Figure 68: **Overview of CloneSig results for the TCGA sub-cohort UVM (n=80).** Panel a: Stratification of patients depending on their pattern of signature change for UVM patients (80 patients, including 24 with a significant signature change). The heatmap represents the difference between the signature activity in the largest subclone (in terms of number of mutations) and the clonal mutations (defined as belonging to the clone of highest CCF).Panel b: Stratification of patients depending on their complete pattern of signature exposure. The heatmap represents the signature activity in the largest subclone (in terms of number of mutations) and the clonal mutations (defined as belonging to the clone of highest CCF). Source data are provided as a Source Data file.

### Supplementary Note 4: Complete overview of PCAWG results

We applied CloneSig to the pan-cancer PCAWG cohort of 2,632 patients, spanning 32 cancer types.

For each sample in the cohort, we estimate with CloneSig the number of subclones present in the tumor, the signatures active in each subclone, and test for the presence of a significant signature change between clones. Based on samples exhibiting a significant signature change, we have attempted to identify the signatures that are the most variant for each cancer type. To that end, we have computed the absolute difference in signature activity between the largest subclone and the set of clonal mutations. We neglected cases where the absolute difference was below 0.05. Figure 69 shows a global summary of the signature changes found in the cohort. For each cancer type, it shows the proportion of samples where a signature change is found, and a visual summary of the proportion of samples where each individual signature is found to increase or to decrease in the largest subclone, compared to the clonal mutations. The thickness of each bar, in addition, indicates the median change of each signature. We retained only signatures found variant in more than 10% of each cohort samples.

We additionally generated similar plots using a log-fold change, a relative variation of signature activity measure.

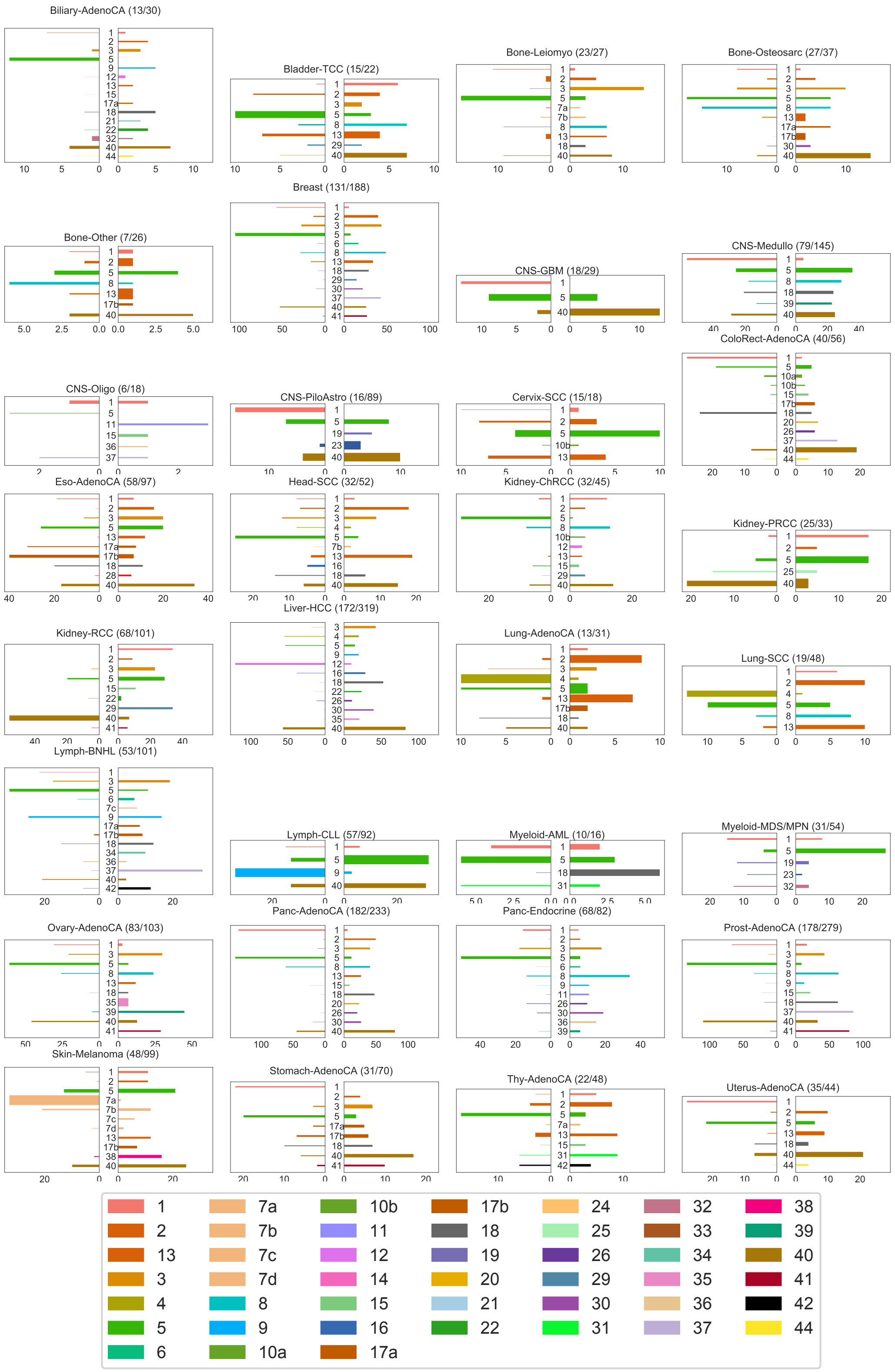

Supplementary Figure 69: **Mutational signature changes in the PCAWG cohort.** Each plot corresponds to one cancer type, indicates the number of samples with a significant signature change compared to the total number of samples, and shows on the right panel an increase of a signature in the largest subclone, compared to clonal mutations, and on the left panel a decrease. The length of each bar corresponds to the number of patients with such changes, and the thickness to the median absolute observed change. Source data are provided as a Source Data file.

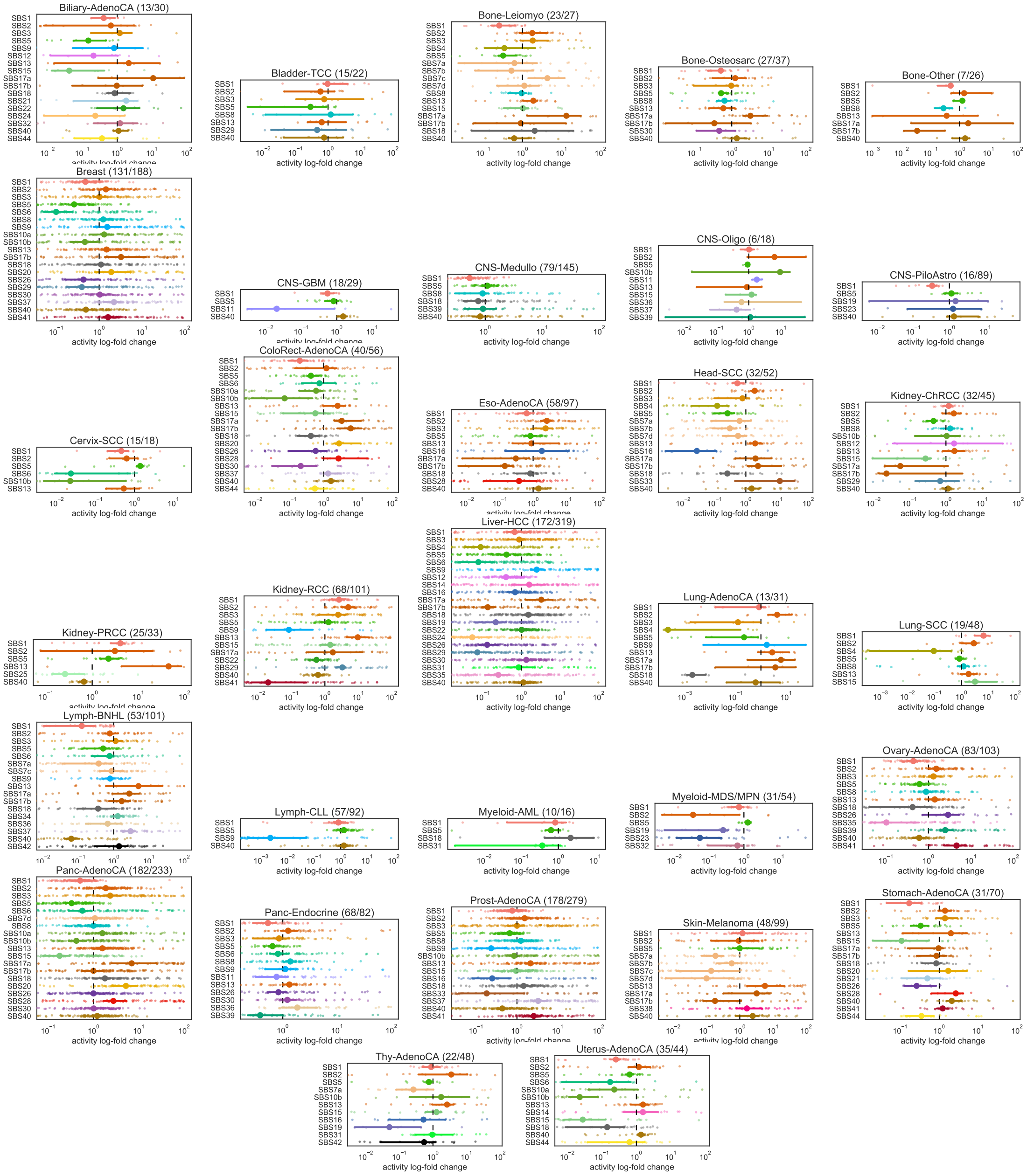

Supplementary Figure 70: **Mutational signature changes in the PCAWG cohort.** Each plot corresponds to one cancer type, indicates the number of samples with a significant signature change compared to the total number of samples, and shows log fold change of signature activity in the largest subclone, compared to clonal mutations. The small dots correspond to individual samples, and the larger dot to the median change. Bootstrap sampling of the scores was used to compute 95% confidence intervals. The vertical dotted line in each plot represent no change.

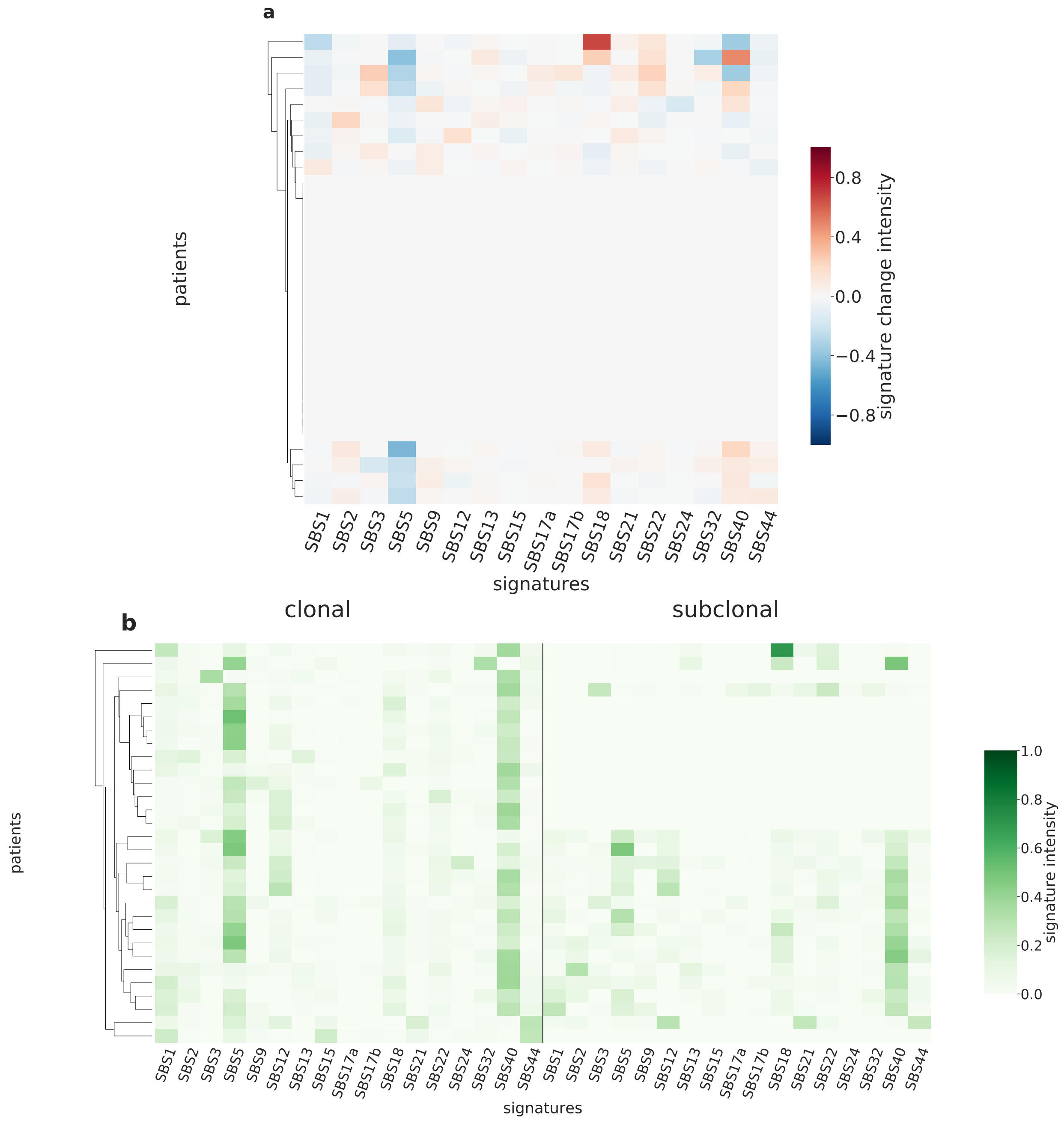

Supplementary Figure 71: **Overview of CloneSig results for the PCAWG sub-cohort Biliary-AdenoCA (n=30).** Panel a: Stratification of patients depending on their pattern of signature change for Biliary-AdenoCA patients (30 patients, including 13 with a significant signature change). The heatmap represents the difference between the signature activity in the largest subclone (in terms of number of mutations) and the clonal mutations (defined as belonging to the clone of highest CCF).Panel b: Stratification of patients depending on their complete pattern of signature exposure. The heatmap represents the signature activity in the largest subclone (in terms of number of mutations) and the clonal mutations (defined as belonging to the clone of highest CCF). Source data are provided as a Source Data file.

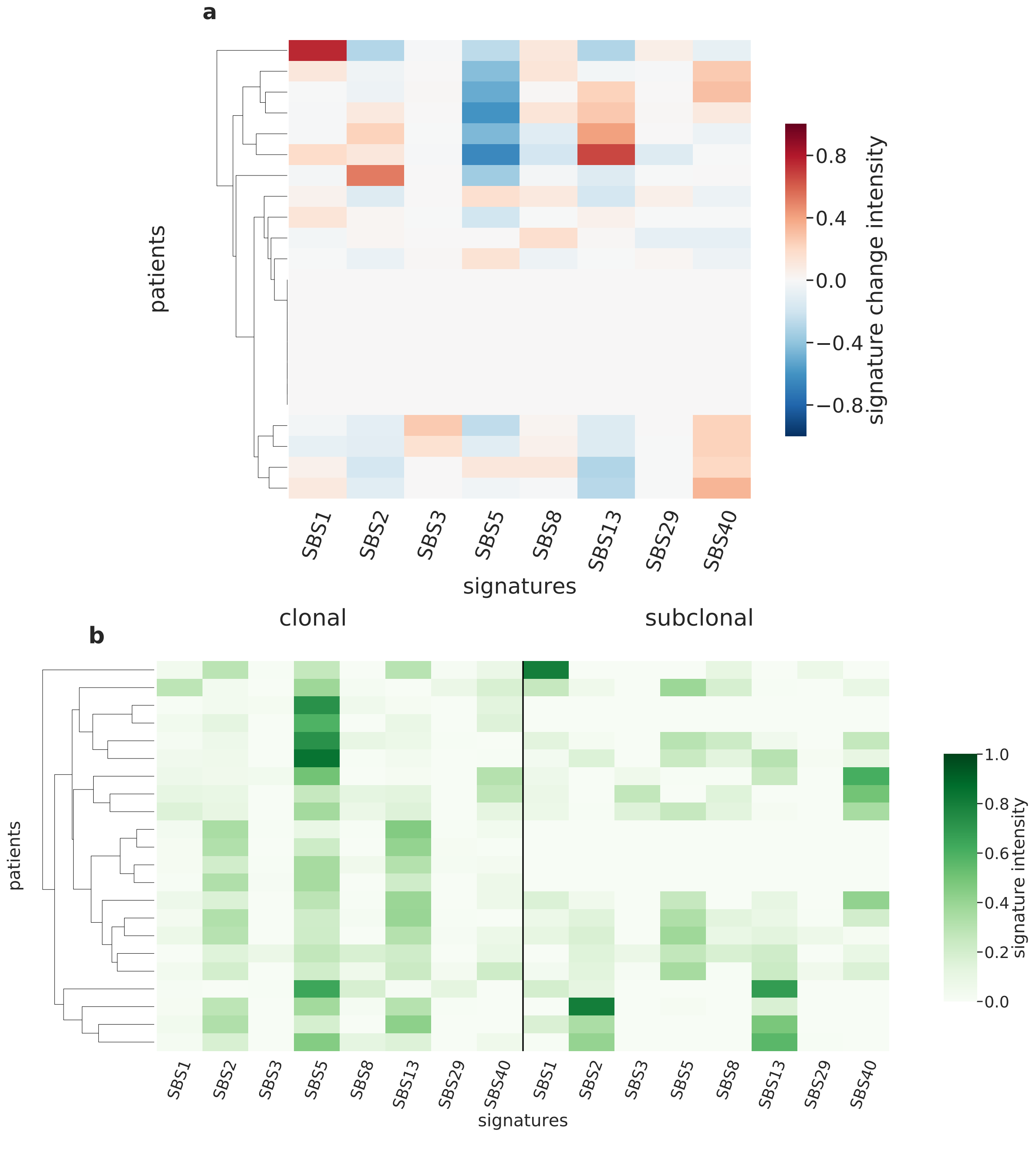

Supplementary Figure 72: **Overview of CloneSig results for the PCAWG sub-cohort Bladder-TCC (n=22).** Panel a: Stratification of patients depending on their pattern of signature change for Bladder-TCC patients (22 patients, including 15 with a significant signature change). The heatmap represents the difference between the signature activity in the largest subclone (in terms of number of mutations) and the clonal mutations (defined as belonging to the clone of highest CCF).Panel b: Stratification of patients depending on their complete pattern of signature exposure. The heatmap represents the signature activity in the largest subclone (in terms of number of mutations) and the clonal mutations (defined as belonging to the clone of highest CCF). Source data are provided as a Source Data file.

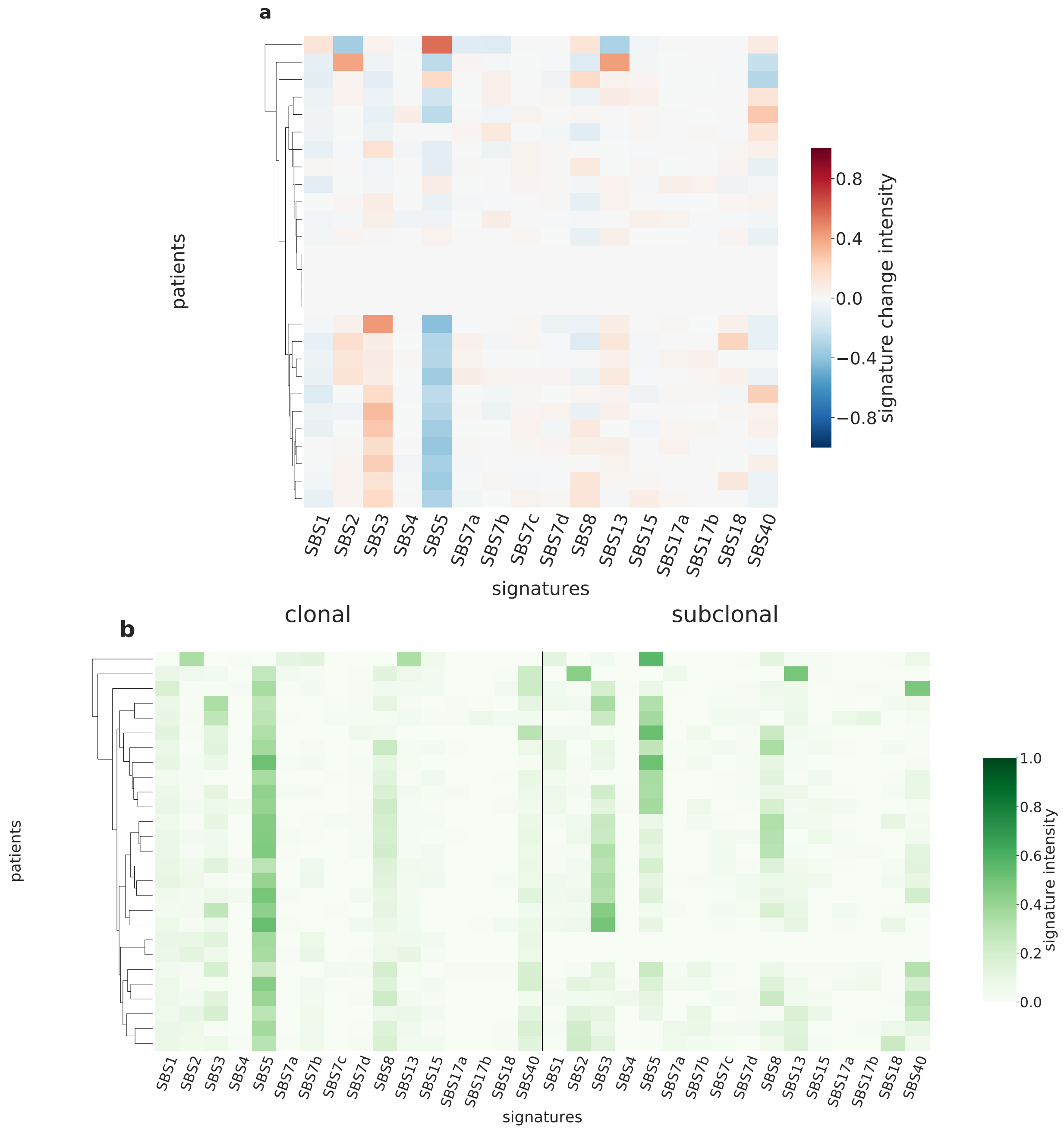

Supplementary Figure 73: **Overview of CloneSig results for the PCAWG sub-cohort Bone-Leiomyo (n=27).** Panel a: Stratification of patients depending on their pattern of signature change for Bone-Leiomyo patients (27 patients, including 23 with a significant signature change). The heatmap represents the difference between the signature activity in the largest subclone (in terms of number of mutations) and the clonal mutations (defined as belonging to the clone of highest CCF).Panel b: Stratification of patients depending on their complete pattern of signature exposure. The heatmap represents the signature activity in the largest subclone (in terms of number of mutations) and the clonal mutations (defined as belonging to the clone of highest CCF). Source data are provided as a Source Data file.

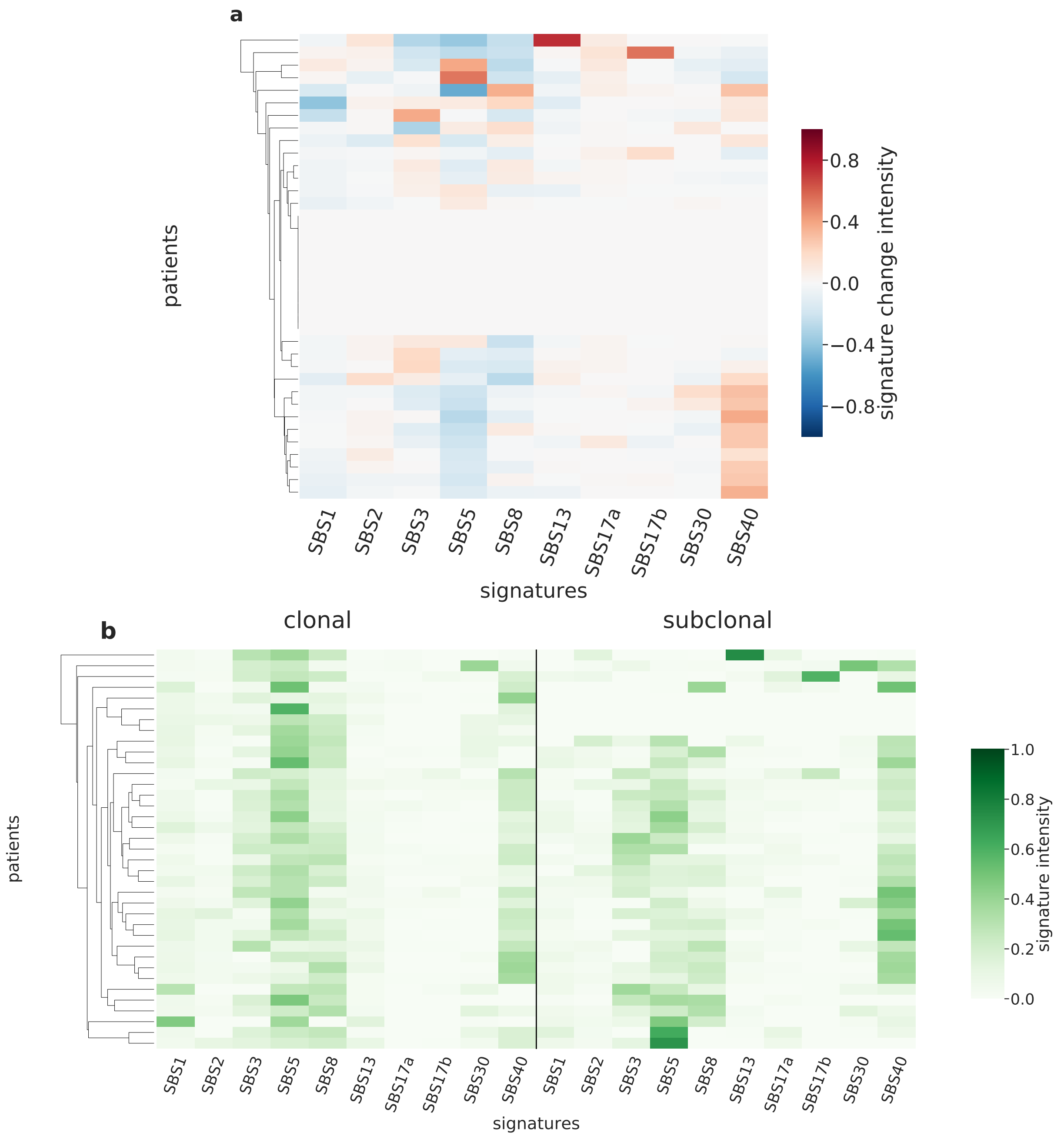

Supplementary Figure 74: **Overview of CloneSig results for the PCAWG sub-cohort Bone-Osteosarc (n=37).** Panel a: Stratification of patients depending on their pattern of signature change for Bone-Osteosarc patients (37 patients, including 27 with a significant signature change). The heatmap represents the difference between the signature activity in the largest subclone (in terms of number of mutations) and the clonal mutations (defined as belonging to the clone of highest CCF).Panel b: Stratification of patients depending on their complete pattern of signature exposure. The heatmap represents the signature activity in the largest subclone (in terms of number of mutations) and the clonal mutations (defined as belonging to the clone of highest CCF). Source data are provided as a Source Data file.

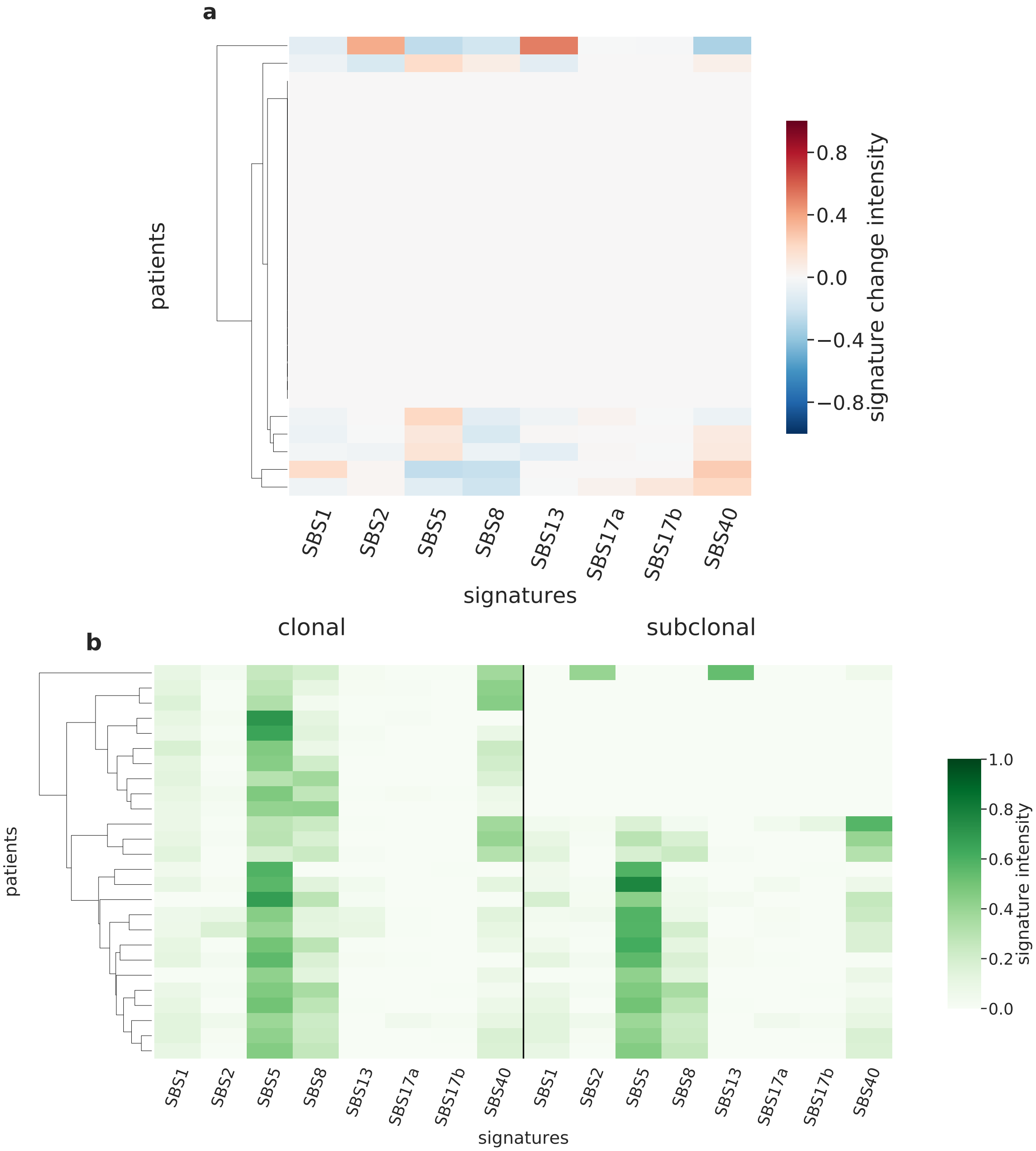

Supplementary Figure 75: **Overview of CloneSig results for the PCAWG sub-cohort Bone-Other (n=26).** Panel a: Stratification of patients depending on their pattern of signature change for Bone-Other patients (26 patients, including 7 with a significant signature change). The heatmap represents the difference between the signature activity in the largest subclone (in terms of number of mutations) and the clonal mutations (defined as belonging to the clone of highest CCF).Panel b: Stratification of patients depending on their complete pattern of signature exposure. The heatmap represents the signature activity in the largest subclone (in terms of number of mutations) and the clonal mutations (defined as belonging to the clone of highest CCF). Source data are provided as a Source Data file.

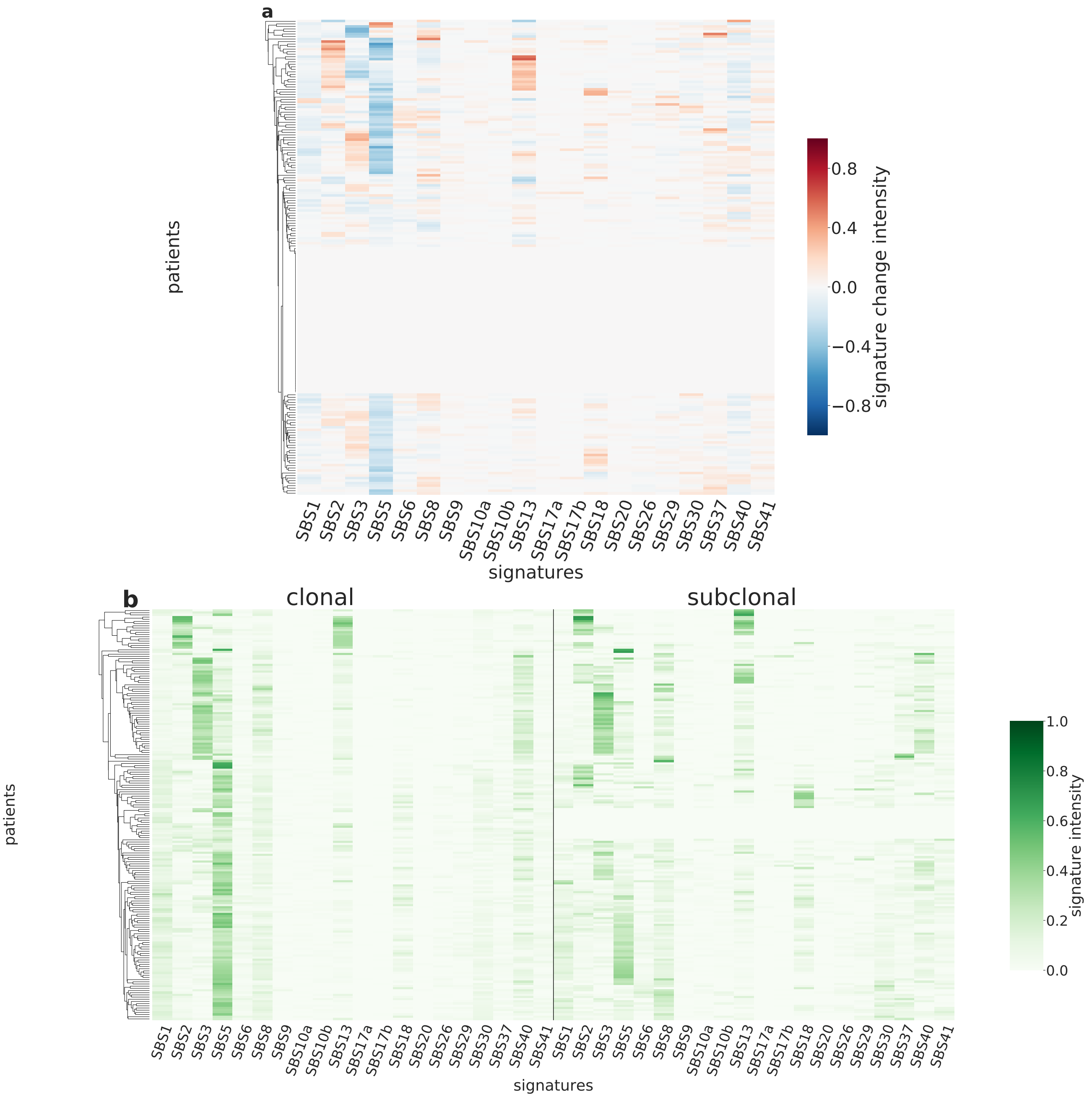

Supplementary Figure 76: **Overview of CloneSig results for the PCAWG sub-cohort Breast (n=188).** Panel a: Stratification of patients depending on their pattern of signature change for Breast patients (188 patients, including 131 with a significant signature change). The heatmap represents the difference between the signature activity in the largest subclone (in terms of number of mutations) and the clonal mutations (defined as belonging to the clone of highest CCF).Panel b: Stratification of patients depending on their complete pattern of signature exposure. The heatmap represents the signature activity in the largest subclone (in terms of number of mutations) and the clonal mutations (defined as belonging to the clone of highest CCF). Source data are provided as a Source Data file.

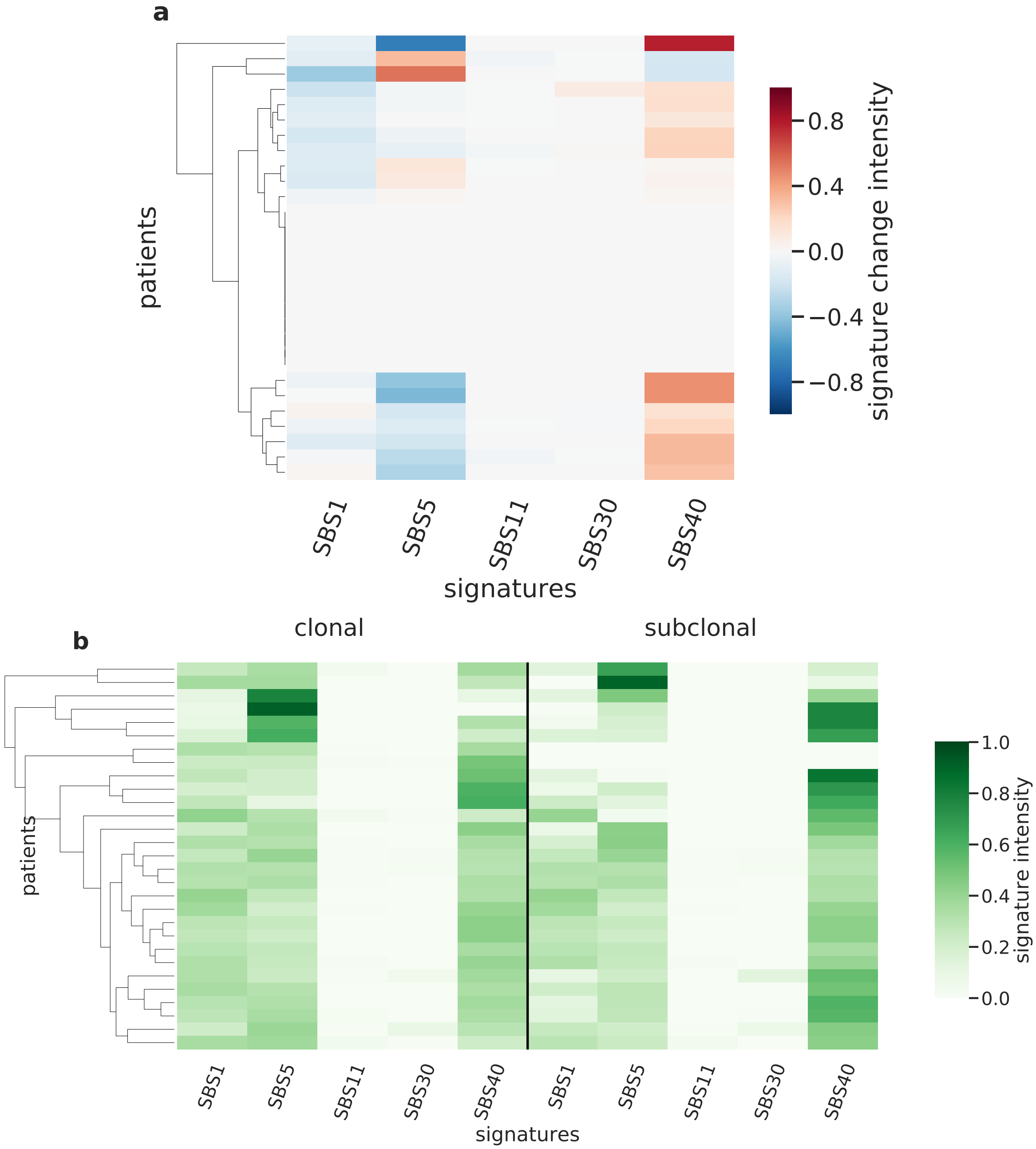

Supplementary Figure 77: **Overview of CloneSig results for the PCAWG sub-cohort CNS-GBM (n=29).** Panel a: Stratification of patients depending on their pattern of signature change for CNS-GBM patients (29 patients, including 18 with a significant signature change). The heatmap represents the difference between the signature activity in the largest subclone (in terms of number of mutations) and the clonal mutations (defined as belonging to the clone of highest CCF).Panel b: Stratification of patients depending on their complete pattern of signature exposure. The heatmap represents the signature activity in the largest subclone (in terms of number of mutations) and the clonal mutations (defined as belonging to the clone of highest CCF). Source data are provided as a Source Data file.

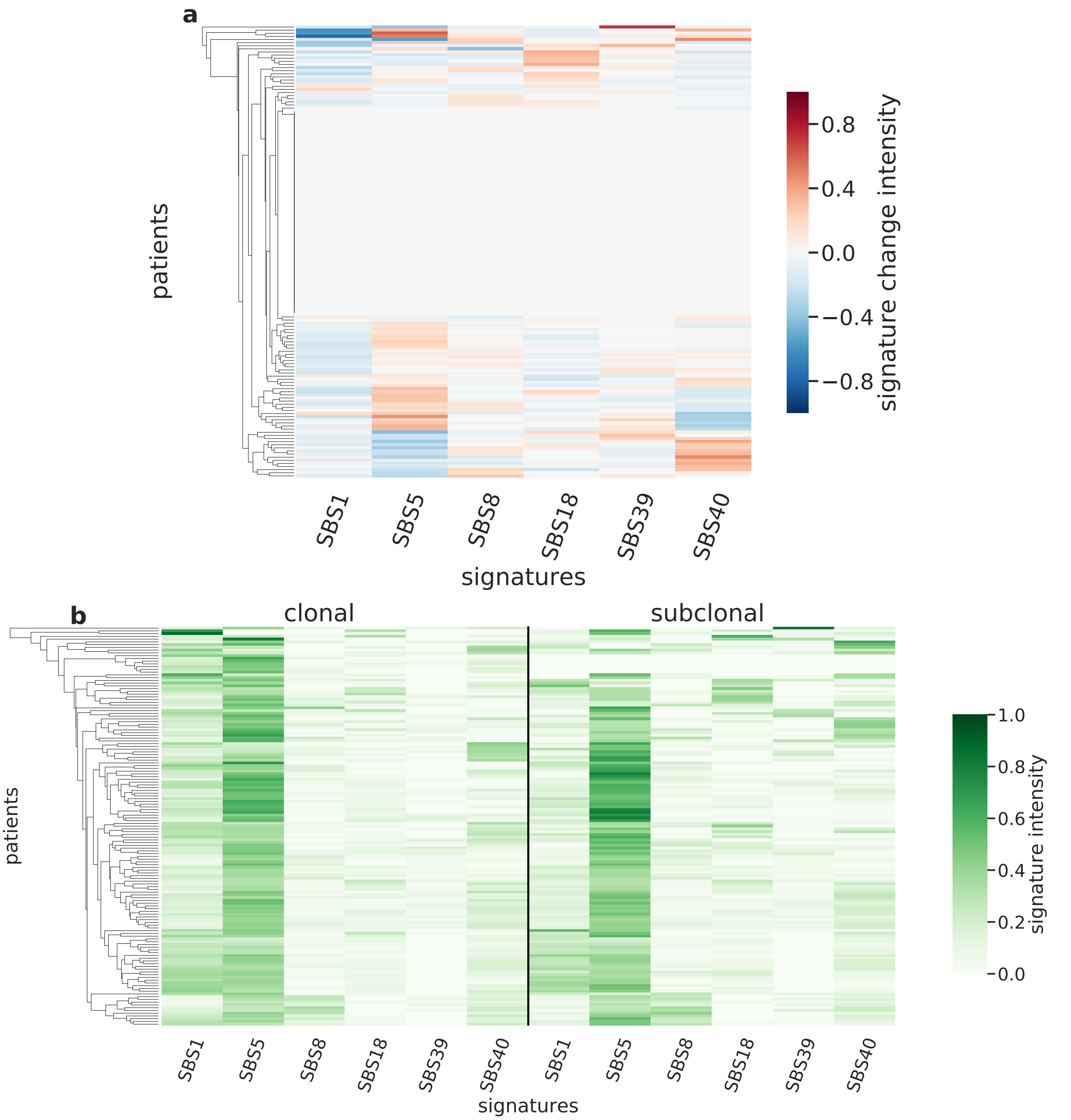

Supplementary Figure 78: **Overview of CloneSig results for the PCAWG sub-cohort CNS-Medullo (n=145).** Panel a: Stratification of patients depending on their pattern of signature change for CNS-Medullo patients (145 patients, including 79 with a significant signature change). The heatmap represents the difference between the signature activity in the largest subclone (in terms of number of mutations) and the clonal mutations (defined as belonging to the clone of highest CCF).Panel b: Stratification of patients depending on their complete pattern of signature exposure. The heatmap represents the signature activity in the largest subclone (in terms of number of mutations) and the clonal mutations (defined as belonging to the clone of highest CCF). Source data are provided as a Source Data file.

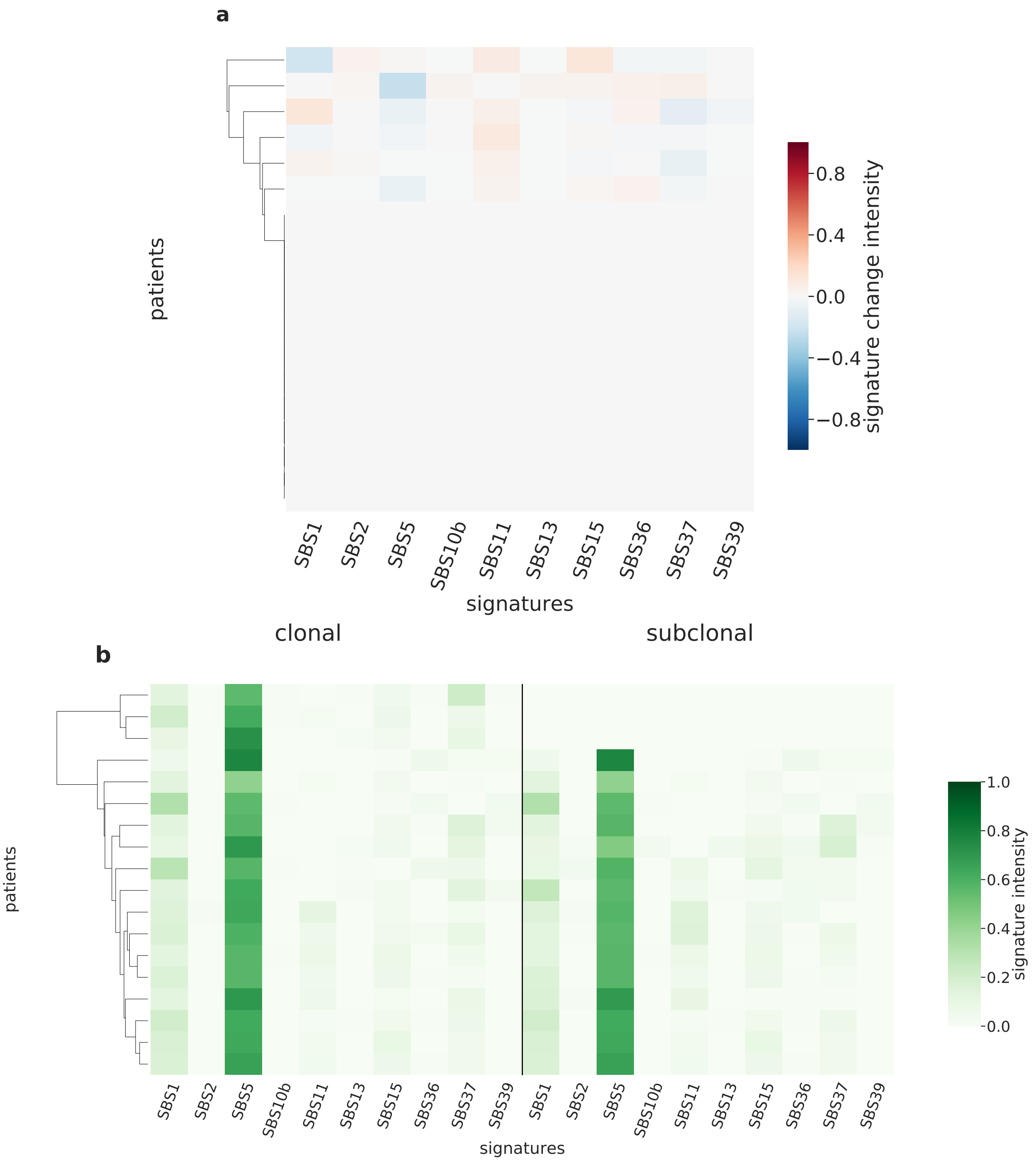

Supplementary Figure 79: **Overview of CloneSig results for the PCAWG sub-cohort CNS-Oligo (n=18).** Panel a: Stratification of patients depending on their pattern of signature change for CNS-Oligo patients (18 patients, including 6 with a significant signature change). The heatmap represents the difference between the signature activity in the largest subclone (in terms of number of mutations) and the clonal mutations (defined as belonging to the clone of highest CCF).Panel b: Stratification of patients depending on their complete pattern of signature exposure. The heatmap represents the signature activity in the largest subclone (in terms of number of mutations) and the clonal mutations (defined as belonging to the clone of highest CCF). Source data are provided as a Source Data file.

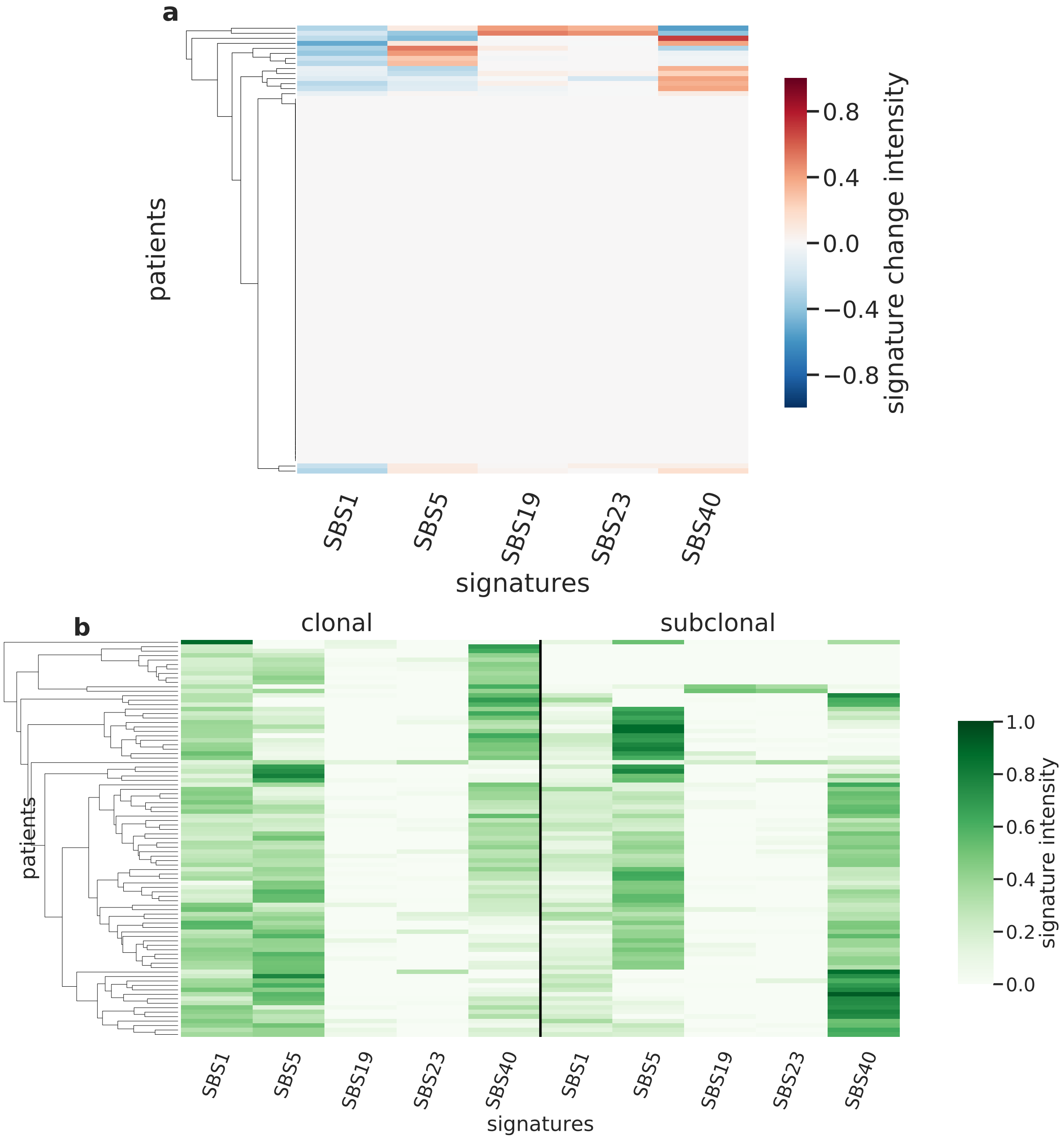

Supplementary Figure 80: **Overview of CloneSig results for the PCAWG sub-cohort CNS-PiloAstro (n=89).** Panel a: Stratification of patients depending on their pattern of signature change for CNS-PiloAstro patients (89 patients, including 16 with a significant signature change). The heatmap represents the difference between the signature activity in the largest subclone (in terms of number of mutations) and the clonal mutations (defined as belonging to the clone of highest CCF).Panel b: Stratification of patients depending on their complete pattern of signature exposure. The heatmap represents the signature activity in the largest subclone (in terms of number of mutations) and the clonal mutations (defined as belonging to the clone of highest CCF). Source data are provided as a Source Data file.

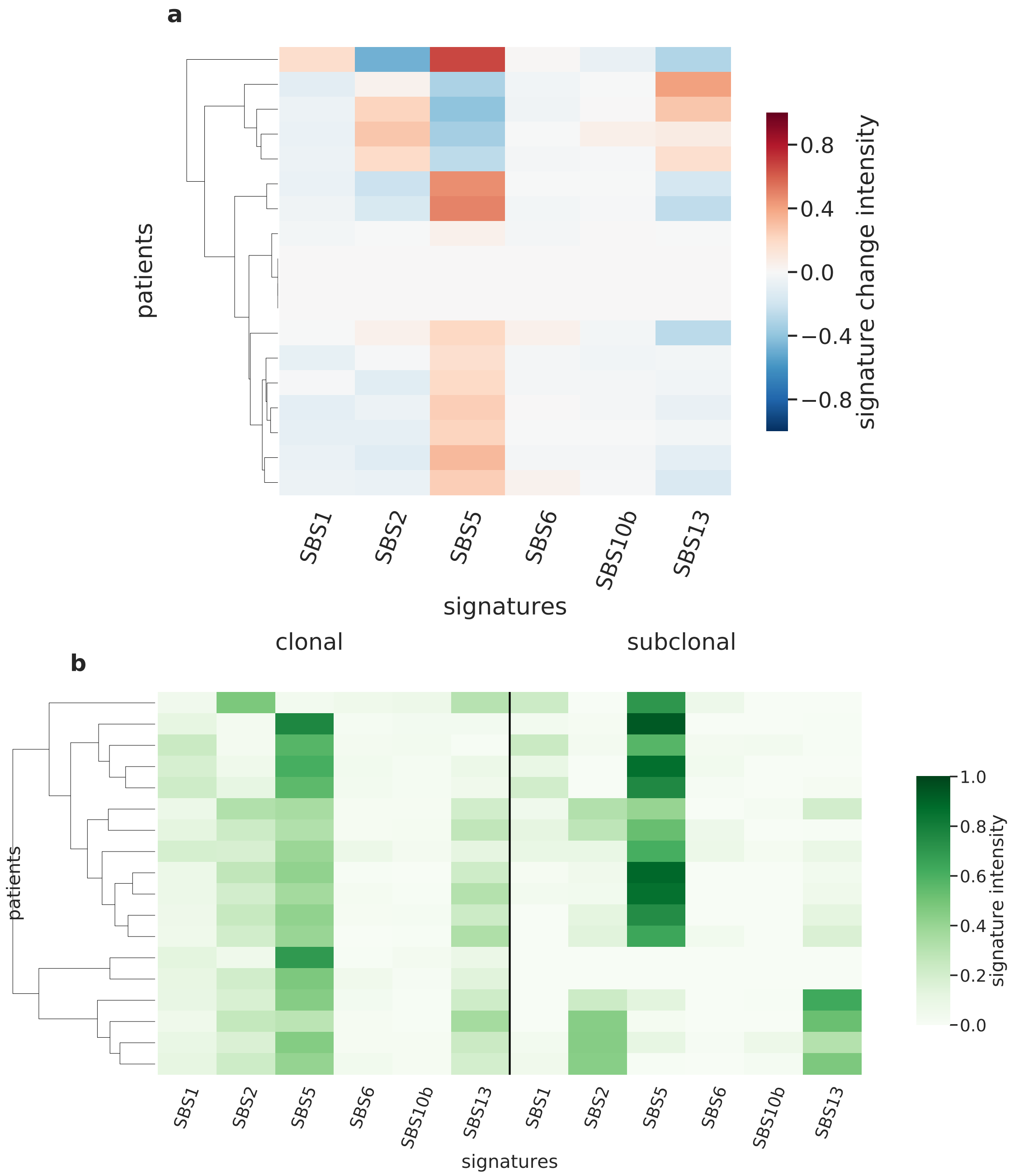

Supplementary Figure 81: **Overview of CloneSig results for the PCAWG sub-cohort Cervix-SCC (n=18).** Panel a: Stratification of patients depending on their pattern of signature change for Cervix-SCC patients (18 patients, including 15 with a significant signature change). The heatmap represents the difference between the signature activity in the largest subclone (in terms of number of mutations) and the clonal mutations (defined as belonging to the clone of highest CCF).Panel b: Stratification of patients depending on their complete pattern of signature exposure. The heatmap represents the signature activity in the largest subclone (in terms of number of mutations) and the clonal mutations (defined as belonging to the clone of highest CCF). Source data are provided as a Source Data file.

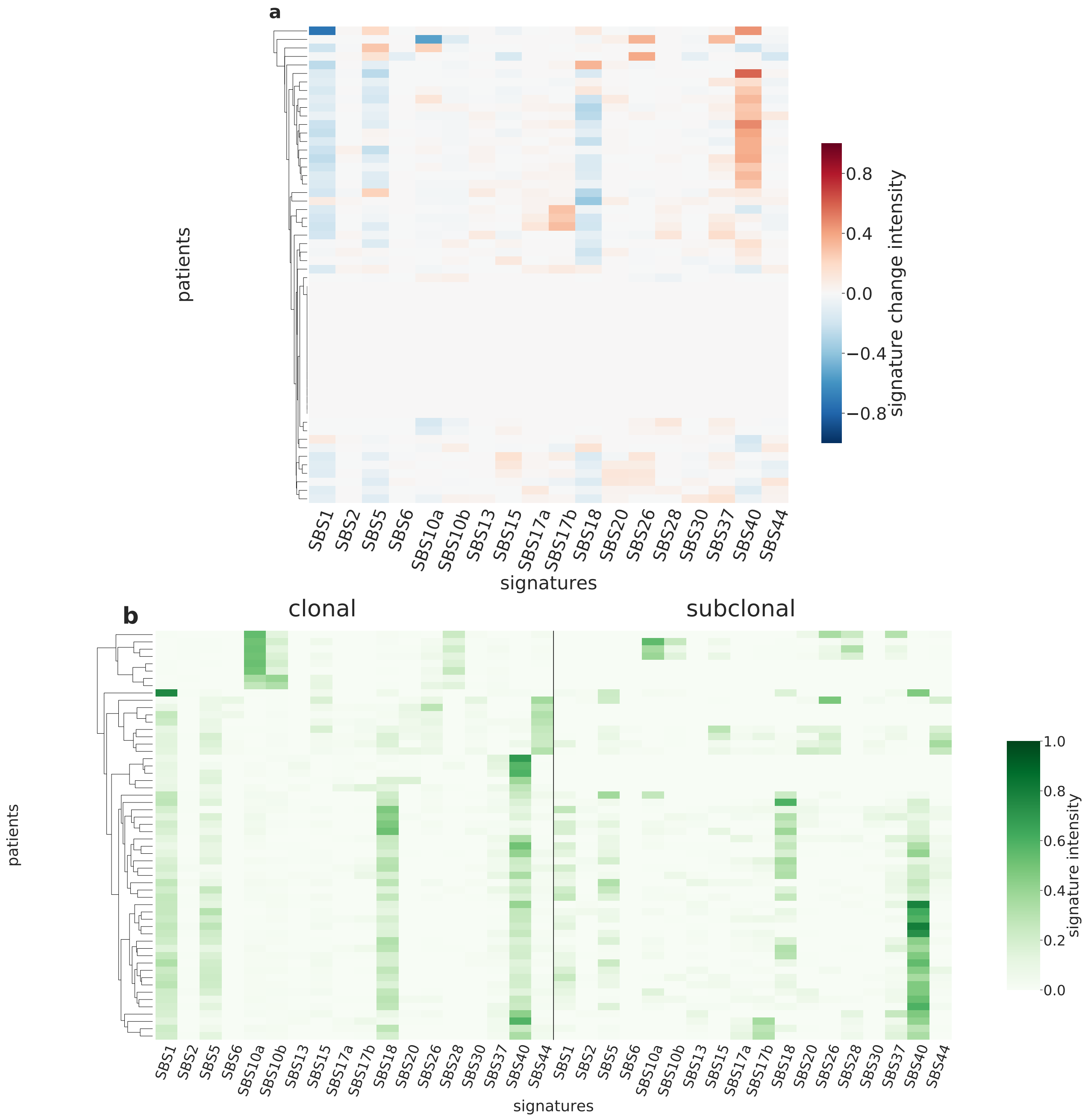

Supplementary Figure 82: **Overview of CloneSig results for the PCAWG sub-cohort ColoRect-AdenoCA (n=56).** Panel a: Stratification of patients depending on their pattern of signature change for ColoRect-AdenoCA patients (56 patients, including 40 with a significant signature change). The heatmap represents the difference between the signature activity in the largest subclone (in terms of number of mutations) and the clonal mutations (defined as belonging to the clone of highest CCF).Panel b: Stratification of patients depending on their complete pattern of signature exposure. The heatmap represents the signature activity in the largest subclone (in terms of number of mutations) and the clonal mutations (defined as belonging to the clone of highest CCF). Source data are provided as a Source Data file.

**a**

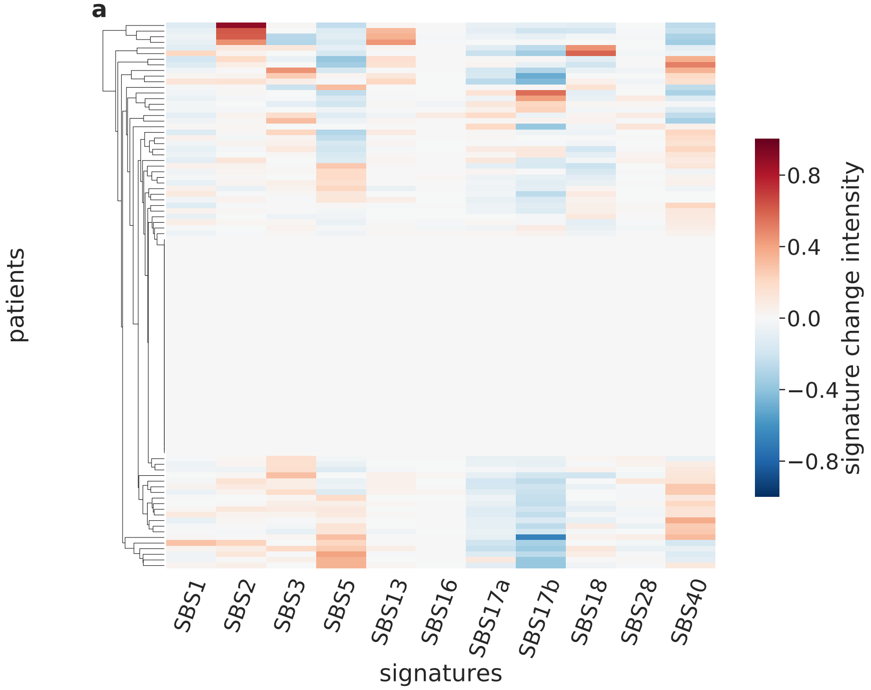

**b**

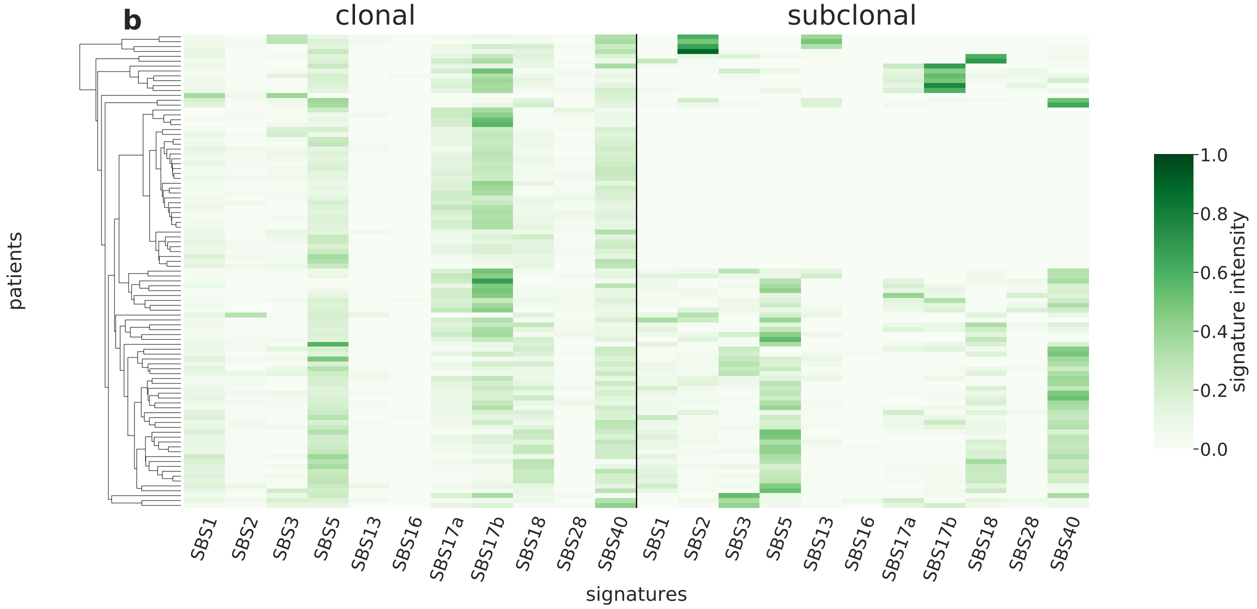

Supplementary Figure 83: **Overview of CloneSig results for the PCAWG sub-cohort Eso-AdenoCA (n=97).** Panel a: Stratification of patients depending on their pattern of signature change for Eso-AdenoCA patients (97 patients, including 58 with a significant signature change). The heatmap represents the difference between the signature activity in the largest subclone (in terms of number of mutations) and the clonal mutations (defined as belonging to the clone of highest CCF).Panel b: Stratification of patients depending on their complete pattern of signature exposure. The heatmap represents the signature activity in the largest subclone (in terms of number of mutations) and the clonal mutations (defined as belonging to the clone of highest CCF). Source data are provided as a Source Data file.

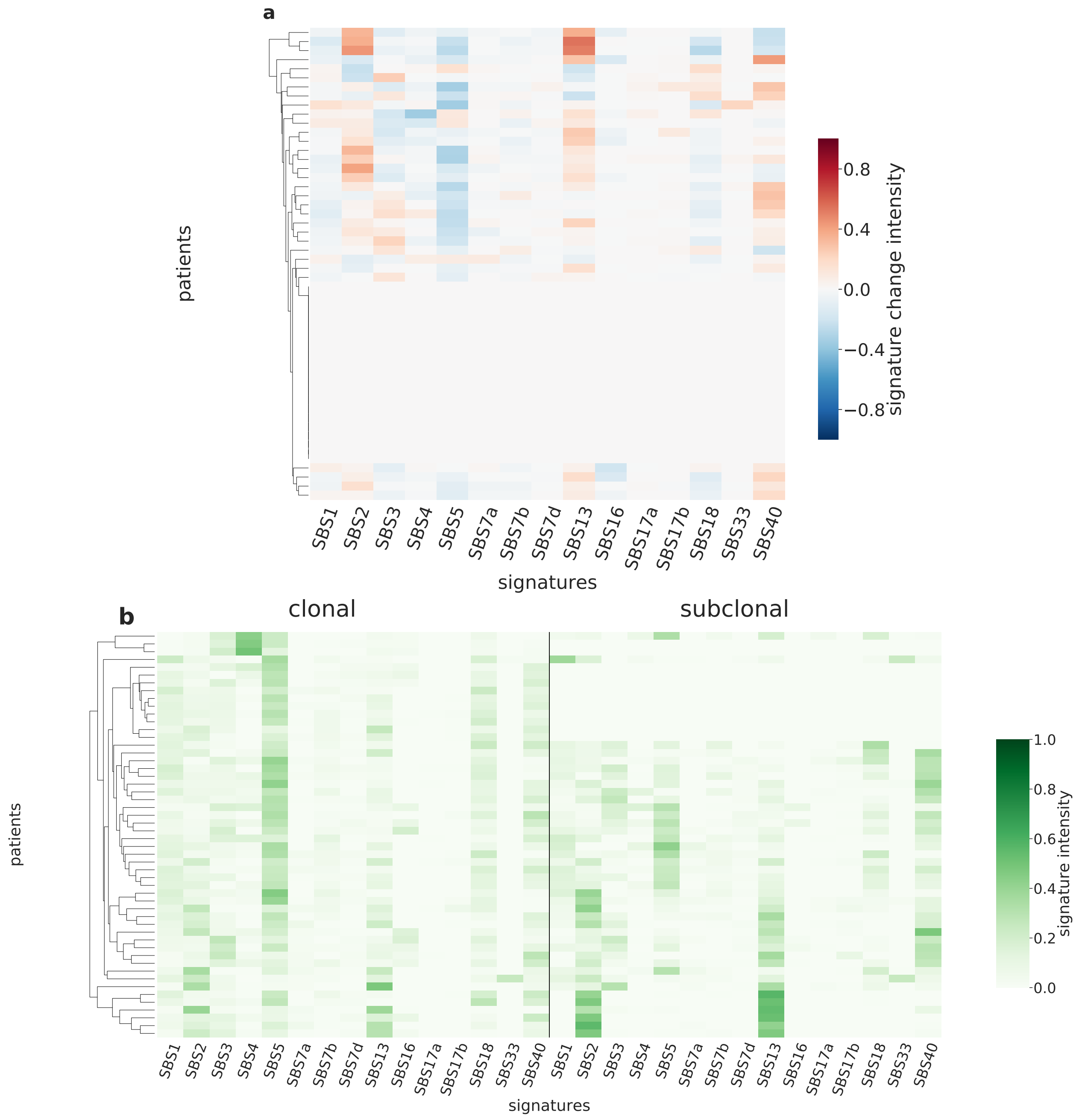

Supplementary Figure 84: **Overview of CloneSig results for the PCAWG sub-cohort Head-SCC (n=52).** Panel a: Stratification of patients depending on their pattern of signature change for Head-SCC patients (52 patients, including 32 with a significant signature change). The heatmap represents the difference between the signature activity in the largest subclone (in terms of number of mutations) and the clonal mutations (defined as belonging to the clone of highest CCF).Panel b: Stratification of patients depending on their complete pattern of signature exposure. The heatmap represents the signature activity in the largest subclone (in terms of number of mutations) and the clonal mutations (defined as belonging to the clone of highest CCF). Source data are provided as a Source Data file.

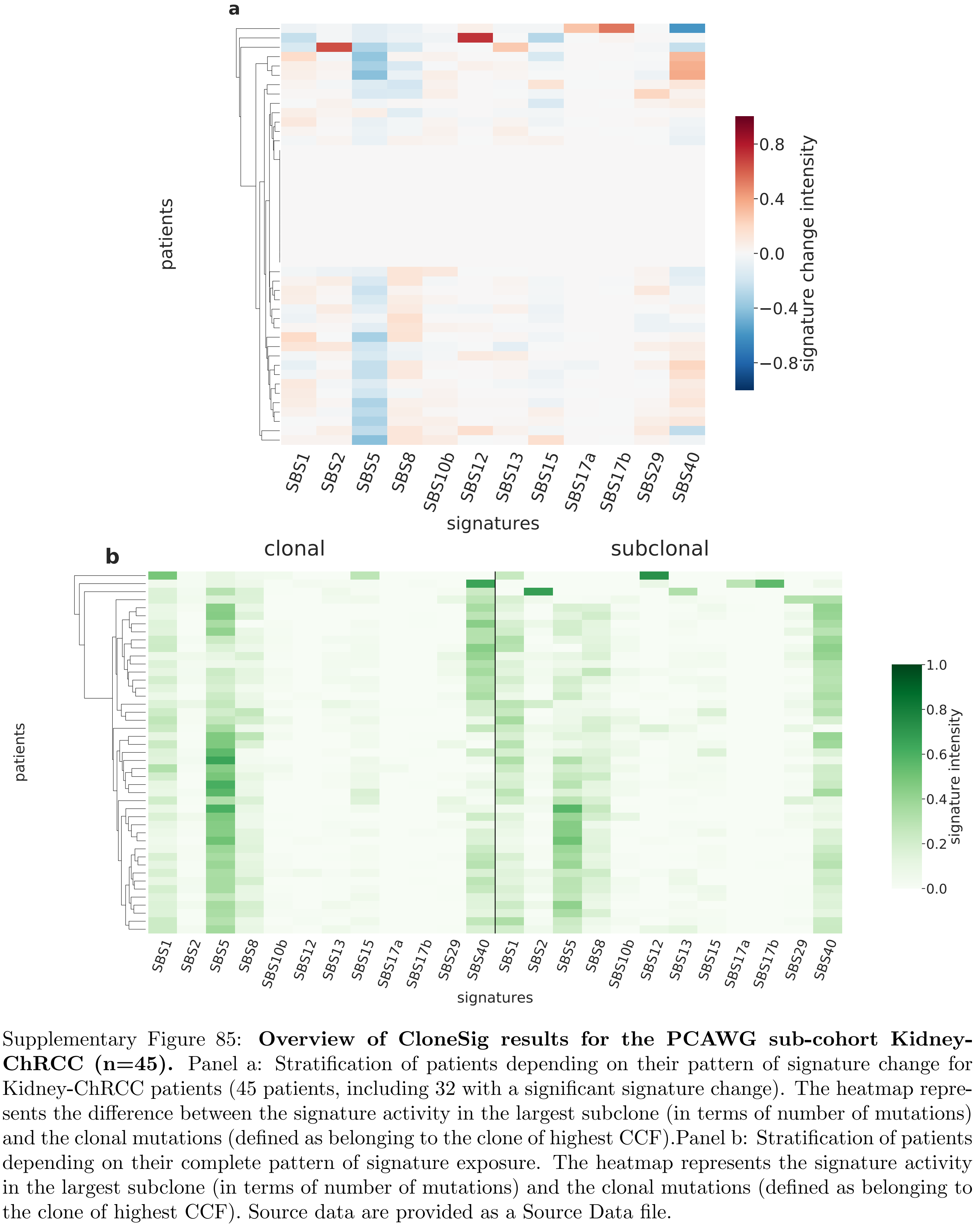

Supplementary Figure 85: **Overview of CloneSig results for the PCAWG sub-cohort Kidney-ChRCC (n=45).** Panel a: Stratification of patients depending on their pattern of signature change for Kidney-ChRCC patients (45 patients, including 32 with a significant signature change). The heatmap represents the difference between the signature activity in the largest subclone (in terms of number of mutations) and the clonal mutations (defined as belonging to the clone of highest CCF).Panel b: Stratification of patients depending on their complete pattern of signature exposure. The heatmap represents the signature activity in the largest subclone (in terms of number of mutations) and the clonal mutations (defined as belonging to the clone of highest CCF). Source data are provided as a Source Data file.

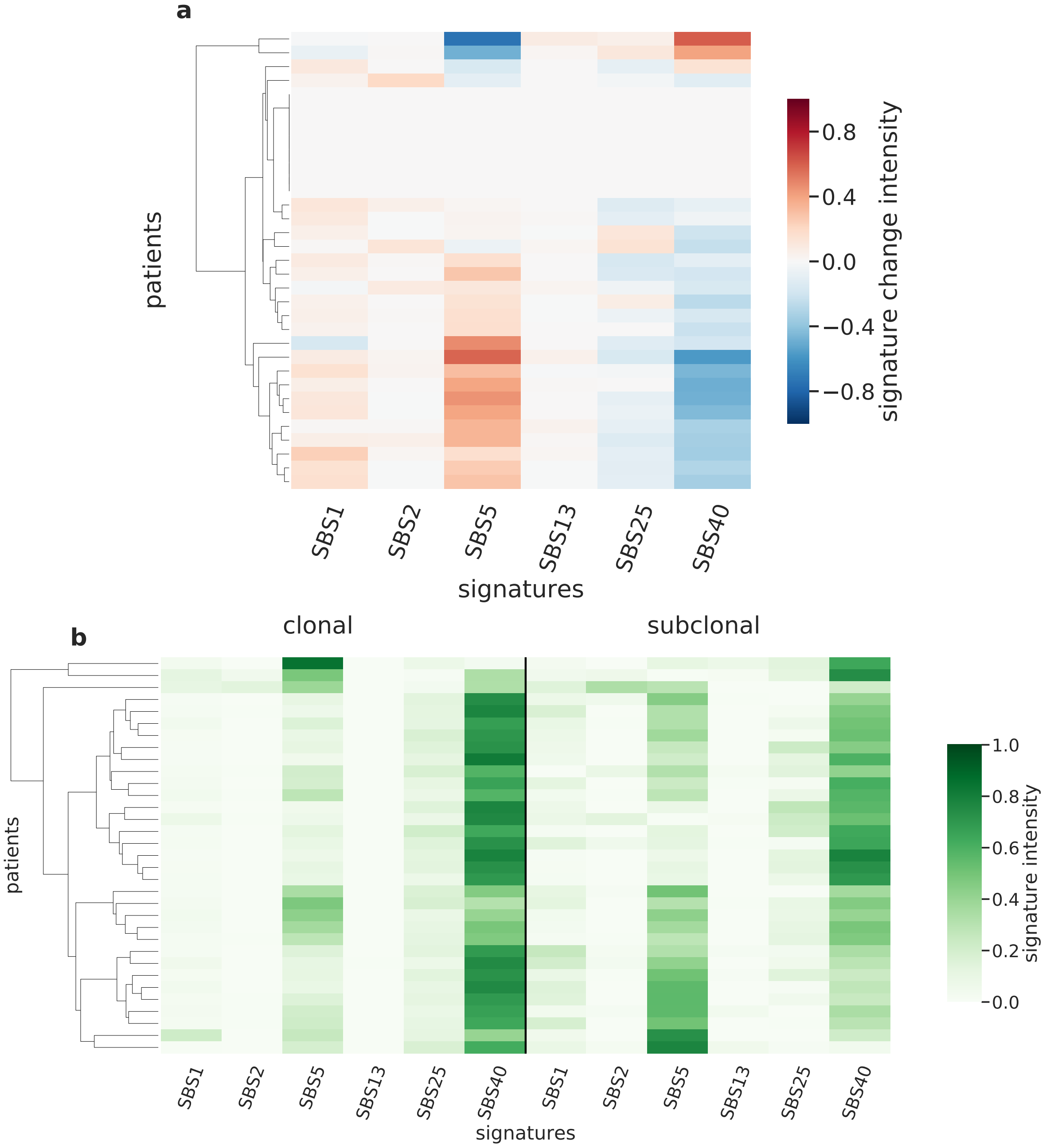

Supplementary Figure 86: **Overview of CloneSig results for the PCAWG sub-cohort Kidney-PRCC (n=33).** Panel a: Stratification of patients depending on their pattern of signature change for Kidney-PRCC patients (33 patients, including 25 with a significant signature change). The heatmap represents the difference between the signature activity in the largest subclone (in terms of number of mutations) and the clonal mutations (defined as belonging to the clone of highest CCF).Panel b: Stratification of patients depending on their complete pattern of signature exposure. The heatmap represents the signature activity in the largest subclone (in terms of number of mutations) and the clonal mutations (defined as belonging to the clone of highest CCF). Source data are provided as a Source Data file.

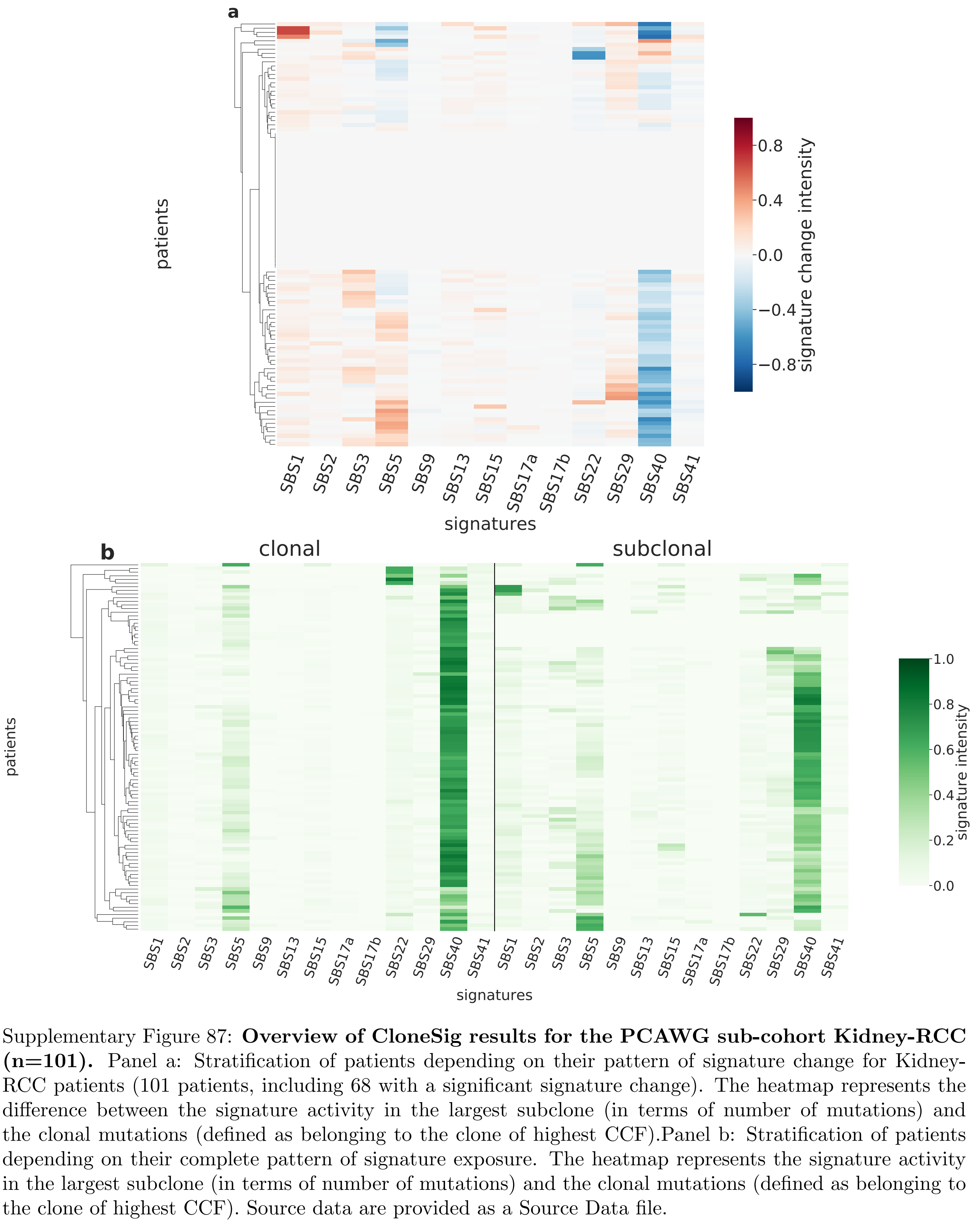

Supplementary Figure 87: **Overview of CloneSig results for the PCAWG sub-cohort Kidney-RCC (n=101).** Panel a: Stratification of patients depending on their pattern of signature change for Kidney-RCC patients (101 patients, including 68 with a significant signature change). The heatmap represents the difference between the signature activity in the largest subclone (in terms of number of mutations) and the clonal mutations (defined as belonging to the clone of highest CCF).Panel b: Stratification of patients depending on their complete pattern of signature exposure. The heatmap represents the signature activity in the largest subclone (in terms of number of mutations) and the clonal mutations (defined as belonging to the clone of highest CCF). Source data are provided as a Source Data file.

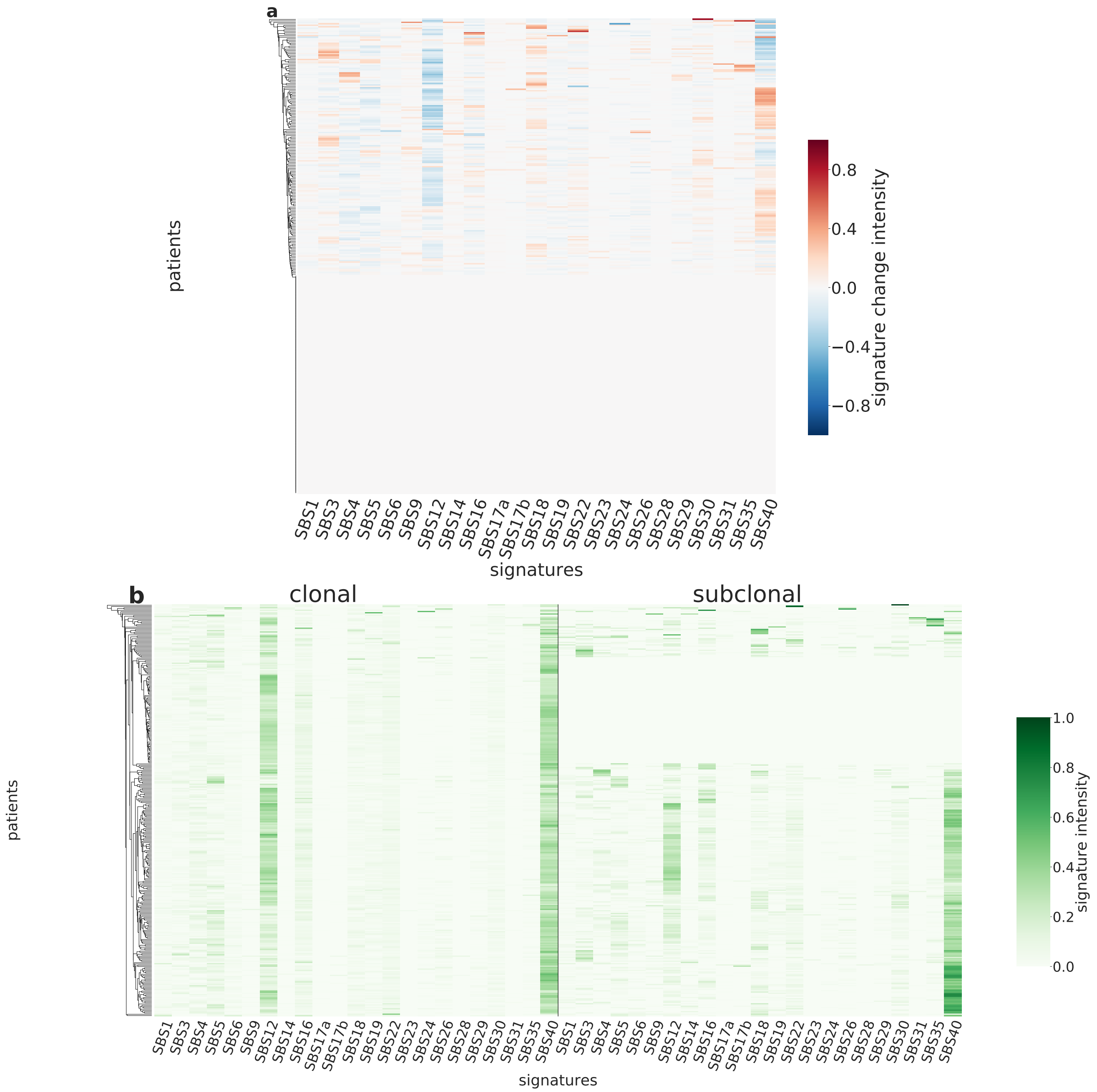

Supplementary Figure 88: **Overview of CloneSig results for the PCAWG sub-cohort Liver-HCC (n=319).** Panel a: Stratification of patients depending on their pattern of signature change for Liver-HCC patients (319 patients, including 172 with a significant signature change). The heatmap represents the difference between the signature activity in the largest subclone (in terms of number of mutations) and the clonal mutations (defined as belonging to the clone of highest CCF).Panel b: Stratification of patients depending on their complete pattern of signature exposure. The heatmap represents the signature activity in the largest subclone (in terms of number of mutations) and the clonal mutations (defined as belonging to the clone of highest CCF). Source data are provided as a Source Data file.

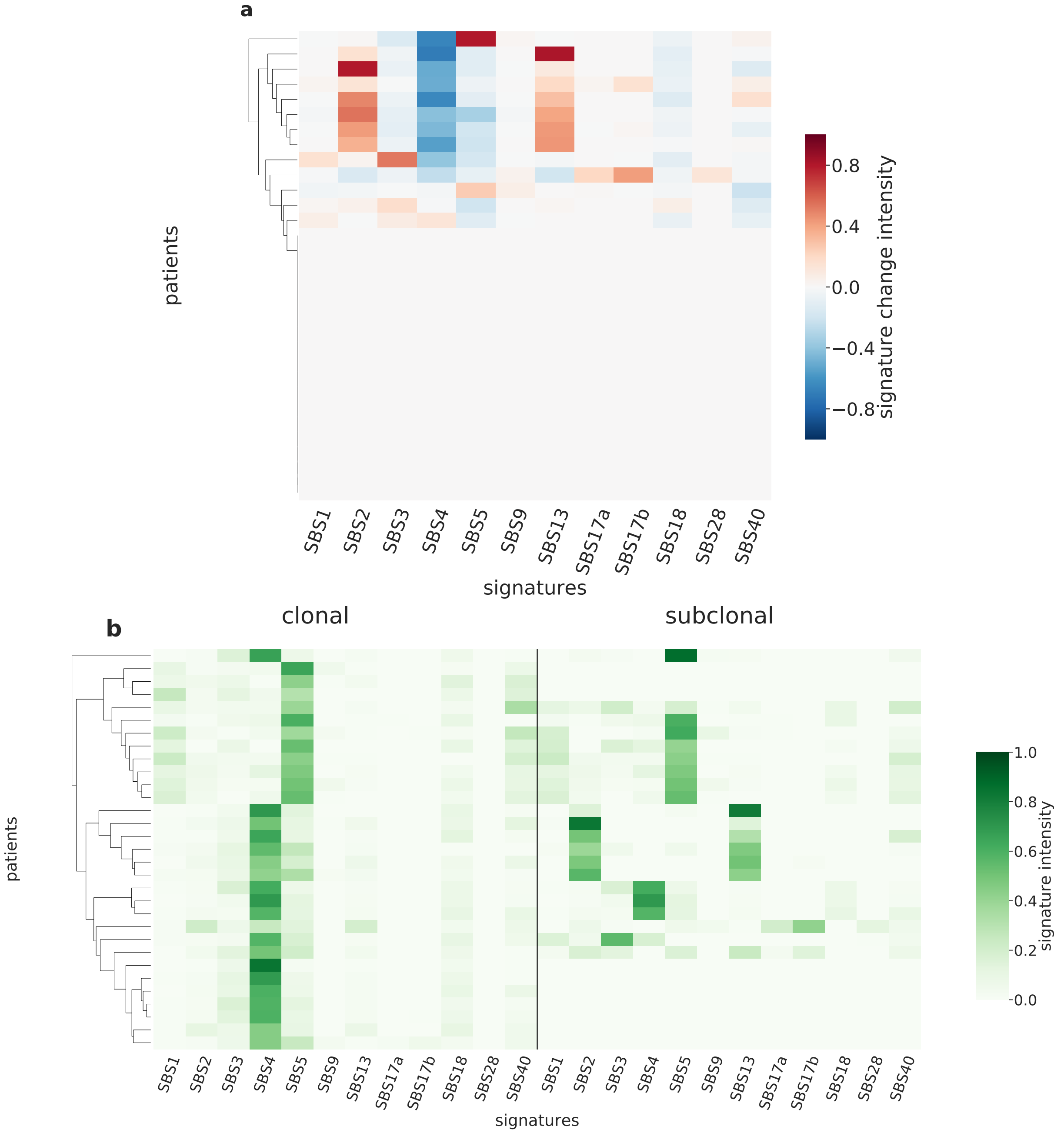

Supplementary Figure 89: **Overview of CloneSig results for the PCAWG sub-cohort Lung-AdenoCA (n=31).** Panel a: Stratification of patients depending on their pattern of signature change for Lung-AdenoCA patients (31 patients, including 13 with a significant signature change). The heatmap represents the difference between the signature activity in the largest subclone (in terms of number of mutations) and the clonal mutations (defined as belonging to the clone of highest CCF).Panel b: Stratification of patients depending on their complete pattern of signature exposure. The heatmap represents the signature activity in the largest subclone (in terms of number of mutations) and the clonal mutations (defined as belonging to the clone of highest CCF). Source data are provided as a Source Data file.

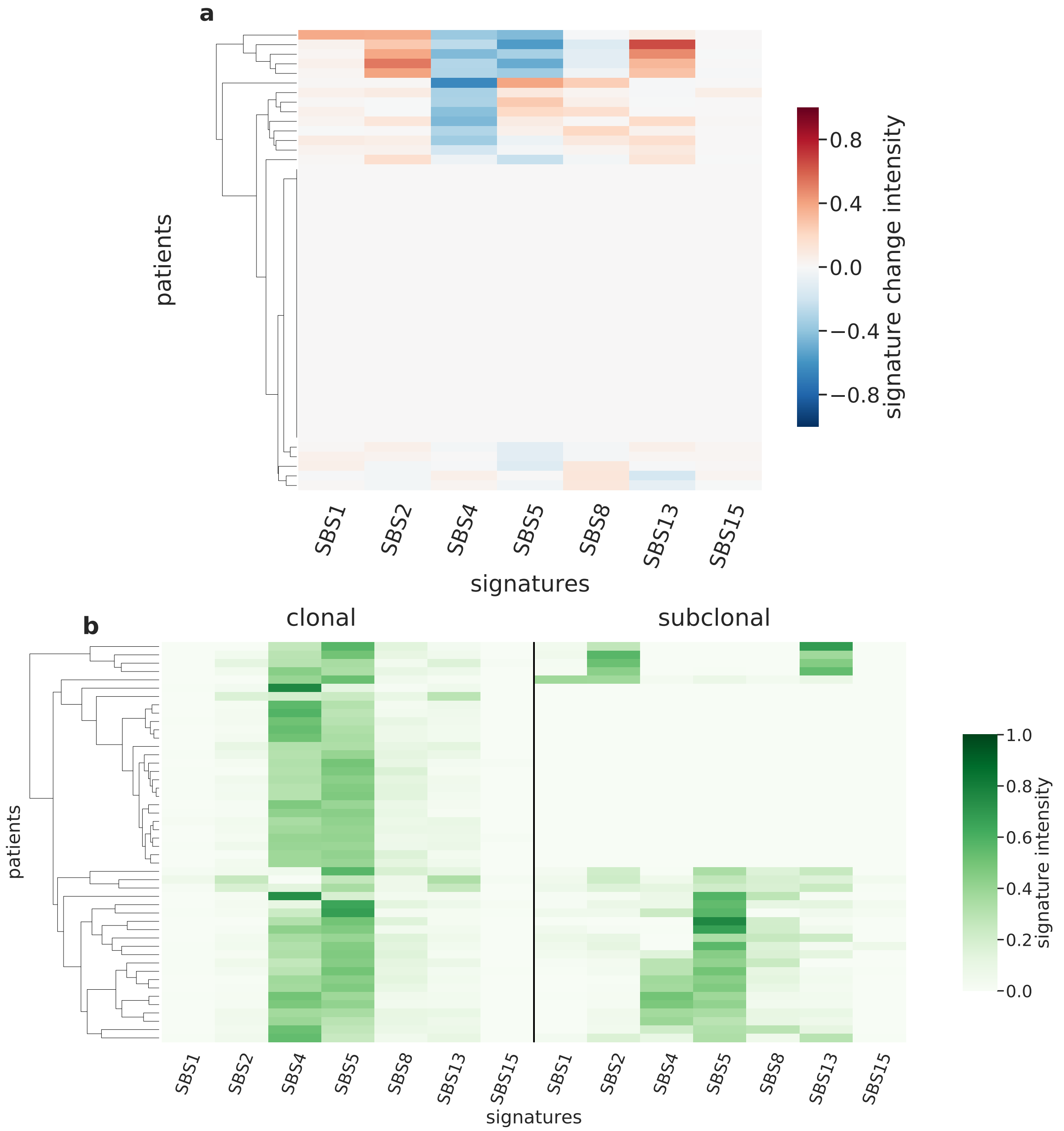

Supplementary Figure 90: **Overview of CloneSig results for the PCAWG sub-cohort Lung-SCC (n=48).** Panel a: Stratification of patients depending on their pattern of signature change for Lung-SCC patients (48 patients, including 19 with a significant signature change). The heatmap represents the difference between the signature activity in the largest subclone (in terms of number of mutations) and the clonal mutations (defined as belonging to the clone of highest CCF).Panel b: Stratification of patients depending on their complete pattern of signature exposure. The heatmap represents the signature activity in the largest subclone (in terms of number of mutations) and the clonal mutations (defined as belonging to the clone of highest CCF). Source data are provided as a Source Data file.

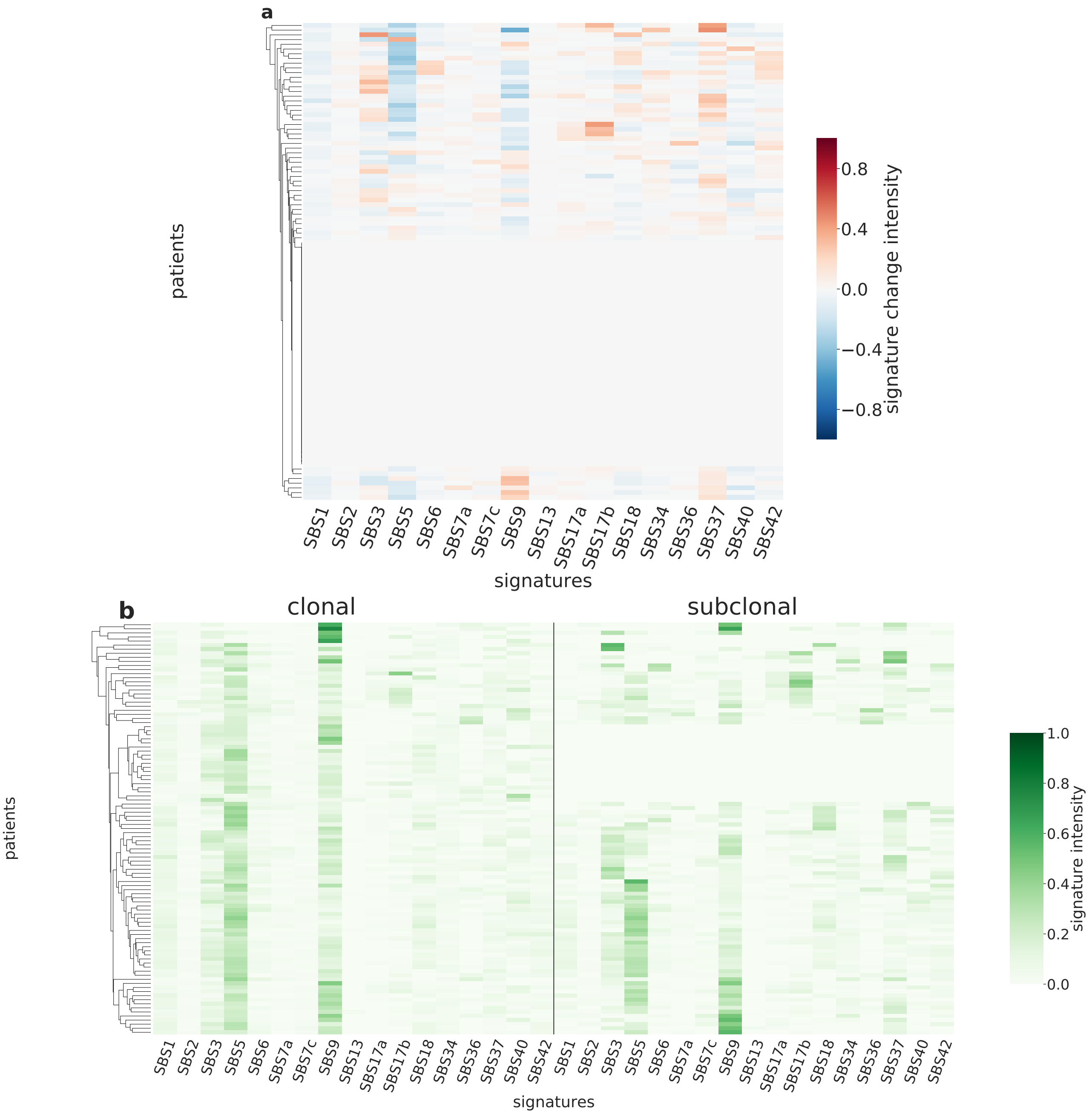

Supplementary Figure 91: **Overview of CloneSig results for the PCAWG sub-cohort Lymph-BNHL (n=101).** Panel a: Stratification of patients depending on their pattern of signature change for Lymph-BNHL patients (101 patients, including 53 with a significant signature change). The heatmap represents the difference between the signature activity in the largest subclone (in terms of number of mutations) and the clonal mutations (defined as belonging to the clone of highest CCF).Panel b: Stratification of patients depending on their complete pattern of signature exposure. The heatmap represents the signature activity in the largest subclone (in terms of number of mutations) and the clonal mutations (defined as belonging to the clone of highest CCF). Source data are provided as a Source Data file.

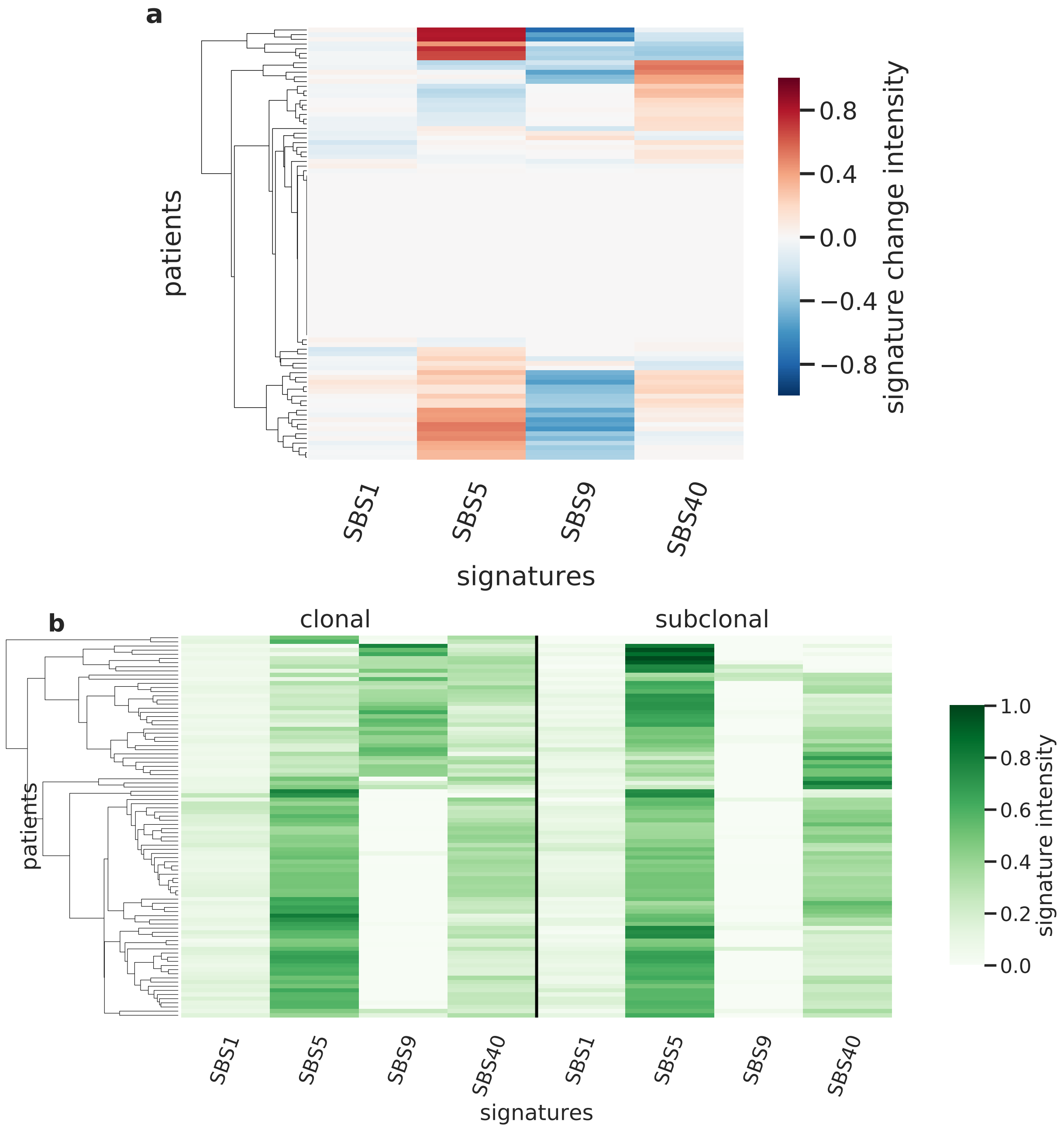

Supplementary Figure 92: **Overview of CloneSig results for the PCAWG sub-cohort Lymph-CLL (n=92).** Panel a: Stratification of patients depending on their pattern of signature change for Lymph-CLL patients (92 patients, including 57 with a significant signature change). The heatmap represents the difference between the signature activity in the largest subclone (in terms of number of mutations) and the clonal mutations (defined as belonging to the clone of highest CCF).Panel b: Stratification of patients depending on their complete pattern of signature exposure. The heatmap represents the signature activity in the largest subclone (in terms of number of mutations) and the clonal mutations (defined as belonging to the clone of highest CCF). Source data are provided as a Source Data file.

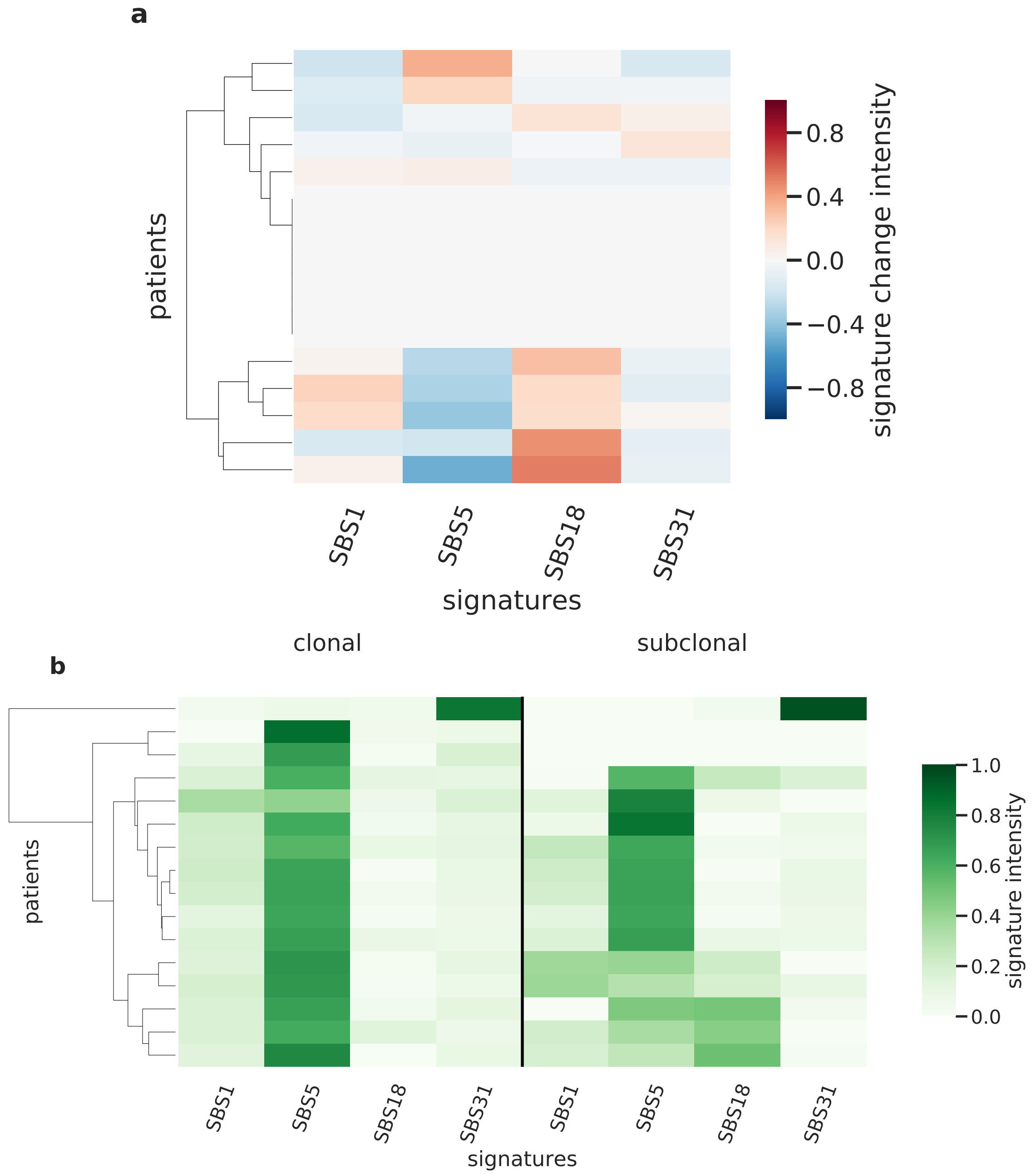

Supplementary Figure 93: **Overview of CloneSig results for the PCAWG sub-cohort Myeloid-AML (n=16).** Panel a: Stratification of patients depending on their pattern of signature change for Myeloid-AML patients (16 patients, including 10 with a significant signature change). The heatmap represents the difference between the signature activity in the largest subclone (in terms of number of mutations) and the clonal mutations (defined as belonging to the clone of highest CCF).Panel b: Stratification of patients depending on their complete pattern of signature exposure. The heatmap represents the signature activity in the largest subclone (in terms of number of mutations) and the clonal mutations (defined as belonging to the clone of highest CCF). Source data are provided as a Source Data file.

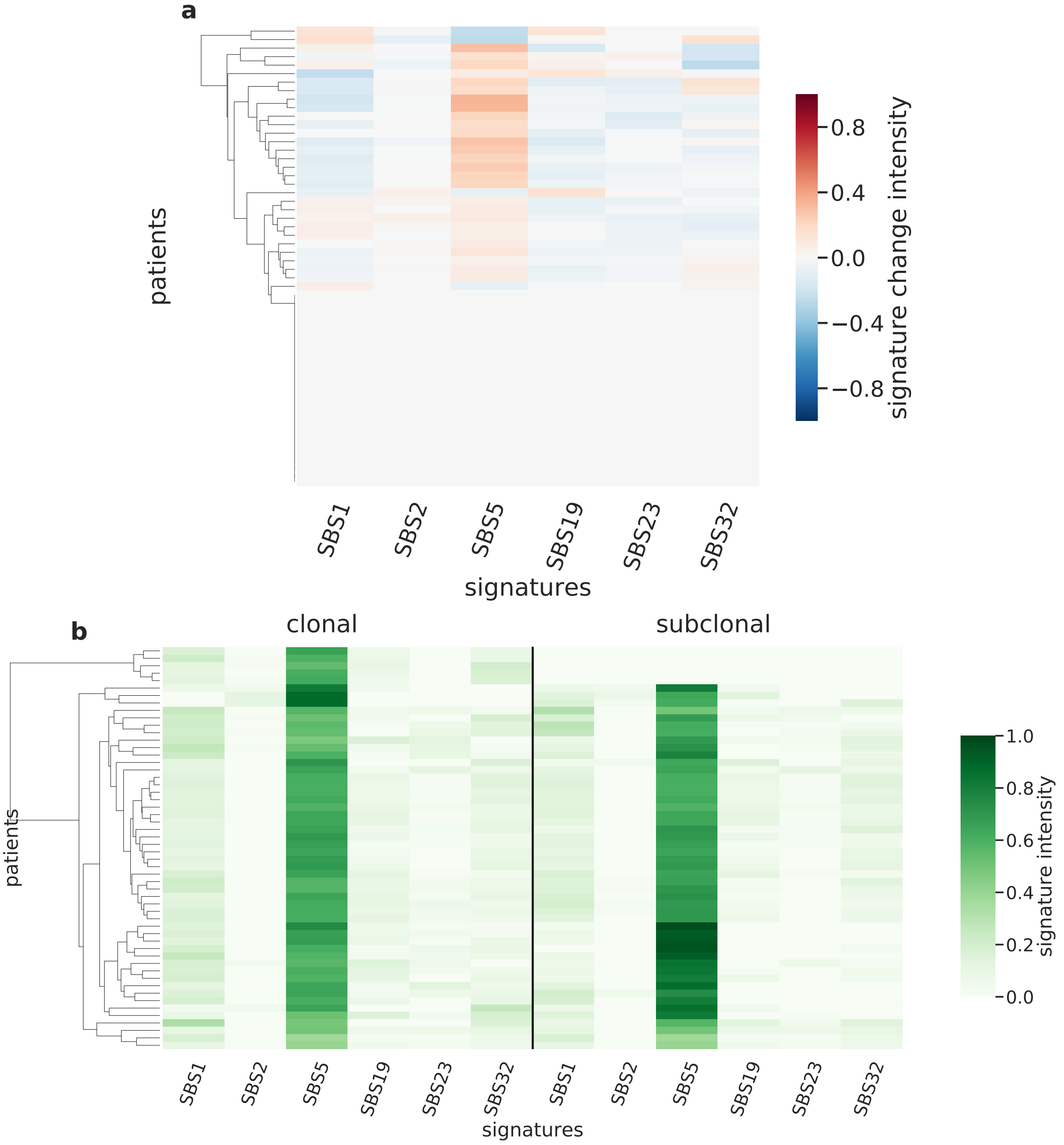

Supplementary Figure 94: **Overview of CloneSig results for the PCAWG sub-cohort Myeloid-MDS/MPN (n=54).** Panel a: Stratification of patients depending on their pattern of signature change for Myeloid-MDS/MPN patients (54 patients, including 31 with a significant signature change). The heatmap represents the difference between the signature activity in the largest subclone (in terms of number of mutations) and the clonal mutations (defined as belonging to the clone of highest CCF).Panel b: Stratification of patients depending on their complete pattern of signature exposure. The heatmap represents the signature activity in the largest subclone (in terms of number of mutations) and the clonal mutations (defined as belonging to the clone of highest CCF). Source data are provided as a Source Data file.

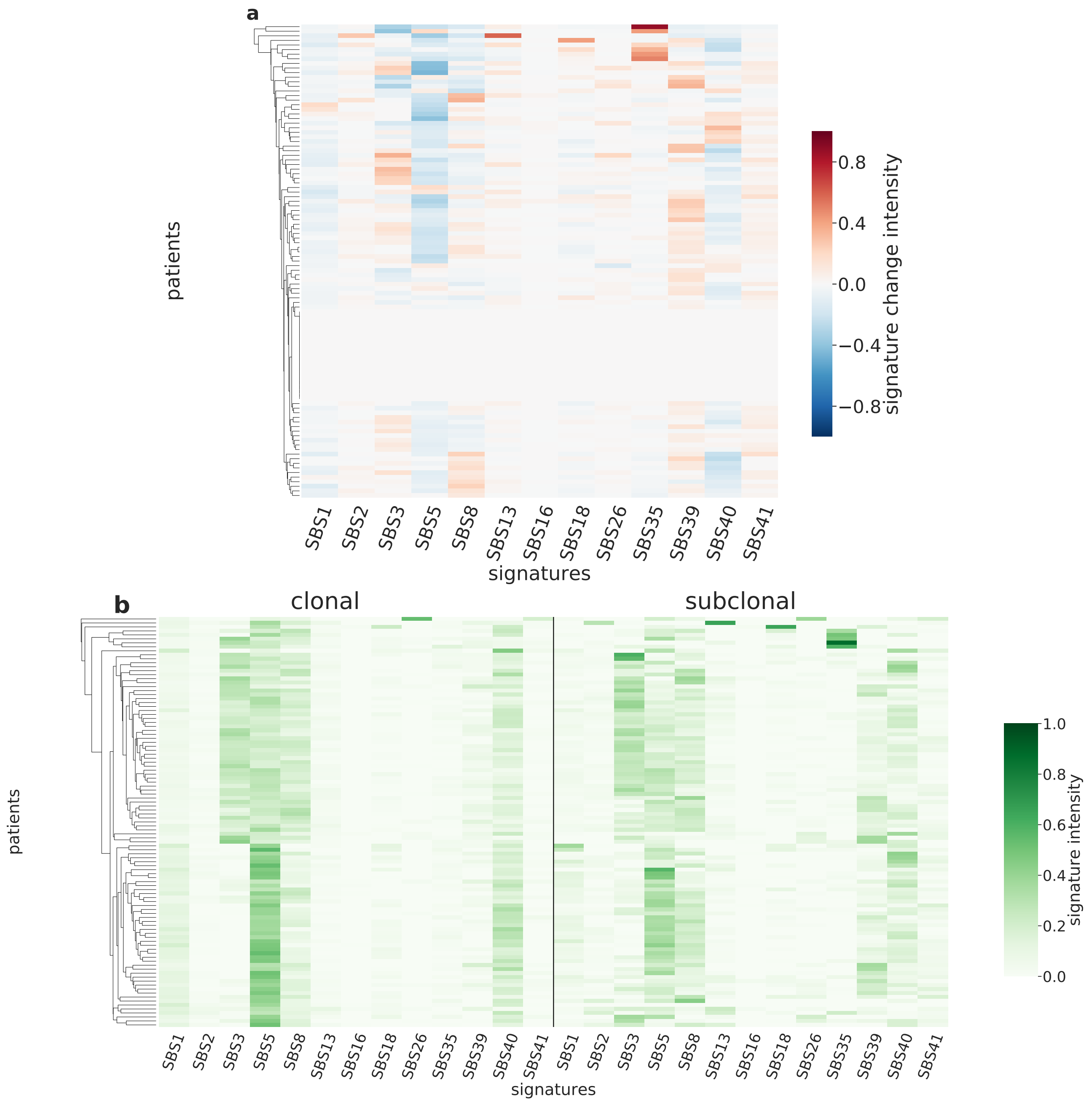

Supplementary Figure 95: **Overview of CloneSig results for the PCAWG sub-cohort Ovary-AdenoCA (n=103).** Panel a: Stratification of patients depending on their pattern of signature change for Ovary-AdenoCA patients (103 patients, including 83 with a significant signature change). The heatmap represents the difference between the signature activity in the largest subclone (in terms of number of mutations) and the clonal mutations (defined as belonging to the clone of highest CCF).Panel b: Stratification of patients depending on their complete pattern of signature exposure. The heatmap represents the signature activity in the largest subclone (in terms of number of mutations) and the clonal mutations (defined as belonging to the clone of highest CCF). Source data are provided as a Source Data file.

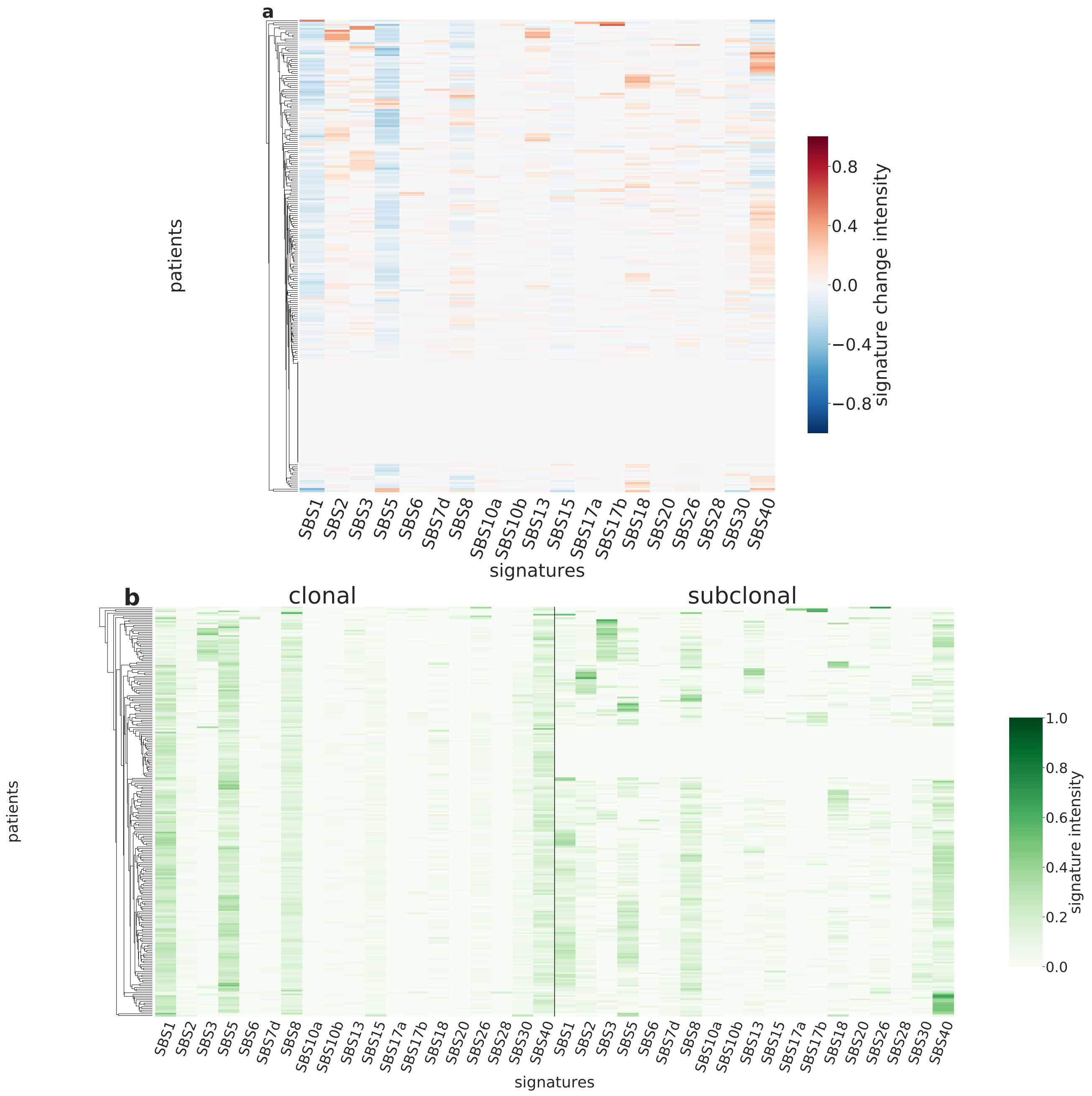

Supplementary Figure 96: **Overview of CloneSig results for the PCAWG sub-cohort Panc-AdenoCA (n=233).** Panel a: Stratification of patients depending on their pattern of signature change for Panc-AdenoCA patients (233 patients, including 182 with a significant signature change). The heatmap represents the difference between the signature activity in the largest subclone (in terms of number of mutations) and the clonal mutations (defined as belonging to the clone of highest CCF).Panel b: Stratification of patients depending on their complete pattern of signature exposure. The heatmap represents the signature activity in the largest subclone (in terms of number of mutations) and the clonal mutations (defined as belonging to the clone of highest CCF). Source data are provided as a Source Data file.

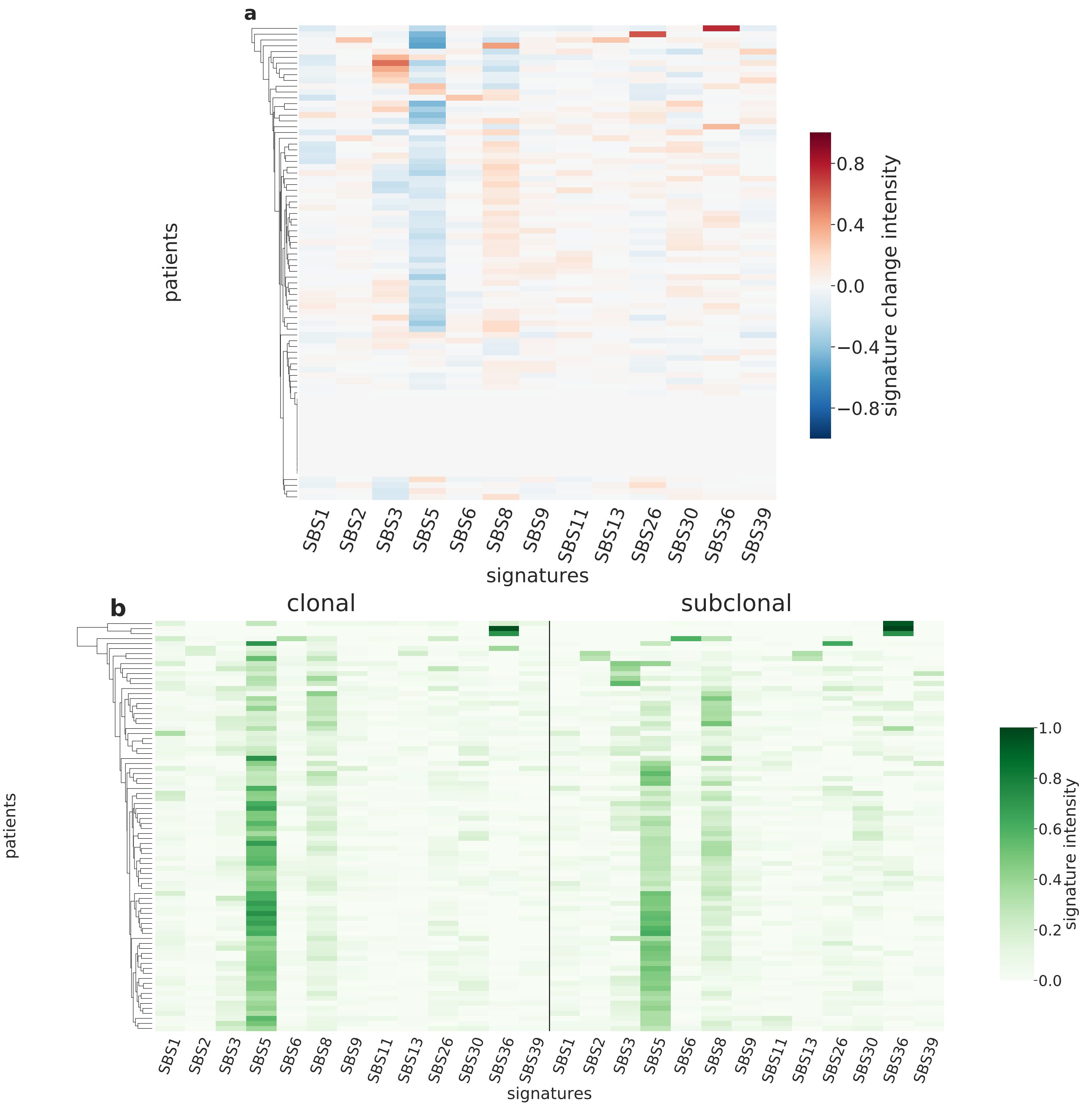

Supplementary Figure 97: **Overview of CloneSig results for the PCAWG sub-cohort Panc-Endocrine (n=82).** Panel a: Stratification of patients depending on their pattern of signature change for Panc-Endocrine patients (82 patients, including 68 with a significant signature change). The heatmap represents the difference between the signature activity in the largest subclone (in terms of number of mutations) and the clonal mutations (defined as belonging to the clone of highest CCF).Panel b: Stratification of patients depending on their complete pattern of signature exposure. The heatmap represents the signature activity in the largest subclone (in terms of number of mutations) and the clonal mutations (defined as belonging to the clone of highest CCF). Source data are provided as a Source Data file.

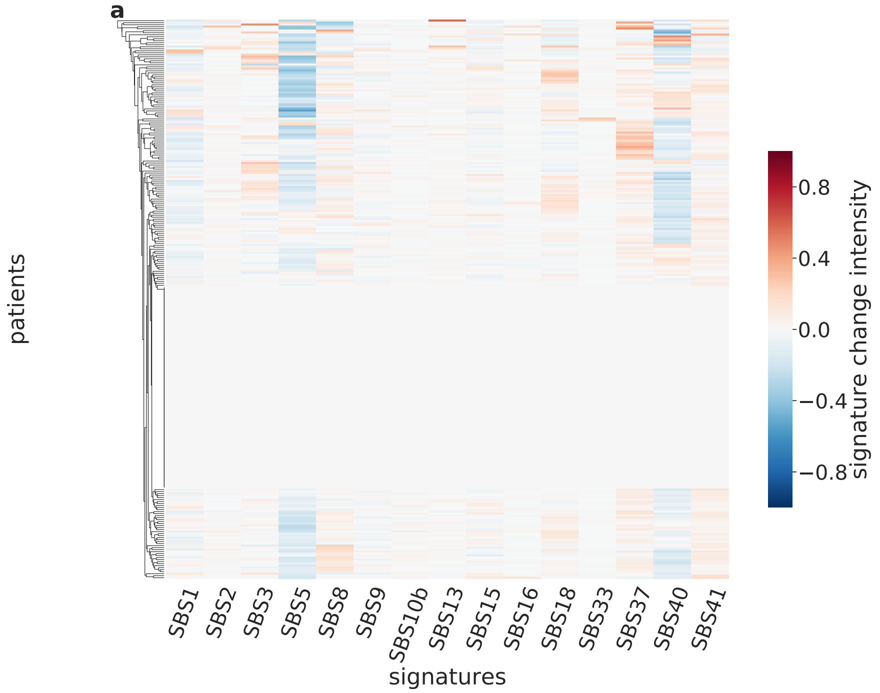

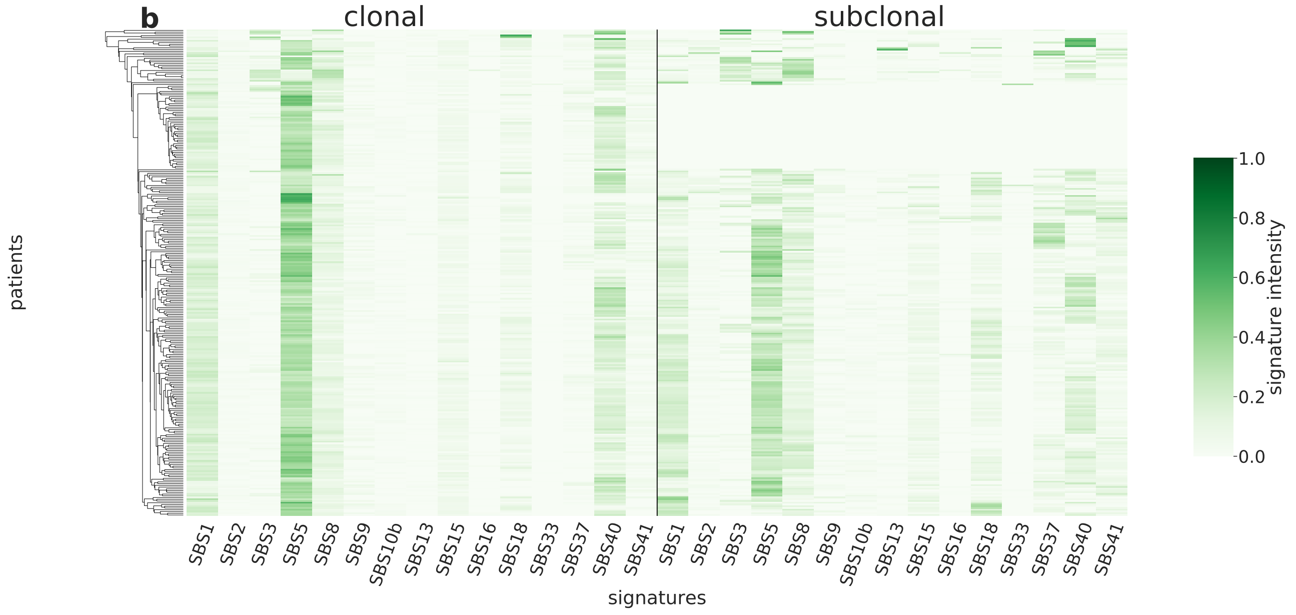

Supplementary Figure 98: **Overview of CloneSig results for the PCAWG sub-cohort Prost-AdenoCA (n=279).** Panel a: Stratification of patients depending on their pattern of signature change for Prost-AdenoCA patients (279 patients, including 178 with a significant signature change). The heatmap represents the difference between the signature activity in the largest subclone (in terms of number of mutations) and the clonal mutations (defined as belonging to the clone of highest CCF).Panel b: Stratification of patients depending on their complete pattern of signature exposure. The heatmap represents the signature activity in the largest subclone (in terms of number of mutations) and the clonal mutations (defined as belonging to the clone of highest CCF). Source data are provided as a Source Data file.

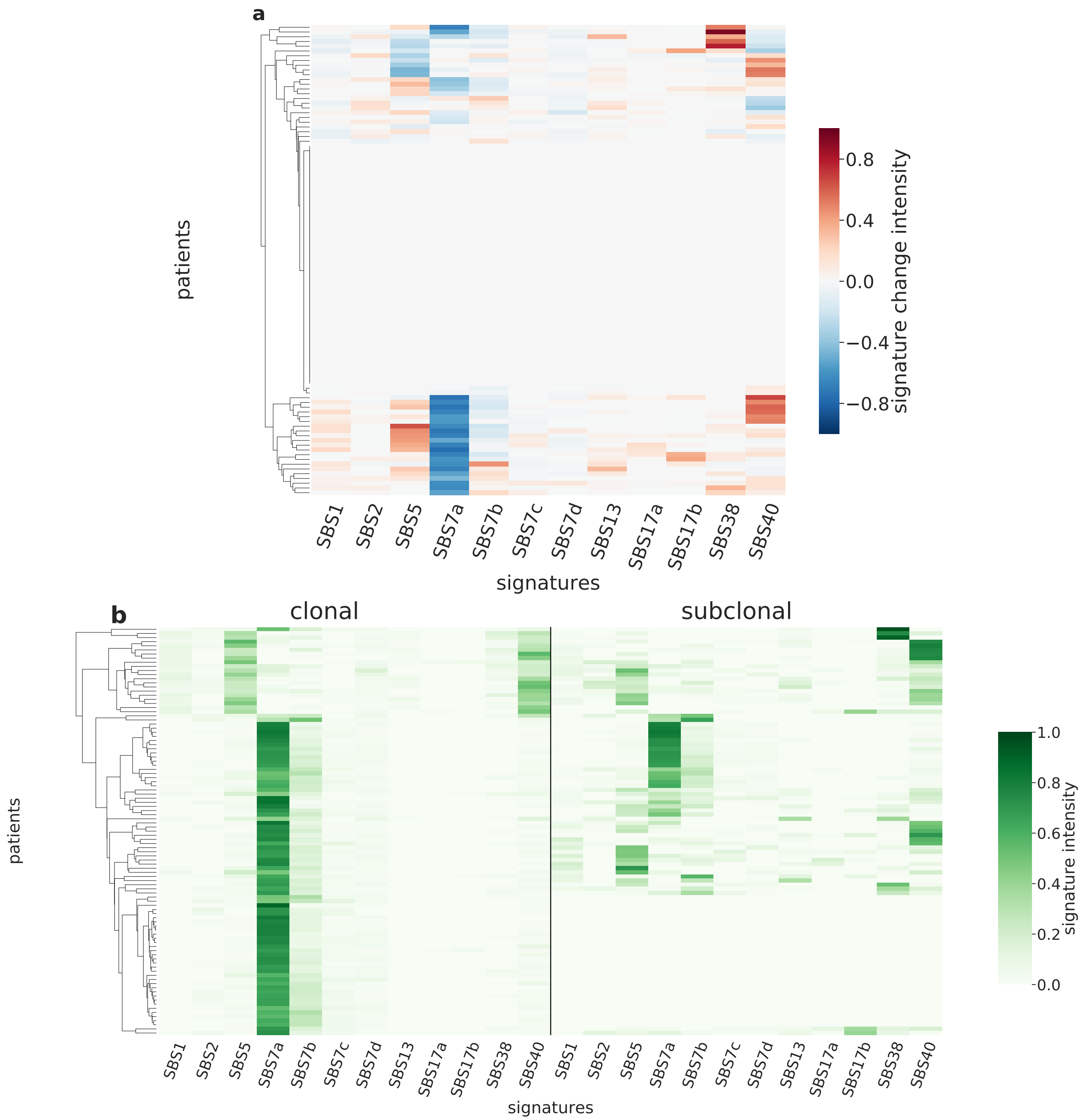

Supplementary Figure 99: **Overview of CloneSig results for the PCAWG sub-cohort Skin-Melanoma (n=99).** Panel a: Stratification of patients depending on their pattern of signature change for Skin-Melanoma patients (99 patients, including 48 with a significant signature change). The heatmap represents the difference between the signature activity in the largest subclone (in terms of number of mutations) and the clonal mutations (defined as belonging to the clone of highest CCF).Panel b: Stratification of patients depending on their complete pattern of signature exposure. The heatmap represents the signature activity in the largest subclone (in terms of number of mutations) and the clonal mutations (defined as belonging to the clone of highest CCF). Source data are provided as a Source Data file.

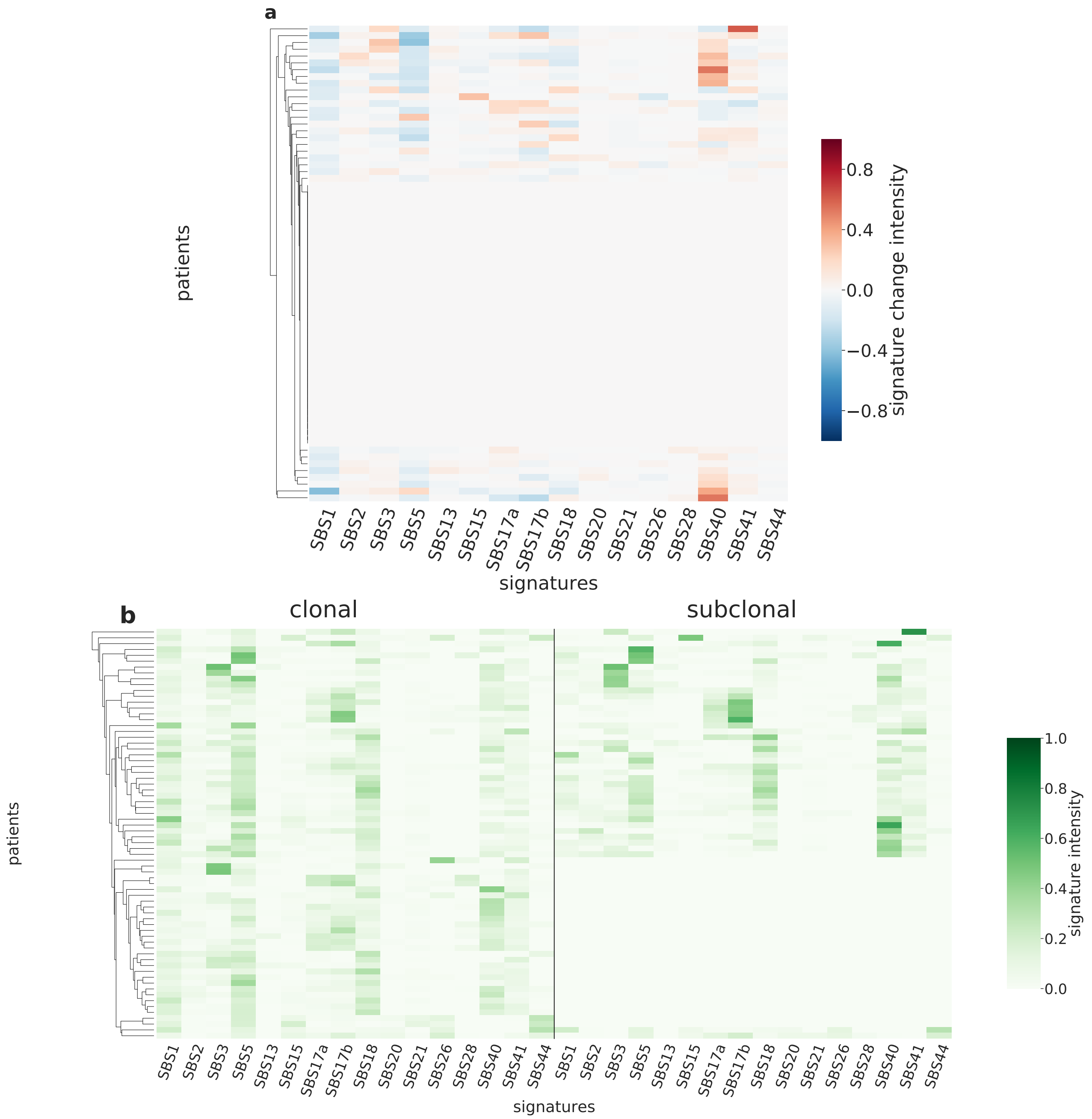

Supplementary Figure 100: **Overview of CloneSig results for the PCAWG sub-cohort Stomach-AdenoCA (n=70).** Panel a: Stratification of patients depending on their pattern of signature change for Stomach-AdenoCA patients (70 patients, including 31 with a significant signature change). The heatmap represents the difference between the signature activity in the largest subclone (in terms of number of mutations) and the clonal mutations (defined as belonging to the clone of highest CCF).Panel b: Stratification of patients depending on their complete pattern of signature exposure. The heatmap represents the signature activity in the largest subclone (in terms of number of mutations) and the clonal mutations (defined as belonging to the clone of highest CCF). Source data are provided as a Source Data file.

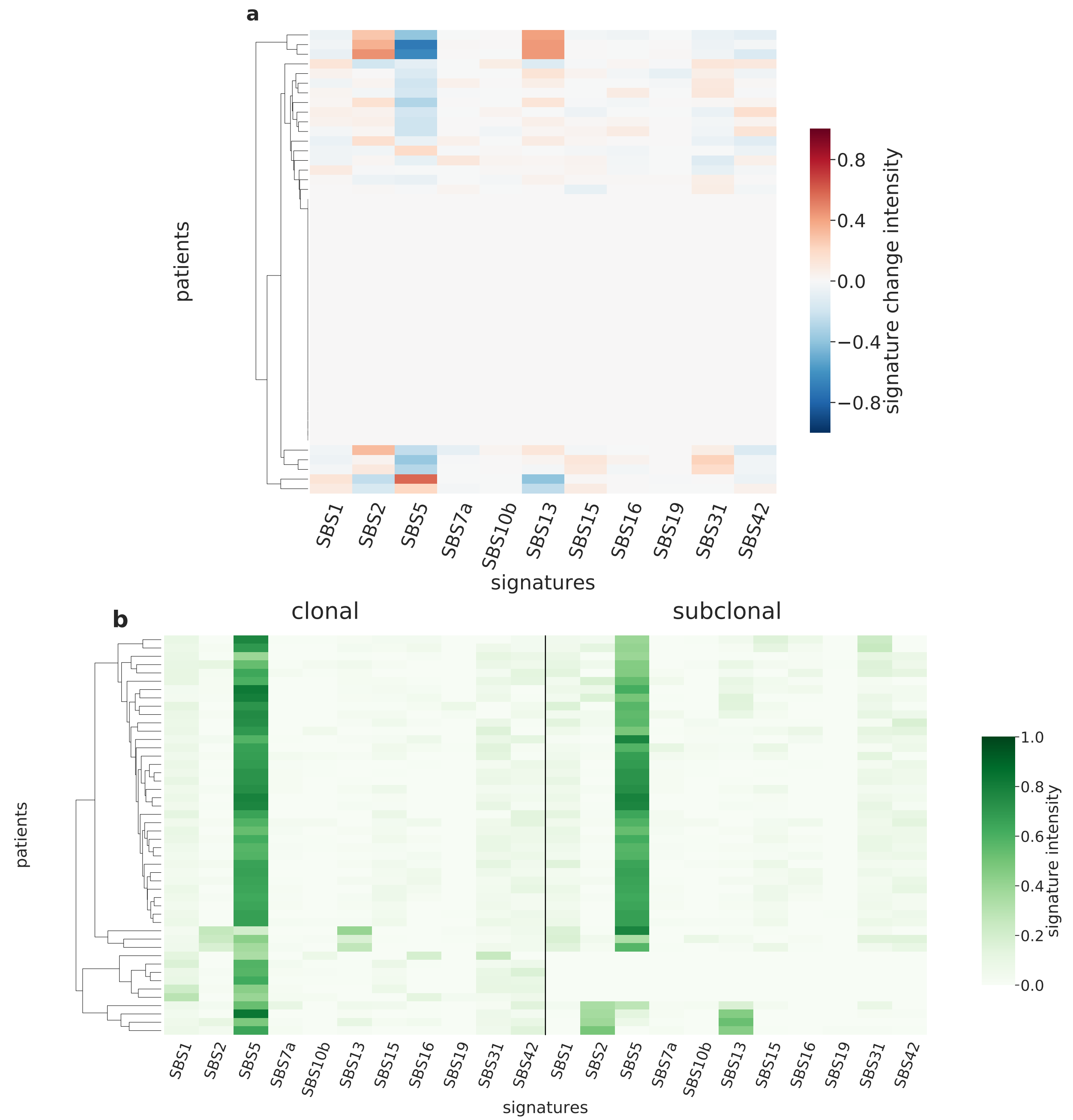

Supplementary Figure 101: **Overview of CloneSig results for the PCAWG sub-cohort Thy-AdenoCA (n=48).** Panel a: Stratification of patients depending on their pattern of signature change for Thy-AdenoCA patients (48 patients, including 22 with a significant signature change). The heatmap represents the difference between the signature activity in the largest subclone (in terms of number of mutations) and the clonal mutations (defined as belonging to the clone of highest CCF).Panel b: Stratification of patients depending on their complete pattern of signature exposure. The heatmap represents the signature activity in the largest subclone (in terms of number of mutations) and the clonal mutations (defined as belonging to the clone of highest CCF). Source data are provided as a Source Data file.

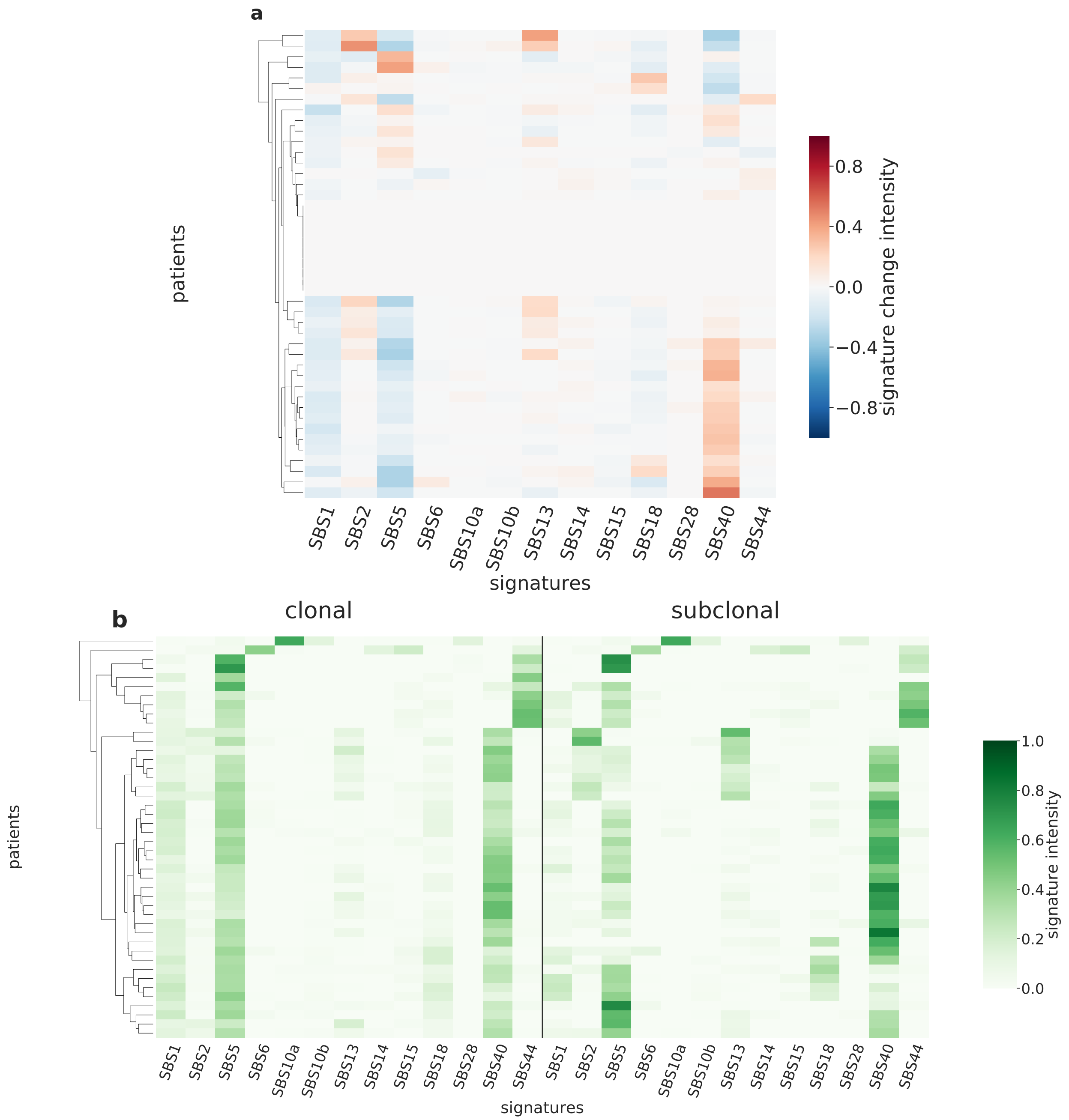

Supplementary Figure 102: **Overview of CloneSig results for the PCAWG sub-cohort Uterus-AdenoCA (n=44).** Panel a: Stratification of patients depending on their pattern of signature change for Uterus-AdenoCA patients (44 patients, including 35 with a significant signature change). The heatmap represents the difference between the signature activity in the largest subclone (in terms of number of mutations) and the clonal mutations (defined as belonging to the clone of highest CCF).Panel b: Stratification of patients depending on their complete pattern of signature exposure. The heatmap represents the signature activity in the largest subclone (in terms of number of mutations) and the clonal mutations (defined as belonging to the clone of highest CCF). Source data are provided as a Source Data file.

### Supplementary Figures

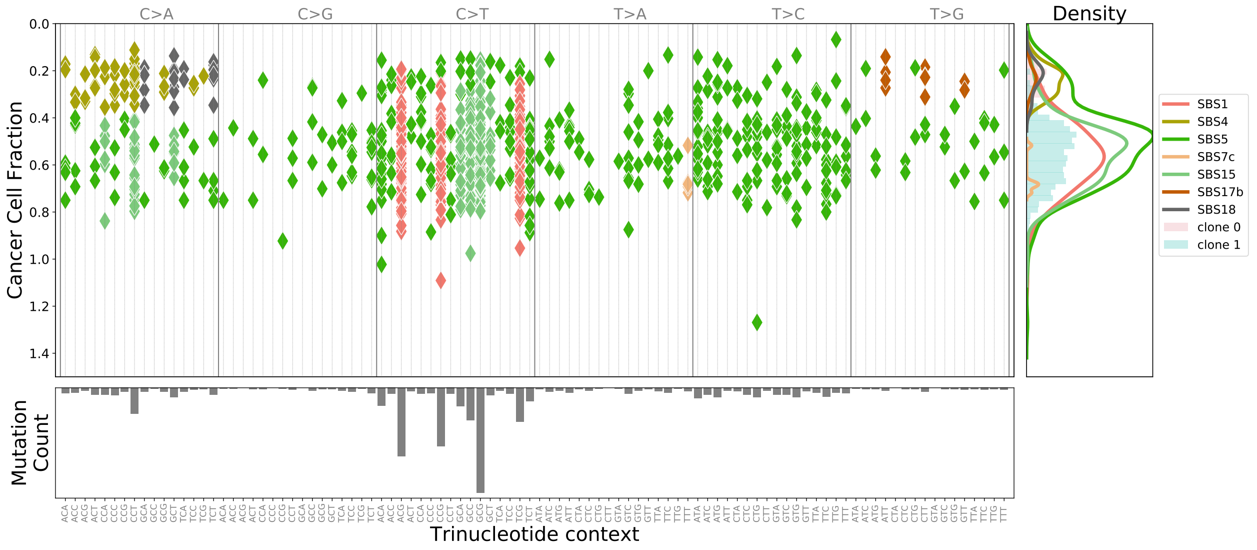

Supplementary Figure 103: **CloneSig analysis of 1002 SNVs obtained by WES of a SARC sample (patient TCGA-3B-A9HT)**

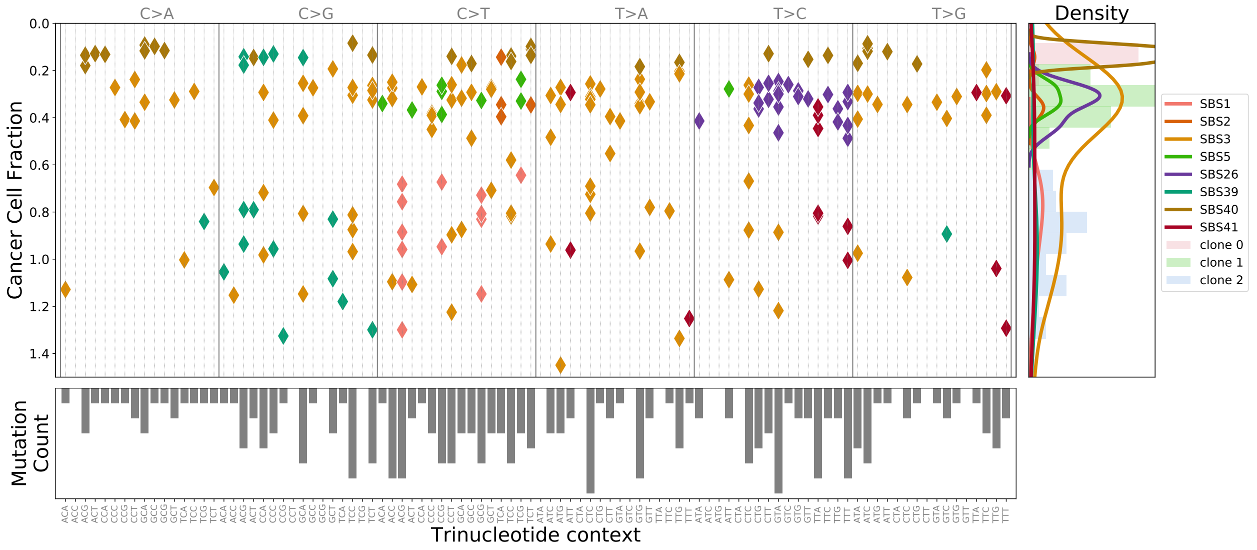

Supplementary Figure 104: **CloneSig analysis of 221 SNVs obtained by WES of a OV sample (patient TCGA-04-1341)**

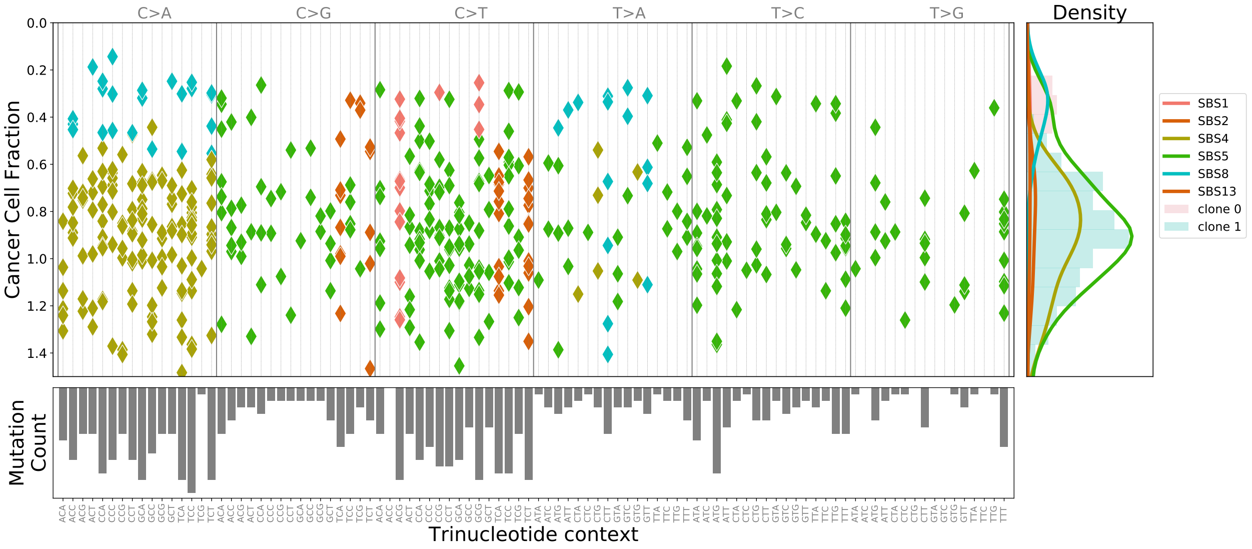

Supplementary Figure 105: **CloneSig analysis of 526 SNVs obtained by WES of a LUSC sample (patient TCGA-18-4721)**

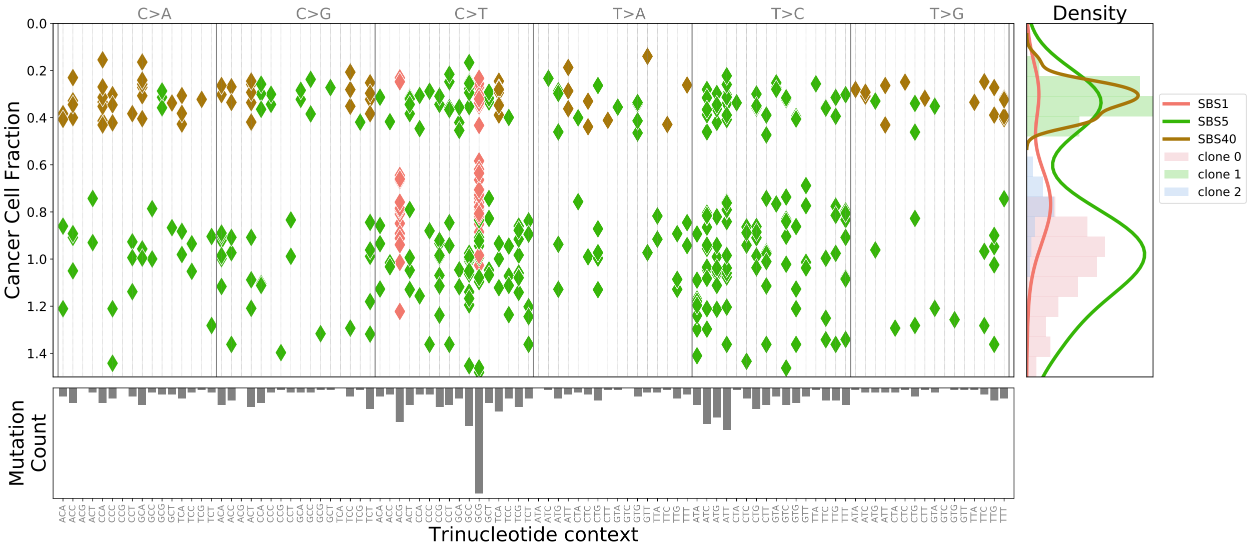

Supplementary Figure 106: **CloneSig analysis of 475 SNVs obtained by WES of a GBM sample (patient TCGA-19-2629)**

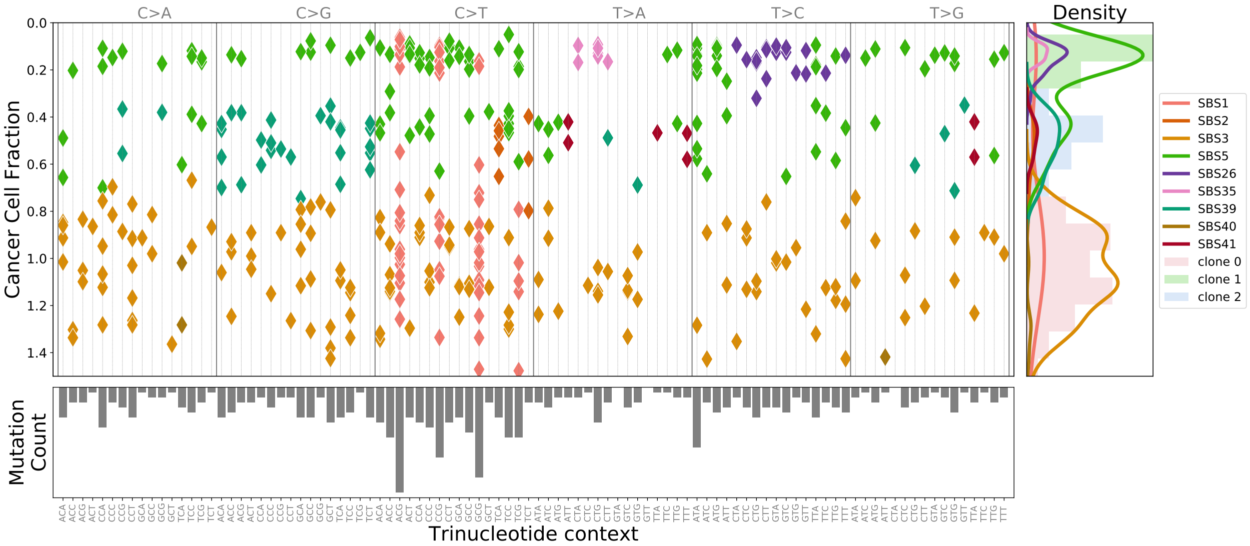

Supplementary Figure 107: **CloneSig analysis of 407 SNVs obtained by WES of a OV sample (patient TCGA-20-0990)**

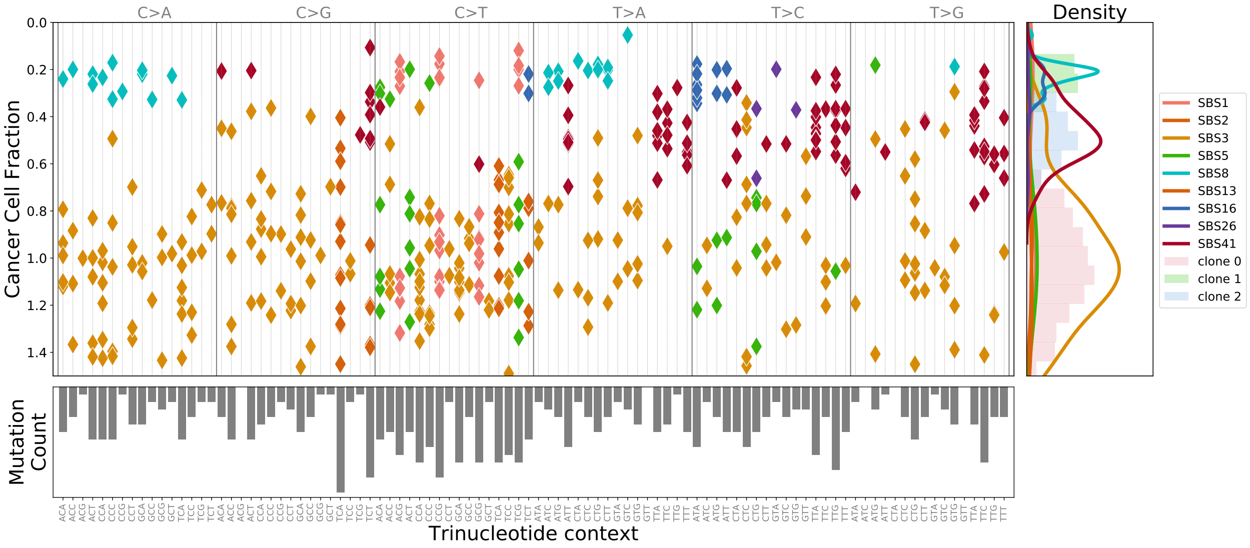

Supplementary Figure 108: **CloneSig analysis of 460 SNVs obtained by WES of a OV sample (patient TCGA-20-1685)**

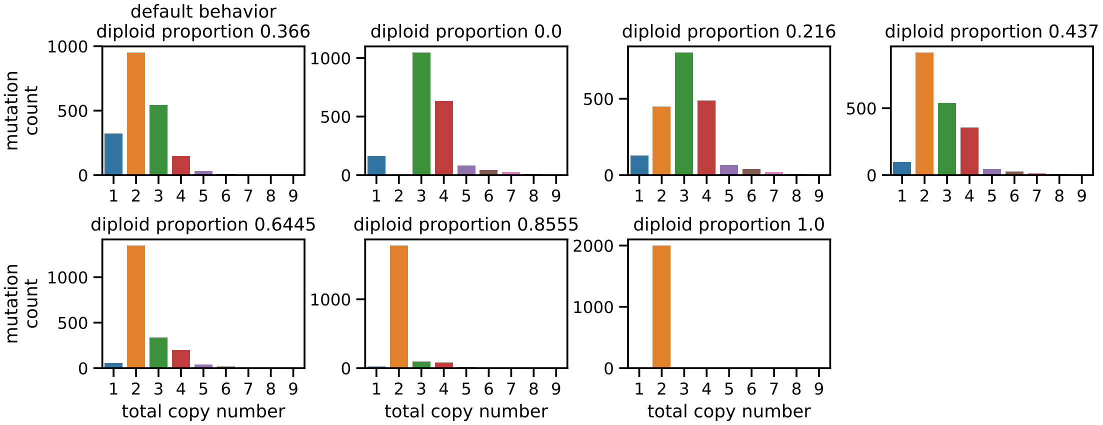

Supplementary Figure 109: **An example of empirical distribution of the total copy number for samples with 2000 mutations.** In the first panel, labeled "default behavior", the user does not specify the percentage of genome that is diploid, and the total copy number values are drawn as specified in the Methods section. On the other panels, the user specifies a desired percentage of genome that is diploid (0, 0.2, 0.4, 0.6, 0.8, 1) respectively for the cases shown. The distribution is slightly different from the default behavior.

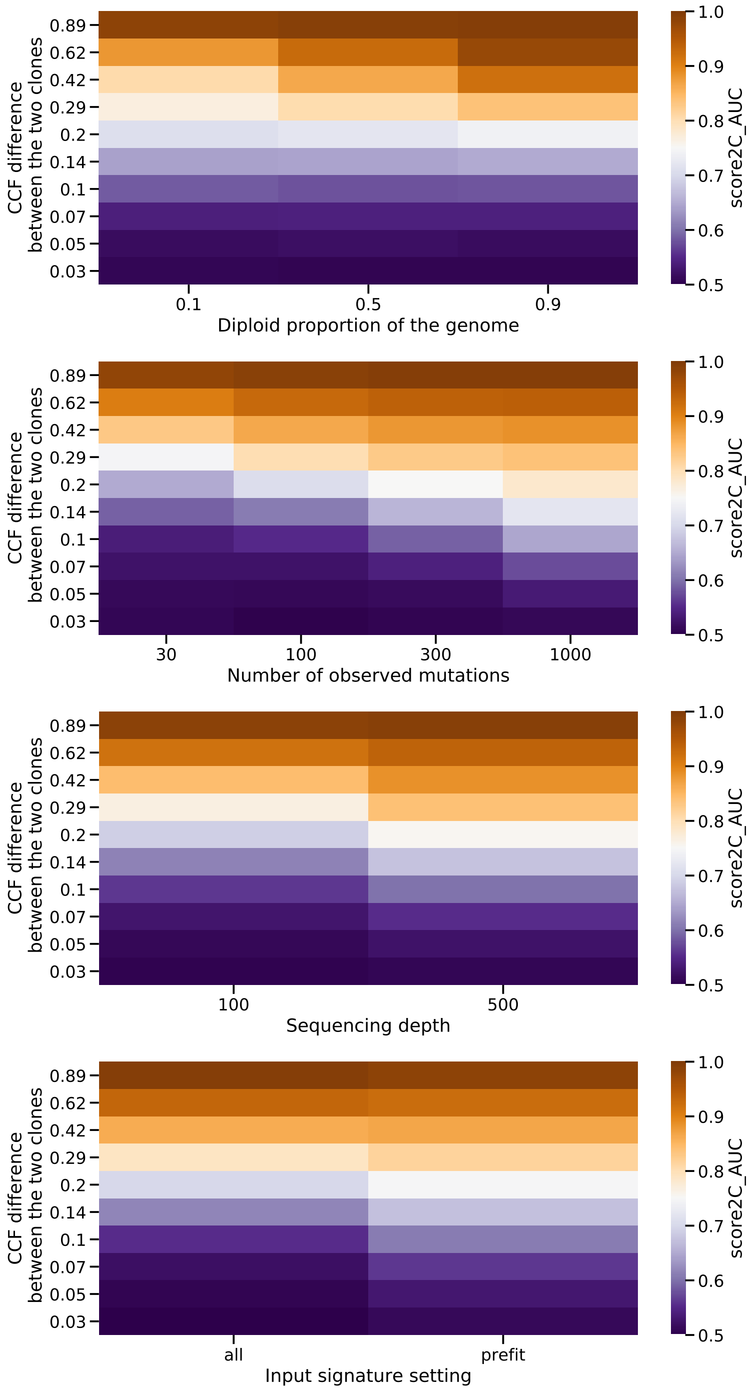

Supplementary Figure 110: **Accuracy of the number of clones estimated by CloneSig.** Those heatmaps illustrate CloneSig's ability to distinguish 2 clones depending on the CCF distance between the two clones, and other relevant variables: number of mutations, percentage of diploid genome, sequencing depth, and the mode of input signature selection. Source data are provided as a Source Data file.

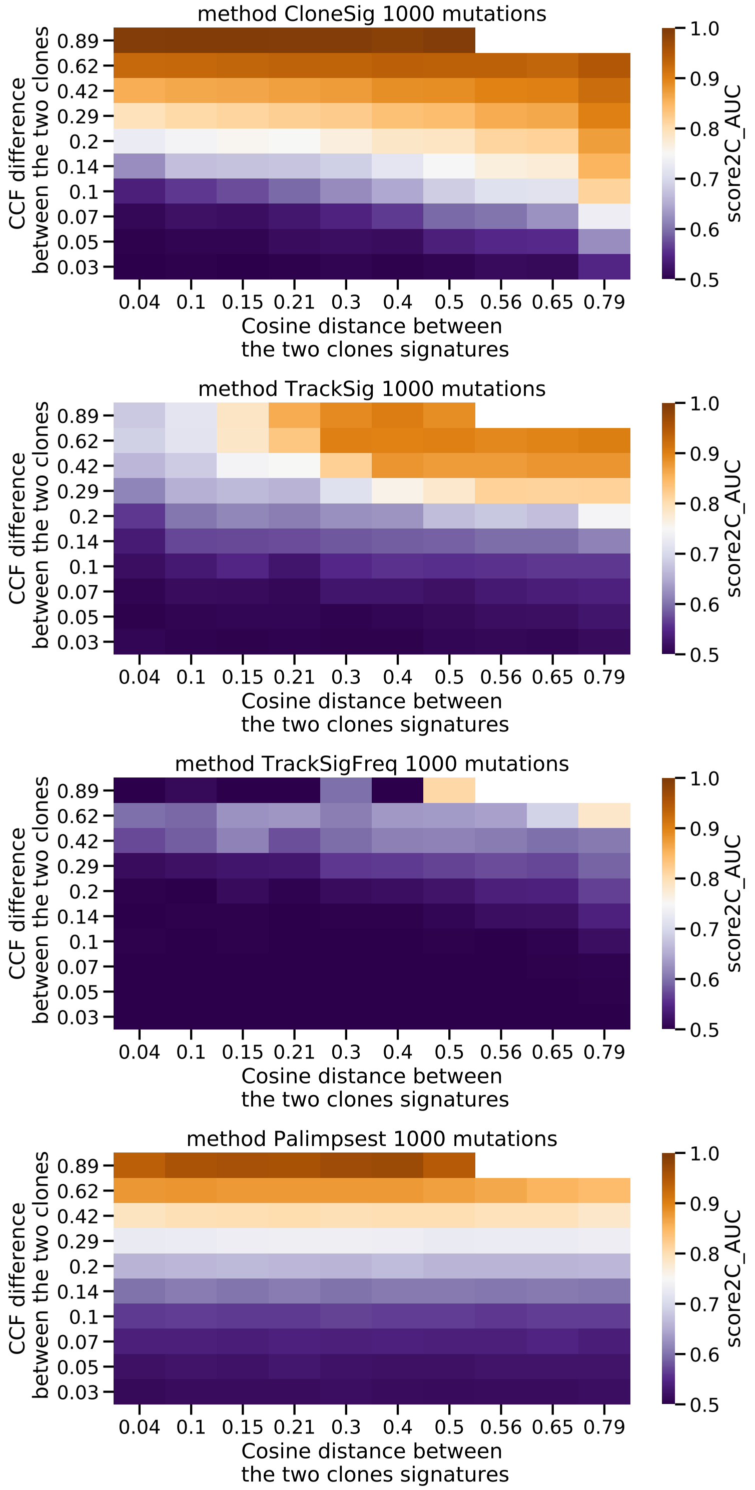

Supplementary Figure 111: **Evaluation of the power of separation of two clones with varying CCF distance and mutation profile distance**, for the different methods considered that account both for CCF mutations, and mutational context. Only simulations with 1000 observed signatures were considered, as TrackSig and TrackSigFreq hardly capture any signal with inferior numbers of signatures. Source data are provided as a Source Data file.

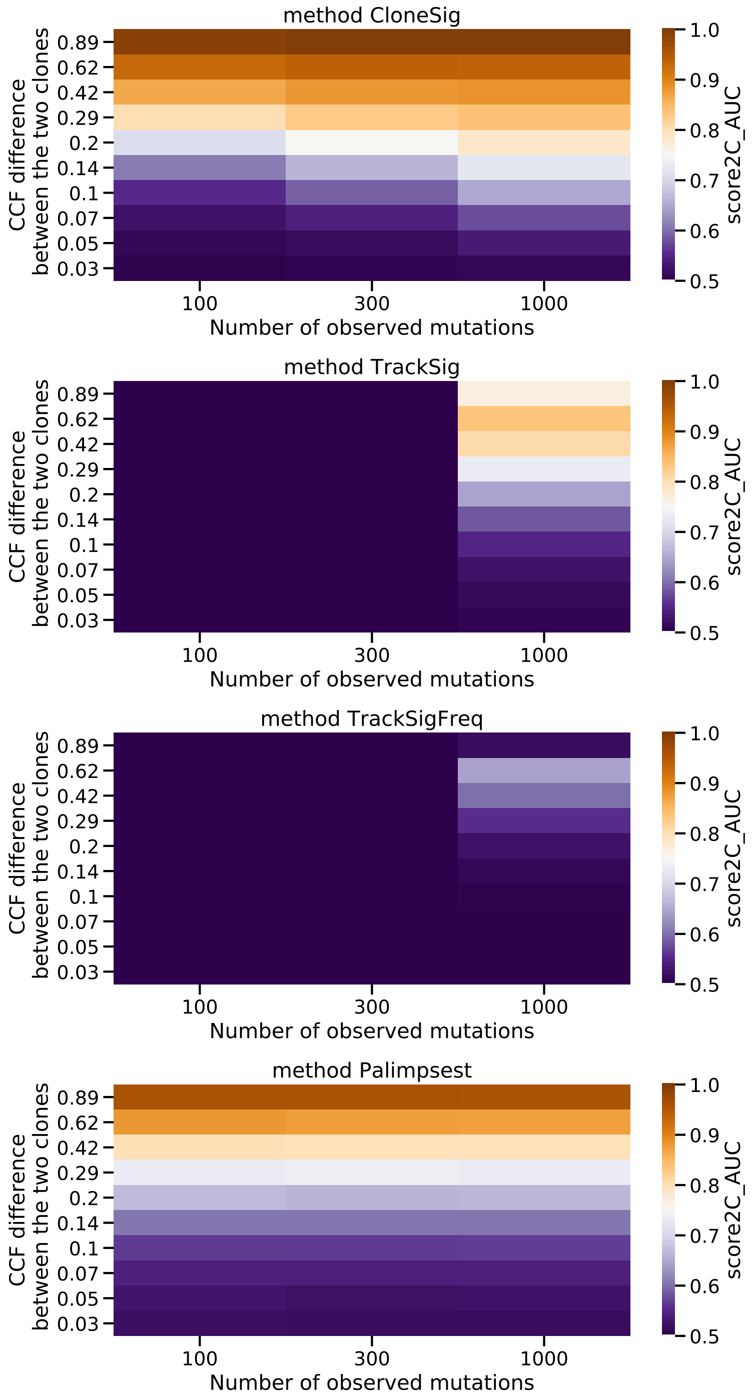

Supplementary Figure 112: **Influence of the number of observed mutations for each method's ability to distinguish clonal and subclonal mutations** for simulations with 2 clones with a varying CCF difference. Source data are provided as a Source Data file.

### Supplementary Tables

| Cancer type | Mean number of mutations (protected) | Mean number of mutations (public) | Standard deviation number of mutations (protected) | Standard deviation number of mutations (public) | Number of samples | Median followup (months) | Number of events |
|---|---|---|---|---|---|---|---|
| ACC | 324.34 | 110.58 | 467.43 | 240.88 | 77 | 39.22 | 27 |
| BLCA | 732.18 | 350.38 | 886.65 | 424.23 | 354 | 17.21 | 153 |
| BRCA | 472.91 | 121.88 | 981.68 | 372.65 | 931 | 27.00 | 123 |
| CESC | 921.07 | 366.51 | 2869.13 | 1323.24 | 275 | 21.42 | 67 |
| CHOL | 358.97 | 100.97 | 505.33 | 225.34 | 35 | 12.65 | 15 |
| COADREAD | 2085.67 | 621.14 | 6247.55 | 1708.77 | 458 | 21.42 | 93 |
| DLBC | 568.30 | 203.68 | 276.72 | 123.15 | 37 | 24.67 | 5 |
| ESCA | 707.39 | 247.19 | 560.34 | 251.02 | 180 | 13.02 | 75 |
| GBM | 790.05 | 245.66 | 2583.80 | 1191.70 | 327 | 11.27 | 246 |
| HNSC | 454.21 | 201.73 | 543.69 | 271.16 | 445 | 20.96 | 185 |
| KICH | 209.32 | 50.12 | 264.68 | 142.42 | 60 | 85.66 | 8 |
| KIRC | 330.32 | 73.08 | 338.69 | 52.35 | 271 | 36.33 | 64 |
| KIRP | 280.86 | 82.57 | 130.50 | 37.90 | 242 | 25.13 | 37 |
| LGG | 212.73 | 76.89 | 1462.32 | 770.54 | 455 | 20.04 | 115 |
| LIHC | 511.53 | 157.00 | 445.93 | 174.27 | 347 | 19.25 | 117 |
| LUAD | 892.89 | 381.00 | 985.35 | 393.91 | 433 | 22.01 | 146 |
| LUSC | 909.66 | 382.83 | 686.09 | 289.58 | 423 | 22.27 | 169 |
| MESO | 203.23 | 47.08 | 114.19 | 48.74 | 78 | NA | 0 |
| OV | 593.79 | 160.15 | 562.30 | 152.24 | 390 | 31.34 | 227 |
| PAAD | 560.23 | 203.05 | 3780.28 | 1828.93 | 150 | 15.14 | 83 |
| PCPG | 78.70 | 14.09 | 16.71 | 7.43 | 141 | NA | 0 |
| PRAD | 171.30 | 61.78 | 824.76 | 467.50 | 458 | 30.80 | 8 |
| SARC | 424.27 | 130.72 | 723.66 | 309.04 | 210 | 30.96 | 81 |
| SKCM | 1876.97 | 886.84 | 2621.06 | 1204.90 | 423 | 35.28 | 184 |
| STAD | 989.96 | 455.07 | 1822.53 | 909.56 | 418 | 14.01 | 163 |
| TGCT | 148.88 | 22.16 | 33.90 | 11.64 | 128 | 43.05 | 3 |
| THCA | 120.98 | 16.22 | 95.88 | 14.59 | 467 | 31.01 | 12 |
| THYM | 253.69 | 36.40 | 175.18 | 83.19 | 121 | 39.17 | 8 |
| UCEC | 4647.22 | 1791.01 | 12341.73 | 4832.10 | 487 | 30.12 | 80 |
| UCS | 680.43 | 198.00 | 1743.77 | 738.28 | 53 | NA | 0 |
| UVM | 88.54 | 24.60 | 96.34 | 62.57 | 80 | 27.52 | 11 |

Supplementary Table 3: **Characteristics of the TCGA cohort used in this study.**

| tcga_name | ACC | BLCA | BRCA | CESC | CHOL | COADREAD | DLBC | ESCA | GBM | HNSC | KICH | KIRC | KIRP | LGG | LIHC | LUAD | LUSC | MESO | OV | PAAD | PCPG | PRAD | SARC | SKCM | STAD | TGCT | THCA | THYM | UCEC | UCS | UVM |
|---|---|---|---|---|---|---|---|---|---|---|---|---|---|---|---|---|---|---|---|---|---|---|---|---|---|---|---|---|---|---|---|
| SBS1 | 1 | 1 | 1 | 1 | 1 | 1 | 1 | 1 | 1 | 1 | 1 | 1 | 1 | 1 | 1 | 1 | 1 | 1 | 1 | 1 | 1 | 1 | 1 | 1 | 1 | 1 | 1 | 1 | 1 | 1 | 1 |
| SBS2 | 1 | 1 | 1 | 1 | 1 | 1 | 1 | 1 | 0 | 1 | 1 | 1[18] | 1 | 1 | 0 | 1 | 1 | 1 | 1 | 1 | 1 | 1[19] | 1 | 1[19] | 1 | 1 | 1 | 1 | 1 | 1 | 1 |
| SBS3 | 1 | 1 | 1 | 0 | 0 | 0 | 1[19] | 1 | 0 | 1[20] | 0 | 1[18] | 0 | 0 | 1 | 1[19] | 0 | 0 | 1 | 1[19] | 1 | 1[19] | 1 | 0 | 1[21] | 1 | 0 | 0 | 0 | 1[19] | 1 |
| SBS4 | 0 | 0 | 0 | 0 | 0 | 0 | 0 | 0 | 0 | 1 | 0 | 0 | 0 | 0 | 1[12] | 1 | 1 | 0 | 0 | 0 | 0 | 0 | 1 | 0 | 0 | 0 | 0 | 1 | 0 | 0 | 0 |
| SBS5 | 1 | 1 | 1 | 1 | 1 | 1 | 1 | 1 | 1 | 1 | 1 | 1 | 1 | 1 | 1 | 1 | 1 | 1 | 1 | 1 | 1 | 1 | 1 | 1 | 1 | 1 | 1 | 1 | 1 | 1 | 1 |
| SBS6 | 0 | 0 | 1[22] | 1 | 1[23] | 1[24] | 1 | 0 | 0 | 0 | 0 | 0 | 0 | 0 | 1 | 0 | 0 | 0 | 0 | 1[19] | 0 | 0 | 0 | 0 | 0 | 0 | 0 | 0 | 1[25] | 1[19] | 1[26] |
| SBS7a | 0 | 0 | 0 | 0 | 0 | 0 | 1 | 0 | 0 | 1[19] | 0 | 0 | 0 | 0 | 0 | 0 | 0 | 0 | 0 | 0 | 1 | 0 | 1 | 1 | 0 | 1 | 1 | 1 | 0 | 0 | 1 |
| SBS7b | 0 | 0 | 0 | 0 | 0 | 0 | 0 | 0 | 0 | 1[19] | 0 | 0 | 0 | 0 | 0 | 0 | 0 | 0 | 0 | 0 | 1 | 0 | 1 | 1 | 0 | 0 | 0 | 0 | 0 | 0 | 0 |
| SBS7c | 0 | 0 | 0 | 0 | 0 | 0 | 1 | 0 | 0 | 0 | 0 | 0 | 0 | 0 | 0 | 0 | 0 | 0 | 0 | 0 | 0 | 0 | 1 | 1 | 0 | 1 | 0 | 0 | 0 | 0 | 0 |
| SBS7d | 0 | 0 | 0 | 0 | 0 | 0 | 0 | 0 | 0 | 1[19] | 0 | 0 | 0 | 0 | 0 | 0 | 0 | 0 | 0 | 1 | 1 | 0 | 1 | 1 | 0 | 0 | 0 | 1 | 0 | 0 | 1 |
| SBS8 | 0 | 1[19] | 1[22] | 0 | 1[23] | 0 | 0 | 0 | 0 | 0 | 1 | 0 | 0 | 0 | 0 | 0 | 1[19] | 0 | 1[19] | 1[19] | 0 | 1[27] | 1 | 0 | 0 | 0 | 0 | 1 | 0 | 0 | 0 |
| SBS9 | 0 | 0 | 1[19] | 0 | 0 | 0 | 1[28] | 0 | 0 | 0 | 0 | 1[18] | 0 | 0 | 1 | 1[19] | 0 | 0 | 0 | 0 | 0 | 1[27] | 0 | 0 | 0 | 0 | 0 | 0 | 0 | 0 | 0 |
| SBS10a | 0 | 0 | 1 | 0 | 0 | 1[19] | 0 | 0 | 0 | 0 | 0 | 0 | 0 | 0 | 0 | 0 | 0 | 0 | 0 | 1 | 0 | 0 | 0 | 0 | 0 | 1 | 0 | 1 | 1 | 1[19] | 0 |
| SBS10b | 1 | 0 | 1 | 1 | 0 | 1[19] | 0 | 0 | 0 | 0 | 1 | 0 | 0 | 1 | 0 | 0 | 0 | 1 | 0 | 1 | 1 | 1 | 0 | 0 | 0 | 1 | 1 | 1 | 1 | 1 | 0 |
| SBS11 | 0 | 0 | 0 | 0 | 0 | 0 | 0 | 0 | 1[19] | 0 | 0 | 0 | 0 | 1 | 0 | 0 | 0 | 0 | 0 | 0 | 0 | 0 | 0 | 0 | 0 | 0 | 0 | 0 | 0 | 0 | 0 |
| SBS12 | 0 | 0 | 0 | 0 | 0 | 0 | 0 | 0 | 0 | 0 | 1 | 0 | 0 | 0 | 1[12] | 0 | 0 | 0 | 0 | 0 | 0 | 0 | 0 | 0 | 0 | 0 | 0 | 0 | 0 | 0 | 1[26] |
| SBS13 | 1 | 1 | 1 | 1 | 1 | 1 | 1 | 1 | 0 | 1 | 1[19] | 1[18] | 1 | 1 | 0 | 1 | 1 | 1 | 1 | 1 | 1 | 1[19] | 1 | 1[19] | 1 | 1 | 1 | 0 | 1 | 1 | 1 |
| SBS14 | 0 | 0 | 0 | 0 | 0 | 0 | 0 | 0 | 0 | 0 | 0 | 0 | 0 | 0 | 1[19] | 0 | 0 | 0 | 0 | 0 | 0 | 0 | 0 | 0 | 0 | 0 | 0 | 0 | 1 | 1[19] | 0 |
| SBS15 | 0 | 0 | 0 | 0 | 0 | 1 | 0 | 0 | 0 | 0 | 1 | 1 | 0 | 1 | 0 | 0 | 1 | 1 | 0 | 1 | 1 | 1 | 1 | 0 | 1 | 1 | 1 | 1 | 1 | 1[19] | 1 |
| SBS16 | 0 | 0 | 0 | 0 | 1[23] | 0 | 0 | 1 | 0 | 1[19] | 0 | 0 | 0 | 0 | 1 | 0 | 0 | 0 | 1[29] | 0 | 1 | 1[27] | 0 | 0 | 0 | 0 | 1 | 1 | 0 | 0 | 1[26] |
| SBS17a | 0 | 0 | 1[22] | 0 | 1[23] | 1 | 1 | 1 | 0 | 1[19] | 1 | 1[18] | 0 | 0 | 1[12] | 1[19] | 0 | 0 | 0 | 1[19] | 1 | 0 | 1[19] | 1[19] | 1 | 1 | 0 | 1 | 0 | 0 | 0 |
| SBS17b | 0 | 0 | 1[22] | 0 | 1[23] | 1 | 1 | 1 | 0 | 1[19] | 1[19] | 1[18] | 0 | 0 | 1[12] | 1[19] | 0 | 0 | 0 | 1[19] | 0 | 0 | 1[19] | 1[19] | 1 | 0 | 0 | 0 | 0 | 0 | 0 |
| SBS18 | 0 | 0 | 1[19] | 0 | 0 | 1[19] | 1[28] | 1 | 0 | 1[19] | 0 | 0 | 0 | 0 | 1[19] | 1[19] | 0 | 0 | 1[19] | 1[19] | 0 | 1[19] | 1 | 0 | 1[19] | 0 | 0 | 1 | 1[19] | 0 | 0 |
| SBS19 | 0 | 0 | 0 | 0 | 0 | 0 | 0 | 0 | 0 | 0 | 0 | 0 | 0 | 0 | 1[19] | 0 | 0 | 1 | 0 | 0 | 0 | 0 | 0 | 0 | 0 | 1 | 1 | 0 | 0 | 0 | 0 |
| SBS20 | 0 | 0 | 1[22] | 0 | 1[23] | 1 | 0 | 0 | 0 | 0 | 0 | 0 | 0 | 0 | 0 | 0 | 0 | 0 | 0 | 1[19] | 0 | 0 | 0 | 0 | 1 | 0 | 0 | 0 | 0 | 0 | 0 |
| SBS21 | 0 | 0 | 0 | 0 | 0 | 0 | 0 | 0 | 0 | 0 | 0 | 0 | 0 | 0 | 0 | 0 | 0 | 0 | 0 | 0 | 0 | 0 | 0 | 0 | 1[19] | 0 | 0 | 0 | 0 | 0 | 0 |
| SBS22 | 0 | 0 | 0 | 0 | 1[23] | 0 | 0 | 0 | 0 | 0 | 0 | 1[19] | 0 | 0 | 1 | 0 | 0 | 0 | 0 | 0 | 0 | 0 | 0 | 0 | 0 | 0 | 0 | 0 | 0 | 0 | 1 |
| SBS23 | 0 | 0 | 0 | 0 | 0 | 0 | 0 | 0 | 0 | 0 | 0 | 0 | 0 | 0 | 1[12] | 0 | 0 | 0 | 0 | 0 | 0 | 0 | 0 | 0 | 0 | 0 | 0 | 0 | 0 | 0 | 0 |
| SBS24 | 0 | 0 | 0 | 0 | 0 | 0 | 0 | 0 | 0 | 0 | 0 | 0 | 0 | 0 | 1 | 0 | 0 | 0 | 0 | 0 | 0 | 0 | 0 | 0 | 0 | 1 | 0 | 1 | 0 | 0 | 0 |
| SBS25 | 0 | 0 | 0 | 0 | 0 | 0 | 0 | 0 | 0 | 0 | 0 | 0 | 1 | 0 | 0 | 0 | 0 | 0 | 0 | 0 | 0 | 0 | 0 | 0 | 0 | 0 | 0 | 0 | 0 | 0 | 0 |
| SBS26 | 0 | 0 | 1[22] | 0 | 0 | 1[24] | 0 | 0 | 0 | 0 | 0 | 0 | 0 | 0 | 1[19] | 0 | 0 | 0 | 1[19] | 1[19] | 0 | 0 | 0 | 0 | 1[19] | 0 | 0 | 0 | 0 | 1[19] | 0 |
| SBS27 | 0 | 0 | 0 | 0 | 0 | 0 | 0 | 0 | 0 | 0 | 0 | 0 | 0 | 0 | 0 | 0 | 0 | 0 | 0 | 0 | 0 | 0 | 0 | 0 | 0 | 0 | 0 | 0 | 0 | 0 | 0 |
| SBS28 | 0 | 0 | 0 | 0 | 0 | 1[19] | 0 | 0 | 0 | 0 | 0 | 0 | 0 | 0 | 1[19] | 1[19] | 0 | 0 | 0 | 1[19] | 0 | 0 | 0 | 0 | 1[19] | 0 | 0 | 0 | 1 | 1[19] | 0 |
| SBS29 | 0 | 1[19] | 1 | 0 | 0 | 0 | 0 | 0 | 0 | 0 | 1[19] | 1[19] | 0 | 0 | 1[19] | 0 | 0 | 0 | 0 | 0 | 0 | 0 | 0 | 0 | 0 | 1 | 0 | 0 | 0 | 0 | 0 |
| SBS30 | 0 | 0 | 1 | 0 | 0 | 1 | 0 | 0 | 1 | 0 | 0 | 0 | 0 | 0 | 1 | 0 | 0 | 1 | 0 | 1[19] | 0 | 0 | 0 | 0 | 0 | 0 | 0 | 0 | 0 | 0 | 0 |
| SBS31 | 0 | 0 | 0 | 0 | 0 | 0 | 0 | 0 | 0 | 0 | 0 | 0 | 0 | 0 | 1[19] | 0 | 0 | 0 | 0 | 0 | 0 | 0 | 0 | 0 | 0 | 1 | 1 | 0 | 0 | 0 | 0 |
| SBS32 | 0 | 0 | 0 | 0 | 0 | 0 | 0 | 0 | 0 | 0 | 0 | 0 | 0 | 0 | 0 | 0 | 0 | 0 | 0 | 0 | 0 | 0 | 0 | 0 | 0 | 0 | 0 | 0 | 0 | 0 | 1 |
| SBS33 | 0 | 0 | 0 | 0 | 0 | 0 | 0 | 0 | 0 | 1[19] | 0 | 0 | 0 | 0 | 0 | 0 | 0 | 0 | 0 | 0 | 0 | 1[19] | 0 | 0 | 0 | 0 | 0 | 0 | 0 | 0 | 0 |
| SBS34 | 0 | 0 | 0 | 0 | 0 | 0 | 1[19] | 0 | 0 | 0 | 0 | 0 | 0 | 0 | 0 | 0 | 0 | 0 | 0 | 0 | 0 | 0 | 0 | 0 | 0 | 0 | 0 | 0 | 0 | 0 | 0 |
| SBS35 | 0 | 0 | 0 | 0 | 0 | 0 | 0 | 0 | 0 | 0 | 0 | 0 | 0 | 0 | 1[19] | 0 | 0 | 0 | 1[19] | 0 | 0 | 0 | 0 | 0 | 0 | 1 | 0 | 0 | 0 | 0 | 0 |
| SBS36 | 1 | 0 | 0 | 0 | 0 | 0 | 1[19] | 0 | 0 | 0 | 0 | 0 | 0 | 1 | 0 | 0 | 0 | 0 | 0 | 0 | 0 | 0 | 0 | 0 | 0 | 0 | 0 | 0 | 0 | 0 | 0 |
| SBS37 | 0 | 0 | 1[19] | 0 | 0 | 1[19] | 1[19] | 0 | 0 | 0 | 0 | 0 | 0 | 1 | 0 | 0 | 0 | 0 | 0 | 0 | 0 | 1[19] | 0 | 0 | 0 | 0 | 0 | 0 | 0 | 0 | 0 |
| SBS38 | 0 | 0 | 0 | 0 | 0 | 0 | 0 | 0 | 0 | 0 | 0 | 0 | 0 | 0 | 0 | 0 | 0 | 1 | 0 | 0 | 0 | 0 | 0 | 1 | 0 | 0 | 0 | 1 | 0 | 0 | 0 |
| SBS39 | 0 | 0 | 0 | 0 | 0 | 0 | 0 | 0 | 0 | 0 | 0 | 0 | 0 | 1 | 0 | 0 | 0 | 0 | 1[19] | 0 | 0 | 0 | 0 | 0 | 0 | 1 | 0 | 0 | 0 | 0 | 0 |
| SBS40 | 0 | 1[19] | 1[19] | 0 | 1 | 1 | 1 | 1[19] | 1 | 1[19] | 1 | 1 | 1 | 0 | 1 | 1 | 0 | 1 | 1 | 1[19] | 0 | 1 | 1 | 1[19] | 1 | 1 | 0 | 1 | 1 | 1[19] | 0 |
| SBS41 | 0 | 0 | 1[19] | 0 | 0 | 0 | 0 | 0 | 0 | 0 | 0 | 1[19] | 0 | 0 | 0 | 0 | 0 | 0 | 1[19] | 0 | 0 | 1[19] | 0 | 0 | 1[19] | 0 | 0 | 0 | 0 | 0 | 0 |
| SBS42 | 0 | 0 | 0 | 0 | 1 | 0 | 1 | 0 | 0 | 0 | 0 | 0 | 0 | 0 | 0 | 0 | 0 | 0 | 0 | 0 | 1 | 0 | 0 | 0 | 0 | 1 | 1 | 1 | 0 | 0 | 1 |
| SBS43 | 0 | 0 | 0 | 0 | 0 | 0 | 0 | 0 | 0 | 0 | 0 | 0 | 0 | 0 | 0 | 0 | 0 | 0 | 0 | 0 | 0 | 0 | 0 | 0 | 0 | 0 | 0 | 0 | 0 | 0 | 0 |
| SBS44 | 0 | 0 | 0 | 0 | 1 | 1 | 0 | 0 | 0 | 0 | 0 | 0 | 0 | 0 | 0 | 0 | 0 | 0 | 0 | 0 | 0 | 0 | 0 | 0 | 1[19] | 1 | 0 | 1 | 1 | 1[19] | 0 |
| SBS45 | 0 | 1 | 0 | 0 | 0 | 0 | 0 | 0 | 1 | 1 | 0 | 1 | 1 | 1 | 0 | 1 | 1 | 0 | 0 | 0 | 0 | 1 | 0 | 1 | 0 | 0 | 1 | 0 | 0 | 0 | 0 |
| SBS46 | 0 | 0 | 0 | 0 | 0 | 0 | 0 | 0 | 0 | 0 | 1 | 0 | 0 | 0 | 0 | 0 | 0 | 0 | 0 | 0 | 0 | 0 | 0 | 0 | 0 | 0 | 0 | 0 | 0 | 0 | 0 |
| SBS47 | 0 | 0 | 0 | 0 | 0 | 0 | 0 | 0 | 0 | 0 | 0 | 0 | 0 | 0 | 0 | 0 | 0 | 0 | 0 | 0 | 0 | 0 | 0 | 0 | 0 | 0 | 0 | 1 | 0 | 0 | 0 |
| SBS48 | 0 | 0 | 0 | 0 | 0 | 0 | 0 | 0 | 0 | 0 | 0 | 0 | 0 | 0 | 0 | 0 | 0 | 0 | 0 | 0 | 0 | 0 | 0 | 0 | 0 | 0 | 0 | 0 | 0 | 0 | 0 |
| SBS49 | 0 | 0 | 0 | 0 | 0 | 0 | 0 | 0 | 0 | 0 | 0 | 0 | 1 | 0 | 0 | 0 | 0 | 0 | 0 | 0 | 0 | 0 | 0 | 1 | 0 | 1 | 1 | 1 | 0 | 0 | 0 |
| SBS50 | 0 | 0 | 0 | 0 | 0 | 0 | 0 | 0 | 0 | 0 | 0 | 0 | 0 | 0 | 0 | 0 | 0 | 0 | 0 | 0 | 0 | 0 | 0 | 0 | 0 | 0 | 0 | 0 | 0 | 0 | 0 |
| SBS51 | 0 | 0 | 0 | 0 | 0 | 0 | 0 | 0 | 0 | 0 | 0 | 0 | 0 | 0 | 0 | 0 | 0 | 0 | 0 | 0 | 0 | 0 | 0 | 0 | 0 | 0 | 0 | 0 | 0 | 0 | 0 |
| SBS52 | 0 | 1 | 0 | 0 | 0 | 0 | 0 | 0 | 0 | 0 | 0 | 1 | 0 | 0 | 0 | 0 | 0 | 0 | 0 | 0 | 0 | 0 | 0 | 0 | 0 | 1 | 0 | 1 | 0 | 0 | 0 |
| SBS53 | 0 | 0 | 0 | 0 | 0 | 0 | 0 | 0 | 0 | 0 | 0 | 0 | 0 | 0 | 0 | 0 | 0 | 0 | 0 | 0 | 0 | 0 | 0 | 0 | 0 | 0 | 0 | 0 | 0 | 0 | 0 |
| SBS54 | 0 | 0 | 0 | 0 | 0 | 0 | 0 | 0 | 0 | 0 | 0 | 0 | 0 | 0 | 0 | 0 | 0 | 0 | 0 | 0 | 1 | 0 | 0 | 0 | 1 | 0 | 0 | 0 | 0 | 0 | 0 |
| SBS55 | 0 | 0 | 0 | 0 | 0 | 0 | 0 | 0 | 0 | 0 | 0 | 0 | 0 | 0 | 0 | 0 | 0 | 0 | 0 | 0 | 0 | 0 | 0 | 0 | 0 | 0 | 0 | 0 | 0 | 0 | 0 |
| SBS56 | 0 | 0 | 0 | 0 | 0 | 0 | 0 | 0 | 0 | 0 | 0 | 0 | 0 | 0 | 0 | 0 | 0 | 0 | 0 | 0 | 0 | 0 | 0 | 0 | 0 | 0 | 0 | 0 | 0 | 0 | 0 |
| SBS57 | 0 | 0 | 0 | 0 | 0 | 0 | 0 | 0 | 0 | 0 | 0 | 0 | 0 | 0 | 0 | 0 | 0 | 0 | 0 | 0 | 0 | 0 | 0 | 0 | 0 | 0 | 0 | 0 | 0 | 0 | 1 |
| SBS58 | 0 | 0 | 0 | 0 | 0 | 0 | 0 | 0 | 0 | 0 | 0 | 0 | 0 | 0 | 0 | 0 | 0 | 0 | 0 | 0 | 1 | 0 | 1 | 0 | 0 | 1 | 0 | 0 | 0 | 0 | 0 |
| SBS59 | 0 | 0 | 0 | 0 | 0 | 0 | 0 | 0 | 0 | 0 | 0 | 0 | 0 | 0 | 0 | 0 | 0 | 0 | 0 | 0 | 0 | 0 | 1 | 0 | 0 | 1 | 0 | 0 | 0 | 0 | 1 |
| SBS60 | 0 | 0 | 0 | 0 | 0 | 0 | 0 | 0 | 0 | 0 | 0 | 0 | 0 | 0 | 0 | 0 | 0 | 0 | 0 | 0 | 0 | 0 | 0 | 0 | 0 | 0 | 0 | 0 | 0 | 0 | 0 |

Supplementary Table 4: **Table of presence of signatures in the different cancer types.** A 1 indicates the presence of the signature, and a 0 an absence. The background color indicates whether the presence of the signature comes from SigProfiler detection in the TCGA (lavender) or from the literature (green), in which case, the matching paper is referenced in the table cell. Signatures suspected to be artefacts were removed and are indicated in dark pink.
